# Microdroplet screening rapidly profiles a biocatalyst to enable its AI-assisted engineering

**DOI:** 10.1101/2024.04.08.588565

**Authors:** Maximilian Gantz, Simon V. Mathis, Friederike E. H. Nintzel, Matthew Penner, Paul J. Zurek, Tanja Knaus, Vasilis Tseliou, Elie Patel, Daniel Boros, Friedrich-Maximilian Weberling, Matthew R. A. Kenneth, Oskar J. Klein, Elliot J. Medcalf, Jacob Moss, Michael Herger, Tomasz S. Kaminski, Francesco G. Mutti, Pietro Lio, Florian Hollfelder

## Abstract

Engineering enzymes for increased efficiency is key to enabling sustainable, ‘green’ biocatalytic production processes in the chemical and pharmaceutical industries. This challenge can be tackled from two angles: by directed evolution, based on labour-intensive experimental testing of enzyme variant libraries, or by computational methods, where data-dependent algorithms relating sequence and function are used to predict biocatalyst improvements. Here, we combine both approaches into a two-week, low-cost workflow, in which ultra-high throughput screening of a library of imine reductases (IREDs) in microfluidic devices provides not only selected ‘hits’, but also long-read sequence data linked to fitness scores of >17 thousand enzyme variants. We demonstrate the engineering of an IRED for chiral amine synthesis by mapping its local fitness landscape in one go, ready to be used for interpretation and extrapolation by protein engineers with the help of machine learning (ML). We calculate position-dependent mutability and combinability scores of mutations and comprehensively illuminate a complex interplay of mutations driven by synergistic, often positively epistatic effects. When interpreted by easy-to-use regression and tree-based ML algorithms designed for random whole-gene mutagenesis data, 3-fold improved ‘hits’ initially obtained from experimental screening are extrapolated further to give another order of magnitude improvement (23-fold in kcat) after testing only a handful of designed mutants. Predictions succeed in >80% of cases. The catalytic features discovered in one IRED are shown to be portable and confer activity on IREDs with ∼50% homology. Our campaigns yield biocatalytically efficient IREDs and are paradigmatic for future enzyme engineering efforts that rely on large sequence-function maps, profiling how a biocatalyst responds to mutation. In the age of predictive biology, these maps will chart the way to improved function by exploiting the synergy of rapid experimental screening combined with ML evaluation and extrapolation.

## Introduction

Protein engineering has advanced enormously over the last decades, successfully employing a range of experimental^1^ and computational approaches^2–8^. Purely computational approaches reduce the laborious iterative cycles of mutagenesis and experimental testing in directed evolution, reliably yielding stabilised proteins, even new folds, as well as to binders with improved or altered functions^9^. The engineering of enzymes, however, remains a challenge because - in addition to stability, folding and binding - the sub-Ångstrom precision of the catalytic machinery (including possible conformational changes as bonds are made and broken) imposes multiple biophysical and chemical demands on an engineered enzyme. The combinatorial space of possible amino acid variations during an engineering task is vast^1^, while relatively few mutations satisfy the complex requirements for catalysis. Beyond identifying single beneficial mutations, combining them in constructive evolutionary trajectories can be hampered by their non-additivity, i.e. negative epistatic interactions between amino acids^10,11^ that pose a major practical challenge in enzyme engineering.

At the same time, the industrial transition to a green bioeconomy urgently requires new enzymes for biocatalysis, to create innovative and sustainable routes to valuable chemicals: atom efficient, with chiral control and with a more benign environmental footprint than current processes based on chemical catalysis^12–15^. Biocatalysts can be sourced from naturally occurring enzymes with promiscuous activities^16^ that accept non-natural substrates. However, these activities usually require further experimental engineering to meet industrial demands for high activity and stability^12^. The success of directed evolution depends heavily on the throughput of experimental screens: the more randomly generated mutants can be experimentally tested, the more likely the identification of an improved catalyst becomes. Consequently, this process is labour and resource intensive, and despite improvements in screening throughput, the outcome of directed evolution remains unpredictable, making it a hit-and-miss approach^17^. Nevertheless, biocatalysts that substantially improve the economic viability and process sustainability of chemical synthesis have been identified in this way. This has been exemplified in the production of high-value intermediates for the pharmaceutical industry, such as chiral amines, by using carbon-nitrogen bond forming enzymes such as transaminases^18^ or imine reductases (IREDs)^19,20^.

Here we showcase a fundamentally different protein engineering approach propelled by the synergy of experimental and computational methods: rapid sequence-function mapping through long-read deep mutational scanning (lrDMS) based on ultrahigh throughput screening, followed by machine learning (ML) and we demonstrate its utility for the engineering of biocatalytic IREDs. First, we perform screening in water-in-oil emulsion microdroplets (formed in microfluidic devices)^21^, in which scaling-down to picolitre volumes minimizes costs (∼10^−5^ cents per assay of a library member^22^) and accelerates the screening process (requiring only two weeks for the entire workflow shown in **Figure 1A**)^21,22^. This gives us access to individual improved variants (as in traditional directed evolution), but more importantly we harvest vast amounts of quantitative sequence-function data to profile the enzyme and gain insights into the importance of individual residues for catalysis, specifically their mutability and combinability. These experiments are designed to reveal each enzyme’s idiosyncratic biophysical and mechanistic persona in response to mutagenesis. Basing a strategy on a comprehensive mutagenic profile, which is unavailable from low-throughput approaches, changes the engineering paradigm from focusing on the effects of individual residues to performing a network analysis that covers the entire protein structure, enabling the protein engineer to navigate complex intra-gene epistasis. The assay-labelled data generated here is selected for catalysis rather than structure^23,24^ and harbours information that purely *in silico* strategies for enzyme engineering with AI^25,26^ cannot access. The direction of the selection pressure towards more efficient catalysis provides excellent training data for ML and enables rapid extrapolation with custom made ML models beyond the improvements encoded in the screened library - by nearly one additional order of magnitude, as demonstrated by a 23-fold improvement in k_cat_. This showcases how combining microfluidic sequence-function mapping and ML-enabled *in silico* extrapolation substantially accelerates enzyme engineering.

**Figure 1:**
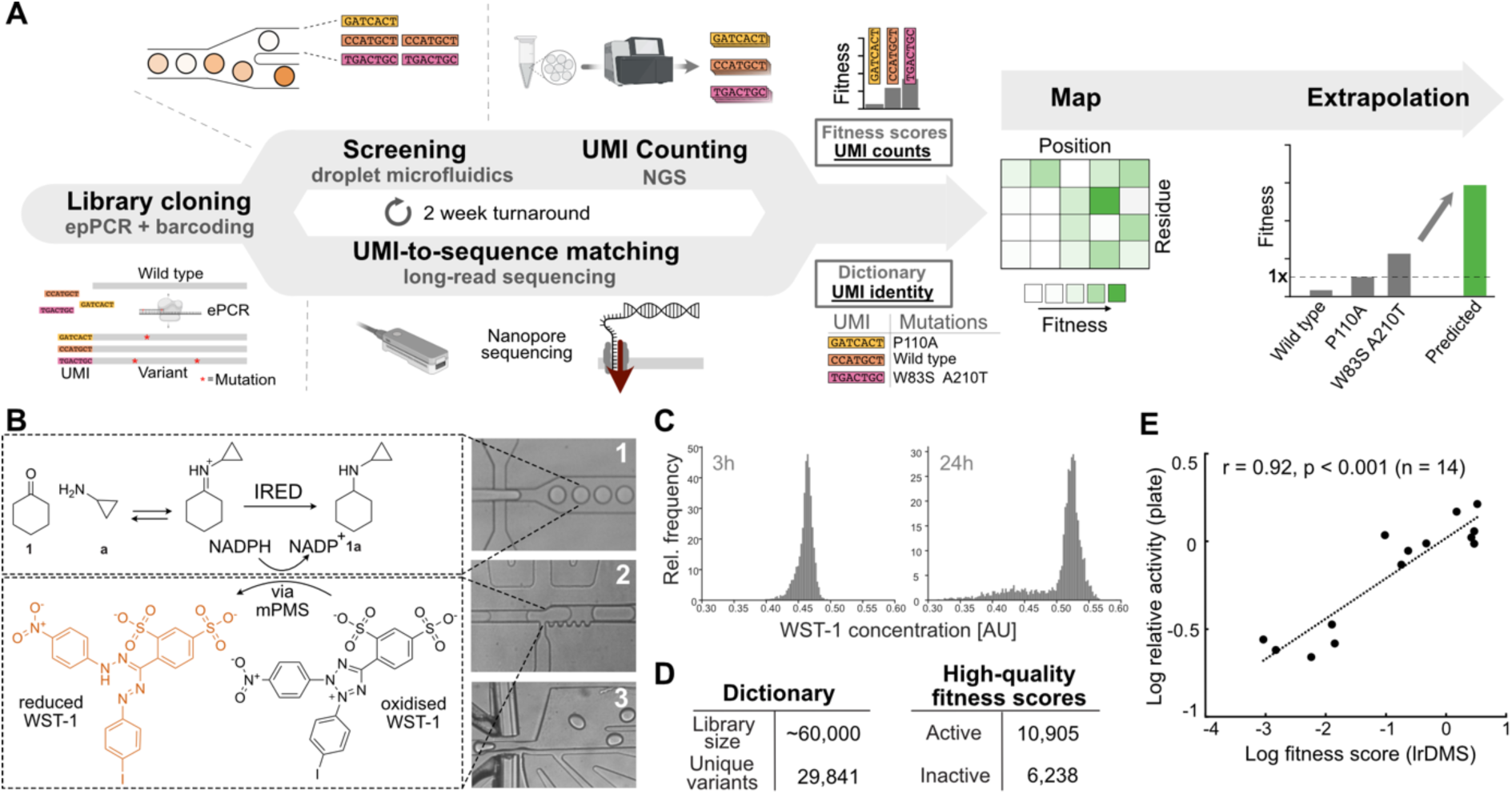
A microfluidics-enabled workflow for rapid large-scale fitness data generation and extrapolation. **(A)** Long read deep mutational scanning workflow. A random mutagenesis library, barcoded with unique molecular identifiers (UMIs), is sequenced using nanopore sequencing to give a dictionary that identifies each library member by linking its UMI sequence to the corresponding variant sequence. The library is screened using a droplet-based uHTS (ultrahigh-throughput screening) assay in multiple replicates. UMIs of the sorted and unsorted library are sequenced using NGS and fitness scores are calculated from the enrichment of individual UMIs upon screening. The fitness information and the identity of the UMIs are combined to give an accurate and quantitative sequence-function map, which serves as the basis for rational and AI-driven extrapolation. **(B)** Ultra-high throughput assay. Substrates (cyclohexanone **1** and cyclopropylamine **a**) are encapsulated with NADPH, *E. coli* expressing the IRED library and lysis agent. After incubation, mPMS and the formazane dye WST-1 are added by picoinjection, producing the colorimetric readout that is used as the criterion of selection by AADS reporting on activity. (**C**) Activity distribution of droplets containing the library after 3h and overnight incubation. The positive population (low absorbance due to consumption of NADPH) increases as the reaction progresses in individual droplets. **(D**) Number of dictionary members and fitness scores obtained (a total of 17,143 high-quality lrDMS scores are calculated from 29,841 unique dictionary members). (**E**) A linear correlation of fitness scores with colorimetric lysate assay data for 14 individual variants chosen randomly from the input library and the library after sorting suggests that the picoliter assay in droplets quantitatively reports on genuine activity levels. *E. coli* lysate for each variant was incubated with cyclohexanone **1** (10 mM), cyclopropylamine **a** (20 mM) and NADPH (5 mM) for 3 h or overnight in Tris buffer (100 mM, pH 8.0). The colorimetric readout was produced by the addition of the formazan dye INT (a less water-soluble alternative to WST-1). A fitness metric for lysate activity was calculated as the logarithm of the ratio of the average variant endpoint to the average wild-type endpoint. [*Abbreviations –* lrDMS: long read deep mutational scanning; epPCR: error-prone PCR; UMI: unique molecular identifier; NGS: next-generation sequencing; IRED: imine reductase; AADS: absorbance-activated droplet sorting; uHTS: ultrahigh-throughput screening]

### Droplet microfluidics rapidly provides a sequence-function map of an imine reductase via deep mutational scanning

Here we address the enzyme engineering challenge of improving the activity of a bacterial IRED from *Streptosporangium roseum* (*Sr*IRED) catalyzing the formation of carbon-nitrogen bonds via reductive amination through large-scale sequence-function mapping. To achieve ultrahigh throughput screening for catalytic function, we use picolitre-sized droplets which act as test tube equivalents (**Figure 1B, Supplementary Figure 1**). These droplets encapsulate single Poisson-distributed *E. coli* cells expressing members of an error-prone PCR (epPCR)-generated DNA library, compartmentalized with lysis agent and substrates, to initiate the enzymatic reaction upon droplet formation in a microfluidic device. The reaction is stopped in a second microfluidic step by picoinjection of the dye WST-1 (to monitor the enzyme cofactor from NADPH to NADP^+^ that stoichiometrically accompanies turnover, **Figure 1C**). The resulting colorimetric readout distinguishes active library members and provides the basis for sorting by absorbance-activated droplet sorting (AADS)^27^(see **Extended Data Figure 1** for a quantification of sorting efficiency by enrichment and **Supplementary Figure 2** showing selective enrichment of IRED activity).

The identity and fitness values of the selected variants were decoded by long-read deep mutational scanning (lrDMS) (**Figure 1A**), allowing the decoding of sequence-function maps in proteins of any size: (i) The *identity* of the variants is encoded by first generating a ‘dictionary’ of 29,841 unique members present before selection (**Figure 1D; Supplementary Figures 3, 4, 5 and 6; Supplementary Note 1**) based on high-quality, 99.5%-accurate long-read sequencing (over the whole length of the gene)^28^, after attaching unique molecular identifiers (UMIs) to their gene sequences. After selection, this dictionary allows the decoding of variant identity based on the short UMI only, saving on sequencing costs and effort (one pooled NextSeq500 run compared to >10 Oxford Nanopore flow cells). (ii) The *fitness* of the selected variants is computed by counting the occurrences of the UMIs (giving a ‘fitness score’) before and after selection (**Supplementary Figure 3 and Supplementary Note 1**; for definition of fitness see **Supplementary Figure 11**): a calibration curve (relating the conversion rates of variants with UMI counts; correlation coefficient r = 0.92) suggests that fitness scores capture quantitative fitness information of variants across a wide range of activities (**Figure 1E**). We performed lrDMS of *Sr*IRED by sorting approximately 1.5% of the most active variants (of 6.9 million droplets screened at a rate of 200 Hz and an occupancy of 50%, a 57.5-fold oversampling of the library), revealing 10,905 unique members to be active (with quantitative activities as UMI-based fitness scores; **Figure 1C, Supplementary Table 1**) and, conversely, 6,238 variants to be inactive (occurring in the input library but not in the output library, i.e. below the selection threshold; **Figure 1D).** This workflow thus quantifies the fitness of >10^4^ randomly selected single and higher-order variants within 2 weeks: a scale, speed and cost not feasible with plate screening or robotic workflows^21^.

### Long-read deep mutational scanning reveals the mutational profile of an IRED

Experimental lrDMS-derived fitness scores for 17,143 variants provide insight into how mutations across the *Sr*IRED sequence and structure affect activity as single mutations (*mutability*) and whether they can be combined in higher-order variants (*combinability*). On average, mutational effects are detrimental to fitness, with a median fitness of −1.2 (0.3-fold wild-type activity), and only 0.9% of mutations showing significantly improved fitness (1 SD > wt). A total of 6,283 sequences (37% of all quantified variants) are classified as inactive because they were present in the input library but never in the output after selection (**Figure 2A**). We calculated the per-position median fitness of all single-variant fitness scores in the IRED sequence (on average 4.3 datapoints per position) as a proxy for mutability, i.e. the capacity of a site to remain functional following mutation (**Figure 2C**). The first shell and the proposed catalytic residues^29,30^ show very low mutability with the exception of T241 that contacts the ketone substrate and is positioned at the bottom of the binding pocket, flanked by all three catalytic residues (**Figure 2D and E**). Positions with high mutability are defined as hotspots and e.g. cluster in specific regions of the IRED sequence (e.g. around position 40). Mapping mutability onto a crystal structure of *Sr*IRED^30^(**Figure 2E**) reveals hotspots, particularly at the N-terminus (**Extended Data Figure 2B**) and in helices and loops of the N-terminal domain **(Extended Data Figure 2C and D**). We also observed a pattern of recurring hotspots in the central helices at the dimer interface (**Extended Data Figure 2E and F**), consistent with previous IRED engineering campaigns that selected mutations in this region^20,31^.

**Figure 2:**
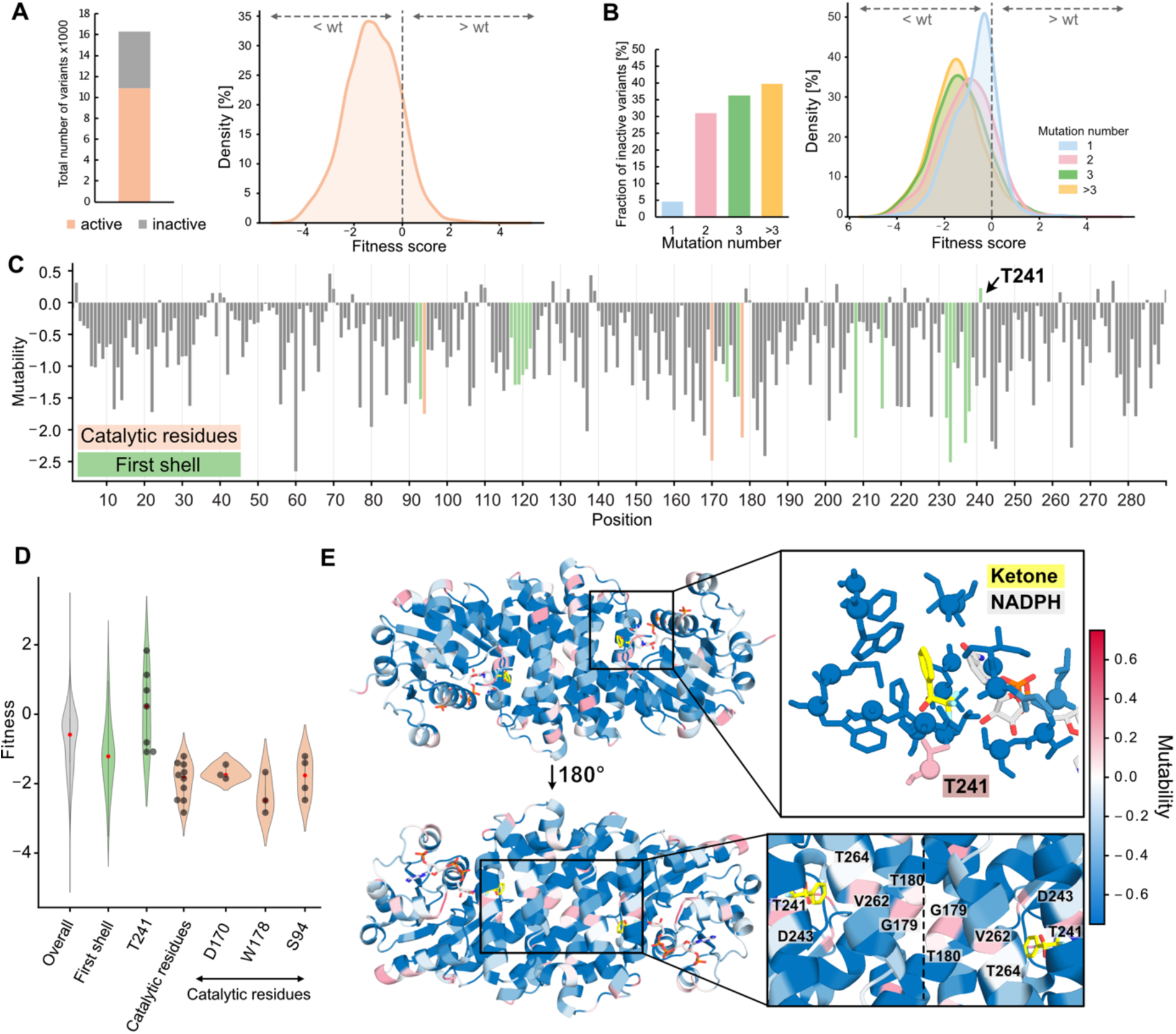
Distribution of mutational effects in an error-prone PCR library. (**A**) 63% of all assessed variants show detectable activity. Distribution of fitness effects of all mutations with detectable activity shows, that most variants are less fit than the wild-type enzyme, with a small fraction of mutations showing an improving effect (**B**) Distribution of fitness effects for single (blue), double (pink), triple (green) and more than triple (yellow) mutations shows, that the mutational effect is dependent on the number of mutations introduced. The more mutations are introduced the more abundant are inactive variants (left) and the more negative the average fitness of functional variants. (**C**) Positional distribution of median fitness shows that there is a large difference in the mutability of different positions in the protein (orange: catalytic residues; green: first shell). (**D**) Violin plot showing the fitness of variants with different mutational backgrounds. Mutants of first shell and active site residues are on average less fit than the average mutations (−1.21 *vs* −1.82). However, there is one exemption with positive mutability and three highly fit mutations: T241. No mutation at any active site position (D170, W178, S94) shows improvements. (**E**) Structural map on median fitness shows distribution of mutability on structure (*blue*: low mutability, *red*: high mutability). The first shell and the dimer interface are highlighted.

To characterize the enzymatic properties of hits, Michaelis-Menten kinetics of 12 single variants with fitness scores >0 and the variant with the top fitness in the library (V6L V67I) were measured (**Supplementary Figure 7 and 8, Supplementary Table 2**). Overall, single mutations at position T241 lead to the highest catalytic efficiency (4-fold improved k_cat_/K_M_ along with a 3-fold improved k_cat_ in T241A and T241S). 8 of 12 variants showed an improved *k_c_*_at_ while only the T241 mutants and E146V showed improved *K*_M_ values. As in previous DMS datasets^32^, along with catalysis, thermostability was improved for 6 (of 12) variants (up to a T_m_ of 50 °C for T241A *vs* 39 °C for wild type), reflecting the need for long term integrity of the catalyst for the entire duration of the reaction in droplets (here up to 24h)^33^.

The ultimate challenge for biocatalysts is their successful application under conditions relevant for larger-scale synthesis. We thus tested *Sr*IRED wild type and the T241A variant in a biotransformation with catalytic amounts of NADPH and a cofactor-recycling system consisting of a variant of the formate dehydrogenase from *Candida boidinii* (FDH-QRN) and formate^34,35^ (**Extended Data Figure 3E**), where product formation of T241A was markedly improved from 83% to 97% after 4h (**Extended Data Figure 3F, Supplementary Figure 10**). We observed excellent correlations between specific activities (based on initial rates of NADPH depletion) and the conversions in biotransformations, suggesting that measuring product formation in our assay reliably identifies catalysts with superior properties for practical applications.

Using nanopore-based long-read DMS to sequence the entire protein, rather than fragments obtained in Illumina sequencing, gives access to the joint effects of multiple mutations and their combinability (i.e. the potential of a mutation for constructive interaction; see **Supplementary Figure 11** for definition) which can be assumed to be a major contributor to evolvability (i.e. the capacity of a mutant to originate improving trajectories in evolution). Our lrDMS data reveal that, along with an increasing number of mutations per variant, a higher fraction becomes inactive resulting in decreasing average fitness (**Figure 2B**). Among the 11,958 available datapoints for higher-order variants, we used 6,188 with measurable fitness scores for all mutations in the higher-order variant to quantify epistasis by comparing their actual fitness to the calculated additive fitness from single-point mutation data alone **(Figure 3A, Supplementary Figure 11)**. As it can be assumed that there is no positive epistasis in the remaining 4,130 inactive variants, the overall fraction of positive epistasis is lower than that for negative epistasis, in line with previous observations^36,37^. However, a large fraction of improved variants (67 of 84, 80%) are positively epistatic (**Figure 3A and B**) and 60 out of 67 (90%) of these contain single mutations that individually are neutral or negative. These mutations would have been disregarded in traditional screening campaigns that iterate combinatorial screening of improving variants from single site saturation libraries.

**Figure 3:**
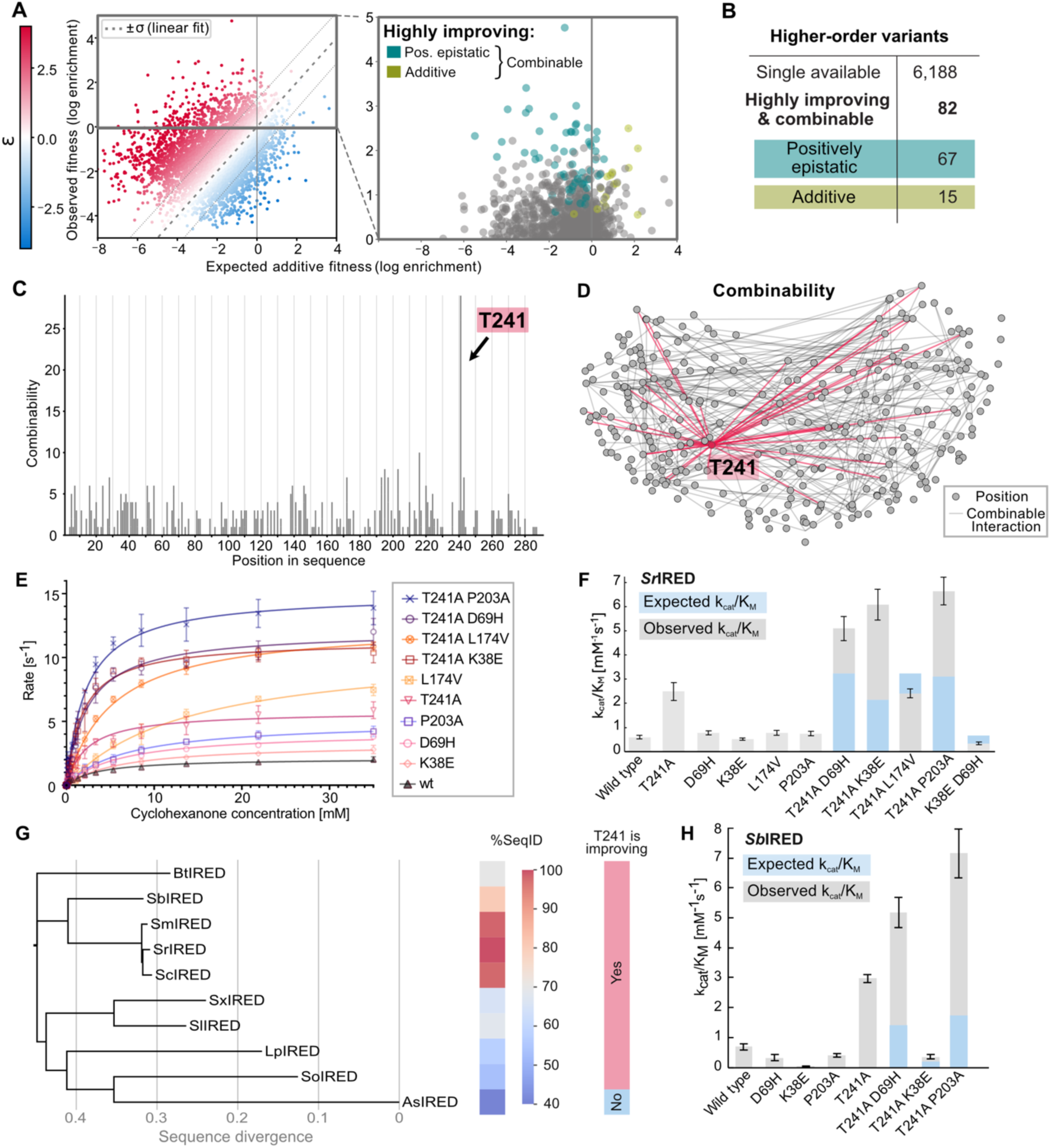
Combinability distribution informs rational engineering. **(A)** Scatter plot showing expected fitness for all higher-order variants from single mutation data (calculated via the product model) versus observed fitness. Positive epistasis is shown in red, negative epistasis in blue. On the right, variants with larger than wild type fitness (observed fitness > 0) are shown. Positively epistatic (1SD > additive) and additive (1 SD > negative epistasis) variants with high fitness (1 SD > wt) are shown in red and **purple**, respectively. **(B)** Of 6188 higher-order mutations in this dataset (2-5 mutations with single mutation data for all mutations), 84 are highly improving (1SD > wt) and 82 are highly improving and combinable. Most of these variants are positively epistatic (67) and only 15 are additive. **(C)** Positional frequency in highly active and combinable variants. T241 is the global hotspot of combinability, followed by V216. **(D)** A combinability graph displays the interactions of mutations at distinct positions that maintain activity above the experimental screening threshold, suggesting they can co-exist. Nodes represent positions in the IRED sequence that are grouped by distance in IRED structure. Connections represent cooccurrence in a higher-order variant the corresponding fitness of which is larger or equal to the combined effect of single mutations: combinable. The high node degree of T241 (edges in red) indicates high potential for combinability and positive epistasis with all parts of the protein. **(E)** Michaelis Menten kinetics with single variants chosen for rational engineering with T241A and their combinations showing high performance of combined variants apart from combinations with L174V and indicates general combinability of T241A. **(F)** Comparing expected k_cat_/K_M_ of single variants (blue) to the observed k_cat_/K_M_ (grey) of rationally engineered higher order variants shows positive epistasis for all rationally engineered variants apart from T241A L174V. Standard errors of the fit and three technical replicates are represented with a 95% confidence interval. **(G)** 9 homologs to *Sr*IRED (with sequence identities ranging 40-98%) were tested in the absence (native) and presence of the T241A mutation. Improved k_ca*t*_/K_M_ values were observed for 8 of 9 homologs (apart from *As*IRED), showing transferability of T241A to variants with as little as 53% sequence identity (*So*IRED; **Extended Data Figure 4**). **(H)** Mutations D69H, K38E and P203A from SrIRED mutability hotspots were transferred to *Sb*IRED (80.3% sequence identity) and kinetic parameters measured for mutants with and without T241A. Comparison of expected additive (blue) and experimentally observed (grey) k_cat_/K_M_ values shows strong positive epistasis with T241A when either D69H (3.7-fold) or P203A (4.1-fold) were present, suggesting transferability (weaker positive epistasis for K38E with 1.8-fold). Standard errors of the fit and three technical replicates are represented with a 95% confidence interval.

Having established that positive epistasis plays a major role in shaping constructive trajectories in *Sr*IRED evolution, we next investigated the sequence background of combinability. Among the 82 higher-order mutants shown to be highly active with no negative epistatic interactions, one highly combinable mutation with high potential for constructive interaction stands out: the global hotspot at position 241 (in the first shell, **Figure 2E**, **Figure 3C, and Supplementary Figure 12**; a further combinability hotspot is evaluated in **Supplementary Figure 13)** with contribution mostly from T241A, the mutation previously identified with the highest improvement in catalytic efficiency (4-fold) and 11°C improvement in T_M._ The central role of position 241 as a combinability hotspot is evident: it exhibits combinable interactions with positions widely distributed across the N-terminal and C-terminal domains (**Figure 3D** and **Supplementary Figure 12, Supplementary Table 3**). Elevated protein stability may explain the high combinability of T241A in cell-based assays as for example our droplet assay, by enhancing mutational robustness, extending the stability threshold of the protein and allowing incorporation of destabilizing mutations^38–40^. By characterizing positive epistatic mutations with T241A and the distant mutations K38E, D69H, or P203A (see next section), we also observed positive epistasis in catalytic parameters (k_cat_/K_M_ and k_cat_). Indeed, the destabilizing variant T193A, for instance, is part of a larger cluster around a hinge region in the C-terminal domain indicating a shared mechanism of action (**Supplementary Figure 12 G and H**). This is consistent with models where combinability is not only influenced by mutational robustness but also protein dynamics, where distant mutations can yield synergistic effects by optimizing the conformational landscape of the enzyme^41,42^.

lrDMS-derived combinability data provide a blueprint for bypassing negative and harnessing positive epistasis throughout the entire protein during rational engineering. We thus matched the combinability hotspot T241A with beneficial mutations distributed throughout the protein, rationally engineering combinations previously not observed in the dataset. We chose P203A, K38E and D69H (hotspots at the dimer interface and in loops interacting with the cofactor), as well as L174V, the second-best mutation in the first shell (**Figure 3G and H**). Three out of four variants exhibit positive epistasis, with the highest improvement being 11-fold in T241A P203A. By contrast, implementing mutations *without* evidence for combinability such as K38E and D69H, leads to double mutant K38E D69H, characterized by strong *negative* epistasis (with a 2-fold drop in k_cat_/K_M_ compared to wild type). The diverging outcomes in rational engineering of higher-order variants underlines the relevance of lrDMS combinability scores for tapping effects that otherwise are impossible to predict (**Figure 3H**).

The value of the fitness information gathered by our lrDMS experiment would be enhanced, if the lessons learned could be transferred to related proteins. We probed this portability of these catalytic features by introducing the T241A mutation - distinguished in our initial analysis by high mutability and combinability - into 9 related IREDs with decreasing sequence identity (98% to 40%; **Figure 3G**, **Extended Data Figure 4, Supplementary Figure 14**). In eight of these IREDs, T241A improved the activity, yielding up to a 7-fold increase in k_cat_/K_M_ (*Bt*IRED) and 5-fold improvement in k_cat_ (*Sb*IRED), as well as enhanced melting temperature accross the board (up to 15 °C; in *Bt*IRED; (**Extended Data Figure 4, Supplementary Table 4)**. Productively transplanting T241A was possible with as little as 53% sequence identity (i.e. beyond that typically covered by patent protection^43^ and corresponding to hundreds of million years of evolution^44^), but it was no longer beneficial once 41% identity was reached.

Not every mutation is as easily portable as T241A: individually transferring the three previously identified mutations K38E, D69H and P203A into *SbI*RED (80.3% sequence identity) improved k_cat_ only marginally (+10%) or had no effect (**Figure 3H**, **Supplementary Table 5**). However, when the same mutations were introduced alongside T241A, they led to improvements of up to 10-fold improvements in k_cat_/K_M_ (*Sb*IRED P203A T241A) and 9-fold in k_cat_ (D69H T241A; **Figure 3H, Supplementary Table 5**), an excess of up to 4-fold (*Sb*IRED P203A T241A k_cat_/K_M_) beyond the effect of introducing T241A alone. The portable feature here is apparently not the direct effect of a mutation, but the positive epistatic synergy uncovered in our original analysis of combinability in *SrI*RED. Thus, key features of the profile recorded for one IRED are transferable to a family of sequences, allowing data gathered in a single deep mutational scan to inform the engineering of other sequences and substrates, thereby replacing semi-empirical intuition in protein engineering with solid evidence-based instructions for mutagenesis.

A second level of portability of the profile of *SrI*RED concerns the universality of enzyme activity improvements to new substrates. Again, we probed the guidance provided by mutability and combinability hotspot T241A with ketone, amine and imine substrates: we observed improvements across 8 of 11 alternative substrates (tested in initial rate assays as well as biotransformations, **Extended Data Figure 3**). For example, total turnover in the conversion to **4a** was improved from 41% to 67% (after 24h; **Extended Data Figure 3F**). Introducing T241A improved the conversion of more challenging bulky ketones such as indanone (from 3% to 5%) and 4-phenyl-2-butanone (from undetected to 0.8%) (**Extended Data Figure 3F**). On the other hand, there were mutations that are uniquely improved the conversion co-substrate **1a** used for screening (K38E, **Extended Data Figure 3D**).

Their substrate promiscuity makes IREDs versatile catalysts for biocatalysis in pharmaceutical synthesis. To isolate generalist vs substrate-specific features, we conducted two additional deep mutational scans with different substrates, the combination of methylamine **b** and cyclohexanone (**1b**, close in structure to **1a**) and the intramolecular imine 2-methyl-1-pyrroline **9** (structurally distant to the **1b** and **1a** imines) (**Extended Data Figures 5 and 3A**) and compared the datasets with our original *Sr*IRED **1a** dataset as well as two other datasets recorded with different IREDs and substrates.

We found that the mutability of individual residues transfers well across all three substrates scanned for *Sr*IRED, with correlation coefficients >0.64. Notably, it also correlates well with mutational scan of a different IRED and a different substrate^45^: the mutability profiles of *Sr*IRED and IRED88 (44% sequence identity) have a Pearson r of 0.44 (**Extended Data Figure 5, Supplementary Figure 15**). With decreasing sequence identity, this correlation ceases to hold. When we carried out a deep mutational scan with *Pc*IRED (30% sequence identity) and a different substrate (see next section), no overall correlation of mutability was apparent, but occasionally hotspots (e.g. 105 *Sr*IRED numbering corresponding to E107K in *PcI*RED) are consistently detected in all three scanning campaigns (**Extended Data Figure 5, Supplementary Figure 15)**. Within a single IRED, we identify “generalist” residues (i.e., those mutable across all substrates: 110, 138 and 290) and “specialists” (i.e., those mutable for *one* substrate but not multiple ones; **Supplementary Figure 16)**. These specificity-conferring positions are found mainly outside the first shell, underlining the notion that substrate specificity has to be engineered globally^46^. The combinability parameter appears to be much more substrate-specific (**Extended Data Figure 5, Supplementary Figure 17**). For example, T241 - the combinability hotspot shown to be portable from one enzyme to another - cannot be transferred to other substrates (neither for **1b** nor for **9**).

### AI-assisted engineering yields highly improved positively epistatic variants for application in biotransformations

Directed evolution relies on sampling and testing the vast combinatorial diversity of sequence space through iterative fixation of mutations, thereby constructing trajectories towards higher fitness. Rational engineering aims to bypass these sometimes long and labor-intensive screening campaigns but is often constrained by the preponderance of negative epistasis, missing out on large leaps in activity provided afforded by positive epistasis, even when information on improving single mutations is available^10^. Machine learning has the potential to discern complex patterns embedded in large datasets that a human protein engineer might overlook. Hypothesizing that the unique long-read sequencing derived combinability score in our lrDMS data can provide a window into future rounds of directed evolution, we use combinability and mutability information from a single lrDMS round for AI-assisted extrapolation to higher-order variants. These higher-order variants have not been observed in the dataset and would otherwise only be accessible by conducting multiple rounds of directed evolution.

#### (i) AI-informed engineering for single mutants

Since not all amino acid changes are accessible via single nucleotide substitutions (in the case of *Sr*IRED only 30% of all single amino acid mutations; **Supplementary Figure 4**), epPCR libraries are biased towards a limited chemical space. We therefore use machine learning to extrapolate to the full chemical space. Specifically, we built a ridge regression model based on physiochemical amino acid embeddings computed from a principal component analysis of the AAIndex database^47,48^, and augmented it with the ESM2^49^ pseudo-log-likelihood ratio of a given mutant sequence vs wildtype, as proposed by Hsu et al.^50^ (**Figure 4A**). We trained this model on the observed single mutants from the lrDMS campaign and extrapolated fitness values for all unobserved single mutations (**Supplementary Figure 18**). We then selected the top five predictions - P203G, T241G, M1P, A251I and E146L (omitting predictions at the same positions to avoid position bias) - for experimental characterization (**Figure 4C**, **Extended Data Figure 6A)**. Of these 5 variants, one showed improved k_cat_/K_M_, three an improved k_cat_, one improved activity in lysate and two of five variants showed an improved melting temperature (improvement > 1SD from wild type). The best predicted variant, T241G, exhibited a significantly higher improvement in k_cat_/K_M_ than all characterized variants in the lrDMS dataset (7-fold, **Figure 4C**, **Extended Data Figure 6A**) and demonstrated substantial conversion improvements in the biotransformation of **1a** (22% to 43% after 30 min; **Extended Data Figure 3E and H and Supplementary Figure 21**). Notably, T241G also increases stability, measured by T_M_ (to 46 °C, up from 39 °C in the wild type, though still below the 50 °C observed for T241A). The learned, extrapolated systematic decrease in side-chain volume at position 241 (threonine in the wild type, alanine and serine in the dataset, glycine in the AI prediction) illustrates the potential of AI-based extrapolation to deliver improved variants that could not themselves have been found given the experimental screening strategy (**Figure 4D**). For T241G to appear in the dataset, two consecutive nucleotide substitutions would have been necessary to reach G (GGN) from T (ACG), a highly unlikely event in epPCR mutagenesis. Learning curves and predictions without ESM (**Extended Data Figure 7**) show that the model’s success is primarily attributable to the experimental dataset: mutant rankings lacking ESM2 information still include 3/5 mutations in the top 5, including T241G (**Supplementary Figure 18**). Conversely, a mutant ranking based on ESM2 alone does not identify T241G until rank 457/5510, and only proposes top-five variants with up to 2-fold improved k_cat_/K_M_ in experimental characterisation, albeit with no variant dropping below the wild type performance (**Supplementary Figure 18 and 20**). While these results indicate that information from ESM2 can serve as important feature or filter to eliminate poorly performing variants, they also show that it cannot replace experimental data in the context of the relevant reaction. This outcome highlights how the synergy of experimental data and data-driven strategies unlock improvements unattainable by either technique in isolation.

**Figure 4:**
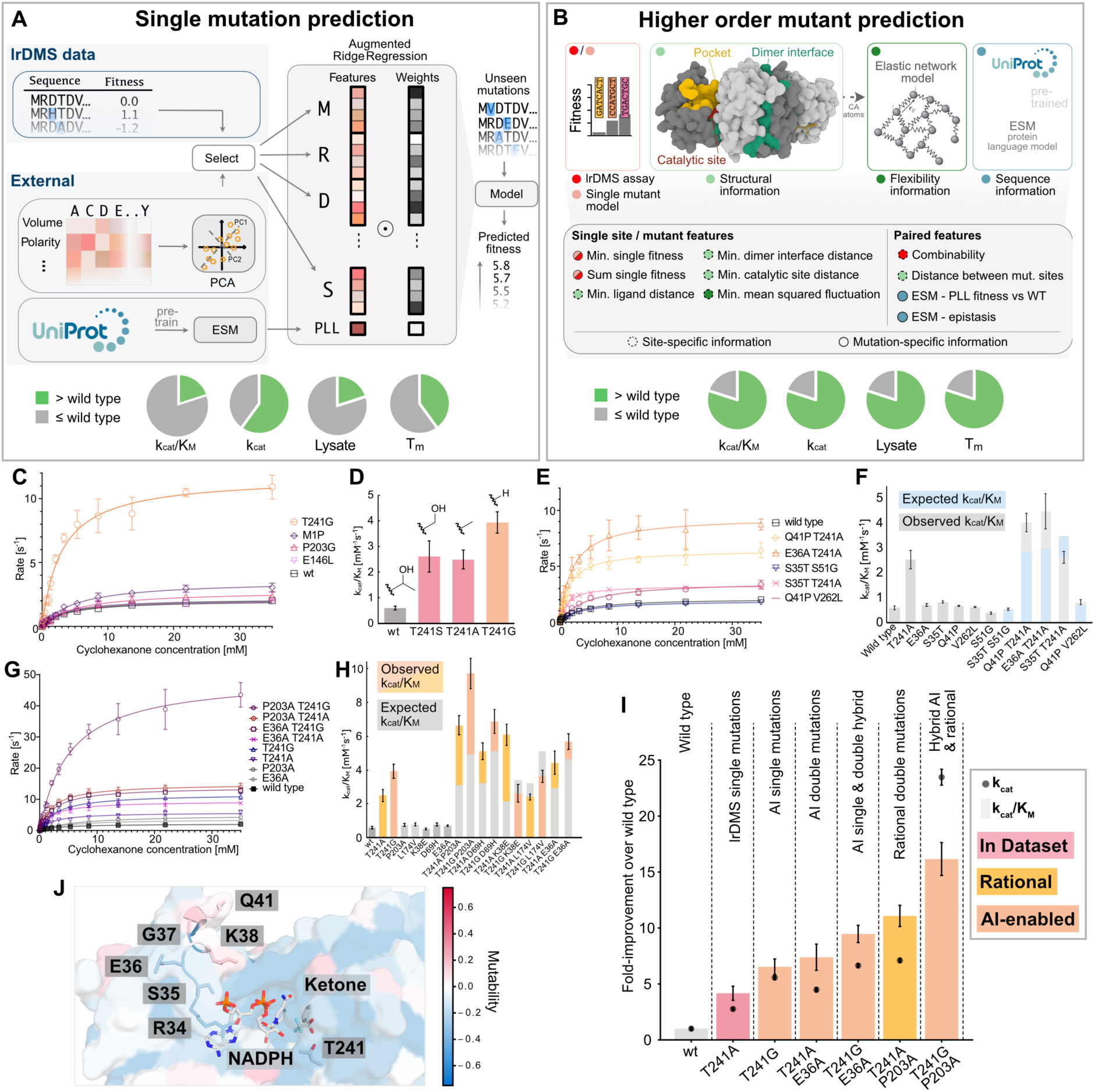
AI and rational engineering enable extrapolation to highly active variants. **(A)** Single mutant model and its performance in predicting improved mutations. lrDMS data and external parameters (ESM pseudo-log likelihood vs wildtype (PLL) and amino acid structure embeddings) are used in an augmented ridge regression model to predict the fitness of single point mutations. The 5 best single point mutations are characterized experimentally (pie chart, **Extended Data Figure 6A**). 2 out of 5 characterized mutations show an improved k_cat_/K_M_ (> 1 SD from wild type), 3 out of 5 an improved *k*_cat_, 1 out of 5 improved lysate activity and 4 out of 5 show an improved melting temperature. (**B**) The double mutant model is based on assay-derived, structural, flexibility and sequence information. Gradient boosted decision trees trained on the lrDMS data provide a predicted fitness estimate for unobserved double-mutants, which is used to rank and select mutants to test. Among the top 5 characterised mutants **(Extended Data Figure 6B**) from the model 4 out of 5 show improved *k*_cat_, *k*_cat_/*K*_M_ and lysate activity. (**C**) Michaelis Menten curves of single point variants predicted by AI showing slight improvements for M1P and E146L and a 6.6-fold improvement for T241G. Standard errors of the fit from three technical replicates are represented with a 95% confidence interval. (**D**) *k*_cat_/*K*_M_ values of T241A mutants (grey: wild type, pink: in dataset, orange: AI-predicted) with side-chain structures highlighting the decrease in steric bulk from threonine to serine, alanine and glycine. (**E**) Michaelis Menten curves of variants derived from the combinability-based model showing improving parameters for 4 out of 5 variants (**F**). The *k*_cat_/*K*_M_ of the top 5 predictions of the higher-order mutant model and corresponding single point mutants (Expected *k*_cat_/*K*_M_ calculated from single mutation data is shown in blue, observed parameters in grey). 2 out of 5 variants show positive epistasis: one negative epistasis and two additivity. The best AI-predicted double mutant shows a 7.4-fold improved *k*_cat_/*K*_M_, better than the best predicted AI single mutant T241G with a 6.6-fold improvement. (**G**) Michaelis Menten plots and (**H**) *k*_cat_/*K*_M_ values for hybrid AI and rational engineering-based strategy where T241G (orange) is transplanted into rationally engineered and AI-predicted variants with T241A (pink). Replacing T241A with T241G has a positive effect in 3 out of 4 cases. A 16-fold improved *k*_cat_/*K*_M_ paired with strong positive epistasis was observed with T241G P203A. The negative epistasis with L174V is conserved but paired with a jump in improvement from 4-fold to 6-fold. Replacing T241A with T241G switches the positive epistatic effect with K38E into a strong negative epistatic effect underlining the influence small residue changes can have on non-linear effects on activity. Replacing T241A with T241G has a similar effect on AI-generated double mutants transforming a 7.4-fold improvement with E36A T241A into a 9.5-fold improvement with E36A T241G. Raw data in **Extended Data Figure 6C**. (**I**) *k*_cat_ (points) and *k*_cat_/*K*_M_ (bars) data for top variants derived from lrDMS data (T241A), the AI single mutation model (T241G), the AI double mutation model (T241A E36A), rational engineering based on both AI models T241G E36A, rational engineering based on single mutation and combinability data (T241A P203A) and a hybrid approach based on AI single mutation and combinability data (T241G P203A). (**J**) Structure of the loop containing residues predicted by the AI model. K38 has been shown to be positively epistatic with T241A in rational engineering, E36A and Q41P show a similar effect.

### (ii) AI-informed engineering for double mutants

Predictive models for higher-order variants must take the combinability of mutations into account to circumvent negative and leverage positive epistasis and are thus dependent on training data that captures long-range interactions of mutations. We developed a tree-based model to automate rational engineering of higher order mutants (**Figure 4B**), focusing on double mutants. The model is trained on lrDMS fitness scores and has access to information on the protein structure (distance between mutated residues, distance to active site residues, dimer interface and first shell, flexibility information computed from an elastic network model^51^). Like in the single mutant model, evolutionary information is incorporated via the protein language model ESM2. Predictions were ranked (**Supplementary Figure 18**) and the 5 best variants were tested experimentally (**Figure 4E and F**, **Extended Data Figure 6B**). We found that 4 out of the 5 tested variants showed considerably higher k_cat_/K_M_, lysate activity and T_M_ than the wild type. The best performing predicted double variant E36A T241A showed a 7-fold improvement in k_cat_/K_M_ (**Figure 4E and F**, **Extended Data Figure 6B**) and improved the conversion in the biotransformation of **1a** from 22% to 35% after 30 min (**Extended Data Figure 3E and 3H and Supplementary Figure 21**). When comparing k_cat_/K_M_ values of the double mutants with respective single mutations, we observed positive epistasis in 2 out of 5 variants, additivity in 2 out of 5 variants, and negative epistasis in 1 variant, showing that the model was able to predict combinability in 4 out of 5 cases (**Figure 4F, Supplementary Figure 19**). The top predictions emerging from this model appear to focus on a loop involved in binding the NADPH phosphate group (**Figure 4J**), consistent with a reorientation of the cofactor that, in combination with mutation T241A, enables a more favorable substrate binding mode that leads to the observed synergistic effects. The successful prediction of multi-residue cooperativity indicates the ability of the model to learn complex structural patterns as the basis for higher-order mutants. This can be primarily attributed to the experimental data. Learning curves (**Extended Data Figure 8**) show that large amounts of data are needed (jump in performance at 3,072 datapoints) and that addition of ESM on top of assay features has minor influence on model performance (as evidenced by ablating ESM features, **Supplementary Figure 18**). Furthermore, a SHAP analysis^52^ (**Extended Data Figure 9, Supplementary Figure 24 and 25**) identifies assay-labeled data as the core feature of the prediction, an observation which is also supported by direct analysis of the learned decision trees. Finally, experimental characterisation of the top 5 purely ESM predicted double mutants shows up to 3-fold improved variants, not rivalling the 7-fold improvement observed when assay labelled data is included (**Supplementary Figure 18 and 20**).

#### (iii) Combination of AI and rational engineering

Access to even greater activity improvements became possible by merging information on the combinability hotspot T241 (from lrDMS, which led to the rationally engineered variant T241A P203A, the best variant so far) with the AI-predicted extrapolation from T241A to T241G, which substantially boosted the single mutant activity. Hypothesizing that T241A’s combinability would be transferable, we replaced T241A with G in all double mutations from rational engineering and the best AI double mutant. In 4 out of 5 cases k_cat_/K_M_ levels increased markedly (**Figure 4G and H, Extended Data Figure 6C**). With a 16-fold improvement in k_cat_/K_M_, driven by a 23-fold improvement in k_cat_, T241G P203A surpasses all previous activity improvements and is the best variant generated in this study. This improvement in catalytic efficiency translates into an improvement in conversion for the biotransformation of **1a** (22% to 42% after 30 min; **Extended Data Figure 3E and H and Supplementary Figure 21**). To evaluate how stereoselectivity changed during AI-assisted engineering, we assayed relevant variants for the reductive amination of the chiral ketone (*R*)-3-methylcyclohexanone (**5**) to install a second stereocenter using cyclopropylamine (**a**) as the amine donor. Whereas the wild type enzyme yielded a diastereomeric ratio (d.r.) of 30:70 at 47% conversion, T241A and all six rational and AI-engineered variants showed enhanced stereoselectivity ranging from 15:85 to 10:90 d.r. at 61% to 90% conversion (**Extended Data Figure 3E and G and Supplementary Figure 22**). This indicates mutating residue T241 to either alanine or glycine principally drives the increased stereoselectivity in the reductive amination of **5** with **a**. Consistent with the catalytic efficiency measurements, the best-performing variant of this study in terms of conversion and stereoselectivity was T241G P203A (90% conversion, 10:90 d.r.; **Extended Data Figure 3E and G and Supplementary Figure 22**) which we obtained *via* a hybrid approach combining rational and AI-informed methods. This outcome demonstrates how merging combinability information into the single-mutant AI model can produce gains that exceed both AI and rational engineering on their own (**Figure 4I**).

#### (iv) Application of lrDMS for the biocatalytic synthesis of the drug Tecalcet

IREDs are versatile enzymes owing to their substrate promiscuity, but bulky and hydrophobic substrates (in particular isopropylamines^53,54^) remain challenging with currently available IREDs, necessitating further protein engineering. We addressed this challenge via large-scale sequence-function mapping using lrDMS for the final assembly step for the reductive coupling between aldehyde **(10)** and amine **(f)** to synthesize the drug Tecalcet for which we selected the pre-engineered IRED variant M3 from *Penicillium camemberti* (*Pc*IRED)^53^ as a starting point (**Extended Data Figure 10A, Supplementary Figure 32, 33, 34 and 35**). Our microfluidics-based workflow requires only 0.8 mg of aldehyde per dataset, thus keeping screening costs low. Compared to *Sr*IRED and the model substrates **1a**, the average fitness in the sequence-function dataset for *Pc*IRED and **10f** is lower (−1.7 compared to −1.2 for *Sr*IRED), consistent with the notion that bulky and hydrophobic substrates are harder to engineer for, and that beneficial mutations are less common (**Extended Data Figure 10B**). Nevertheless, fitness information on 6897 variants was collected, with 4286 variants showing detectable activity (62%, comparable to SrIRED’s 63%) and 2611 were inactive. Overall mutability in *Pc*IRED is decreased (to −1.3 from −0.6 for *Sr*IRED), making it a more challenging engineering target, but mutational hotspots remain apparent, especially abundant in the C-terminal domain (**Extended Data Figure 10C, Supplementary Figure 36**). Whereas in *Sr*IRED, the N-terminal domain was more mutable than the C-terminal domain (−0.5 vs −0.7), *Pc*IRED shows the opposite trend: higher mutability in the C-terminal domain (−1.1) than in the N-terminal domain (−1.4). Although no global combinability hotspot comparable to T241 for *Sr*IRED could be identified for *Pc*IRED, positions with high combinability (e.g. R61) and mutability stand out (e.g. A181 in the first shell) (**Extended Data Figure 10C and D**). We employed this data with the engineering framework established for *Sr*IRED to engineer *Pc*IRED: (i) Rational engineering based on combinability suggested to join R61L at the most combinable position with the first shell mutability hotspot A181T (**Extended Data Figure 11A**). The resulting double mutant exhibited strong positive epistasis with a K_M_ 2.3-fold lower and a k_cat_/K_M_ 2.5-fold higher than additive, respectively. This synergy is remarkable given that R61 is 25.9 Å away from the active site (site of hydride transfer) and 37.7 Å from A181, defying traditional protein engineering approaches focused on first-shell residues. Remarkably, R61L on its own is nearly neutral (15% increase in k_cat_/K_M_) so it would have been disregarded in traditional directed evolution campaigns which focus on recombining improving mutations. (ii) We also were successful in AI-based extrapolation: Our single mutant AI model identified one hit, A181S (**Extended Data Figure 11B, Supplementary Figure 37**) which gave a 2-fold increase in k_cat_ (albeit a 3-fold rise in K_M_). Our double mutant AI models led to up to 3-fold improved K_M_ values (K249N K296R; 5 out of 9 predictions successful originating from two separate models) (**Extended Data Figure 11C**) and a 2-fold increase in k_cat_ (H199L K249N, 3 out of 9 predictions successful). Success was concentrated in the one of the models that most strongly relied on assay-derived features (*model 2*, **Supplementary Figure 38)**, as compared to the alternative model (*model 1*) that relied more on evolutionary signal from ESM and structural features (**Supplementary Figures 38-40**). As observed for *Sr*IRED, the double mutant models had a higher prediction success compared to the single mutant modeeven though no significant combinability hotspot such as T241A was present.

The mutations derived from combinability and mutability were validated in biotransformations at a 1-mg scale using the top hit, R61L A181T. This engineered variant achieved a 3-fold yield improvement over a previously optimized M3 variant (representing the final outcome of an ultimately stalled evolution campaign^53^; **Supplementary Note 3**). This enhancement after only one round of engineering is comparable to published yield improvements for similar substrates following three rounds of directed evolution (**Supplementary Note 3**). By contrast, when we deliberately disregarded the lrDMS insights and combined A181T with the mutant S262N, characterized by low combinability, yet high improvement (alone bringing about an excellent yield of 62%), we observed *negative* epistasis: the S262N A181T *double* mutant achieved only 50% conversion, whereas additivity would have predicted 93% (**Supplementary Figure 52D and E**). This showcases the applicability of lrDMS in engineering biocatalysts for challenging, pharmaceutically relevant substrates.

## Discussion

Generating sequence-function maps in merely two weeks enables a radically different approach to enzyme engineering. Instead of simply isolating one improved mutant in traditional directed evolution, large datasets are generated that profile the idiosyncrasies of a protein in a way that eludes enzyme engineering based on single site-directed mutagenesis. We obtain a transparent picture of the mutational potential of a protein and its ability to adapt to enhanced functionality. The complex interplay of biophysical properties and biochemical reactivity that has made enzymes harder to engineer than binders is illustrated by insights into (i) hotspots vs regions that should not be touched by engineering (from single mutations), described by mutability, (ii) cooperative vs detrimental epistatic interactions across the entire protein (from higher-order mutations), described by combinability and (iii) a sense of the tolerable mutational burden (for maintaining stability while introducing catalytically beneficial mutations). Previously, these complex interactions were seen as hard to dissect, but algorithmic help from ML allowed for the extraction of useful and interpretable information from sequence-function datasets, as shown by enhancing the activity of our biocatalyst beyond the improvements achieved by hits isolated from the experimental screen.

Several key features of our approach will make this a general method for enzyme engineering:

(i) The use of droplet microfluidics is crucial to assay large libraries to produce datasets with > 10^4^ variants, an achievement that would be prohibitively expensive and time-consuming in plate screening or robotic workflows^45,55,56^. Droplet microfluidics is fast, affordable (even for public sector labs) and scalable, with the throughput depending on the detection method (AADS ∼1 kHz^57^; FADS: >5 kHz or FACS with double emulsion droplets: >10 kHz)^58^. While miniaturizing assays for use in droplet format requires an optical readout, assays for all 7 EC classes are already available^21^, and coupled assays^27,59,60^ or turnover detection with product sensors^61^ cover reactions for which chromo- or fluorogenic substrates are lacking. Methods that do not require an optical readout, e.g. based on mass spectrometry^62^ or Raman spectroscopy^63^, will further expand the range of accessible reactions further in the future.
(ii) Global profiling of the entire enzyme via epPCR offers a hypothesis-free and cheap alternative to targeted mutagenesis (based on proximity to the active site^64^ or phylogeny^65^), which requires the synthesis of designed libraries. Because epPCR mutations are applied randomly (unbiased by position, order of mutation or mechanistic role), the catalytic improvements we observed were brought about by mutations that would not have been targets of a design-led approach. This empirical approach based on mutations in random positions is essentially biomimetic, thus as general as natural evolution.
(iii) The central challenge in enzyme engineering lies in the complexity of epistatic interactions^10^, which are not fully captured by short-read Illumina-based approaches^66–68^ (limited to ∼600 bps, i.e. ∼200 amino acids). Remedial stitching read approaches are poorly scalable^69,70^, unless the proteins are exceptionally short. Most bacterial and eukaryotic proteins are longer (on average 320-472 amino acids^71^), so capturing long-range epistasis on this scale requires the lrDMS sequencing workflow outlined here. Previously, a scalable approach of this sort has been successfully employed for a binding domain^72^, but not for an enzyme. Elucidating long-range epistatic interactions requires sequence-function datasets containing full-length gene sequences (see **Supplementary Table 4** for an overview of enzyme DMS studies highlighting the limited availability of quantitative sequence-function datasets over the whole gene length). Ignoring this crucial information will make datasets biased and less useful for AI interpretation. Indeed, our observed 23-fold rate (*k*_cat_) improvement in the *Sr*IRED engineering case was an order of magnitude higher than the rate improvements (1.3-fold) achieved by the only published previous AI-informed IRED engineering campaign, mostly based on single mutation data^45^. Although *Sr*IRED is an already proficient IRED for the substrate pair used (**1a**) and good catalysts are typically harder to improve, the evolution campaign has yielded a substantial improvement that created a variant amongst the fastest IREDs on record (**Supplementary Tables 10 and 11**). Similar success was achieved in the engineering campaign for *Pc*IRED for a substrate acknowledged to be challenging^54^. By identifying positive long-range epistatic interactions inaccessible via short-read sequencing, we rescued a stalled evolution campaign in only one round of engineering.
(iv) Finally, high hit rates are crucial in typical enzyme engineering campaigns to reduce time and cost of experimentation. The high success rate of predictions of catalytic improvements in our *Sr*IRED work (12 out of 15 tested variants, or 80%, showed an improved k_cat_, **Extended Data Figure 6**) means that very few mutants must be experimentally tested. This compares favorably with ML-based engineering approaches in the low or zero input data regime with hit rates of 2-3%^23,73^ and design approaches where testing of thousands of mutants was necessary to identify three functional catalysts^74^. We thus achieve significant reductions in time and cost for enzyme engineering by pinpointing markedly improved hits among only a handful of designed sequences after only a single round of screening in microdroplets while recent ML-based approaches operating with less sequence-function data still require multiple rounds of optimization^75,76^.

Beyond achieving specific improvements of this IRED, the ability to quickly and cheaply generate datasets that enable AI-based extrapolation for hard-to-predict phenotypes like enzymatic activity addresses a key unmet need for AI-assisted protein engineering^77^. Structure prediction by AlphaFold^78^ and design of protein backbones^9^ are now feasible and routinely used, but do not address function directly. They benefit from the availability of large-scale structure repositories (e.g. the PDB), but equivalent training datasets for protein function still need to be created to enable prediction with similar ease. For data-driven models, the outcome is determined by the nature of the input datasets (and the way they were generated). Previously enzyme stability was enhanced and appeared to be the main driver, e.g. in zero-shot AI approaches based e.g. on PDB data^23,24^. However, catalysis is a function of biophysical properties, specificity *and* biochemical reactivity. Hence, for catalytic enhancement in particular, an assay-labeled experimental dataset based on an assessment of turnover in a target reaction will be crucial for a comprehensive description of enzyme proficiency. When all assay labels directly relate to function, ML interpretation should reveal a clearer ‘response’ to trigger improvements in all properties, including biocatalytic turnover. Moreover, the portability of features of lrDMS sequence function maps to other enzymes, substrates and even completely new engineering scenarios shows its potential to inform future zero-shot tools for enzyme engineering. Equipped with multiple profiles, protein engineers can take account of these evidence-based rules to make informed strategic choices when tailoring biocatalysts for applications, avoiding the effort of a directed evolution campaign. In this context, AI can be instructive in unraveling the complex interdependent combinatorial matrix of multiple substrates, mutability and combinability. Starting with a catalytic selection in our system, it will be interesting to follow up with a complementary microscale assay system, HT-MEK^79^, which can analyse fewer variants (on the order of thousands instead of tens of thousands in microfluidic droplets) but provides non-aggregated, high accuracy kinetic, thermodynamic and biophysical profile. Using these systems in succession may gradually refine the functional information to improve outcomes of AI extrapolations and complement summary ‘fitness’ assessment in initial droplet screening.

The combination of a high throughput assay and AI is important as unbiased approaches are data hungry: most published AI models utilizing experimental data use focused site-saturation libraries (affording data depth in a smaller combinatorial matrix) for model improvement and extrapolation^77,80^. Indeed, analysing the performance of our models with increasing data input shows that large ground truth datasets are needed (**Extended Data Figure 7 and 8**) and substantial improvements of the models may require experimental input on a scale that only *ultra*high throughput techniques such as droplet microfluidics offer.

Our demonstration of the portability of catalytic features rooted in a rational framework based on mutability and combinability makes it tempting to speculate whether the characterisation of individual proteins via sequence-function datasets will provide a growing empirical body that allows further extrapolation and meta-analysis – perhaps revealing complex, yet unifying features for mechanistic classification by ML analysis. This large-scale molecular information would in turn trigger testable hypotheses that should reinvigorate mechanistic studies and, apart from being useful, bring us closer to a fundamental understanding of function. The language of physical-organic chemistry has provided a mechanistic framework for engineering small molecule reactivity and catalysis. Now a hitherto elusive language for describing enzyme catalysis in its enormous complexity seems in reach, and can be developed and probed systematically.

## Supporting information

Combined SI

## Data and code availability

Machine learning models and python code for analysis are available via https://github.com/Hollfelder-Lab/lrDMS-IRED. Fitness scores are available via https://zenodo.org/records/14544486. Raw sequencing data will be made available via the European Nucleotide Archive.

## Acknowledgments

M.G. holds a Trinity College Benn W Levy studentship, S.V.M. received funding from the Vice-Chancellor’s award from Cambridge Trust and from the UKRI Centre for Doctoral Training in Application of Artificial Intelligence to the Study of Environmental Risks (EP/S022961/1). The European Union Marie-Curie networks ‘Oligomed’ (956070), and ES-Cat (722610), and an Individual Fellowship (MSCA-IF 750772) provided support for F.E.H.N., P.Z. and T.S.K, respectively. F.M.W. holds scholarships of the Cambridge Trust, Stiftung der Deutschen Wirtschaft (sdw) and Evonik Stiftung. E.J.M. holds scholarships of the BBSRC (BB/M011194/1) and O.J.K of the EPSRC (EP/R513180/1). F.G.M. and T.K thank NWO Sector Plan for Chemistry and Physics and ERC StG (638271) for funding. F.H. was an ERC Advanced Grant holder (695669) and acknowledges support by an Ignition grant from the ACS CGI Pharmaceutical Roundtable.

## Methods

### Ultra-high throughput screening in droplets

*Droplet screens of SrIRED with substrates 1a:*_All solutions were filtered before they were applied to microfluidic chips. Aqueous solutions without protein components were filtered using Millex-GV filters (Merck), Novec HFE-7500 (3M) with 2% (w/v) 008-Fluorsurfactant (RAN Biotechnologies) was filtered using GD/X 13 filters (Whatman), and aqueous solutions containing protein components were filtered using Minisart syringe filters (Sartorius). 0.5×1mm (I.DxO.D.) PTFE tubing (Bola), glass syringes (Hamilton, SGE) and a syringe pump system (Nemesys, Cetoni) were used to inject solution onto polydimethylsiloxane (PDMS) chips. A high-speed camera (Phantom, Miro eX4, Vision Research) was used to determine droplet size and frequency. Electric currents were applied via a function generator (TG2000, TTi), a pulse generator (TGP110M 10MHz, TTi) and an amplifier (Trek, Model 601C). Droplets were stored in custom-made droplet storage devices^21^. Droplets were generated at 2000 Hz using a flow-focusing device (**Supplementary Figure 1, Supplementary methods**) with cell solutions (OD_600_ 0.01, to reach an average occupancy of 0.5 cells per droplet, in 20% Percoll, Sigma) and substrate solution (0.2 mg/mL polymyxin B (Sigma), 2 µl/ml R-lysozyme (Merck), 10 mM NADPH (Carbolution), 20 mM cyclohexanone (Fluka), 40 mM cyclopropylamine (from pH-adjusted stock of 500 mM cyclopropylamine (Sigma) in 100 mM Tris/HCl, pH 8.0) and HFE (Fluorochem) with 2% RAN surfactant. Flowrates were adjusted to yield droplets with a volume of approximately 170 pl. Droplets were incubated for either 3 or 22h. Picoinjection was performed at 250-500 Hz (1.0 - 2.0 V constant sine wave) with 15 mM WST-1 (NBS biologicals), 5 μg/mL mPMS (Sigma) and 6 mM Tartrazine (Sigma) in a pico-injection device (**Supplementary Figure 1, Supplementary methods**)). 85 pL were injected and flowrates at injection speed were carefully adjusted to rule out droplet fusion. Sorting was performed at 150-200 Hz using custom Arduino and LabVIEW scripts ^27^ with 4-7 V voltage. A custom-made sorting device was used (**Supplementary Figure 1, Supplementary methods**)) that was fabricated as previously described. Droplets were collected in a DNA LoBind tube (Eppendorf). The emulsion of collected droplets was broken by adding 1H,1H,2H-2H-Perfluoro-octanol in a 1:1 (v/v) ratio with HFE-7500 (3M) and 100 µl 2 ng/µL salmon sperm DNA (Thermo), and subsequent vortexing (1 min, full-speed). After centrifugation (14000xg, 1 min), the aqueous solution was carefully pipetted off to avoid transferring the oil phase and DNA was extracted three times by repeated addition of 2 ng/µL Salmon sperm DNA, vortexing and centrifugation. The resulting aqueous solution was concentrated using a spin column (Monarch, NEB) following the manufacturer’s instructions. DNA was either eluted in 6 µL nuclease-free water (Invitrogen) and transformed into electrocompetent *E. coli* (E cloni, Lucigen) for secondary screening, or eluted in 20 µl nuclease-free water for NGS.

*Droplet screens of SrIRED with substrates 1b:* The general protocol for *Sr*IRED with substrate 1a was followed but using a substrate solution with 0.2 mg/ml polymyxin B (Sigma), 2 µl/ml R-lysozyme (Merck), 10 mM NADPH (Carbolution), 20 mM cyclohexanone **(1)** (Fluka), and 40 mM methylamine **(b)** (from pH-adjusted stock of 500 mM methylamine, Sigma) in 100 mM Tris/HCl, pH 8.0 and incubation times of 22 and 92 h, respectively. The final substrate concentrations for the reaction in droplets were 10 mM **(1)** and 20 mM **(b)**.

*Droplet screens of SrIRED* with substrate ***9***: The general protocol for *Sr*IRED with substrate **1a** was followed but using a substrate solution with 0.2 mg/ml polymyxin B (Sigma), 2 µl/ml r-lysozyme (Merck), 10 mM NADPH (Carbolution), and 20 mM 2-methyl-1-pyrroline (Sigma) in 100 mM Tris/HCl, pH 8.0 (final substrate concentration of 10 mM 2-methyl-1-pyrroline) and the incubation times of 3 h and 24 h, respectively.

*Droplet screens of PcIRED* with substrates **10f**: The general protocol for *Sr*IRED with substrates **1a** was followed with the following alterations. A substrate solution with 0.2 mg/ml polymyxin B (Sigma), 2 µl/ml r-lysozyme (Merck), 1 mM EDTA (Sigma), 4 mM NADPH (Carbolution), 4 mM 3-(2-chlorophenyl)propanal **(10)** (Key Organics; from a 250 mM stock in 100 mM TrisHCL pH 8.0 with 80% DMSO), and 4 mM (1*R*)-1-(3-methoxyphenyl)ethan-1-amine hydrochloride **(f)** (Enamine, from a 500 mM pH-controlled stock in 100 mM Tris pH 8.0 with 50% DMSO) was prepared. Final conditions in droplets: 2 mM **(10)**, 4 mM **(f)**, 2 mM NADPH, 0.5 mM EDTA, 0.1 mg/ml polymyxin B, 1 µl/ml r-lysozyme, 100 mM Tris/HCL, 2.5% DMSO. Flow rates were adjusted to produce droplets with a volume of 185 pl. Droplets were incubated for either 20h or 40h. Picoinjection of 92.5 pL was performed with 6 mM WST-1 (NBS biologicals), 5 μg/mL mPMS (Sigma) and 3 mM tartrazine (Sigma).

### Cloning of UMI-tagged error-prone PCR libraries

The *Sr*IRED gene was amplified with error-prone PCR using the GeneMorph II random mutagenesis kit (Agilent) following the manufacturer’s instructions using 2.4 µg template plasmid and 25 PCR cycles. The product was subjected to DpnI digest (NEB) overnight, extracted from an agarose gel and purified again using SPRIselect beads (Beckman coulter) following the manufacturer’s instructions. The UMI was introduced from randomised oligonucleotides (IDT) and 3 cycles of PCR with a single primer annealing to its 3’-end were performed to produce double-stranded DNA (see **Supplementary Methods 7** for primer sequences). Two more fragments (vector and promoter region) were produced by PCR. Gibson assembly mix^81^ was added on ice, and the reaction was immediately transferred to be incubated at 50°C for 1h. After purification using SPRIselect beads (Beckman Coulter), plasmid DNA was transformed into electrocompetent *E. coli* (Lucigen). A plate with 60000 transformants was obtained by conducting a dilution series (overall number of colonies: 2*10^6^). The *Pc*IRED epPCR library was constructed with the same protocol but using Golden Gate assembly (**Supplementary Methods 7**).

### Oxford Nanopore sequencing

After library transformation into *E. coli*, DNA for sequencing was prepared by adding 10 ml LB media to the agar plate covered with grown cells, scraping the colonies using a sterile L-scraper, recovering the cell suspension, harvesting the cells by centrifugation (4000xg), and minipreping following the kit manufacturer’s instructions (Genejet). For the *Sr*IRED library, a restriction digest with Acl1 (NEB) and Bsu36I (NEB) was performed to yield a DNA fragment with UMI and the *Sr*IRED gene (for scheme see **Supplementary Methods 8**). The fragment was purified via agarose gel electrophoresis and SPRIselect beads. 500 fmol of DNA were sequenced on three Minion R10.4 flowcells (Oxford nanopore technologies) using the Oxford Nanopore ligation sequencing kit following the manufacturers protocol. For the *Pc*IRED library, plasmids were linearized with the restriction enzyme HpaI (NEB) (**see SI methods section 8** for a scheme) and sequenced on four Minion R10.4.1 flow cells (ligation sequencing kit, V14 with 300 fmol of SPRI-purified DNA).

### Preparation of libraires for NGS

For NGS library preparation, primers including Illumina flow-cell binding sequences and indices in overhangs were used to amplify the UMI region of the recovered DNA (**Supplementary Table 1** for SrIRED and **Supplementary Table 7** for *Pc*IRED). The different microfluidics runs were multiplexed using primer pairs with different NGS indices. Upon preparation of the PCR mix (NEBNext Q5 mix, NEB) following the instructions of the manufacturer, PCR conditions were optimized using a 15% fraction of the recovered DNA and qPCR to determine the cycle number at 10% of the maximum fluorescence. 18-22 cycles were employed for subsequent recovery. DNA was purified using SPRIselect beads (Beckman Coulter), with a beads-to-sample volume ratio of 0.7 for *Sr*IRED and 0.65 for *Pc*IRED to remove primer dimers. The DNA concentration was determined using NEBnext library quant kit (NEB) and capillary electrophoresis (Bioanalyzer, Agilent) following the manufacturers protocols. The samples were pooled and prepared into Illumina TruSeq libraries by the University of Cambridge, Department of Biochemistry sequencing facility according to the manufacturer’s instructions. Next-generation sequencing was performed using an Illumina next-seq 2×150bp run (with 20% PhiX spike in).

### Generation of dictionaries and fitness scores

#### Pipeline for SrIRED

A flow scheme of the pipeline is provided in **Supplementary figure 3.** Basecalling of Oxford Nanopore sequencing data was performed using guppy (ONT), quality control was performed using NanoPlot and filtering using NanoFilt (length 1,250 --maxlength 1,400 --quality 8). A custom python script was used to extract the UMI sequences ^28^ and the generated fasta file was clustered using the mmseqs2 suite^82^ (min-seq-id 0.8, seq-id-mode 2, cov-mode 0 -c 0.8). The files were then processed in a custom pipeline using minimap2 to align the sequences in every cluster, racon^53^ to generate a draft sequence assembly, minialign to align draft assembly and reference (with NNNN in variable regions of the UMI) and medaka (ONT) for generation of a consensus file. The vcf files generated by medaka were converted into .fasta files using custom python scripts. A variant identifier file for downstream data generation was generated by custom python scripts. The custom python scripts used in dictionary generation are available at https://github.com/Hollfelder-Lab/lrDMS-IRED/tree/main/scripts. Fitness scores were calculated from the variant identifier file and the raw NGS foward reads using the DiMSum suite^83^ with the following parameters: retainIntermediateFiles T, cutadaptMinLength 20, vsearchMinQual 10, cutadaptErrorRate 0.6, numCores 8, paired F, indels all, maxSubstitutions 100, mixedSubstitutions T, barcodeErrorRate 0.1, fitnessMinInputCountAll 0.

#### Pipeline for PcIRED

The same pipeline as for *Sr*IRED dictionary and fitness score generation was used and any alterations are outlined here. Basecalling was performed using dorado basecaller (ONT) and NanoFilt was applied with the following parameters: length 4,250 -- maxlength 5,250 --quality 15. Parameters for mmseqs2 suite were the following: min-seq-id 0.85, seq-id-mode 2, cov-mode 0 -c 0.8. Parameters for DiMSum suite were the following: retainIntermediateFiles T, cutadaptMinLength 20, vsearchMinQual 10, cutadaptErrorRate 0.6, numCores 2, paired F, indels all, maxSubstitutions 100, mixedSubstitutions T, barcodeErrorRate 0.1, fitnessMinInputCountAll 0.

### Cloning, expression and purification

IRED variants (N-terminal His_6_-tag, under control of a T7 promoter in a pRSF vector, see SI section “sequences”) were either cloned using site-directed ligation-independent mutagenesis (SLIM)^84^, Gibson assembly^81^ or using dial out PCR with primers specific to the UMI region from the library^28^. Cloning success was confirmed by Sanger sequencing. Variants were transformed into BL21(DE3) cells (NEB), a single clone was picked and grown to saturation in LB media supplemented with kanamycin (50 µg/mL) (37°C, 200rpm). A culture was diluted to an OD_600_ of 0.05, grown to an OD_600_ of 0.4-0.6 and protein expression was induced with 0.2 - 1 mM IPTG. Protein was expressed overnight (20°C, 200rpm) and cells were harvested by centrifugation (4000xg). Cells were resuspended in lysis buffer (100 mM Tris/HCl pH 8, 100 mM NaCl, 1 mM DTT, one Complete Protease Inhibitor tablet for 200 mL buffer, Roche) and lysed by sonication for 7 min (Vibracell, 50%, 1 sec on, 1 sec off,). The lysate was cleared by centrifugation (14000xg, 4 °C) and incubated with Ni-NTA agarose (NeoBiotech) on a roll mixer at 4°C for 1h. The beads were washed three times in a plastic gravity column with a frit with 10 ml wash buffer (100 mM Tris/HCl pH 8, 150 mM NaCl, 1 mM DTT, 30 mM imidazole). Protein was eluted in 10 ml elution buffer (100 mM Tris/HCl pH 8, 150 mM NaCl, 1 mM DTT, 300 mM imidazole). The protein was rebuffered into storage buffer (100 mM Tris/HCl pH 8, 150 mM NaCl, 1 mM DTT, 10% glycerol) using Amicon centrifugation filters (10,000 NMWL; Merck), shock frozen with liquid nitrogen and stored at −80°C. Enzyme concentration was determined in a Bradford assay (Sigma) and purity was controlled by SDS PAGE.

### Lysate activity assays

After transformation into chemically competent BL21(DE3) cells (NEB), colonies were picked into deep-well plates (Greiner) and filled with 1 ml LB media supplemented with kanamycin. Plates were incubated overnight (37 °C, 750 rpm). 30 µL of the culture were diluted in 920 µL of fresh LB media supplemented with kanamycin in a fresh deep-well plate and grown for 2h (37 °C, 750 rpm). Protein expression was induced with 0.2 mM IPTG, and plates were incubated overnight (20-25 °C, 750 rpm). Cells were harvested by centrifugation (30 min, 4000g), the supernatant was removed, and cells were resuspended in 200 µL lysis buffer (100 mM Tris pH 8, 4 mg/ml egg white lysozyme, 0.5 mM EDTA, 1 mg/ml polymyxin B) per well. After incubation (25°C, 1h, 750 rpm), lysate was cleared by centrifugation (30 min, 4000g, 4°C). Assays were performed in Nunc 96-well plates (Thermo). To determine the initial rate of the reaction, the reaction was induced by adding 10 µL lysate to 90 µl reaction buffer (10 mM carbonyl compound, 20 mM amine, 0.5 mM NADPH in 100 mM Tris pH 8) and the absorption at λ of 340 nm was followed in a plate reader (Spectramax 190, Molecular Devices). The initial rate in µM/s was calculated using a calibration curve of known NADPH concentrations at 340 nm. To determine the conversion of the reaction in an assay similar to the droplet assay, cleared lysate was used to carry out a reaction with 5 mM NADPH, 10 mM cyclohexanone and 20 mM cyclopropylamine in 100 mM Tris/HCl pH 8.0. After 3h and 22h incubation at 25°C, 5 µl of the reaction mix were added to 95 µl 5 mM Iodonitrotetrazolium chloride which is a cheaper alternative to WST-1 (INT, Sigma) and 5 µg/ml mPMS in DMSO. The reaction was diluted 1:20 and the absorbance was measured in a spectrophotometer in Nunc 96-well plates (ThermoFisher). In analogy to the calculation of a fitness score, the natural logarithm of the average fold-change of INT absorbance over wild type (proportional to the remaining NADPH concentration) after 3 and 22h was calculated.

### Michaelis Menten kinetics for *Sr*IRED and homologs

Kinetic constants of *Sr*IRED for cyclohexanone were determined in 100 mM Tris pH 8.0 with 0.5 mM NADPH and varying concentrations of cyclohexanone (0.06 mM - 35 mM) in transparent Nunc 96-well plates (ThermoFisher) and by monitoring the depletion of NADPH concentration at 340 nm with 30 mM cyclopropylamine and 0.001 - 0.01 mg/mL enzyme at T = 25 °C. Initial rates were measured in 3 replicates and curves were fitted using Prism.

### Michaelis Menten kinetics for *Pc*IRED

Michaelis Menten curves were measured by following the decline of NADPH absorbance at 340 nm with 0.5 mM NADPH and 20 mM amine **f** in 100 mM Tris/HCl pH 8, 50% DMSO with varying aldehyde **10** concentrations (between 10 and 0.014 mM) in three technical replicates (T=25 °C). Standard errors of the fit and three technical replicates are represented with a 95% confidence interval.

### Soluble expression and thermal shift assays

Soluble expression of variants was determined by SDS PAGE analysis of the pellet and the supernatant by densitometry. Variants were expressed analogous to expression for the lysate assay and the lysate was cleared by centrifugation. The pellet was resuspended in 200 µl Laemmli buffer (NuPAGE, invitrogen) and samples of the supernatant were collected; SDS PAGE analysis was conducted and band intensity was compared using ImageJ. For thermal shift assays, 10 µM enzyme with 2x SyproOrange (invitrogen) in 100 mM Tris/HCl pH 8 were analysed in a Bio-Rad CFX Connect device.

### Heuristics for generation of a mutational profile

To guide rational engineering, we define two metrics – mutability and combinability – reflecting how well individual positions accept mutations and how well mutations can be combined into improved higher-order variants. Our discussion is restricted to substitution mutations only. We use (*p*, *a*) to denote the substitution of position *p* in the wild type sequence to the amino acid *a*. A variant *v* is then defined as a collection of *K* substitutions 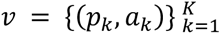, where each position may occur at most once. The mutation order |*v*| is the number of mutations *K* in variant *v*. For example, the single mutant T241A would be denoted {(241, *A*)}. The double mutant T241A P203A would be {(203, *A*), (241*A*)}, and so on. Given these definitions, we define mutability of a position *p*_0_ as

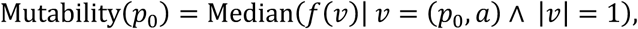

where *f*(*v*) is the logarithmic lrDMS fitness score vs wildtype. In words, the mutatability of position *p*_0_ is the median fitness over all single mutants in the dataset that occur at position *p*_0_.

Further, we define combinability of position *p*_0_ as the empirical combination

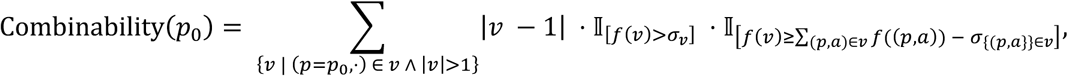

with 𝕀 the indicator function, which is 1 if the condition is fulfilled and 0 otherwise. I.e. the combinability of position *p*_0_ is the weighted sum over all higher-order mutants, which contain a mutation at *p*_0_ and have confidently positive and confidently non-negatively epistatic fitness *f*(*v*). *Confidence* here is calibrated against the standard deviation *σ*_*v*_ of the fitness value of variant *v* across the replica of the experiment as well as the uncorrelated estimate of the standard deviation 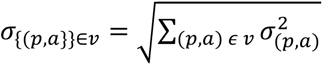 of the additive fitness of the single mutants (*p*, *a*) that make up the variant *v* across the replicas. The weighting is given by the mutation order |*v*|, excluding the mutation at the position itself (|*v* − 1|). Note that this definition can be extended naturally to a combinability score for individual mutations (*p*, *a*) and not just for positions *p*.

A combined plot of the *mutability* and *combinability* heuristics for rational engineering along the *Sr*IRED sequence can be found in **Supplementary Figure 23**.

For the *Pc*IRED campaign, less sequence function data was available, so the definition of combinability was relaxed to the following equation:

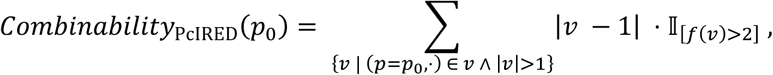

A combined plot for the heuristics with *Pc*IRED can be found in **Supplementary Figure 36**.

### Machine learning modelling

The single and double mutant AI predictions come from two simple models which were trained on the obtained lrDMS fitness data. The single mutant model is based on a ridge-regression model using fixed, biochemically informed amino acid embeddings and pre-trained language model likelihood while the double mutant model is a gradient boosted tree model on a few curated features deemed important for predicting the fitness of double mutants and tailored to the nature of lrDMS data. We chose these simple models for their relative robustness and interpretability, allowing us to better understand the value of the lrDMS data and to better mitigate the effects of noise in the data, which we expected to be larger than data from plate-based assays on which many current models are evaluated. We note that in the future the predictive performance could be further enhanced by utilizing more complex models which leverage pre-trained information from sequence, structure, or related assay data more extensively, provided they are paired with approaches to reduce model overfitting.

### Single mutant machine learning model

The single mutant model is a ridge regression model, which is augmented with an evolutionary density from a large language model as proposed in Hsu et al.^50^, to predict fitness scores for unobserved single mutants. For an amino acid sequence *x* = (*x*_,1_*x*_2,_ ⋅⋅⋅, *x*_*L*_) we parameterize the fitness as

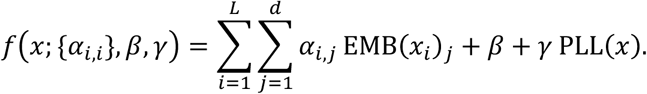

The embedding EMB: {Ala, Arg, …, Tyr, Val} → ℝ^*d*^ for a given amino acid is computed from a principal component analysis of the AAINDEX database^47^, which contains reference data on over 500 physiochemical and biochemical properties, such as e.g. hydrophobicity, for each natural amino acid. As in Gelman et al.^48^ we reduce the dimensionality via principal component analysis to a 19-dimensional feature vector (*d* = 19) for each amino acid, still capturing over 98% of the variance. In comparison to the one-hot encoding used by Hsu et al.^50^, this chemically informed embedding better captures the similarity between amino acids. This chemical similarity between amino acids is crucial to help the model extrapolate the effects of unseen mutations at positions: The top prediction T241G for example substitutes glycine (G) instead of alanine (A) as in T241A, which is chemically similar but smaller than alanine.

We further include evolutionary information by adding the pseudo-log-likelihood (PLL) of a pre-trained large language model. Based on its successful use in prior work^50,85^ we use the ESM2 masked language model^49^. The PLL for masked language models,

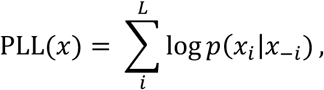

where *x*_–*i*_ denotes masking at position *i*, requires *L* function evaluations, making it expensive to compute for many variant sequences *x*. To reduce the computational cost, we instead use the PLL difference versus the wildtype sequence, which only differs from PLL(*x*) by a constant and can further be approximated as^50,86^.

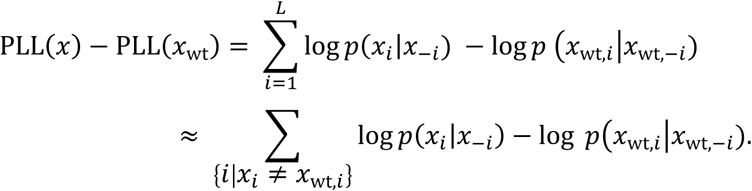

This approximation assumes that the conditional log-likelihood of an amino acid at any position is almost the same regardless of whether the wild type or mutant background are given, an assumption which we observed holds well when the mutation order is low for a random subsample of mutants. We used the ESM2 3B parameter model in our experiments.

The ‘augmented’ ridge-regression model *f* is then fitted to the assay-labelled single-mutant data using 5-fold cross validation to tune the *L*^2^ regression hyperparameter *λ*. Regularization was only applied to the amino acid features, but not the PLL component of the model. To compute the final predictions, the model is re-fitted on the entire single-mutant data using the hyperparameters obtained from cross-validation. A detailed evaluation of the single mutant model performance can be found in **Extended Data Figure 7** and **Supplementary Figure 26**.

### Double mutant machine learning model

The double mutant model is a gradient boosted tree model (xgboost^87^) on a selected group of features. The features can be grouped into four categories:

(1) *Assay derived features:* These include the (1) fitness of the least fit mutation in the double mutant, as well the (2) additive fitness of both single mutations assuming no epistatic effects. This data can be derived directly from the dataset or extrapolated from a single mutant model when the single mutations in question were not observed. Finally, we also include combinability information (3). For historic reasons, the combinability information used in the double mutant model differs slightly from the *combinability* heuristic used for rational engineering. In the double mutant model, two combinability features, defined as (using the same notation as above)

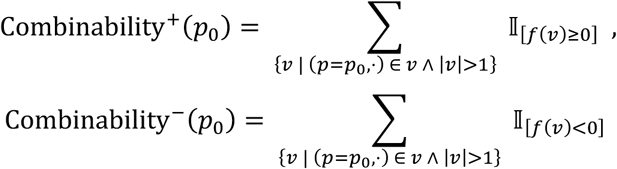 are used. Compared to the rational engineering heuristic, the weighting by mutation order |*v*| and the indicator function for better-than-additive mutations are removed and we instead define a positive or negative combinability which count the number of times a mutation at the given position *p*_0_ was observed in a better/worse than wildtype variant. The combinability-type information benefits particularly from the epPCR library generation step in the lrDMS workflow, as this step tends to generate a *stacked* library where higher order mutants are based off generated lower order mutants.
(2) *Structural information:* To give the model a rough idea of the structural context of a given double mutation we provide (1) the minimum distance of the singles that comprise the double mutant to the NADPH binding pocket, (2) the minimum distance of said singles to the dimer interface and (3) the proposed catalytic site residues. We also include (4) the pairwise distance between the Cα atoms of the two mutants. A profile of the structural features along the *Sr*IRED sequence is shown in **Supplementary Figure 31** and of *Pc*IRED in **Supplementary Figure 41**.
(3) *Coarse grained dynamics information:* We use a coarse-grained normal mode analysis using and invariant elastic network model on the wild type structure (`5OCM`) to derive the mean squared expected fluctuation, a value that mimics b-factors, of each position. We use provide the mean squared fluctuation of both single mutants (min & max) as a feature. A profile of the mean squared residue fluctuation along the *Sr*IRED/*Pc*IRED sequence is shown in **Supplementary Figure 31 / 41**.
(4) *Language model information:* Finally, we include sequence and evolutionary information through (1) the pseudo-log likelihood (PLL) of the ESM2 language model as well as an ESM2 ‘epistasis’ feature (2) which is defined as the difference between ESM2 PLL for the double mutant versus the sum of the PLL predictions for the single mutants. We use the same PLL approximations described above for the single mutant model and cache intermediate results where possible, to reduce the computational burden of these features when computing them for a library. As demonstrated in the ablations in **Extended Data Figure 8** and in the top double mutant predictions when removing language model features (**Supplementary Figure 18**), the model can perform well even without these more computationally expensive language model features.

We performed feature selection and hyperparameter tuning of the double-mutant model using cross validation. Our final double-mutant model is composed of 100 estimators, each of maximum depth 3, and was obtained by training with a squared error regression objective, although we also explored training with ranking and multiclass objectives. Detailed learning curves and a cross-validation evaluation of the final model can be found in **Extended Data Figure 8** and **Supplementary Figure 27**. To obtain predictions for assaying in the lab we re-trained the models on all higher order lrDMS data with the fixed hyperparameters found during tuning and predicted the fitness of all 721,870 double mutants that could be composed by combining single mutants observed in lrDMS. We chose to restrict ourselves to double mutants obtained from single mutants to increase robustness given our low experimental budget (5 assayed mutants per model). Consequently, T241G, the best *predicted* single mutant for example, was not within the scope of double mutant predictions. However, this choice can easily be relaxed, and the model can take extrapolated single mutant predictions as input to build higher order mutants. To understand how the various features influenced the final double mutant model’s predictions we investigated model internal feature importance indicators (gain) and performed a SHAP analysis, detailed in **Extended Data Figure 9** and **Supplementary Figures 24 and 25**.

#### Comparison of zero-shot models

Besides ESM, Tranception^88^ is another well established, zero-shot language model trained on evolutionary data that has been evaluated for protein fitness prediction^25^. In **Supplementary Figures 28, 29 and 30** we show that ESM and Tranception’s predictions are highly correlated (Spearman: 0.84) and have essentially the same performance on our dataset. We therefore proceeded to use ESM as a representative of this class of models.

## Extended data figures

**Extended Data Figure 1:**
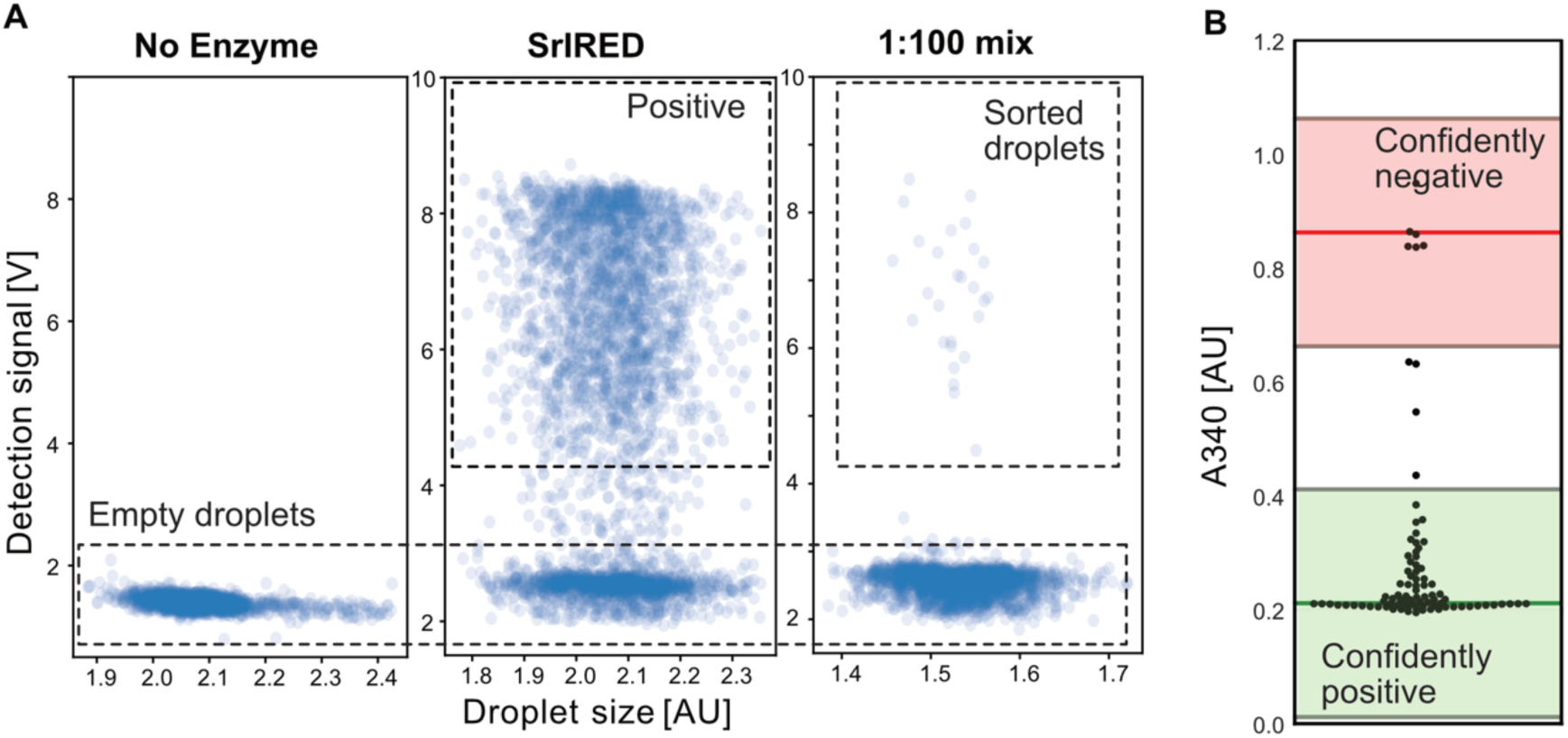
An ultra-high throughput assay for screening imine reductases. A mock-sorting experiment with an IRED from *Streptosporangium roseum* (*Sr*IRED in pRSF vector) diluted with *E. coli* expressing no IRED (pRSF vector without insert) shows the capability of the assay to enrich IRED activity substantially (93-fold) over background. (**A**) Droplets with cells not expressing enzyme, cells expressing *Sr*IRED and a 1 to 100 dilution of cells expressing *Sr*IRED with cells expressing no enzyme in the AADS assay. The population with high detection signal represents droplets with IRED-expressing cells. The second population with a low detection signal represents empty droplets (due to single-cell encapsulation of a diluted cell suspension 60% of droplets contain no cell as a consequence of Poisson distribution) and droplets with non-expressing cells. **(B)** For an enrichment experiment, the positive population was sorted from a 1 to 100 dilution of cells expressing *Sr*IRED and cells not expressing enzyme. The plasmid DNA was recovered by transformation and a secondary assay was performed with 83 clones. The reductive amination activity was determined in lysate with 0.5 mM NADPH, 10 mM cyclohexanone and 20 mM cyclopropylamine. An endpoint was measured after 3 h at 340 nm. Positive and negative clones are assigned using an interval of 0.2 from the positive and negative controls (green and red lines). 6 negative and 77 positive clones were found, giving an enrichment factor of 93.

**Extended Data Figure 2:**
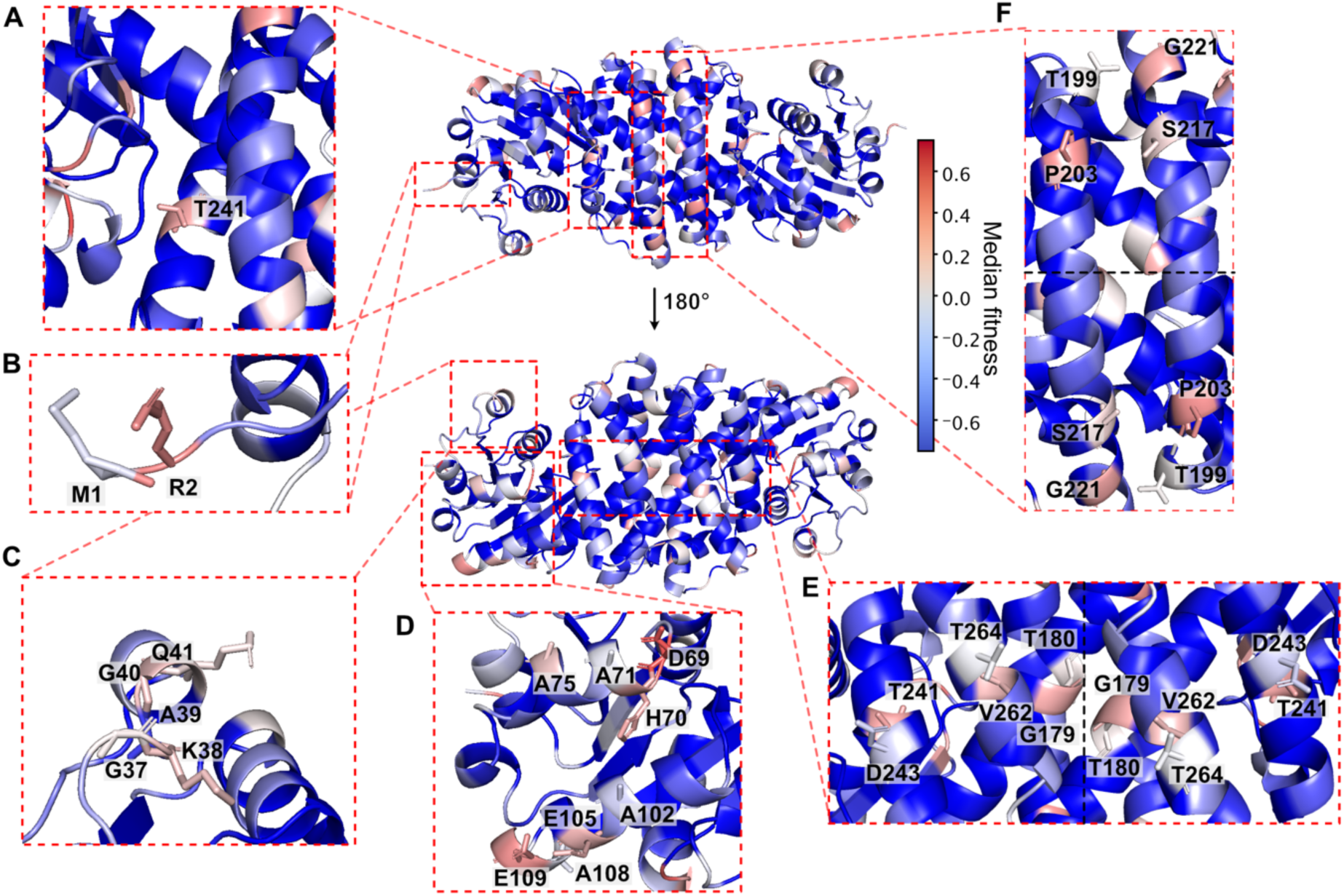
Structural mapping of mutability hotspots. **(A)** T241 is the only residue in the first shell at which mutations show improving median fitness. (**B**) Mutations at the two N-terminal residues M1 and R2 show high average fitness values indicating a high potential for improvement. M1 can be mutated without a high fitness penalty due to missed translation as a His_6_-tag with a second start codon is located at the N-terminus. (**C**) A loop and the beginning of a helix at the edge of the cleft forming the active site shows a pattern of highly mutable positions interspaced by an alanine with low mutability pointing to the inside of the N-terminal domain. The loop is involved in NADPH binding e.g. via K38 interacting with the NADPH phosphate. **(D**) Two helices in the N-terminal domains show high mutability especially at residues pointing to the outside of the protein. (**E**) Patches of residues on a horizontal axis in the central helices show high mutability (the pattern duplicated because of the dimeric structure of the protein as indicated by the black dotted line). Similar to the patches shown in D, a role of the mutations in orientating the central helices at the dimer interface of the protein can be hypothesized. (**F**) In two helices in the central domain two patches of four positions with high mutability were identified (the pattern duplicated because of the dimeric structure of the protein as indicated by the black dotted line). It is conceivable that mutations at these positions play a similar role, e.g. in orientating the central helices at the dimer interface could be hypothesized.

**Extended Data Figure 3:**
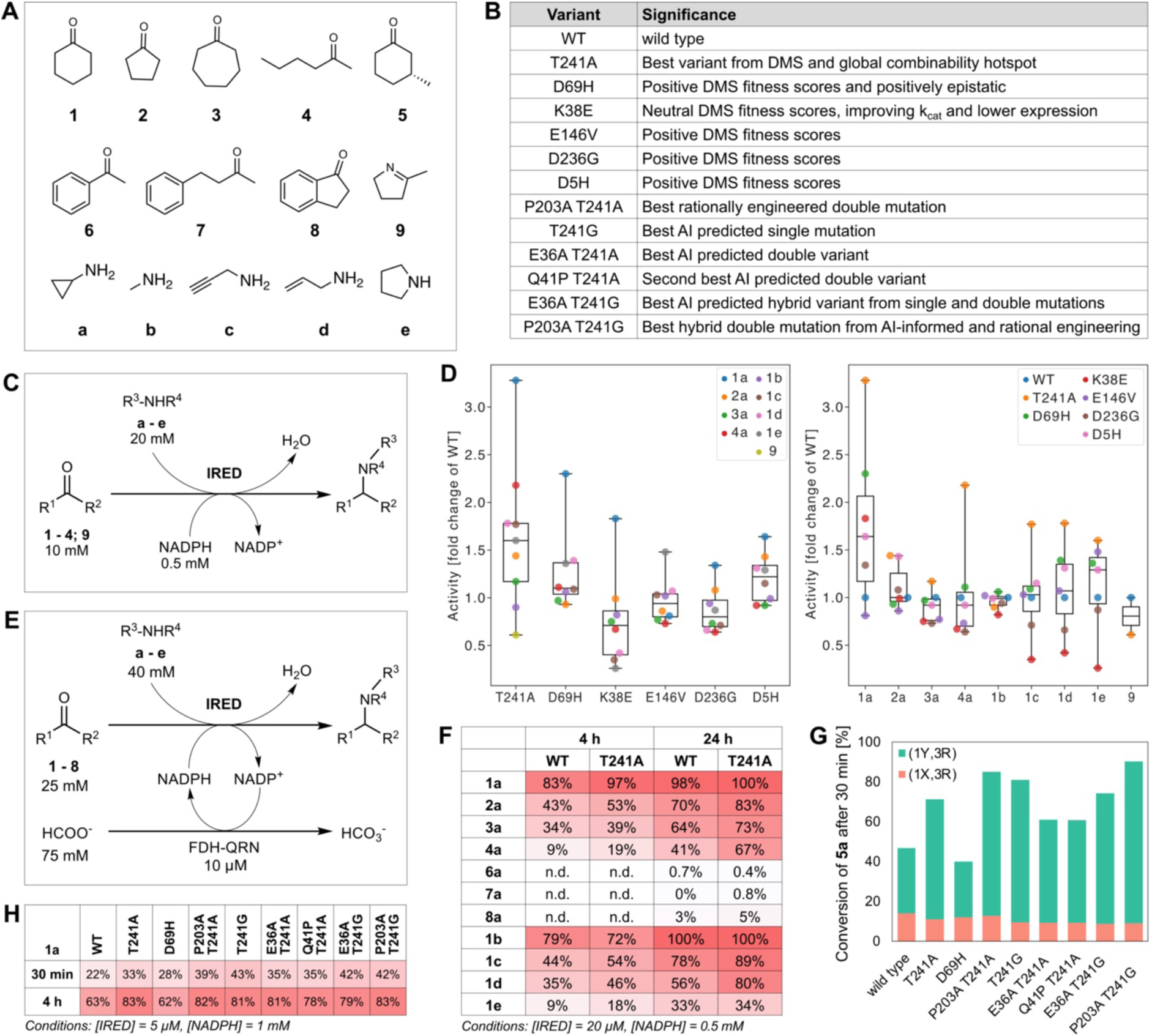
Portability of features of the mutagenic profile of lrDMS to other substrates improves application of *Sr*IRED in biotransformations. **(A)** *Sr*IRED wild type and selected mutants were tested with a panel of ketones and amines, as well as an imine, with varying degrees of similarity to the model substrate combination (ketone parameters: ring size, structure, aromaticity, bulkiness, reactivity; amine parameters: size, secondary/primary, bulkiness at the reaction center, influence of a π-system on reactivity). (**B**) Criteria for choosing variants for characterization of substrate scopes. **(C)** The single variants were characterized by using 0.5 mM NADPH and following the progress of the reaction by monitoring the depletion of NADPH absorbance at 340 nm (**Supplementary methods**). **(D)** Fold-change in initial rate compared to wild type with six chosen mutants and a set of substrate combinations (color-coded) with ketone (10 mM) and amine (20 mM) or imine only (10 mM), and NADPH (0.5 mM) in Tris-HCl buffer (100 mM pH 8.0). Absolute activity values are listed in **Supplementary Figure 9**. T241A increases activity for a broad range of different ketones and amines, showing a 3-fold activity improvement with **1a**, but lower increases with smaller and larger rings. K38E is strictly specialized to **1a** and shows decreasing or unchanged rate with all other substrates. D69H shows significant improvements only with the model substrates **1a** but slight improvements with **1e** and **1d.** E146V mainly improves lysate activity by increasing the expression level (**Supplementary Figure 8 and Table S2**) and a slight decrease in *K*_M_ (1.5-fold) but also shows slight activity improvements with **1e** (1.5-fold) that go beyond expression and binding. The relative changes in initial rate (compared to wild type) for all substrate combinations shows that T241A is the best variant for all substrate combinations apart from **1b** and **9** and the only variant significantly improving activity with 2-hexanone (**4a)**. Plotting the activity improvements for different mutants for all substrates (right) reveals significant changes in median activity (one standard deviation > 1) only for (**1a)** showing that the discovered mutations overall specialize for the substrate combination we screened with. With methylamine (**1b**) and 2-methyl-1-pyrroline (**9**) mutations are at best neutral: even the mutability and combinability hotspot T241A is not improving, leading to the conclusion of low adaptability of *Sr*IRED for these substrates. Interestingly, with pyrrolidine (1**e**) E146V and D69H are rivalling the improvement with T241A which might represent alternative potential trajectories in adaptation of SrIRED towards these substrates. (**E**) Reaction conditions used for monitoring conversion during biotransformations, quantified by GC. A variant of the formate dehydrogenase from *Candida boidinii* from (FDH-QRN) was used as a recycling-enzyme. Sodium formate served as the ultimate hydride donor (**Supplementary Methods**). (**F**) *Sr*IRED wild type and T241A were tested in biocatalytic reactions for the amination of ketone substrates **1** to **4** and **6** to **8** with amine donors **a** to **e**. T241A improves conversions with all tested substrates apart from **1b** and **6a**. Raw values with standard deviations are listed in **Supplementary Figure 11**. **(G)** The top variants’ conversion with the chiral substrate (*R*)-3-methyl cyclohexanone with nonchiral amine **a** and the stereoselectivity at the newly formed stereogenic centre (green and red). It shows that T241A and T241G significantly improve conversion on their own and combined with other mutations (up to 90% conversion with a 90:10 diastereomeric ratio compared to a 70:30 diastereomeric ratio for wild type with P203A T241A). All T241A and T241G mutants only accelerate the reaction towards one diastereomer with conversion to the other diastereomer staying nearly constant or even decreasing. Raw values with standard deviations are listed in **Supplementary Figure 22. (H)** Conversions of model substrate **1a** in biotransformations. The top variants show improvements that are reflected in the kinetic parameters as well as in conversion values under reaction conditions relevant for biocatalytic application. Raw values with standard deviations are listed in **Supplementary Figure 21**.

**Extended Data Figure 4:**
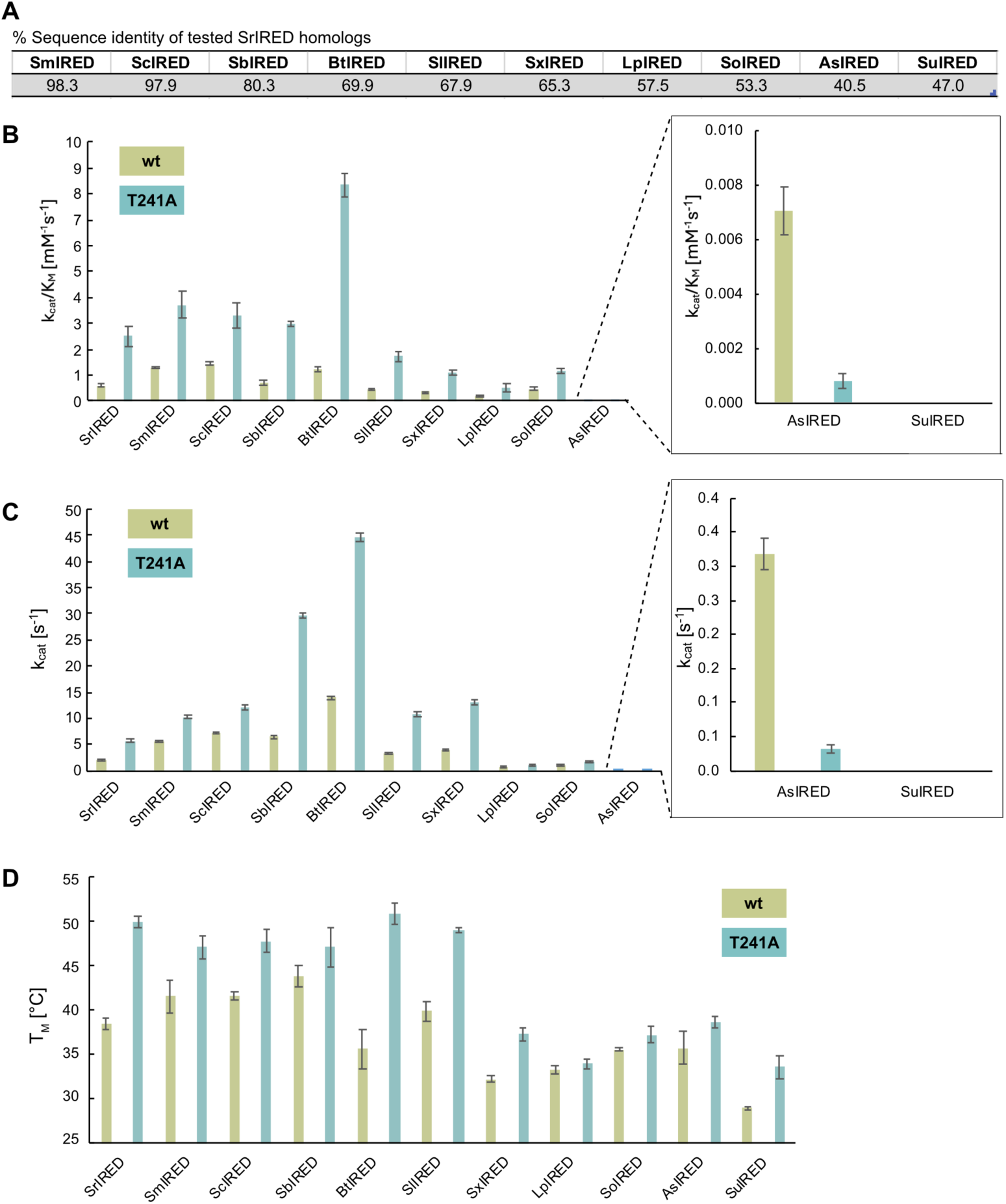
T241A-mediated improvements in activity and melting temperature are transferable to *Sr*IRED homologs. **(A)** Percent sequence identity of *Sr*IRED homologs **(B)** k_cat_/K_M_ values of *Sr*IRED homologs tested with substrate pair **1a** (wild type: green, T241A: blue). Improvements in k_cat_/K_M_ are transferable for IREDs with sequence identities as low as 53% (*So*IRED). For *As*IRED (sequence identity: 41%), no improvement but a drop in k_cat_/K_M_ was observed. No activity was observed with *Su*IRED. Michaelis-Menten kinetics were conducted with variable concentrations of ketone substrate **1** (0.05 - 35 mM) and a constant concentration of amine **a** (30 mM) in Tris/HCl pH 8 with NADPH (0.5 mM) and variable enzyme concentrations (from 0.03 µM to 0.3 µM, adjusted to the assay timescale) at T = 25 °C. Raw data in **Supplementary Table 4. (C)** A similar pattern of improvement as a function of sequence identity was observed with k_cat_ values. **Supplementary Table 4. (D)** The introduction of T241A leads to improvements in melting temperature T_M_ across all tested IREDs, the highest improvement of 15 <u>°</u>C being observed for *Bt*IRED. Even IREDs that do not show an improvement in activity upon introduction of T241A (*As*IRED) and IREDs with no measurable activity with **1a** showed improvements in melting temperature, indicating that the improvements in stability conferred by T241A are more readily transferred than activity improvements. Melting temperatures were measured by thermal shift assays (see example curves in **Supplementary Methods Section 9, data in Supplementary Table 4**). Error bars show standard deviations of three technical replicates.

**Extended Data Figure 5:**
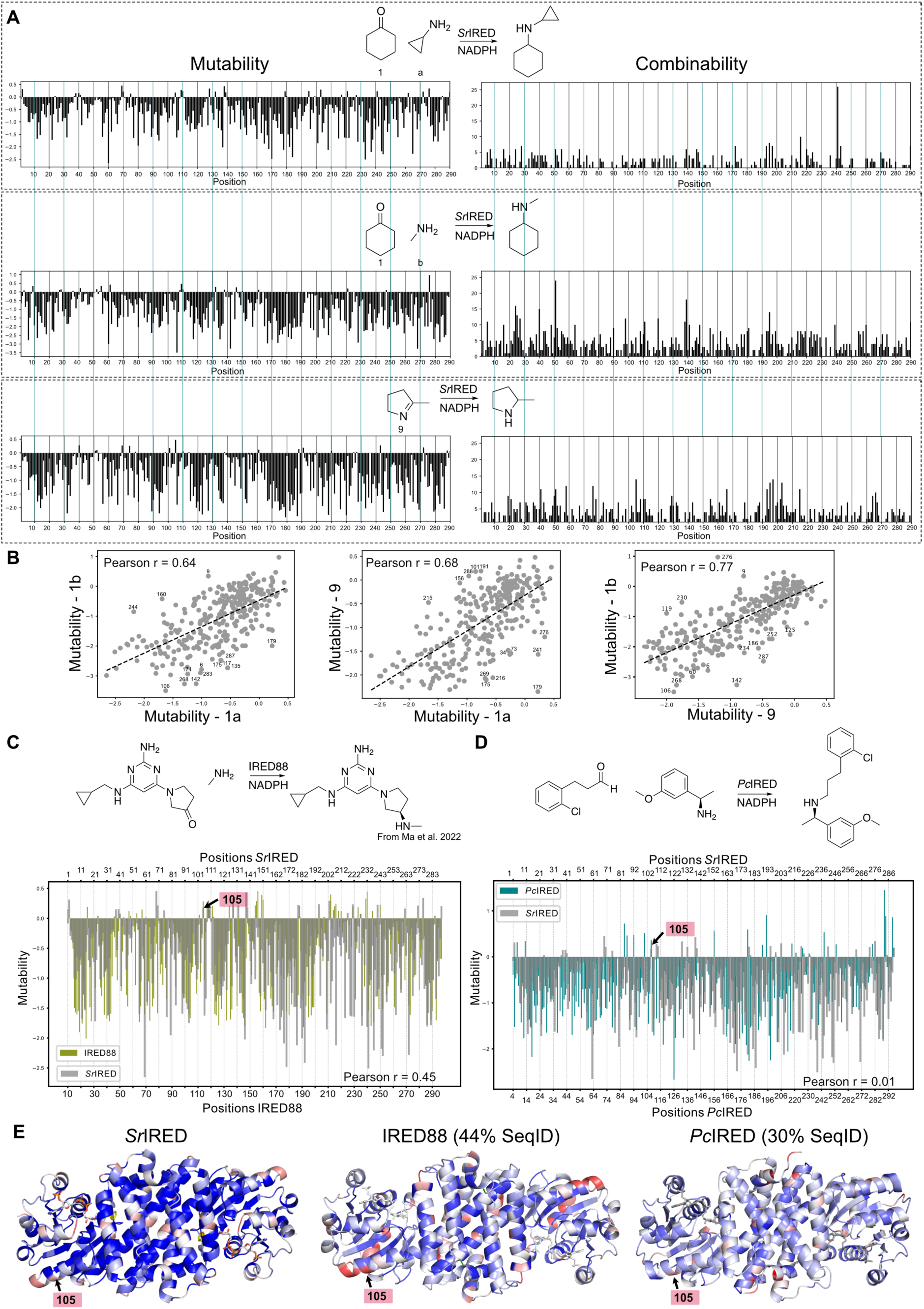
*Sr*IRED mutability is transferable to different substrates and IREDs. **(A)** Mutability and combinability profiles of *Sr*IRED for the deep mutational scanning with three different substrates: cyclopropylamine (**a**) and cyclohexanone (**1**), methylamine (**b**) and cyclohexanone (**1**), and 2-methyl-1-pyroline (**9**). While mutability correlates well, combinability does not. T241 is only a combinability hotspot with **1a** (Combinability correlations in **Supplementary Figure 17**). Deep mutational scanning with **1b** reveals a combinability hotspot at position 51, while no clear combinability hotspot could be observed with **9**. Structural insight into the specificity-conferring residues can be found in **Supplementary Figure 16**. **(B)** *Mutability correlates well between the three deep mutational scans*. Residues that are significantly more mutable with one substrate than the other (“specificity conferring residues”) are labeled. One example is T241, which is improving with **1a** (but not with **1b** and **9)**. **1b** and **9** are also among the only substrates which were not improving for T241A when tested experimentally (**Extended Data Figure 3**). **(C)** *Overlay of mutability between SrIRED and IRED88* (44% sequence identity) from a deep mutational scan conducted by Ma *et al.*^45^ shows significant overlay of the mutability profile with an overall correlation with a Pearson r of 0.45 **(Supplementary Figure 15**). This shows that mutability information can also be transferred to other IREDs for engineering activity with different substrates **(D)** *Comparison of the deep mutational scan* with *Pc*IRED (30% sequence identity) and *Sr*IRED shows no global correlation of mutability (**Supplementary Figure 15**). However, there are still similarities in mutability in the N-terminal domain that might guide engineering: The mutability hotspot at position 105 (*SrI*RED numbering) is conserved across all three IREDs with different substrates **(Supplementary Figure 15)**. The corresponding E107K mutation in *Pc*IRED shows a 1.8-fold improvement in K_M_ (**Supplementary Table 6**). (E) *Structural mutability profiles* of *Sr*IRED, *Pc*IRED and IRED88 shows the shared mutability hotspot at position 105 on the surface of a helix in the N-terminal domain.

**Extended Data Figure 6:**
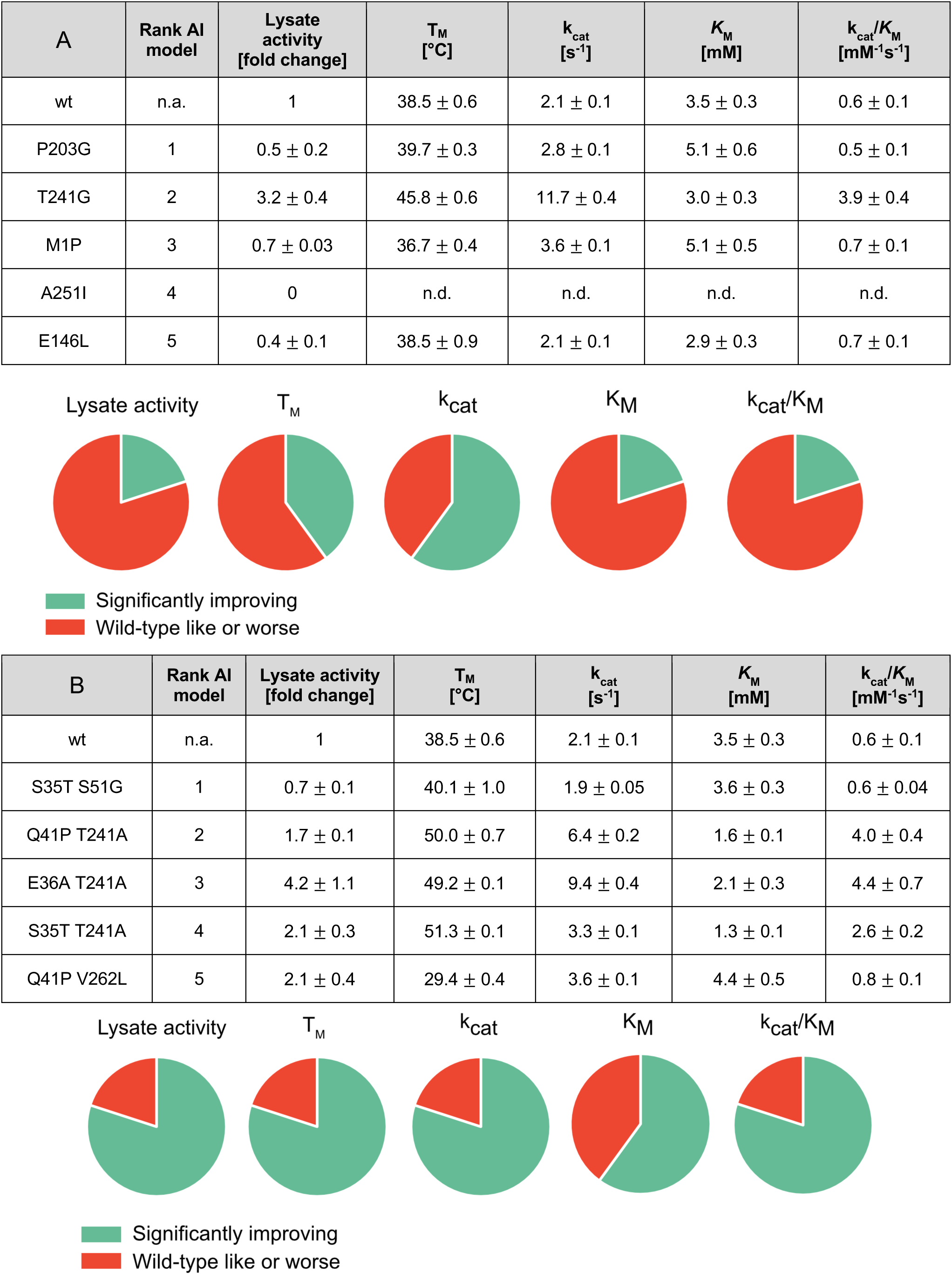
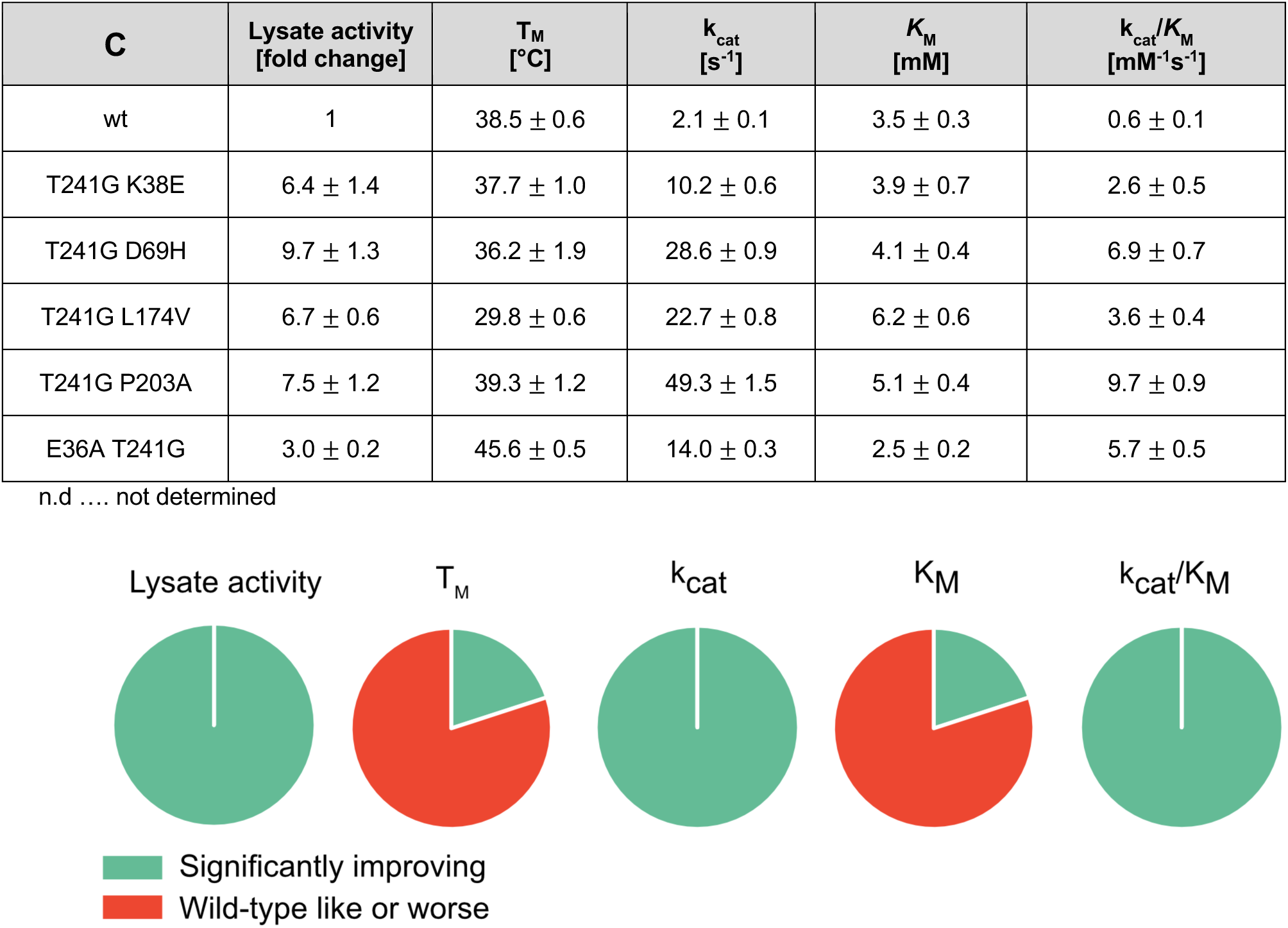
Characterization of AI-predicted and hybrid variants. Characterization of top-ranked predictions of the single mutant AI model (**A**), double mutant AI model **(B),** and transplantation of AI-generated mutants into rationally engineered variants **(C)**. In total, 12 out of 15 tested variants improve k_cat_, giving a 80% success rate. The top variant T241G P203A (bold), shows a 23-fold improved k_cat_ with a 16-fold improved k_cat_/K_M_ and is also the best variant tested in biotransformations under conditions relevant for application (Extended Data Figure 3G and H). Lysate activity was measured as initial rates in three replicates (errors represent standard deviations) with **1** (10 mM), **2** (20 mM), NADPH (0.5 mM) in 100 mM Tris pH 8.0 following the decline of NADPH absorbance at 340 nm and normalized to lysate without IRED. Soluble expression was determined by densitometry of SDS-PAGE bands of the soluble and insoluble fraction after lysis. Values are normalized to wild type. Melting temperatures were measured in thermal shift assays in three replicates, errors represent standard deviations. Michaelis Menten kinetics were conducted with variable concentrations of ketone substrate (0.05 - 35 mM) and a constant concentration of amine (30 mM) in Tris pH 8 (T = 25°C) with NADPH (0.5 mM) and variable enzyme concentrations (from 0.03 µM to 0.3 µM) depending on the rate. Each curve was measured in three replicates. Errors represent standard errors of the fit and three technical replicates with a 95% confidence interval.

**Extended Data Figure 7:**
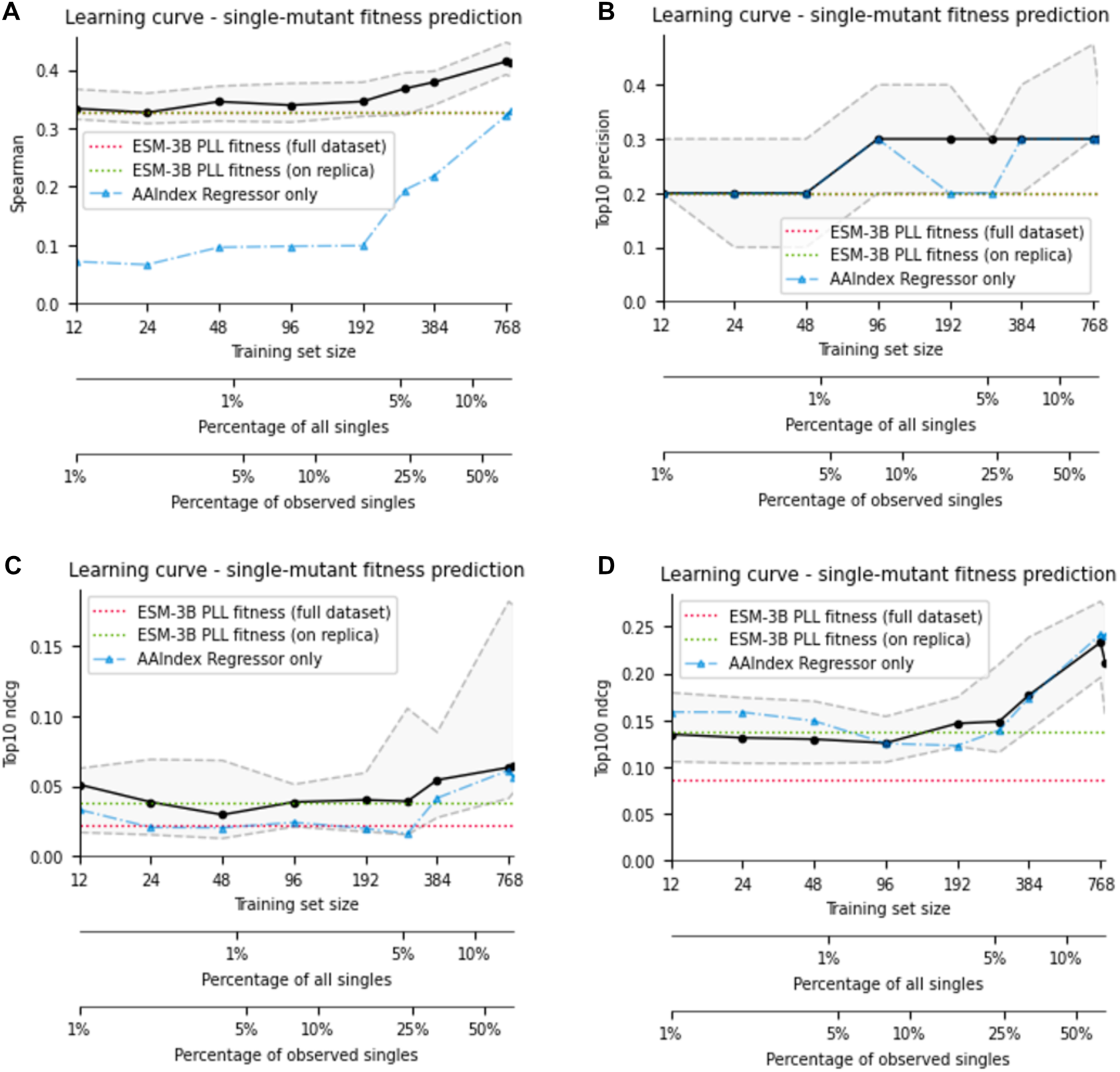
Single mutant model learning curves for four key metrics: Spearman, Top10 precision and NDCG and Top 100 NDCG. Learning curves for the single mutant model and several ablations are computed on 10 replicas, using 3-fold cross validation with a different random seed per replica, resulting in 30 experiments. The black line indicates the single mutant model’s median across all experiments, while the top/bottom dashed gray line indicates the 75^th^ and 25^th^ percentile – corresponding to a ‘good’ and ‘bad’ case respectively. The blue line ablates the ESM features and shows the median performance of the assay-trained linear regression model based on AAIndex embeddings alone. The red and green lines respectively indicate ESM PLL zero-shot performance on all single mutants and the median across the 30 folds used to test the other models, for fair comparison in the presence of batch effects. **(A)** *Spearman correlation.* For small dataset sizes (<200 fitness scores) the model’s ranking performance is dominated by the performance of ESM, as expected: by construction the single mutant model treats each position independently and can therefore not generalize to unseen positions. From dataset sizes above ∼200 (*Sr*IRED is 290 AA long) the model has on average observed 2 amino acids (wild type and a mutant) per position and the droplet data starts to clearly improve global ranking performance. The overall Spearman coefficient (∼0.41) is modest, but not uncommon for noisy biological datasets^24,46^. Note that while Spearman is a useful measure of overall model performance, it poorly reflects the performance on the few top mutants, which are most relevant for engineering. **(B)** *Top 10 precision*. The model’s hit-rate for discerning variants better than wild type (WT) (fitness > 0) among the top 10 variants does not significantly improve with increasing lrDMS training dataset size apart from the uptick of the 25^th^ percentile. On average, 2-3 of the top 10 predicted variants are consistently better than wildtype, which is comparable to the hit-rate of ESM-PLL ranking (2). This is likely due to the limited resolution of lrDMS data for variants of similar activity, and because most single mutants are of WT-like fitness (c.f. Figure 2B, single-mutants are strongly peaked around WT-fitness compared to higher order mutants). Instead, lrDMS data contains a more coarse-grained signal for when a variant is significantly better or worse than WT. As a result, if the test data for a given replica only contains mediocre fitness variants, the model is unlikely to be able to say with confidence if this variant is slightly better or worse than WT, limiting precision improvements. (**C-D)** *Top 10/Top 100 normalised discounted cumulative gain (NDCG).* The NDCG computes the quality of recommendations by encompassing both their relevance (fitness) and their rank in the results (ranking). The restriction to the top 10/top 100 predictions further focuses the NDCG on the top few predictions which are most relevant for protein engineering at low test budgets. Similarly to the Spearman correlation, for small training dataset sizes below the sequence length (<290) the Top 10/100 NDCG are mostly constant. For training dataset sizes higher than the sequence length, NDCG starts to improve. Note that not only the median, but also the 25^th^ percentile (“bad luck scenario”) exceeds the both zero-shot baselines. Overall, the NDCG values clearly show improvements for larger dataset sizes. Similar magnitudes for NDCG are not uncommon for noisy biological datasets, and the low NDCG value achieved by ESM points to the single mutant data in this dataset being a more challenging than most others in ProteinGym^25^. Further detailed cross-validation analysis of the single mutant model is available in

**Extended Data Figure 8:**
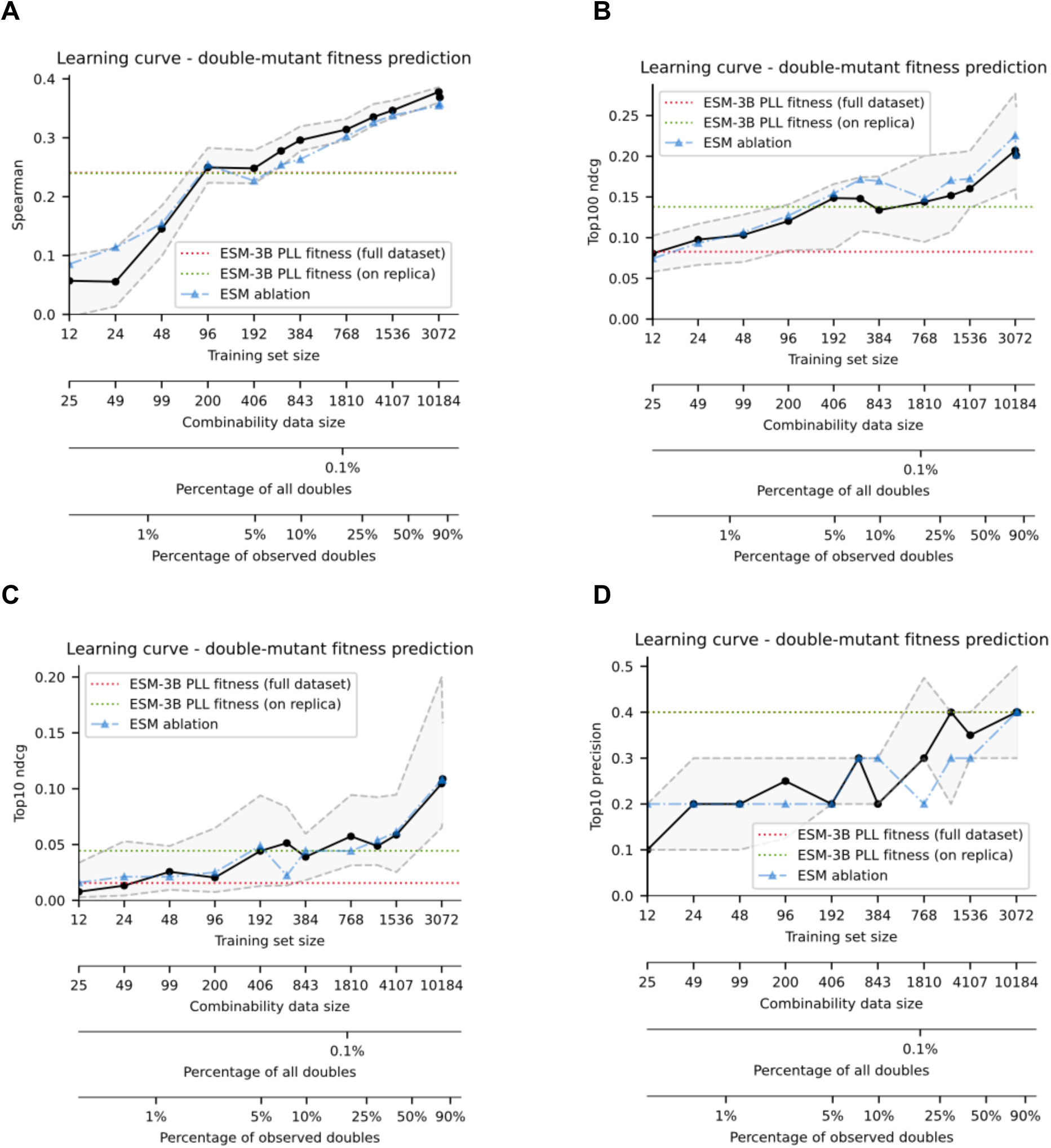
Double mutant model learning curves for 4 key metrics: Spearman, Top10 precision and NDCG and Top 100 NDCG. Learning curves for the double mutant model and several ablations were computed on 6 replicas using 5-fold cross validation with a different random seed per replica, resulting in 30 overall experiments. The black line indicates the double mutant model’s median performance across all experiments, while the top/bottom dashed gray line indicates the 75^th^ and 25^th^ percentile – corresponding to a ‘good’ and ‘bad’ case respectively. The blue line ablates the ESM features, which are the most expensive to compute, and demonstrate that our model is performant even when omitting ESM features, e.g. when no GPUs but only a table-top computer is available. It also demonstrates that most useful information is carried by the lrDMS data itself, rather than the language model. The red and green lines respectively indicate ESM PLL zero-shot performance on all single mutants and the median across the 30 folds used to test the other models, for fair comparison in the presence of batch effects. The two x-axes (“Training set size” & “Combinability data size”) correspond to the number of double mutants available for training the model and the number of higher order mutants available to inform combinability features respectively. **(A)** Spearman correlation. The model’s ranking performance consistently increases with increasing dataset size. Notably, even the 25^th^ percentile outperforms the zero-shot baselines as soon as about 300 datapoints are available. **(B-C)** Top 100 and Top 10 Normalized discounted cumulative gain (NDCG). The NDCG computes the quality of recommendations by encompassing both their relevance (fitness) and their rank in the results (ranking). The restriction to the top 10/top 100 predictions further focuses the NDCG on the top few predictions which are most relevant for protein engineering at low test budgets. **(D)** Top 10 precision. The model’s hit-rate for discerning variants better than wild type (WT) (fitness > 0) among the top 10 variants also shows a consistent uptick as dataset size is increased. It is the only metric for which the “bad case” (25^th^ percentile) is not consistently better than the zero-shot ESM performance. Similarly to the single mutant case, we attribute this behaviour to the limited resolution of lrDMS assay data for variants of similar fitness but note that the double mutant case is less severe than in the single mutant case. For all metrics we observe significant improvements in the quality of the model with increasing amount of training data, emphasizing why ultrahigh throughput assays are crucial in making sensible predictions, and observe that ablating ESM features appears not to affect performance. Further detailed cross-validation analysis of the double mutant model is available in **Supplementary Figure S27**.

**Extended Data Figure 9:**
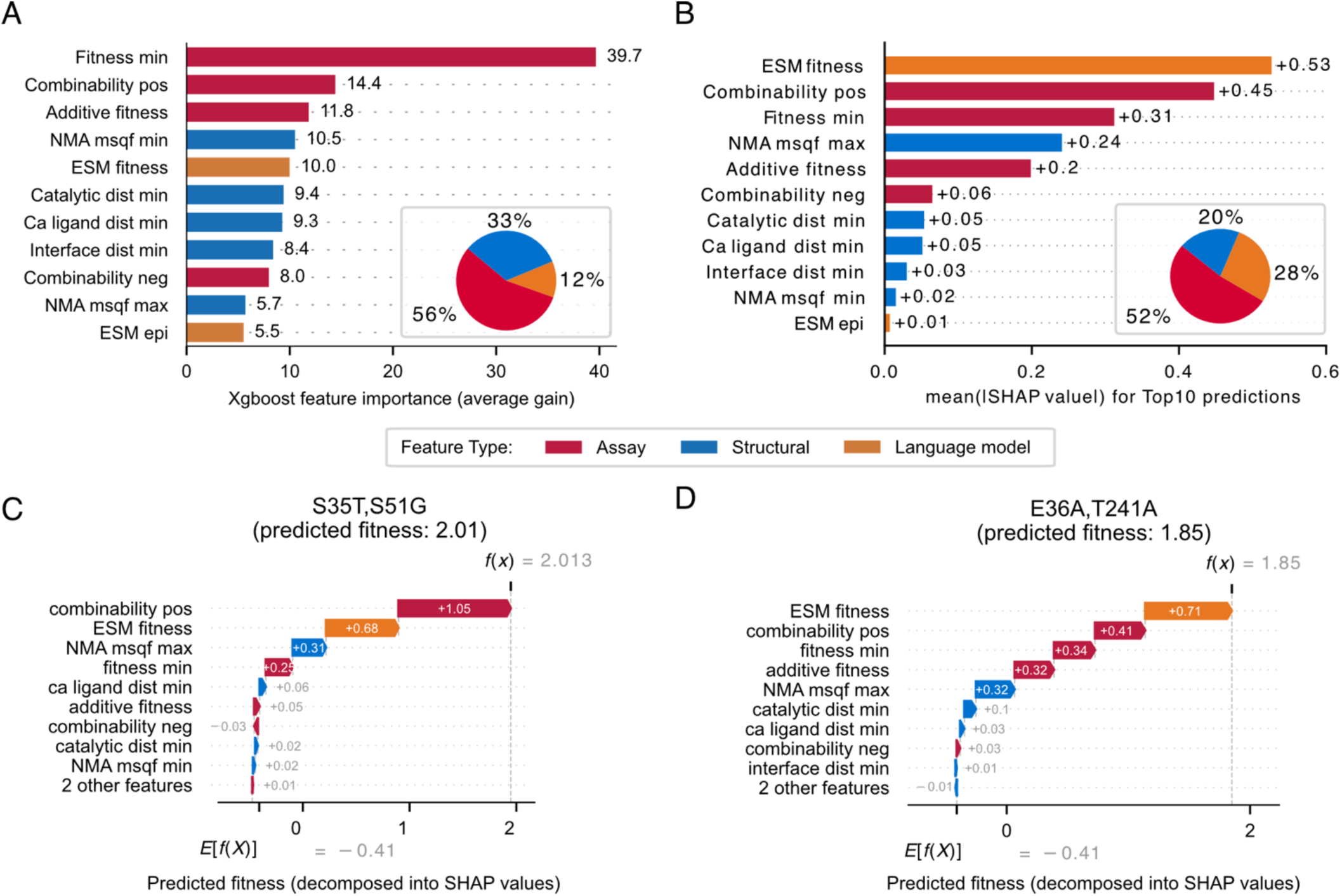
Evaluation of the influence of different features in the double mutant model. Irrespective of which feature attribution method is used, the assay derived features consistently have the highest influence on the model’s predictions. We investigate the extent to which different features influence the double mutant model that was used for hit combination with AI. (**A**) The features are ranked by their average gain^89^ across all splits in which that feature is used within the decision trees constructed by xgboost. Loosely, this gain corresponds to the extent of improvement in training error when the training data is split at a threshold value of the given feature. The so-derived ‘gain’-feature importance values indicate that assay derived features (fitness min, combinability and additive fitness) have the strongest impact for enhancing model performance during training. (**B**) To assess each feature’s the impact on our top predictions, we plot the absolute SHAP values^52^ for each feature, averaged across the top 10 predicted mutants. SHAP values are a well-studied, alternative way of assigning feature importances based on cooperative game theory and provide insight into the contribution of each feature to the prediction for an individual instance. As in (A), assay derived features are responsible the largest contribution to the predicted scores of the top10 mutants, followed by language model features and structural features. (**C**) and (**D**) show a detailed breakdown for double mutant predicted to have the highest score (C) and the model-predicted double mutant with the highest improvement in post-hoc characterization (D). Breakdowns for the other predicted variants and a more detailed SHAP analysis are available in **Supplementary Figures 24 and 25**. Note that Combinability, NMA, Catalytic site distance and Ligand distance features only influence the position of the chosen mutant. The amino acid identity is purely informed via the fitness min, additive fitness, and ESM fitness features, therefore at least one of these three features must give significant input to select the amino acid identity at the selected position, which may overweight the importance of these features over the position-only features.

**Extended Data Figure 10:**
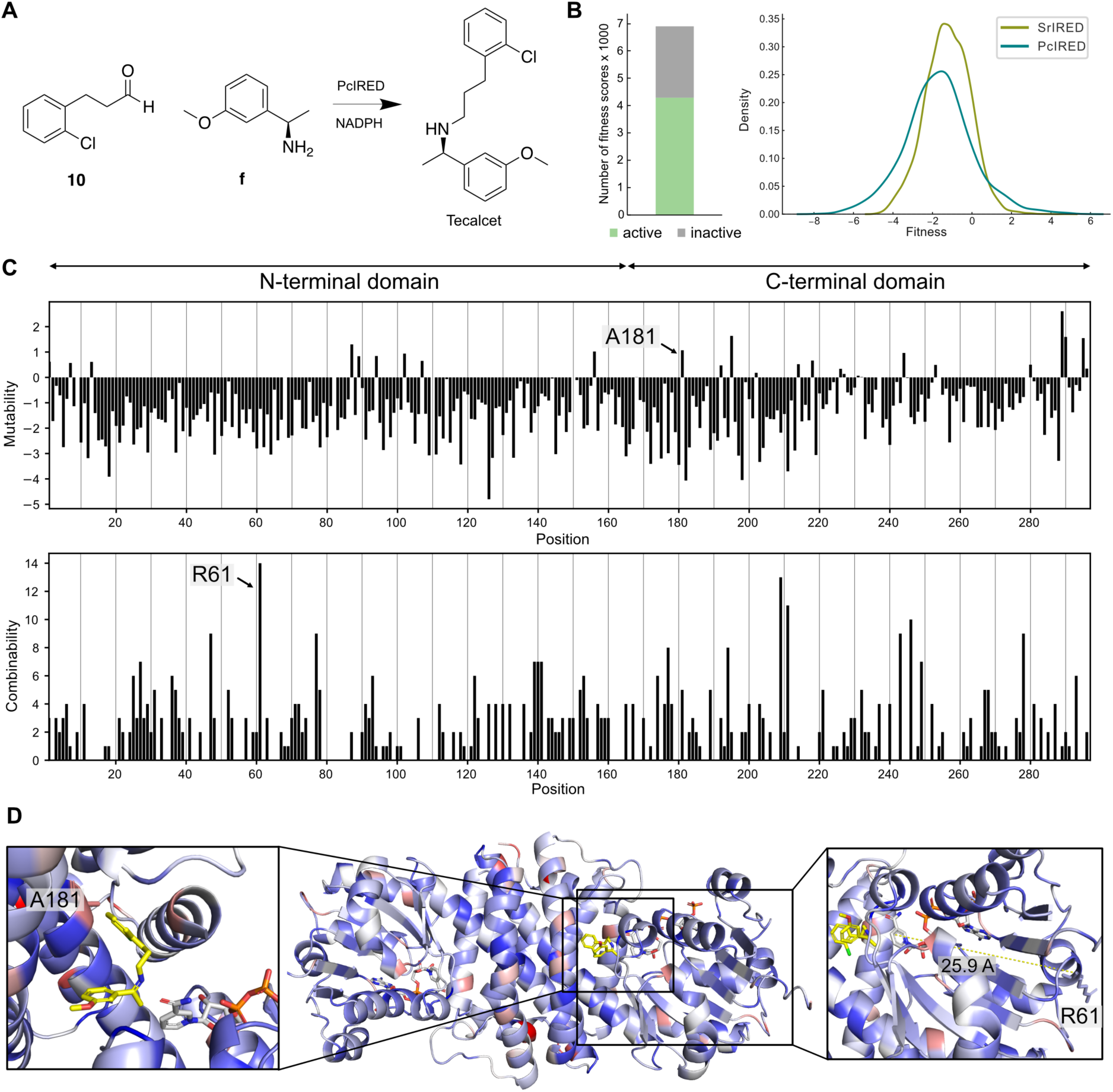
Mutability and combinability profile of *PcI*RED for the synthesis of Tecalcet. **(A)** Reaction scheme of Tecalcet synthesis from aldehyde **10** and amine **f (B)** Distribution of fitness scores. Fitness information is available for 6897 variants (compared to 17143 for *SrI*RED). For 4286 variants detectable activity was observed (62%, a similar proportion of 63% with *Sr*IRED), while 2611 were inactive. The activity distribution of *Pc*IRED is shifted to lower fitness values with an average fitness of −1.7 compared to −1.2 in *SrI*RED. **(C)** The PcIRED mutability and combinability profile. There are overall less data available for *Pc*IRED than for *SrI*RED. For mutability, therefore, on average only 2.5 fitness scores per position were used for mutability calculation. The definition of combinability was slightly adjusted to be more robust towards smaller datasets (see **Methods**). Overall mutability in *Pc*IRED is reduced to −1.3 from −0.6 for *SrI*RED. However, hotspots still emerge and are especially abundant in the C-terminal domain. While in *SrI*RED, the N-terminal domain was more mutable than the C-terminal domain (−0.5 vs −0.7), the opposite effect is observed in *Pc*IRED with higher mutability in the C-terminal (−1.1) than in the N-terminal domain (−1.4). While no global hotspot comparable to T241 for *Sr*IRED could be observed for *Pc*IRED, there are still positions standing out with higher combinability than the rest of the sequence (e.g. R61). **(D)** Mutability mapped on the *Pc*IRED Alphafold3 model docked with the Tecalcet imine. The mutability hotspot at position A181 in the first shell and the combinability hotspot R61 (located 25.9 Å from the active site) are highlighted.

**Extended Data Figure 11:**
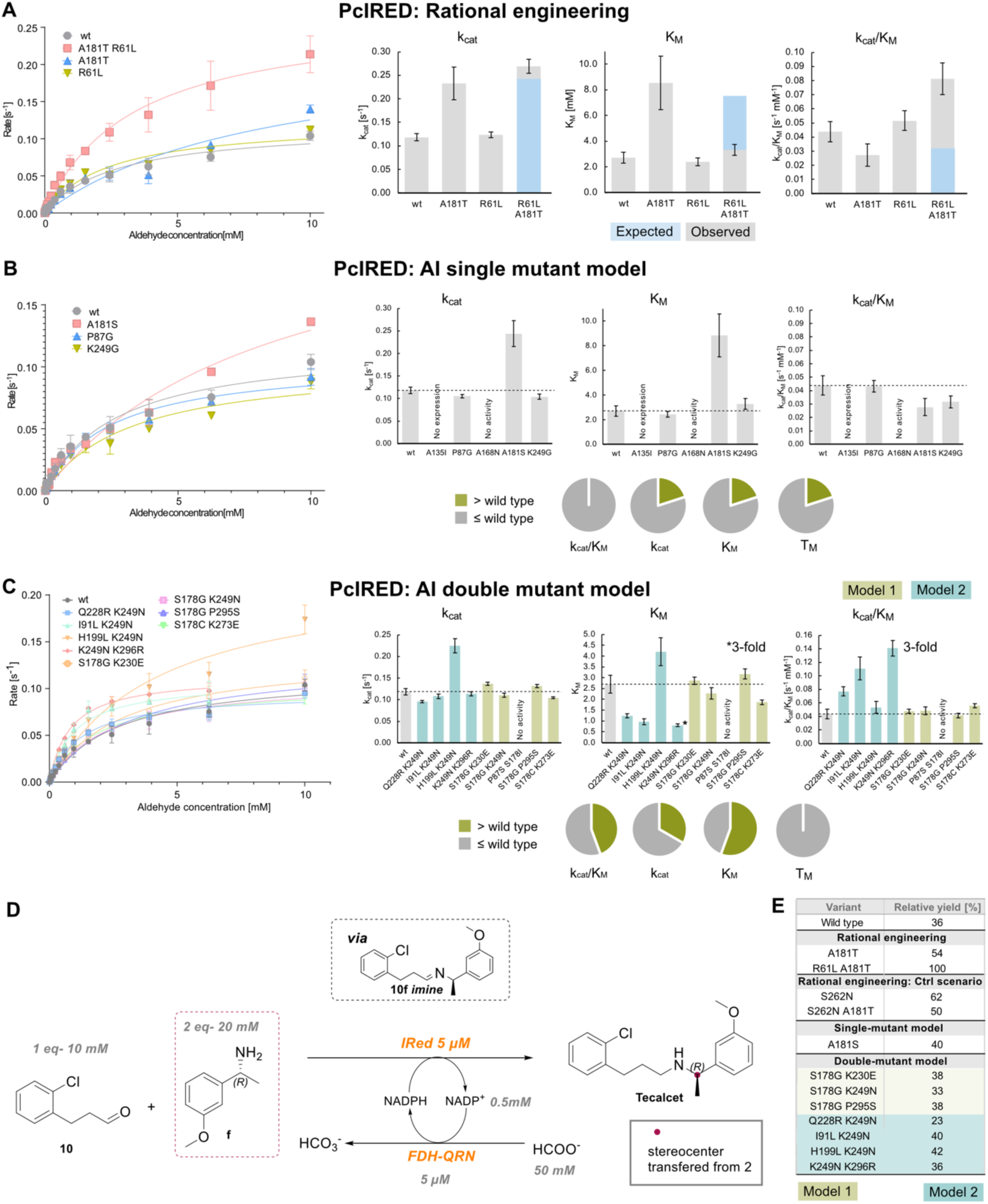
Engineering *Pc*IRED using the lrDMS profile and AI tools. **(A)** *Rational engineering using mutability and combinability information*. R61L from the position with the highest combinability was combined with A181T from a mutability hotspot in the first shell of interaction. The combination shows strong positive epistasis for K_M_ and k_cat_/K_M_ (2.3-fold and 2.5-fold lower than expected, respectively). This is especially remarkable given the long distance between R61L and the active site and its interaction partner: R61 is 25.9 Å apart from the active site (i.e. the site of hydride transfer) and 37.7 Å apart from A181, defying active-site focused protein engineering strategies. Raw data in **Supplementary Table 6**. **(B)** *The AI single mutant model was applied to PcIRED*. Due to the small single mutant dataset (on average 2.5 fitness scores per residue) the design space was restricted to residues with at least 3 fitness scores and with non-zero count in at least three sorting replicates. Moreover, mutations to proline were excluded and we limited ourselves to characterizing one mutation per site (ranked by ESM if predicted fitness is very close). The model performed significantly worse than in the case of *Sr*IRED, with only one hit experimentally verified: A181S (compared to three k_cat_-improving variants for *SrI*RED). A181S shows a 2-fold improvement in k_cat_ which comes at the expense of a 3-fold increase in K_M_. Raw data in **Supplementary Table 6**. (**C**) Given the sparser higher-order mutant data for *Pc*IRED to increase our chances of success we characterised mutants from two slightly different double mutant AI model runs: with different weights on mutation order and buffer around additive fitness (For details see **Supplementary Figure 39**). Criteria similar to the single mutant model case were applied for choosing mutants: at least three sorting replicas per position and exclusion of mutations to proline. The model was much more successful in predicting hits than the single mutant model: 3 out of 9 tested variants are improving in kcat with an up to 2-fold improvement for H199L K249N. The model performed best at predicting mutations with improved K_M_: 5 out of 9 mutations showed significant improvement with an up to 3-fold improvement for K249N K296R that also carried over to catalytic efficiency k_cat_/K_M_. Raw data in **Supplementary Table 6**. **(D)** Biotransformations were conducted with selected mutants under the following conditions: 10 mM aldehyde **10**, 20 mM amine **f** 5 µM IRED. A variant of the formate dehydrogenase (FDH-QRN) from *Candida boidini w*as used as a recycling-enzyme for the NADPH coenzyme. Sodium formate served as the ultimate hydride donor (**Supplementary Methods**). **(E)** Yields in biotransformations (in %) after 1h incubation relative to the highest-intensity peak (R61L A181T) show a 3-fold improvement in relative yield for the rationally engineered variant R61L A181T. Conversely, combining A181T with another single mutant with low combinability S262N that shows excellent yields (62%) results in a negatively epistatic interaction: S262N A181T gives only 50% conversion, whereas 93% would have been expected.

