## Supplementary material for "Microdroplet screening rapidly profiles a biocatalyst to enable its AI-assisted engineering": Combined SI

#### Supporting information

Maximilian Gantz[1], Simon V. Mathis[2#], Friederike E. H. Nintzel[1#], Matthew Penner[1], Paul J. Zurek[1], Tanja Knaus[3], Vasilis Tseliou[3], Elie Patel[1], Daniel Boros[1], Friedrich-Maximilian Weberling[1], Matthew R. A. Kenneth[1], Oskar J. Klein[1], Elliot J. Medcalf[1], Jacob Moss[2], Michael Herger[1], Tomasz S. Kaminski[1,4], Francesco G. Mutti[3], Pietro Lio[2], Florian Hollfelder[1*]

[1]Department of Biochemistry, University of Cambridge, 80 Tennis Court Road, Cambridge, CB2 1GA, UK

[2]Department of Computer Science, University of Cambridge, 15 JJ Thomson Avenue Cambridge CB3 0FD, UK

[3]Van´t Hoff Institute for Molecular Sciences, HIMS-Biocat, University of Amsterdam, Science Park 904, 1098 XH Amsterdam, Netherlands

[4]Department of Molecular Biology, Institute of Biochemistry, Faculty of Biology, University of Warsaw, Miecznikowa 1, 02-096 Warsaw, Poland

[#]Equal contribution

**Figure S30: Correlation analysis between Tranception log likelihoods and ESM pseudo log likelihood (PLL) vs wild type (WT). The scatterplot between ESM[4] and Tranception[5] fitness values (PLL for ESM, sequence log likelihood for Tranception) shows that ESM and Tranception fitness predictions are highly correlated, indicating that both models likely capture the similar aspects of the underlying evolutionary patterns from related sequences. ...45**

**Figure S39 Feature importance analysis for the double mutant *model 1* in the PcIRED campaign. While assay derived features are still attributed about one third of impact on the model's predictions, the PcIRED model relies more heavily (but not exclusively, about ~33% each) on a priori evolutionary (ESM) and structural biases than was the case *Sr*IRED (compare Extended Data Figure 9). This is consistent with what would be expect based on the learning curves for *Sr*IRED in Extended Data Figure 8, which, for a dataset of this size (490 mutation-stacked double mutants), suggest that evolutionary features (*ESM*) hold similar predictive power to the assay-derived features. For more details and analysis of these trends see Supplementary Figure 38. ....56**

#### Supplementary Figures

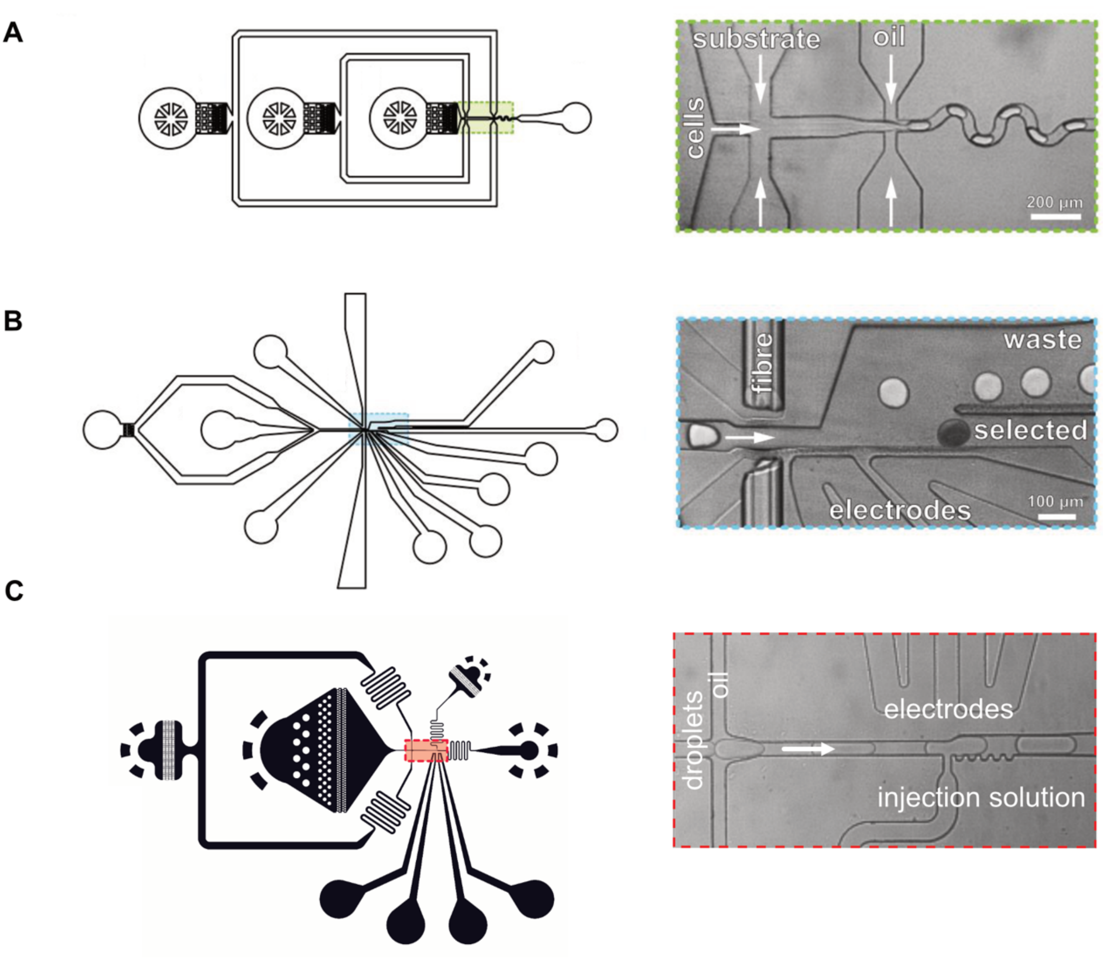

**Figure S1: Microfluidic chip designs.** All designs are available as CAD files from our repository DropBase (https://openwetware.org/wiki/DropBase:Devices). (**A**) Flow-focusing device for droplet generation https://openwetware.org/wiki/DropBase:droplet_generation_3_inlets (**B**) Sorting device https://openwetware.org/wiki/DropBase:AADS **(C)** Picoinjection device https://openwetware.org/wiki/DropBase:Picoinjection_chip

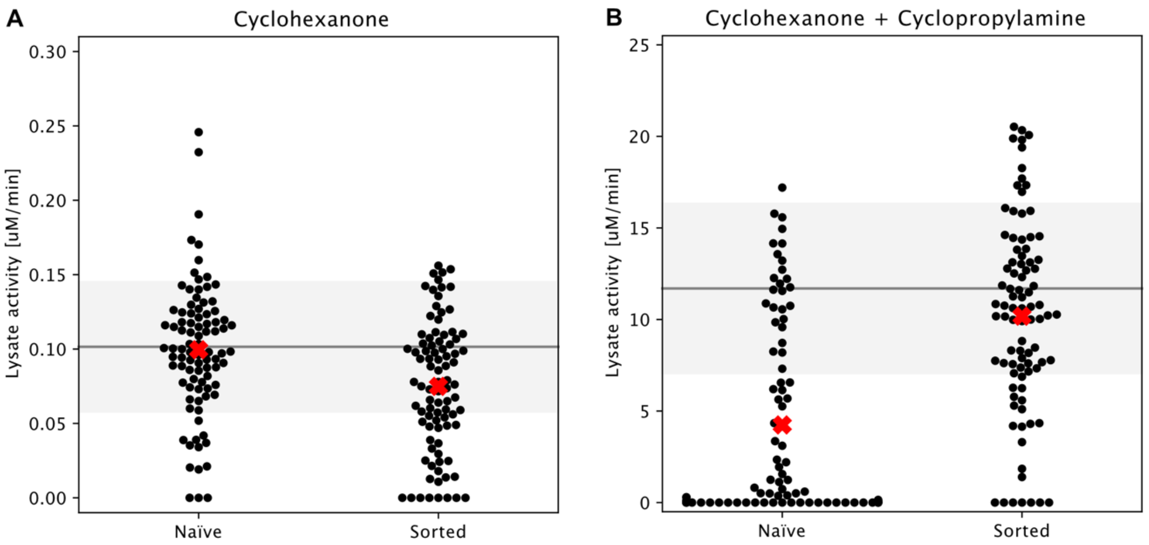

**Figure S2: Enrichment after absorbance-activated droplet sorting (AADS).** Secondary assays were performed for the naïve library (before sorting) and for the sorted library (after AADS, recovery, and retransformation). Both libraries were analysed (**A**) for enrichment of ketoreductase activity using 10 mM cyclohexanone as sole substrate and (**B**) for enrichment of imine reductase activity using 10 mM cyclohexanone and 20 mM cyclopropylamine as substrates. For each library, 90 random variants were grown in 96-well plates and expressed for 18 hours at 20 °C in duplicates. The cells were lysed and cleared lysate was recovered by centrifugation. Activities were determined as the initial rate of NADPH depletion at 340 nm in a spectrophotometer (SpectraMax 190, Molecular Devices). Reactions were carried out with 0.5 mM NADPH in 100 mM Tris-HCl buffer at pH 8. Any NADPH background depletion was subtracted, and the lysate activity was calculated using the Beer-Lambert law. The average activity of the wild type (duplicates ± standard deviation) is shown in grey, and the average activity of the whole population is marked with a red cross. The decrease in average ketoreductase activity after sorting **(A)** suggests no enrichment during AADS, while the increase in imine reductase activity after sorting **(B)** is consistent with a successful selection that enriches clones with higher activity.

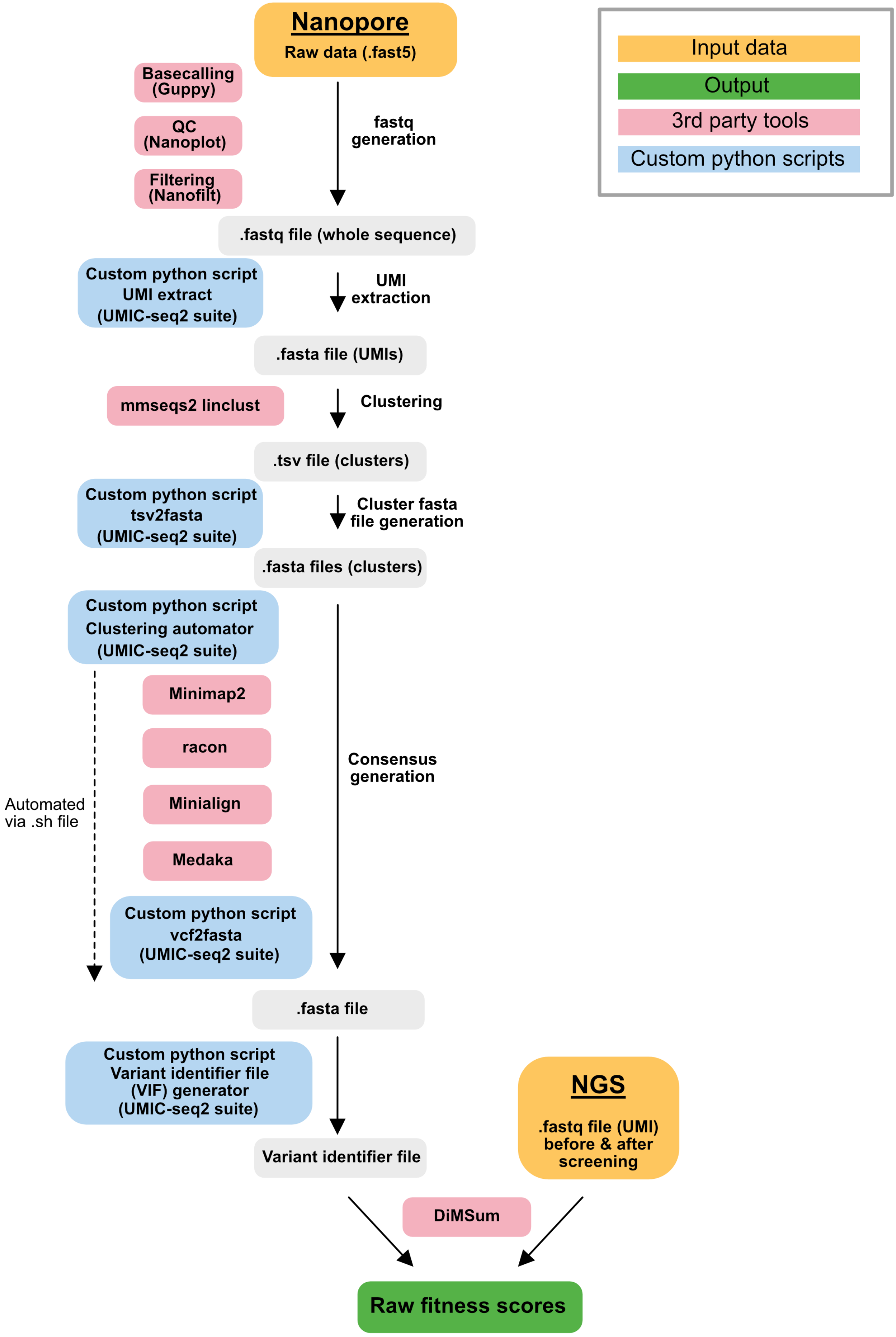

**Figure S3: Bioinformatic workflow for fitness data generation.** Workflow illustrating the generation of the dictionary (here "variant identifier file" for direct use by the DiMSum pipeline) and fitness scores. Custom python scripts are available via https://github.com/fhlab/IRED.

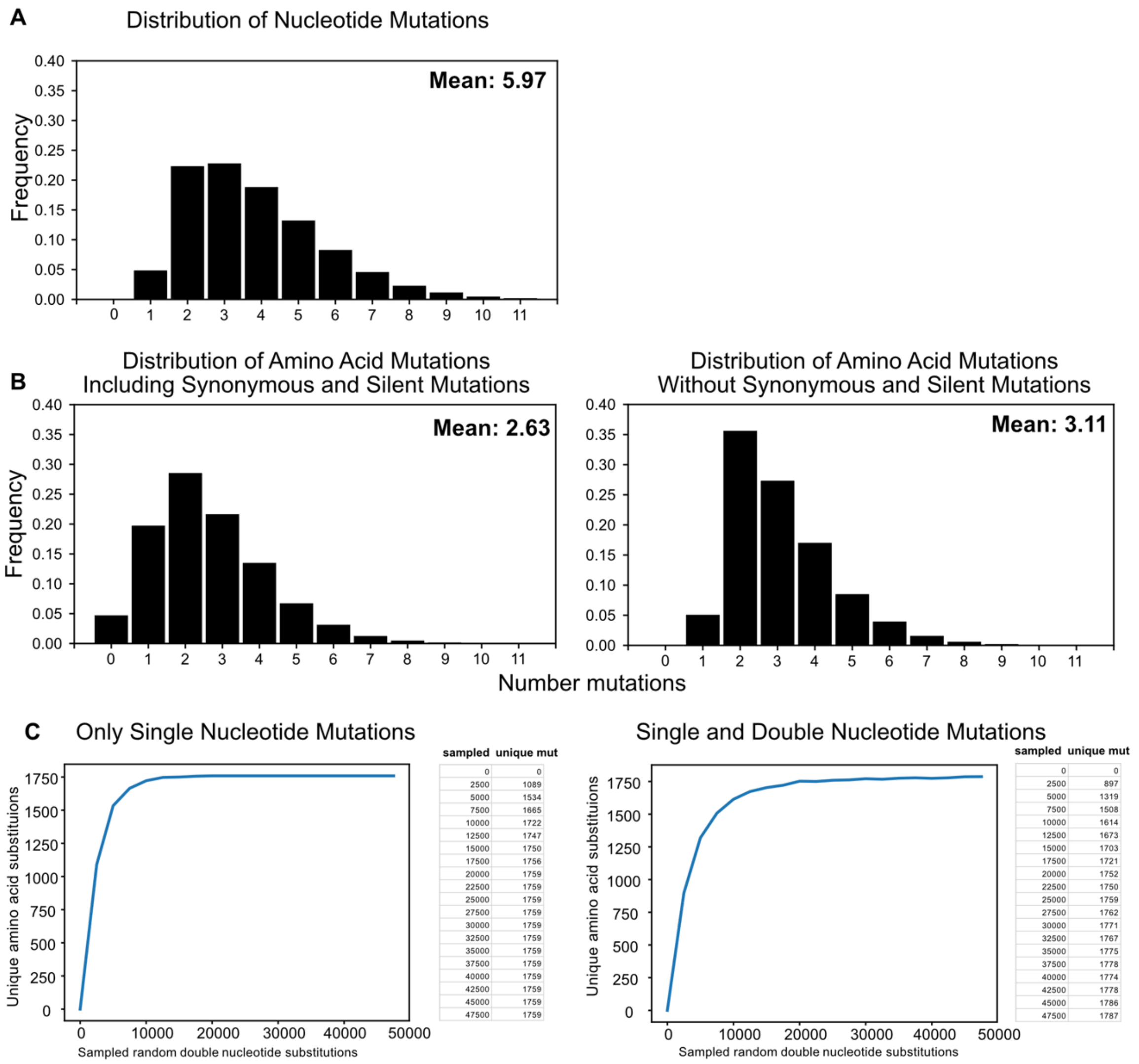

| sampled | unique mut |
|---|---|
| 0 | 0 |
| 2500 | 1089 |
| 5000 | 1534 |
| 7500 | 1665 |
| 10000 | 1722 |
| 12500 | 1747 |
| 15000 | 1750 |
| 17500 | 1756 |
| 20000 | 1759 |
| 22500 | 1759 |
| 25000 | 1759 |
| 27500 | 1759 |
| 30000 | 1759 |
| 32500 | 1759 |
| 35000 | 1759 |
| 37500 | 1759 |
| 40000 | 1759 |
| 42500 | 1759 |
| 45000 | 1759 |
| 47500 | 1759 |

| sampled | unique mut |
|---|---|
| 0 | 0 |
| 2500 | 897 |
| 5000 | 1319 |
| 7500 | 1508 |
| 10000 | 1614 |
| 12500 | 1673 |
| 15000 | 1703 |
| 17500 | 1721 |
| 20000 | 1752 |
| 22500 | 1750 |
| 25000 | 1759 |
| 27500 | 1762 |
| 30000 | 1771 |
| 32500 | 1767 |
| 35000 | 1775 |
| 37500 | 1778 |
| 40000 | 1774 |
| 42500 | 1778 |
| 45000 | 1786 |
| 47500 | 1787 |

**Figure S4: SrIRED - Distribution of mutation number in the dictionary.** The error-prone PCR (ePCR) library of *Sr*IRED was sequenced using UMIC-seq to generate a dictionary of 29,841 unique variants present before selection. Analysis of the obtained dictionary shows that the available number of single-mutation datapoints number is capped by codon bias. **(A)** Distribution of mutation number of all unique nucleotide mutations in the dictionary. **(B)** Distribution of amino acid mutation number before and after redundant and synonymous variants are removed. The main effect is a loss of apparent diversity in single point mutations and an upward shift in the average number of mutations from 2.63 to 3.11. In total we observe 1514 unique single, 10626 unique double and 17695 unique variants with more than 2 mutations (**C**) Simulation of the number of expected unique amino acid mutations with different numbers of sampled nucleotide mutations. With only single nucleotide mutations (e.g. C533G; *left*) and with double nucleotide mutations (e.g. C533G C630T; *right*). If only single nucleotide mutations are allowed, the simulation quickly reaches a plateau of 1759 unique single mutations, saturating the available diversity (the long tail on the plateau are silent mutations). When we additionally allow double nucleotide mutations, a higher amino acid mutation diversity can be reached, but only after including significantly more nucleotide mutations (i.e. a slight increase instead of levelling off). With approximately 60,000 single nucleotide mutations in the library, the 1514 observed single amino acid mutants in the dictionary therefore represent a high portion (86%) of the achievable theoretical diversity.

These considerations show how the architecture of the genetic code restricts the sequence space available to directed evolution using ePCR. Due to the observed codon bias nearly all higher order mutations are combinations of single point mutants also observed in the dataset providing rich information on the combinability and epistasis in these variants.

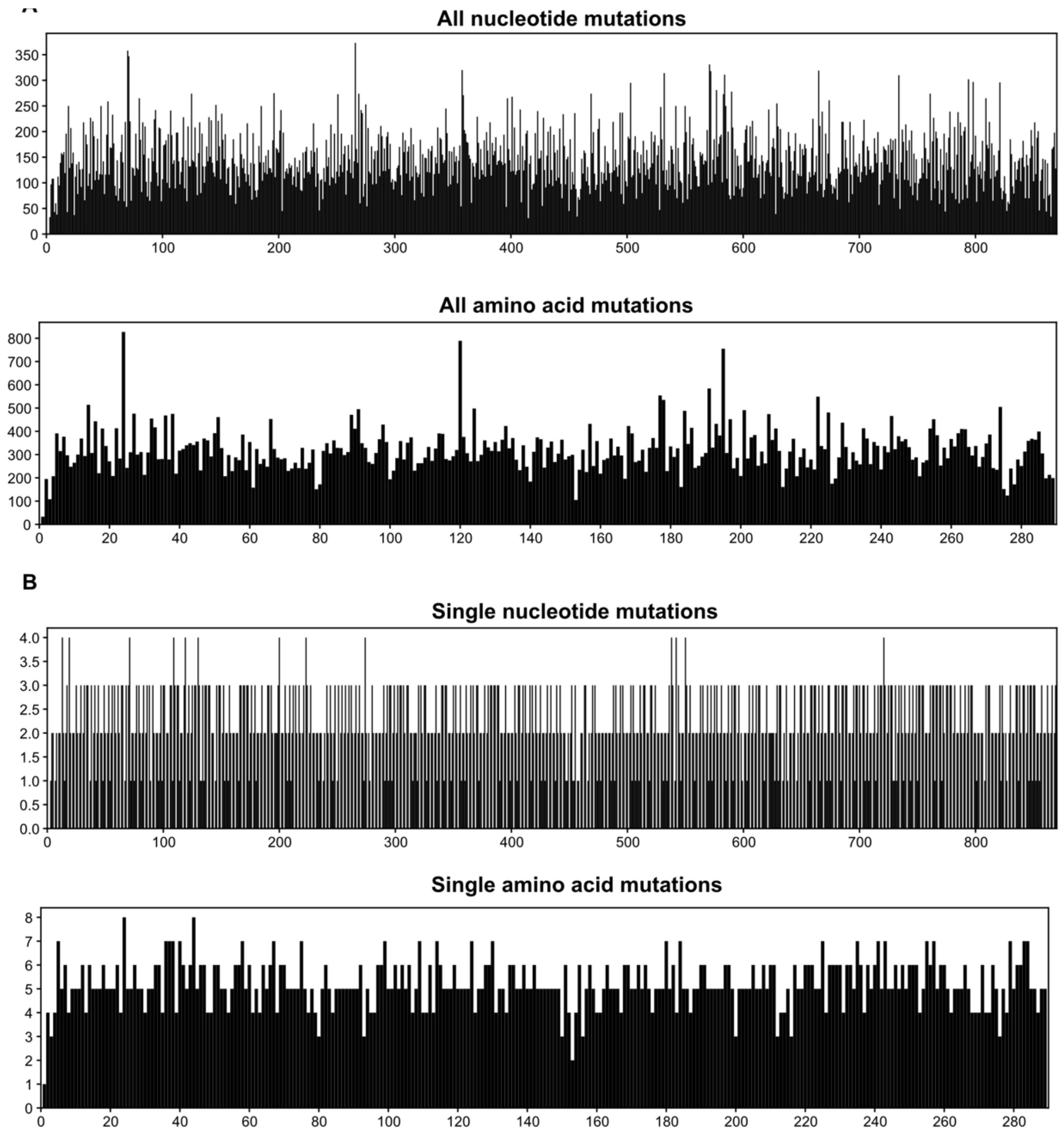

**Figure S5: Distribution of mutations in the random mutagenesis library of *Sr*IRED prior to selection.** The error-prone PCR library of *Sr*IRED was sequenced using UMIC-seq to generate a_dictionary of 29,841 unique variants (quantified at the level of *amino acid* mutations) present before selection. The dictionary thus obtained was used to decode the distribution of mutations. **(A)** Distribution of all unique nucleotide and amino acid mutations (single and higher order mutations) in the dictionary. Nucleotide and amino acid mutations are not equally distributed over the dictionary implying that errors caused by error-prone PCR randomisation are not evenly distributed over the sequence. However, the positions with high occurrence of nucleotide or amino acid mutations do not overlap with the calculated combinability hotspots, suggesting that those are independent of observed biases in the library before sorting. **(B)** Distribution of single nucleotide and amino acid mutations (single mutations only). Single mutations are much more evenly distributed than higher-order mutations which can be attributed to an upper limit of 1759 (**Extended Data Figure 2**) amino acid mutations that are accessible via single nucleotide changes in *Sr*IRED, capping potential diversity. This analysis suggests that observed enrichment and depletion of single mutations after sorting informs on mutability. The analysis also suggests that calculated mutability scores (based on single mutations) are not influenced by any biases in the library before sorting. The low amount of observed single mutations can be explained by a high number of higher-order mutations including silent mutations that lead to single amino acid mutations.

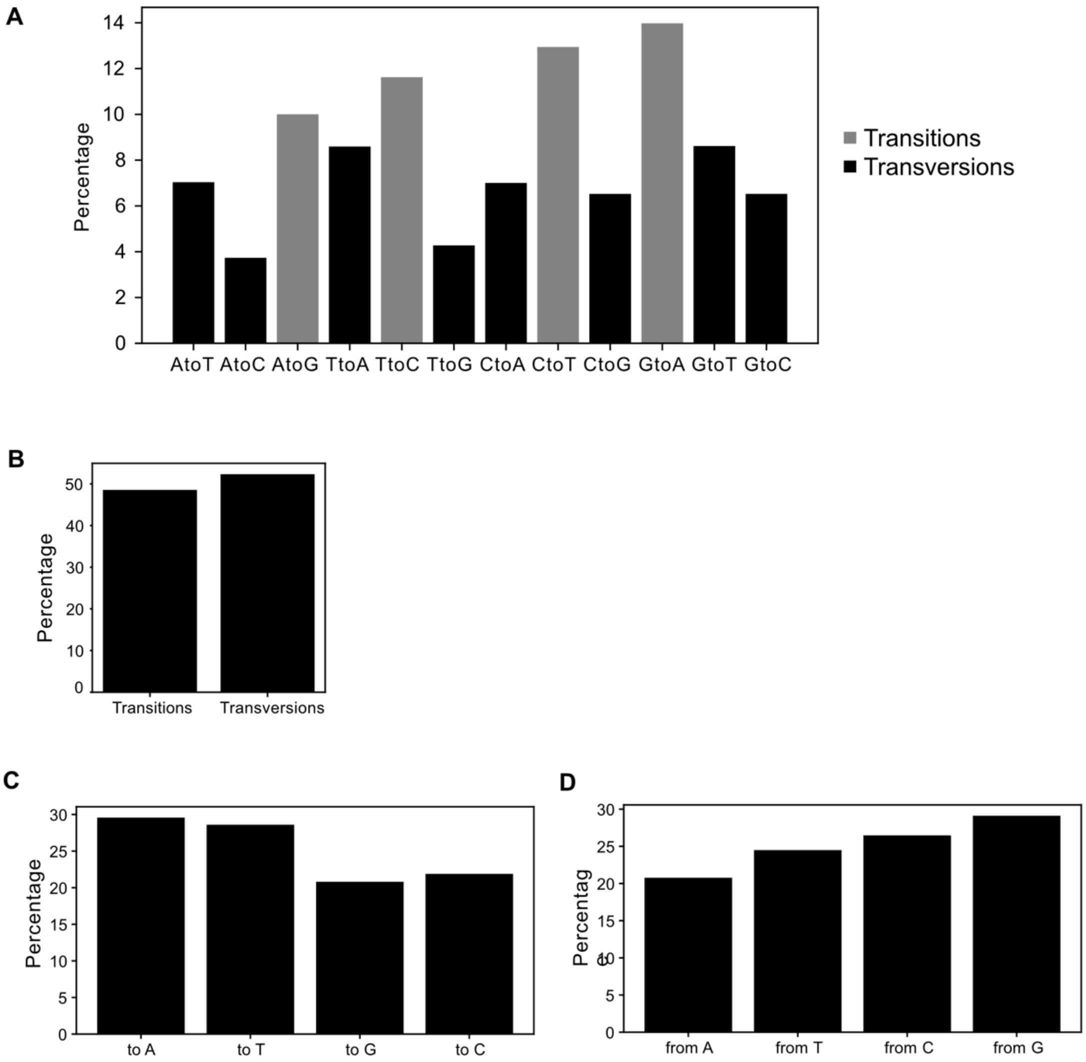

**Figure S6: Analysis of nucleotide mutations by type based on codon usage.** The error-prone PCR library of *Sr*IRED was sequenced using UMIC-seq to generate a_dictionary of 29,841 unique variants present before selection and the obtained dictionary was used to analyse potential substitution biases of error-prone PCR. (**A**) Consideration of all possible nucleotide mutations reveals an intrinsic bias towards transition mutations. (**B**) Transitions are as likely as transversions, even though there are 8 possible transversions and only 4 possible transitions. (**C**) Mutations to A and T (58%) are slightly more likely than mutations to G or C (42%). (**D**) Also, mutations from C or G (55%) are slightly more likely than mutations from A or T (45%). These results are consistent with the slight GC to AT bias observed previously observed in Mutazyme polymerase[1]. Appearance of single mutations however seems to be capped much more by lacking appearance of several nucleotide substitution in the same codon (Extended Data Figure 2) as - assuming no biases at all - only 1759 amino acid mutations can be reached, and we observe 1514 unique amino acid mutations in the dictionary.

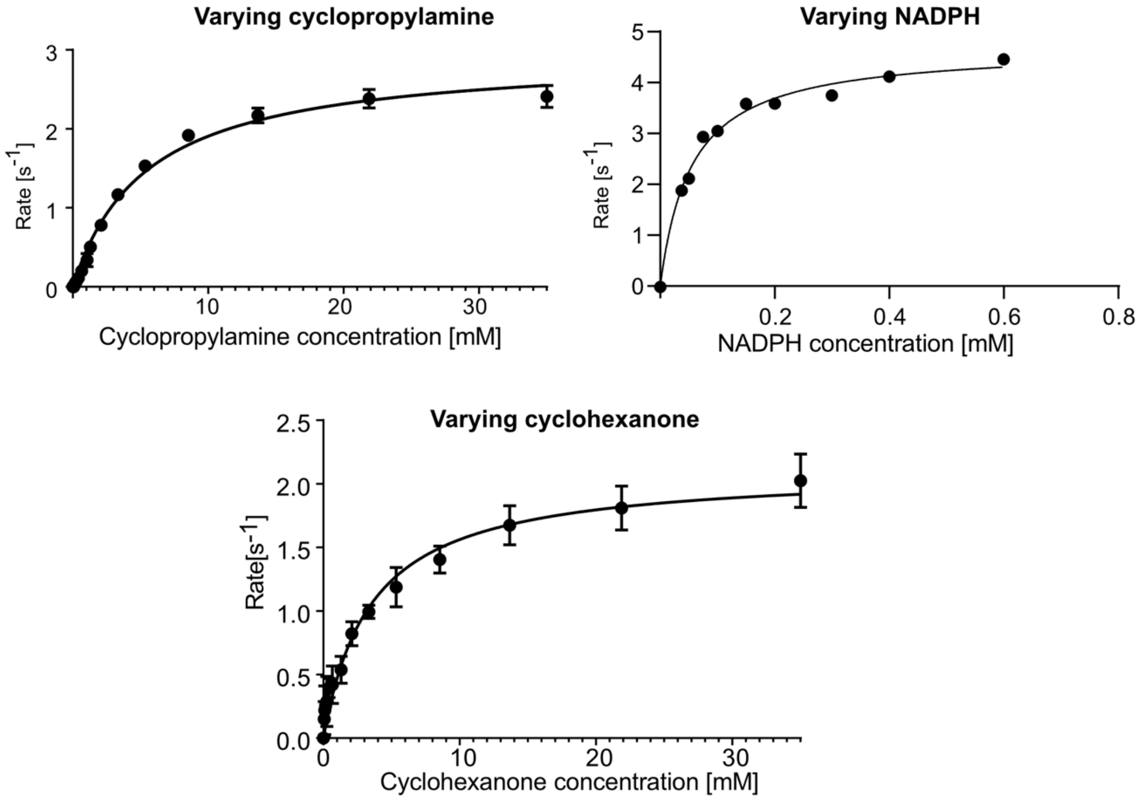

| IRED variant | Variable substrate | Constant substrates | $k_{cat}$ [$s^{-1}$]* | $k_{cat}$ error | $K_M$ [mM]* | $K_M$ error | $k_{cat}/K_M$ [$s^{-1}$ $mM^{-1}$]* | $k_{cat}/K_M$ error |
|---|---|---|---|---|---|---|---|---|
| wt | **1** | **a**<br>NADPH | 2.11 | 0.08 | 3.52 | 0.44 | 0.60 | 0.08 |
| wt | **a** | **1**<br>NADPH | 2.95 | 0.07 | 5.47 | 0.35 | 0.54 | 0.04 |
| wt | NADPH | **1**<br>**a** | 4.70 | 0.13 | 0.05 | 0.006 | 94.0 | 11.6 |

**Figure S7: Michaelis-Menten kinetics of wild type *Sr*IRED (wt).** For these multi-substrate-reactions, kinetics were approximated by varying one of the three components (cyclohexanone **1**, cyclopropylamine **a**, or the NADPH cofactor) in the presence of a constant amount of the other two in 100 mM Tris pH 8.0 (when kept constant, concentrations were the following: [cyclohexanone] = 30 mM (9 x $K_M$), [cyclopropylamine] = 30 mM (5 x $K_M$), [NADPH] = 0.5 mM (10 x $K_M$)). Initial rates were determined in transparent Nunc 96-well plates (ThermoFisher) and by monitoring the depletion of NADPH at 340 nm (SpectraMax 190, Molecular Devices, at 25°C) with 0.001 - 0.01 mg/ml enzyme. Initial rates were measured in 3 replicates and curves were fitted using Prism (error bars in the reported Michaelis-Menten plots are small and sometimes not visible). Equivalent experimental conditions were used for all ketone kinetics of the characterised *Sr*IRED variants.

| | Fitness | Lysate activity [fold-change] | Soluble expression [fold-change] | $T_M$ [°C] | $k_{cat}$ [$s^{-1}$] | $K_M$ [mM] | $k_{cat}/K_M$ [$mM^{-1}s^{-1}$] |
|---|---|---|---|---|---|---|---|
| wild type | 0 | 1 | 1 | 38.5 ± 0.6 | 2.1 ± 0.1 | 3.5 ± 0.3 | 0.6 ± 0.1 |
| **T241A** | **1.8 ± 0.5** | **1.3 ± 0.5** | **0.8** | **50.0 ± 0.7** | **5.8 ± 0.2** | **2.3 ± 0.3** | **2.5 ± 0.4** |
| **T241S** | **0.7 ± 0.6** | **2.8 ± 0.7** | **1.2** | **37.6 ± 0.6** | **6.0 ± 0.4** | **2.3 ± 0.5** | **2.6 ± 0.6** |
| M120L | 0.3 ± 0.4 | 3.7 ± 0.9 | 1.0 | 33.3 ± 0.1 | 2.1 ± 0.1 | 5.3 ± 0.7 | 0.4 ± 0.1 |
| L174V | 1.1 ± 1.4 | 3.8 ± 1.2 | 0.7 | 27.6 ± 0.2 | 10.7 ± 0.5 | 13.8 ± 1.5 | 0.8 ± 0.1 |
| D69H | 0.5 ± 2.4 | 3.3 ± 1.0 | 1.2 | 39.5 ± 0.6 | 4.1 ± 0.2 | 5.3 ± 0.6 | 0.8 ± 0.1 |
| D69G | 0.5 ± 0.5 | 5.7 ± 1.8 | 1.4 | 38.8 ± 0.6 | 3.3 ± 0.1 | 6.0 ± 0.4 | 0.5 ± 0.04 |
| D236G | 0.4 ± 0.6 | 2.5 ± 0.7 | 1.4 | 27.9 ± 0.4 | 2.0 ± 0.1 | 3.1 ± 0.4 | 0.7 ± 0.1 |
| P203A | 2.9 ± 1.3 | 0.2 ± 0.1 | n.d. | 40.8 ± 0.2 | 5.0 ± 0.2 | 6.7 ± 0.8 | 0.7 ± 0.1 |
| P203S | 0.7 ± 0.5 | 1.6 ± 0.4 | 0.9 | 43.7 ± 1.7 | 2.5 ± 0.1 | 3.2 ± 0.2 | 0.8 ± 0.1 |
| E146V | 0.9 ± 0.6 | 4.2 ± 0.9 | 1.7 | 31.1 ± 0.5 | 1.7 ± 0.1 | 2.4 ± 0.3 | 0.7 ± 0.1 |
| D267G | 0.2 ± 0.5 | 5.4 ± 1.2 | 1.9 | 43.3 ± 0.6 | 2.7 ± 0.1 | 4.3 ± 0.3 | 0.6 ± 0.1 |
| D5H | 1.5 ± 2.3 | 3.0 ± 1.1 | 1.8 | 39.6 ± 0.05 | 1.9 ± 0.1 | 5.1 ± 0.9 | 0.4 ± 0.1 |
| V6L V67I | 4.8 ± 2.3 | n.d. | n.d. | 29.1 ± 0.6 | 1.7 ± 0.0 | 2.3 ± 0.2 | 0.7 ± 0.1 |

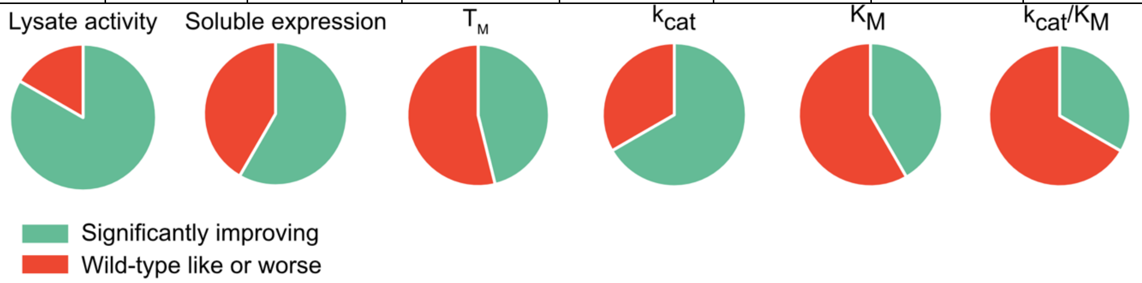

**Figure S8: Characterization of 12 selected single-point mutants with positive fitness and the variant with the highest fitness in the library V6L V67I.** (**A**) Lysate activity is given as initial rates measured in three replicates (error bars represent standard deviations; with **1** (10 mM), **2** (20 mM), NADPH (0.5 mM) in 100 mM Tris pH 8.0) by following the decline of NADPH absorbance at 340 nm and normalized to lysate without IRED. Soluble expression was determined by densitometric analysis of SDS-PAGE bands of the soluble and insoluble fractions after lysis. Values were normalized to wild type. Melting temperatures $T_M$ were measured in thermal shift assays in three replicates, errors represent standard deviations. Michaelis Menten kinetics were conducted at T = 25 °C with variable concentrations of ketone substrate (0.05 - 35 mM) with a constant concentration of amine (30 mM) in Tris pH 8.0 with NADPH (0.5 mM) and with variable enzyme concentrations from 0.03 μM to 0.3 μM (adjusted to be convenient to measure). Each curve was measured in three replicates. Errors represent standard errors of the fit and three technical replicates with a 95% confidence interval. Amine and NADPH kinetics as well as Michaelis-Menten curves for wild type are provided in

**Supplementary Figure 7**. (**B**) Pie charts representing the fraction of variants significantly improved (1 standard deviation over wild type).

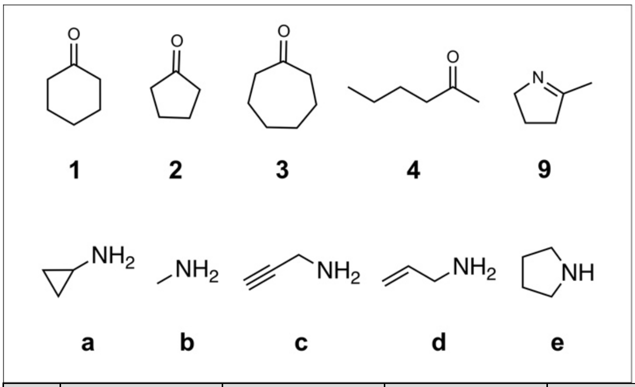

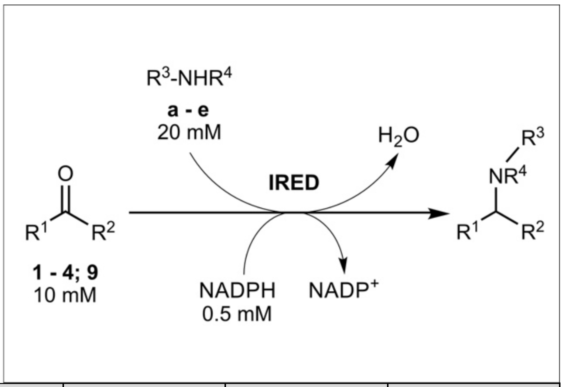

| | WT | | T241A | | D69H | | K38E | | E146V | | D236G | | D5H | |
|---|---|---|---|---|---|---|---|---|---|---|---|---|---|---|
| | specific activity | ± | specific activity | ± | specific activity | ± | specific activity | ± | specific activity | ± | specific activity | ± | specific activity | ± |
| | [mU/mg] | | [mU/mg] | | [mU/mg] | | [mU/mg] | | [mU/mg] | | [mU/mg] | | [mU/mg] | |
| **1a** | 3978 | 298 | 13062 | 485 | 9133 | 253 | 7287 | 197 | 3219 | 890 | 5339 | 987 | 6531 | 1795 |
| **2a** | 341 | 72 | 492 | 189 | 317 | 53 | 339 | 57 | 293 | 37 | 369 | 62 | 487 | 96 |
| **3a** | 308 | 9 | 360 | 7 | 299 | 7 | 230 | 25 | 238 | 7 | 225 | 11 | 283 | 10 |
| **4a** | 27 | 7 | 58 | 2 | 29 | 4 | 18 | 0 | 19 | 3 | 17 | 2 | 24 | 2 |
| **9** | 2280 | 205 | 1380 | 152 | n.d. | | n.d. | | n.d. | | n.d. | | n.d. | |
| **1b** | 471 | 26 | 422 | 41 | 499 | 34 | 384 | 57 | 482 | 38 | 444 | 49 | 467 | 23 |
| **1c** | 259 | 34 | 461 | 56 | 360 | 18 | 110 | 1 | 278 | 37 | 172 | 35 | 339 | 20 |
| **1d** | 77 | 3 | 136 | 2 | 83 | 9 | 27 | 1 | 79 | 5 | 54 | 5 | 88 | 8 |
| **1e** | 7 | 1 | 11 | 1 | 10 | 1 | 2 | 0 | 11 | 0 | 6 | 1 | 9 | 0 |

**Figure S9: Substrate scoping of wt *Sr*IRED and selected mutants based on initial rates.** Specific activities in mU/ mg were determined at room temperature using 10 mM ketones **1 – 4** and 20 mM amines **a – e** or 10 mM imine **9** as well as 0.5 mM NADPH in 100 mM Tris-HCl buffer at pH 8. Enzyme concentrations varied depending on the used substrates between 0.05 µM and 3 µM (see **Supplementary Methods**). Activities were measured in triplicate in 96-well plates by following the initial rate of NADPH depletion at 340 nm in a spectrophotometer (Spectramax 190, Molecular Devices). Substrate depletion was quantified using the Beer-Lambert law and specific activity was derived by taking into account the enzyme concentration for normalisation (n.d. = not determined). Activity values listed in the table were used for **Extended Data Figure 4D**.

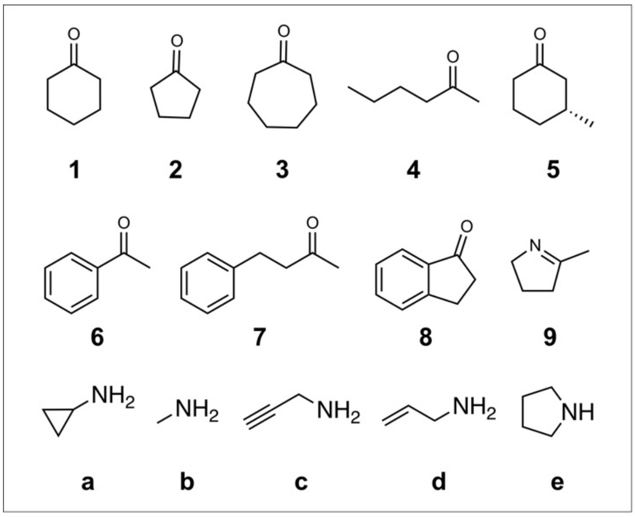

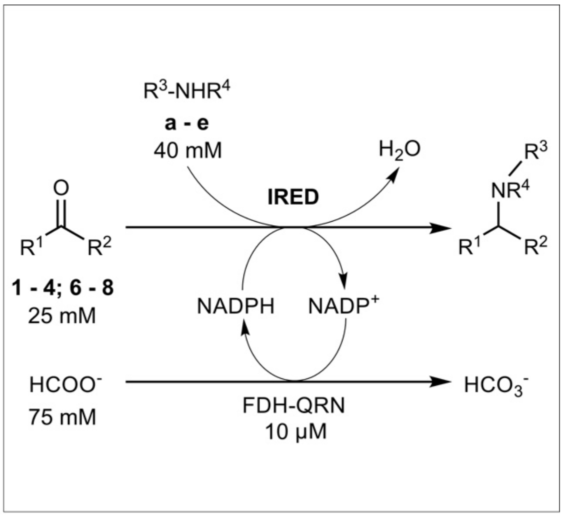

| | | Conversion 4h [%] | | | | Conversion 24h [%] | | | |
|---|---|---|---|---|---|---|---|---|---|
| Ketone | Amine donor | WT | ± | T241A | ± | WT | ± | T241A | ± |
| **1** | **a** | 83 | 0.0 | 97 | 0.4 | 98 | 1.0 | 100 | 0.1 |
| **2** | **a** | 43 | 2.5 | 53 | 1.7 | 70 | 1.3 | 83 | 2.7 |
| **3** | **a** | 34 | 0.1 | 39 | 0.2 | 64 | 2.3 | 73 | 1.6 |
| **4** | **a** | 9 | 1.0 | 19 | 1.0 | 41 | 0.8 | 67 | 2.1 |
| **6** | **a** | n.d. | | n.d. | | 0.7 | | 0.4 | |
| **7** | **a** | n.d. | | n.d. | | 0 | | 0.8 | |
| **8** | **a** | n.d. | | n.d. | | 3 | | 5 | |

| | | Conversion 4h [%] | | | | Conversion 24h [%] | | | |
|---|---|---|---|---|---|---|---|---|---|
| Ketone | Amine donor | WT | ± | T241A | ± | WT | ± | T241A | ± |
| **1** | **a** | 83 | 0.0 | 97 | 0.4 | 98 | 1.0 | 100 | 0.1 |
| **1** | **b** | 79 | 3.2 | 72 | 3.8 | 100 | 0.1 | 100 | 0.1 |
| **1** | **c** | 44 | 5.2 | 54 | 3.5 | 78 | 9.3 | 89 | 6.7 |
| **1** | **d** | 35 | 1.7 | 46 | 3.2 | 56 | 1.3 | 80 | 15.3 |
| **1** | **e** | 9 | 1.8 | 18 | 0.9 | 33 | 8.0 | 34 | 2.9 |

**Figure S10: Biotransformations of wt *Sr*IRED and T241A with different substrates.** *Sr*IRED wild type and its variant T241A were tested in biocatalytic reactions for the amination of ketone substrates **1** to **4** and **6** to **8** with amine donors **a** to **e**. A variant of the formate dehydrogenase from *Candida boidinii* (FDH-QRN) was used as a recycling enzyme for NADPH. Sodium formate served as the ultimate hydride donor. Biocatalytic reactions were performed with purified enzyme (reaction conditions: see **Supplementary Methods**) and stopped after 4 h and 24 h, respectively (n.d. = not determined). As the corresponding reference amine products were not commercially available, the reported conversions were calculated based on the substrate consumption and using a calibration curve with the substrate and an internal standard. Moreover, the identity of the formed products was confirmed by GC-MS. Conversion values listed in the table were used for **Extended Data Figure 4F**.

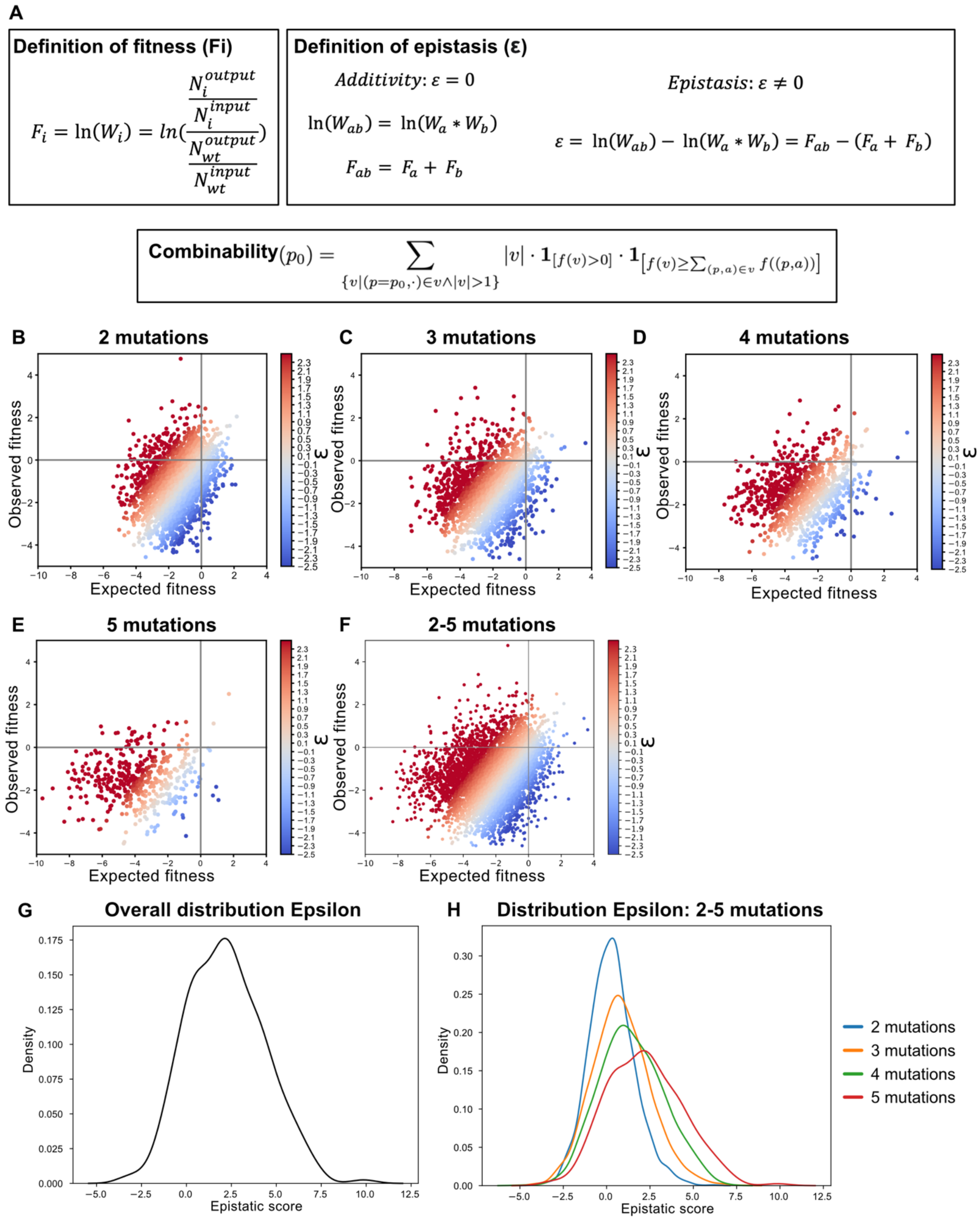

**Figure S11: Epistasis model. (A)** Definitions of fitness, epistasis and combinability from sequencing counts. *Fitness definition:* The fitness (F) of a variant (i) was calculated from the sequencing counts (N). *Epistasis definition:* The product model[2] was used for the calculation of hypothetical additive fitness by summing up the fitness values for single mutation (calculated from sequencing counts). ε is the epistatic score of a mutation a with b (i.e. the double mutation ab), Fitness (F) and non-log fitness W. *Combinability definition*: the combinability of position p0 is the weighted sum over all higher order mutants, which contain a mutation at p0 and have positive and non-negatively epistatic fitness. The weighting is given by the mutation order |v|. The indicator function is 1 if the condition is fulfilled and 0 otherwise. Note that this definition can be extended naturally to a combinability score for individual mutations (p, a) and not just for positions p. (**B-F**) Scatter plots of hypothetical additive vs. observed fitness for all higher order variants with 2 - 5 mutations. Hypothetical additive as well as observed fitness decreases with increasing mutation order. **(G)** Overall, $\varepsilon$ (positive for positive epistasis, negative for negative epistasis) shows a distribution tilted to positive epistasis (disregarding inactive higher mutants). This suggests that in variants with residual activity there is more positive than negative epistasis in *Sr*IRED. As it can be assumed that there is no positive epistasis in the residual 4,130 inactive variants, the overall fraction of positive epistasis is lower than negative epistasis (**H**) The distribution of epistatic scores by mutation number suggests that the higher the mutation order the higher the tendency of positive epistasis which can be expected due to accumulating effects of pairwise positive epistasis. This, however, disregards potential epistatic effects arising from effects visible only in co-occurrence of mutations, for example epistatic effects linked to the stability threshold, and should not lead to the conclusion that higher-order mutations in general tend to exhibit more positive epistasis.

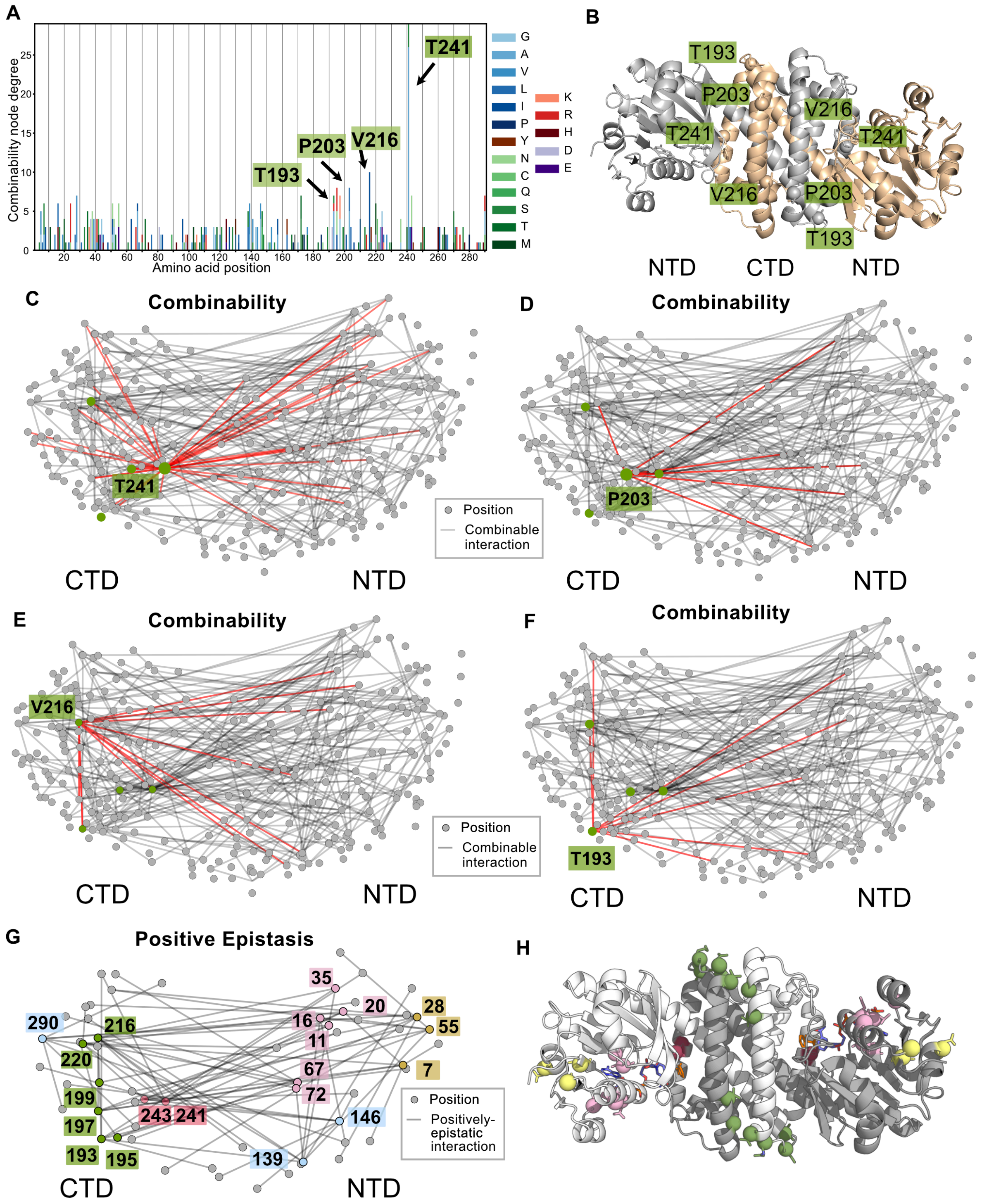

| I | Fitness | Combinability score | Lysate activity [fold-change] | $T_M$ [°C] | $k_{cat}$ [$s^{-1}$] | $K_M$ [mM] | $k_{cat}/K_M$ [$mM^{-1}s^{-1}$] |
|---|---|---|---|---|---|---|---|
| wt | 0 | | | 38.5 ± 0.6 | 2.1 ± 0.1 | 3.5 ± 0.3 | 0.6 ± 0.1 |
| T241A | 1.8 ± 0.5 | 29 | 1.3 ± 0.5 | **50.0 ± 0.7** | 5.8 ± 0.2 | 2.3 ± 0.3 | 2.5 ± 0.4 |

| | | | | | | | |
|---|---|---|---|---|---|---|---|
| T241S | 0.7 ± 0.6 | 29 | 2.8 ± 0.7 | 37.6 ± 0.6 | 6.0 ± 0.4 | 2.3 ± 0.5 | 2.6 ± 0.6 |
| V216I | -0.6 ± 0.6 | 10 | 1.1 ± 0.3 | 38.8 ± 0.9 | 3.5 ± 0.1 | 4.3 ± 0.4 | 0.8 ± 0.1 |
| P203A | 2.9 ± 1.3 | 8 | 0.2 ± 0.1 | **40.8 ± 0.2** | 5.0 ± 0.2 | 6.7 ± 0.8 | 0.7 ± 0.1 |
| P203S | 0.7 ± 0.5 | 8 | 1.6 ± 0.4 | **43.7 ± 1.7** | 2.5 ± 0.1 | 3.2 ± 0.2 | 0.8 ± 0.1 |
| T193A | 0.1 ± 0.4 | 8 | 0.8 ± 0.3 | 33.7 ± 0.5 | 2.6 ± 0.1 | 2.0 ± 0.2 | 1.3 ± 0.2 |
| K38E | 0.0 ± 0.7 | 5 | 0.6 ± 0.1 | 37.7 ± 1.1 | 3.2 ± 0.1 | 6.3 ± 0.6 | 0.5 ± 0.05 |
| D69H | 0.5 ± 2.4 | 0 | 3.3 ± 1.0 | 39.5 ± 0.6 | 4.1 ± 0.2 | 5.3 ± 0.6 | 0.8 ± 0.1 |
| L174V | 1.1 ± 1.4 | 2 | 3.8 ± 1.2 | 27.6 ± 0.2 | 10.7 ± 0.5 | 13.8 ± 1.5 | 0.8 ± 0.1 |
| T241A P203A | - | | 1.5 ± 0.3 | 50.6 ± 0.4 | 15.0 ± 0.4 | 2.3 ± 0.2 | 6.6 ± 0.6 |
| T241A D69H | - | | 8.7 ± 1.7 | 47.0 ± 0.6 | 12.1 ± 0.3 | 2.4 ± 0.2 | 5.1 ± 0.5 |
| T241A K38E | - | | 3.1 ± 0.8 | 50.8 ± 0.4 | 11.3 ± 0.3 | 1.9 ± 0.2 | 6.1 ± 0.6 |
| T241A L174V | - | | 2.8 ± 0.2 | 37.2 ± 0.2 | 12.7 ± 0.3 | 5.3 ± 0.4 | 2.4 ± 0.2 |
| V216I D69H | - | | 3.3 ± 0.7 | 37.8 ± 0.1 | 5.4 ± 0.2 | 4.3 ± 0.5 | 1.3 ± 0.2 |
| V216I K38E | - | | 1.0 ± 0.3 | 37.5 ± 0.8 | 5.2 ± 0.2 | 5.3 ± 0.7 | 1.0 ± 0.1 |
| K38E D69H | - | | 0.3 ± 0.01 | 36.7 ± 0.1 | 4.3 ± 0.3 | 12.7 ± 2.0 | 0.3 ± 0.1 |
| T241A K38E D69H | - | | 1.4 ± 0.1 | 50.2 ± 0.9 | 24.5 ± 1.0 | 7.1 ± 0.8 | 3.4 ± 0.4 |
| T241A V216I | - | | 2.9 ± 0.7 | 48.8 ± 0.6 | 7.7 ± 0.2 | 4.9 ± 0.4 | 1.6 ± 0.1 |

**Figure S12: Combinability hotspots. (A)** Positional frequency in highly active and combinable variants. Amino acid identity of mutations is colour-coded (blues: non-polar aliphatic, brown: aromatic, greens: polar, uncharged, reds: positively charged, violets: negatively charged). T241 is the global hotspots of combinability, followed by V216. **(B)** *Sr*IRED structure (monomers indicated in grey and orange; PDB ID: 5ocm) with hotspots represented as spheres identifies them in the C-terminal domain and first shell. **(C-F)** Combinability graphs showing combinable interactions of mutations at distinct positions (CTD = C-terminal domain; NTD = N-terminal domain). Nodes represent positions in the IRED sequence that are grouped by distance in the IRED structure. Connections represent cooccurrence in a higher-order variant whose corresponding fitness is larger or equal to the combined effect in single mutations (i.e. combinability). The high node degree of T241 (edges in red) indicates high potential for combinability and positive epistasis with all parts of the protein while interactions of P203, V216 and T193 are more focused on the N-terminal domain. **(G)** A simplified graph only including positively epistatic interactions grouped by distance of nodes in the SrIRED structure shows structural grouping of positive epistasis hotspots. Positions with larger than 2 positively epistatic interactions are colored (light blue, green, purple, pink and yellow). **(H)** Epistatic hotspots are marked on the *Sr*IRED crystal structure (5ocm). **(I)** Characterization of combinability and positively epistatic hotspots (yellow) and combined variants (white). While combinability hotspots T241A and P203A show improvements in melting temperature along with improvements in rates, V216I is neutral in melting temperature and T193A is destabilizing. This, along with positive epistasis in catalytic parameters with T241A and V216I shows that combinability is not linked to the stability threshold and mutational robustness. Lysate activity was measured as initial rates in three replicates (errors represent standard deviations) with **1** (10 mM), **a** (20 mM), NADPH (0.5 mM) in Tris buffer (100 mM pH 8.0) following the decline of NADPH absorbance at 340 nm and normalized to lysate without IRED. Soluble expression was determined by densitometry of SDS-PAGE bands of the soluble and insoluble fraction after lysis. Values are normalized to wild type. Melting temperatures were measured in thermal shift assays in three replicates, errors represent standard deviations. Michaelis Menten kinetics were conducted with variable concentrations of ketone substrate (0.05 - 35 mM) and a constant concentration of amine (30 mM) in Tris pH 8 with NADPH (0.5 mM) and variable enzyme concentrations (from 0.03 µM to 0.3 µM) depending on the rate at T = 25 °C. Each curve was measured in three replicates. Errors represent standard errors of the fit and three technical replicates with a 95% confidence interval.

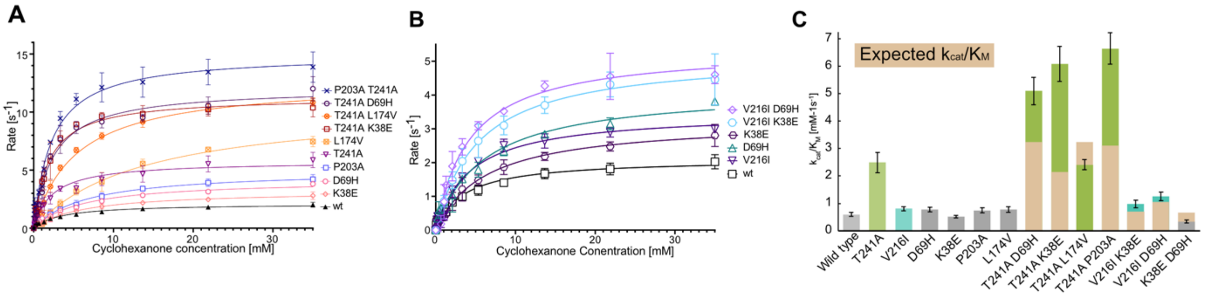

**Figure S13: Analysis of the combinability hotspot V216I.** The variant V216I was identified as second most combinable mutation in **Extended Data Figure 6C** and gives rise to improving mutations with slight positive epistasis when combined with K38E and D69H. Raw Michaelis Menten Plots are shown in (**A**) and **(B)** and $k_{cat}/K_M$ is shown in **(C). (A)** Effect of T241A upon combination with three mutations from hotspots (K38E, D69H, P203A) and the second-best mutation from the first shell (L174V). Strong positive epistasis can be observed with K38E, D69H and P203A. **(B)** Effect of combining V216I with K38E and D69H shows improved double mutations with slightly higher than additive $k_{cat}/K_M$. **(C)** Expected $k_{cat}/K_M$ is shown in brown, combinations with T241A in green and with V216I in blue. A 1.6-fold improvement is observed for V216I K38E and a 2-fold improvement for V216I D69H, showing that combinability analysis can also be used for mutations beyond T241A.

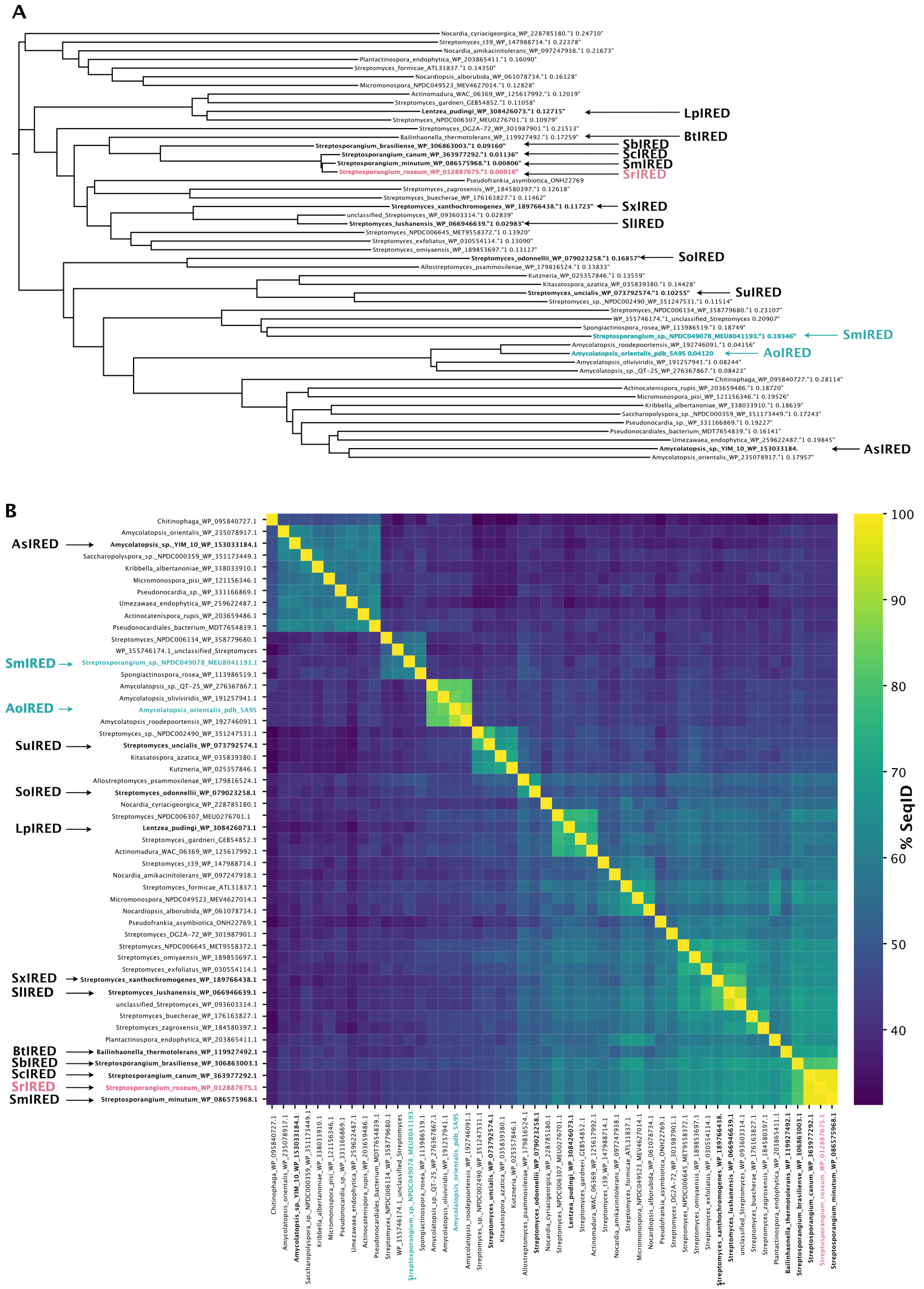

**Figure S14: Discovery of SrIRED homologs.** Homologs of SrIRED were identified by NCBI BLAST search of the uniprot database (07/2024). For sequences see "**Sequences**" section where threonine residues corresponding to T241 in SrIRED are labelled in bold. For gene accession numbers and organism name see **Table S8**. Hits were filtered for presence of threonine at position 241 (SrIRED numbering) in a multiple sequence alignment. A phylogenetic tree **(A)** and a percent identity matrix **(B)** are shown here. IREDs chosen for experimental analysis (Figure 3G & H) with and without the T241A mutation that enhanced $k_{cat}/K_M$ 4-fold in SrIRED (red) are marked in black. Blue IREDs could not be expressed in sufficient yield to enable kinetic analysis using the standard expression and purification protocols outlined in the Methods section.

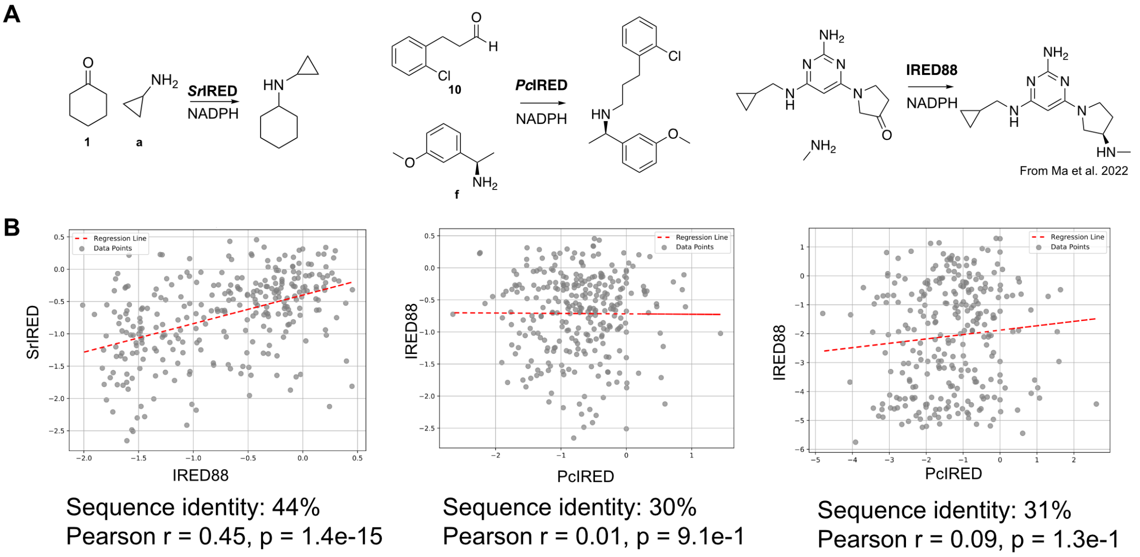

C

| IRED88 | mutability_IRED88 | SrIRED | mutability_SrIRED | position_PcIRED | mutability_PcIRED |
|---|---|---|---|---|---|
| S114 | **0.3** | E105 | **0.2** | E107 | **0.4** |
| L119 | 0.1 | R110 | 0.2 | M114 | 0.0 |
| N238 | 0.1 | A227 | 0.1 | D231 | 0.0 |

**Figure S15: Transferability of mutability to other IRED engineering campaigns. (A)** Reaction schemes of reactions sequence function data is available for SrIRED, PcIRED and IRED88[3]. **(B)** Correlation of mutability between datasets from different engineering campaigns. Despite recorded with two different substrate pairs and for IREDs with only 44% sequence similarity there is still a global correlation of mutability between the IRED88 and SrIRED deep mutational scans. No correlation can be observed between SrIRED and PcIRED and PcIRED and IRED88 which all show significantly lower sequence identity (30 and 31%) compared to the SrIRED IRED88 pair (44%). **(C)** Positions with high mutability in all three campaigns. Position 105 (SrIRED numbering) shows positive mutability in all three campaigns.

**A**

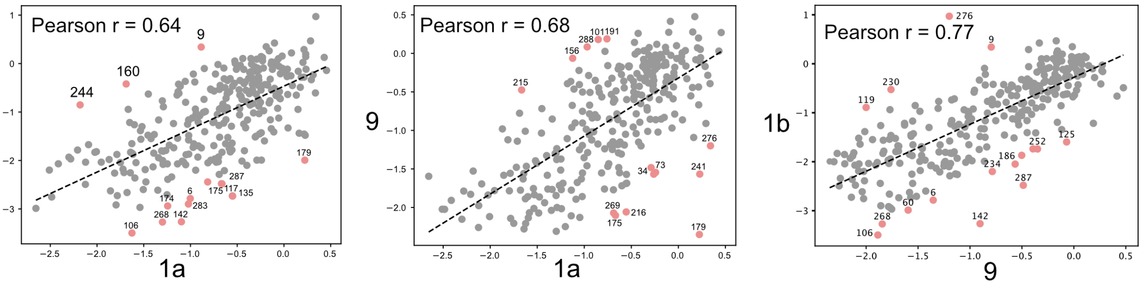

**B Mutability: Specificity-determining residues**

| Position | 1a | 1b | 9 |
|---|---|---|---|
| 2 | 0.31 | -0.23 | -0.28 |
| 9 | -0.89 | 0.34 | -0.80 |
| 70 | 0.22 | -1.46 | -0.67 |
| 74 | -0.43 | -0.41 | 0.26 |
| 128 | 0.34 | -0.61 | -0.39 |
| 179 | 0.22 | -1.99 | -2.35 |
| 203 | 0.29 | -0.33 | -0.91 |
| 241 | 0.23 | -1.48 | -1.57 |

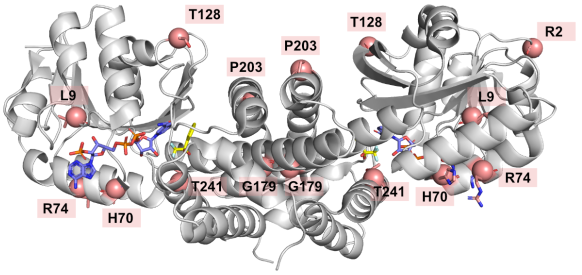

**C Mutability: Generalist residues**

| Position | 1a | 1b | 9 |
|---|---|---|---|
| 110 | 0.23 | 0.14 | 0.05 |
| 138 | 0.43 | 0.13 | 0.03 |
| 290 | 0.20 | 0.02 | 0.07 |

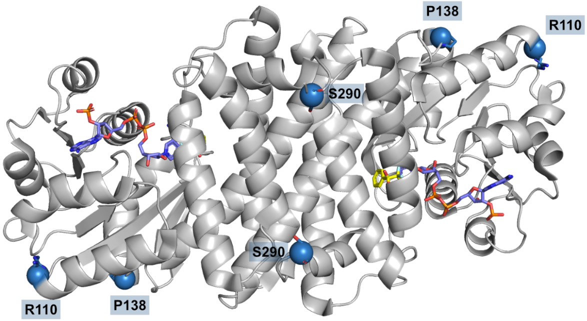

**Figure S16: Specificity-determining residues are spread all over the protein structure: Substrate-specificity is globally encoded. (A)** Mutability shows good correlation between deep mutational scans with different substrates (**B**) List of residues with mutability > 0.2 in one replicate but < 0.2 in two other replicates: Specificity-determining positions. Map on crystal structure (red) shows that specificity-determining positions are spread all across the protein structure. **(C)** Positions with improving mutability for all three substrate combinations and their position in the structure (blue).

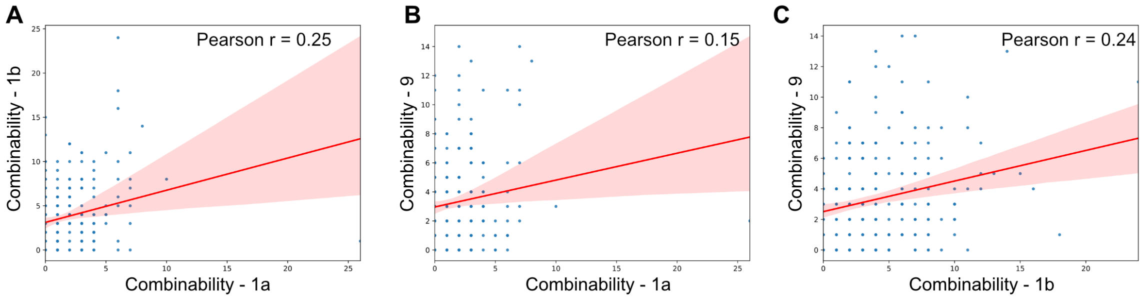

**Figure S17: Combinability is not transferable between substrates**. Correlations between combinability with: (**A**) cyclohexanone (1) with methylamine (b) and cyclohexanone (1) with cyclopropylamine (a); (**B**) (1) with (a) and 1-methyl-2-pyroline (9); (**C**) (1) with (b) and (9). Results only showed very poor correlation.

**A** **Single mutation model**

| | Full model | Full model without ESM (ESM ablation) | ESM prediction alone |
|---|---|---|---|
| | variant | variant | variant |
| 0 | P203G | P203G | S183G |
| 1 | T241G | T241G | K195G |
| 2 | M1P | M1D | Q41P |
| 3 | A251I | A251I | E36P |
| 4 | M1N | M1R | H163P |
| 5 | M1G | M1N | L231D |
| 6 | M1D | M1E | Q16S |
| 7 | M1R | M1P | L231E |
| 8 | E146L | M1G | L231G |
| 9 | M1S | P61I | A154G |
| 10 | P61A | M1S | Q274H |
| 11 | M1E | P203E | N206T |
| 12 | E36P | M1Q | H163A |
| 13 | P61V | P61F | H163V |
| 14 | P203E | A251F | F132V |
| 15 | T241E | T241E | E36A |

**B** **Double mutation model**

| | Full model | Full model without ESM (ESM ablation) | ESM prediction alone |
|---|---|---|---|
| | variant | variant | variant |
| 0 | S35T,S51G | P203L,T241A | Q41P,H163P |
| 1 | Q41P,T241A | P203A,T241K | Q41P,A154G |
| 2 | Q41P,T241S | P203Q,T241A | E36A,Q41P |
| 3 | E36A,T241A | P203A,T241R | Q41P,Q274H |
| 4 | S35T,T241A | P203L,T241P | A154G,H163P |
| 5 | Q41P,V262L | P203L,T241S | Q41P,F132L |
| 6 | P203Q,T241P | P203Q,T241P | H163P,Q274H |
| 7 | P203Q,T241A | P203S,T241K | K24R,Q41P |
| 8 | Q41P,V67L | P203Q,T241S | E36A,H163P |
| 9 | Q41P,T180V | P203R,T241A | Q41P,H247R |
| 10 | P203L,T241S | P203S,T241P | F132L,H163P |
| 11 | P203L,T241A | P203A,T241A | K24R,H163P |
| 12 | K24R,S51G | P203A,T241S | H163P,H247R |
| 13 | P203S,T241K | P203S,T241A | Q41P,T200A |
| 14 | P203A,T241K | P203A,T241P | Q41P,L231P |
| 15 | S51G,T193S | T193S,T241A | Q41P,S166A |

**Figure S18: Raw predictions from a single mutant and double mutant AI models for SrIRED.** (**A**) Raw predictions from single mutant model show that ESM has minimal influence on the top predictions, only changing the identity of the substitution at position 1. The predictions match very well with the predictions that also leverage ESM (the top-performing variant T241G is still ranked second) only replacing M1P with M1D and loosing E146L which are not our top-performing variants. This shows that ESM has minimal influence on the prediction of top hits with the model mostly relying on lrDMS data for its prediction. We also inspected the position of T241G and T241A in the pure ESM ranking and found them ranked 457/5510 and 180/5510, respectively. Therefore, the best single mutation (T241G) would only have been found when screening a relatively large number of single mutants (457+, ~10% of single mutant space) and while the good single mutant (T241A) was ranked 180$^{th}$ (~3% of single mutant space), it would not have been synthesized in our experimental setup where only the top 5 variants are selected. The ESM prediction alone - without assay labelled data - suggests two mutations in a NADPH binding loop (**Figure 4J**) ranked among the top 5 variants. T241G is not ranked among the top predictions (**B**) The ESM ablation focuses the top predictions of the model to positions P203 and T241A, which can be rationalised by the high combinability and mutability at both positions. The top rationally engineered mutant T241A P203A (surpassing the activity of the top prediction of the double mutant model) is at these predictions and is ranked at position 11. These data support the hypothesis that even without leveraging ESM features the highly performant double mutants would have been predicted, and that using zero-shot ESM information alone misses the importance of key positions such as 203 and 241 identified only from the assay data, focussing instead on the NADPH binding loop (**Figure 4J**).The double mutant model, which uses ESM features, correctly predicts synergistic effects with T241A in this loop, but does not reach the high synergistic effect with P203A at the dimer interface in its top 5 predictions (though positions 203 and 241 appear from rank 6 onwards).

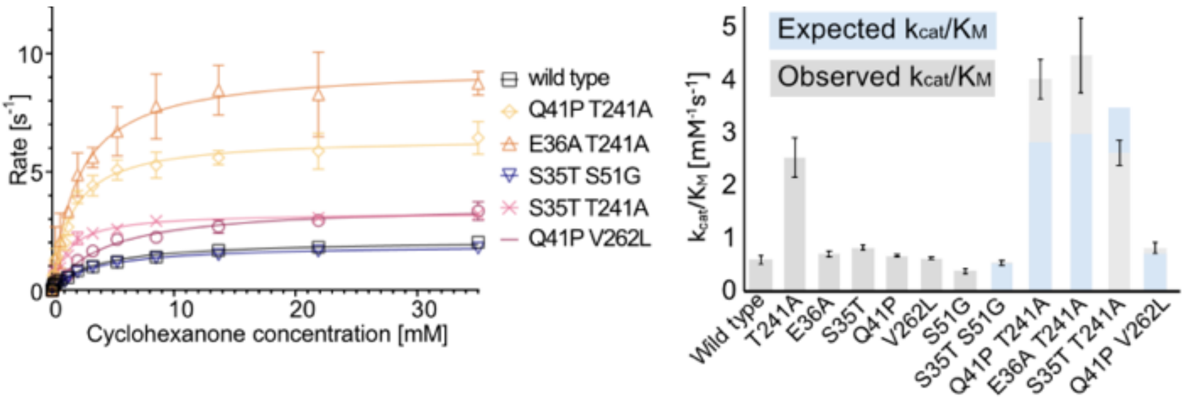

| | Lysate activity [fold-change] | $T_M$ [°C] | $k_{cat}$ [$s^{-1}$] | $K_M$ [mM] | $k_{cat}/K_M$ [$mM^{-1}s^{-1}$] |
|---|---|---|---|---|---|
| wt | 1 | 38.5 ± 0.6 | 2.1 ± 0.1 | 3.5 ± 0.3 | 0.6 ± 0.1 |
| S35T | 0.6 ± 0.04 | 33.4 ± 0.7 | 2.8 ± 0.04 | 3.4 ± 0.2 | 0.8 ± 0.04 |
| S51G | n.d. | n.d. | 2.1 ± 0.1 | 5.5 ± 0.5 | 0.4 ± 0.04 |
| Q41P | 1.6 ± 0.2 | 34.5 ± 1.5 | 2.3 ± 0.2 | 3.6 ± 0.8 | 0.6 ± 0.3 |
| T241A | 1.3 ± 0.5 | 50.0 ± 0.7 | 5.8 ± 0.2 | 2.3 ± 0.3 | 2.5 ± 0.4 |
| E36A | 0.7 ± 0.1 | 32.0 ± 0.8 | 3.7 ± 0.1 | 5.2 ± 0.3 | 0.7 ± 0.05 |
| V262L | 1.4 ± 0.1 | 30.3 ± 0.5 | 5.9 ± 0.1 | 9.5 ± 0.4 | 0.6 ± 0.03 |

**Figure S19: Single mutations occurring in ML-predicted double mutants for epistasis analysis.** Lysate activity as initial rates was measured in three replicates (errors represent standard deviations) with **1** (10 mM), **2** (20 mM), NADPH (0.5 mM) in 100 mM Tris pH 8.0, 25 °C following the decline of NADPH absorbance at 340 nm and normalized to lysate expressing no IRED. Soluble expression was determined by densitometry of SDS-PAGE bands of the soluble and insoluble fraction after lysis. Values are normalized to wild type. Melting temperatures were measured in thermal shift assays in three replicates, errors represent standard deviations. Michaelis Menten kinetics were conducted with variable concentrations of ketone substrate (0.05 - 35 mM) with a constant concentration of amine (30 mM) in Tris pH 8 with NADPH (0.5 mM) and variable enzyme concentrations (from 0.03 µM to 0.3 µM) depending on the rate (T = 25 °C). Each curve was measured in three replicates. Errors represent standard errors of the fit with a 95% confidence interval.

| | Lysate activity [fold-change] | $T_M$ [°C] | $k_{cat}$ [$s^{-1}$] | $K_M$ [mM] | $k_{cat}/K_M$ [$mM^{-1}s^{-1}$] |
|---|---|---|---|---|---|
| wt | 1 | 38.5 ± 0.6 | 2.1 ± 0.1 | 3.5 ± 0.3 | 0.6 ± 0.1 |
| E36P | 1.7 ± 0.3 | 33.1 ± 1.0 | 3.0 ± 0.2 | 4.1 ± 0.5 | 0.7 ± 0.2 |
| Q41P | 1.6 ± 0.2 | 34.5 ± 1.5 | 2.3 ± 0.2 | 3.6 ± 0.8 | 0.6 ± 0.3 |
| H163P | 1.4 ± 0.3 | 34.1 ± 1.5 | 4.5 ± 0.2 | 5.7 ± 0.8 | 0.8 ± 0.2 |
| S183G | 1.2 ± 0.3 | 33.4 ± 0.7 | 4.2 ± 0.2 | 3.4 ± 0.5 | 1.2 ± 0.2 |
| K195G | 0.6 ± 0.2 | 34.7 ± 0.9 | 6.2 ± 0.2 | 5.5 ± 0.4 | 1.1 ± 0.1 |
| E36P S183G | 0.2 ± 0.2 | 33.7 ± 1.3 | 2.4 ± 0.1 | 1.2 ± 0.1 | 2.0 ± 0.1 |
| Q41P S183G | 1.2 ± 0.4 | 34.6 ± 1.7 | 4.8 ± 0.2 | 2.7 ± 0.3 | 1.8 ± 0.2 |
| Q41P K195G | 0.6 ± 0.3 | 35.4 ± 1.8 | 5.9 ± 0.1 | 7.2 ± 0.4 | 0.8 ± 0.1 |
| H163P S183G | 0.9 ± 0.4 | 33.5 ± 2.5 | 5.4 ± 0.2 | 3.6 ± 0.5 | 1.5 ± 0.2 |
| S183G K195G | 0.7 ± 04 | 33.2 ± 0.7 | 4.2 ± 0.2 | 3.3 ± 0.6 | 1.3 ± 0.2 |

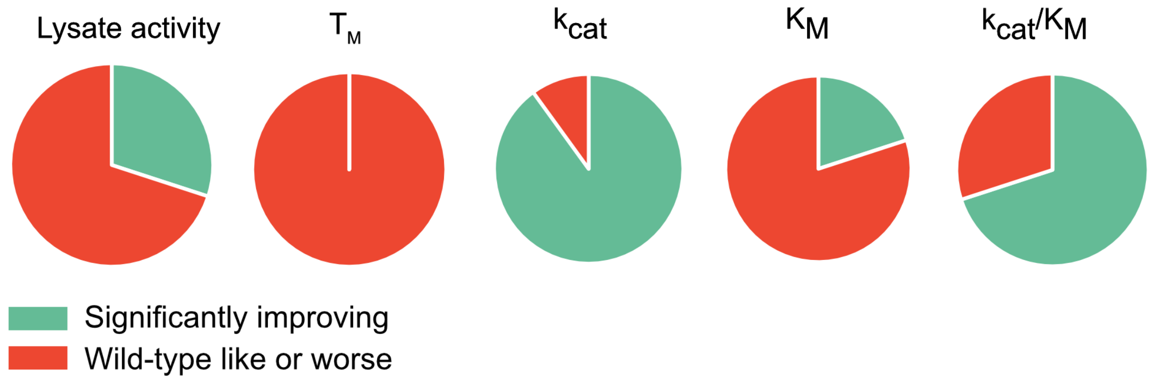

**Figure S20: ESM-predicted mutants by using ESM PLL in zero-shot mode to rank single and double mutations.** Lysate activities as initial rates were measured in three replicates (errors represent standard deviations) with **1** (10 mM), **2** (20 mM), NADPH (0.5 mM) in 100 mM Tris pH 8.0 following the decline of NADPH absorbance at 340 nm and normalized to lysate expressing no IRED. Soluble expression was determined by densitometry of SDS-PAGE bands of the soluble and insoluble fraction after lysis. Values are normalized to wild type. Melting temperatures were measured in thermal shift assays in three replicates, errors represent standard deviations. Michaelis Menten kinetics were conducted with variable concentrations of ketone substrate (0.05 - 35 mM) with a constant concentration of amine (30 mM) in Tris pH 8 with NADPH (0.5 mM) with variable enzyme concentrations (from 0.03 µM to 0.3 µM) depending on the rate (T = 25 °C). Each curve was measured in three replicates. Errors represent standard errors of the fit with a 95% confidence interval.

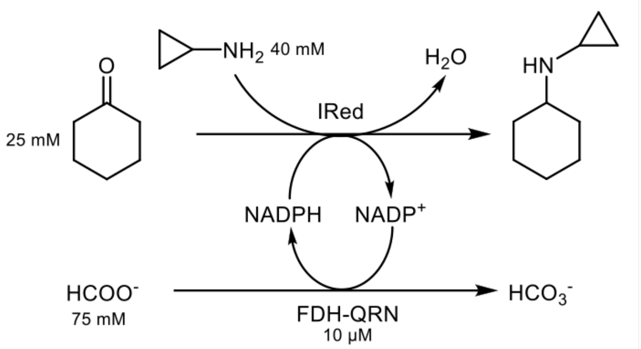

| | WT | | T241A | | D69H | | T241G | | T241A P203A | | T241G P203A (5 µM) | | T241G P203A (2.5 µM) | | T241A E36A | | T241G E36A | | T241A Q41P | |
|---|---|---|---|---|---|---|---|---|---|---|---|---|---|---|---|---|---|---|---|---|
| time (h) | conv (%) | ± | conv (%) | ± | conv (%) | ± | conv (%) | ± | conv (%) | ± | conv (%) | ± | conv (%) | ± | conv (%) | ± | conv (%) | ± | conv (%) | ± |
| 0.5 | 22 | 3 | 33 | 4 | 28 | 3 | 43 | 6 | 39 | 4 | 42 | 3 | 38 | 4 | 35 | 4 | 42 | 1 | 35 | 1 |
| 1 | 35 | 5 | 52 | 5 | 40 | 0 | 64 | 4 | 59 | 3 | 63 | 2 | 55 | 3 | 56 | 3 | 61 | 1 | 56 | 0 |
| 2 | 47 | 1 | 73 | 1 | 53 | 1 | 77 | 4 | 75 | 1 | 77 | 0 | 68 | 1 | 74 | 2 | 75 | 1 | 72 | 1 |
| 4 | 63 | 1 | 83 | 1 | 62 | 4 | 81 | 2 | 82 | 0 | 83 | 1 | 72 | 1 | 81 | 2 | 79 | 2 | 78 | 3 |

**Figure S21: Biotransformations of wt and selected mutants with substrates 1 and a.** The tested variants were from experimental screening (T241A, D69H), rational engineering (T241A-P203A), AI-based engineering (T241G, T241A-E36A, T241A-Q41P), or hybrid engineering (T241G-P203A (highlighted in green) and T241G-E36A). Reactions were carried out with cyclohexanone **1** and cyclopropylamine **a** as substrates. Reaction conditions: buffer (100 mM $K_3PO_4$, pH 8), amine donor **a** (1.6 eq., 40 mM final concentration; preparation stock solution: 80 mM of the amine donor **a** were dissolved in 100 mM $K_3PO_4$ buffer, pH 8; the pH was then re-adjusted to 8.0 with phosphoric acid). Reactions contained reagents for cofactor regeneration [sodium formate (3 eq. 75 mM, 68.01 g/mol), $NADP^+$ (0.5 mM or 1 mM, 801.4 g/mol), FDH-QRN (10 µM)] and 5 µM *Sr*IRED wild type (or one of the variants, respectively). The substrate **1** was added last (25 mM final concentration as a 1 M stock solution in DMSO). The reactions were incubated in a horizontal shaker at 30 °C. The reported conversions were calculated based on substrate consumption and using a calibration curve with the substrate and an internal standard. Moreover, the identity of the formed product was confirmed by GC-MS. Activity values listed in the table were used for **Extended Data Figure 5H**.

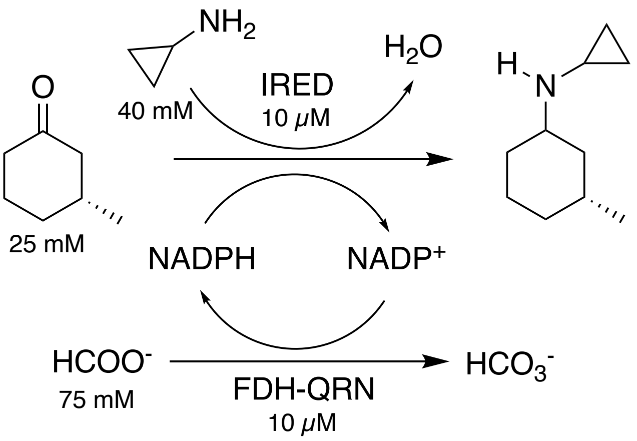

| IRED variant | conv (%) | (1*X*,3*R*) (%) | (1*Y*,3*R*) (%) | diastereomeric ratios (1*X*,3*R*) (%) | (1*Y*,3*R*) (%) |
|---|---|---|---|---|---|
| wt | 47 | 14.0 | 32.7 | 30.0 | 70.0 |
| T241A | 71 | 11.0 | 60.4 | 15.4 | 84.6 |
| D69H | 40 | 12.1 | 28.0 | 30.1 | 69.9 |
| T241G | 81 | 9.7 | 71.4 | 12.0 | 88.0 |
| T241A P203A | 85 | 12.8 | 72.3 | 15.0 | 85.0 |
| T241G P203A | 90 | 9.0 | 81.4 | 10.0 | 90.0 |
| E36A T241A | 61 | 9.4 | 51.7 | 15.3 | 84.7 |
| E36A T241G | 74 | 8.9 | 65.5 | 11.9 | 88.1 |
| Q41P T241A | 61 | 9.4 | 51.4 | 15.4 | 84.6 |

Note: the absolute configuration of C-1 was not determined; X and Y indicate the first and the second eluting diasteromer, respectively.

**Figure S22: Stereoselectivity of wt and selected mutants with (*R*)-3-methylcyclohexanone.** The tested variants were from experimental screening (T241A, D69H), rational engineering (T241A-P203A), AI-based engineering (T241G, T241A-E36A, T241A-Q41P), or hybrid engineering (T241G-P203A (highlighted in green), T241G-E36A). Reactions were carried out as described before with 25 mM (*R*)-3-methylcyclohexanone, 40 mM cyclopropylamine, 10 µM enzyme, and 1 mM $NADP^+$ at pH 8 and incubated in a horizontal shaker at 30 °C. The reported conversions were calculated based on substrate consumption and using a calibration curve with the substrate and an internal standard. Moreover, the identity of the formed product was confirmed by GC-MS. Activity values listed in the table were used for **Extended Data Figure 5G**.

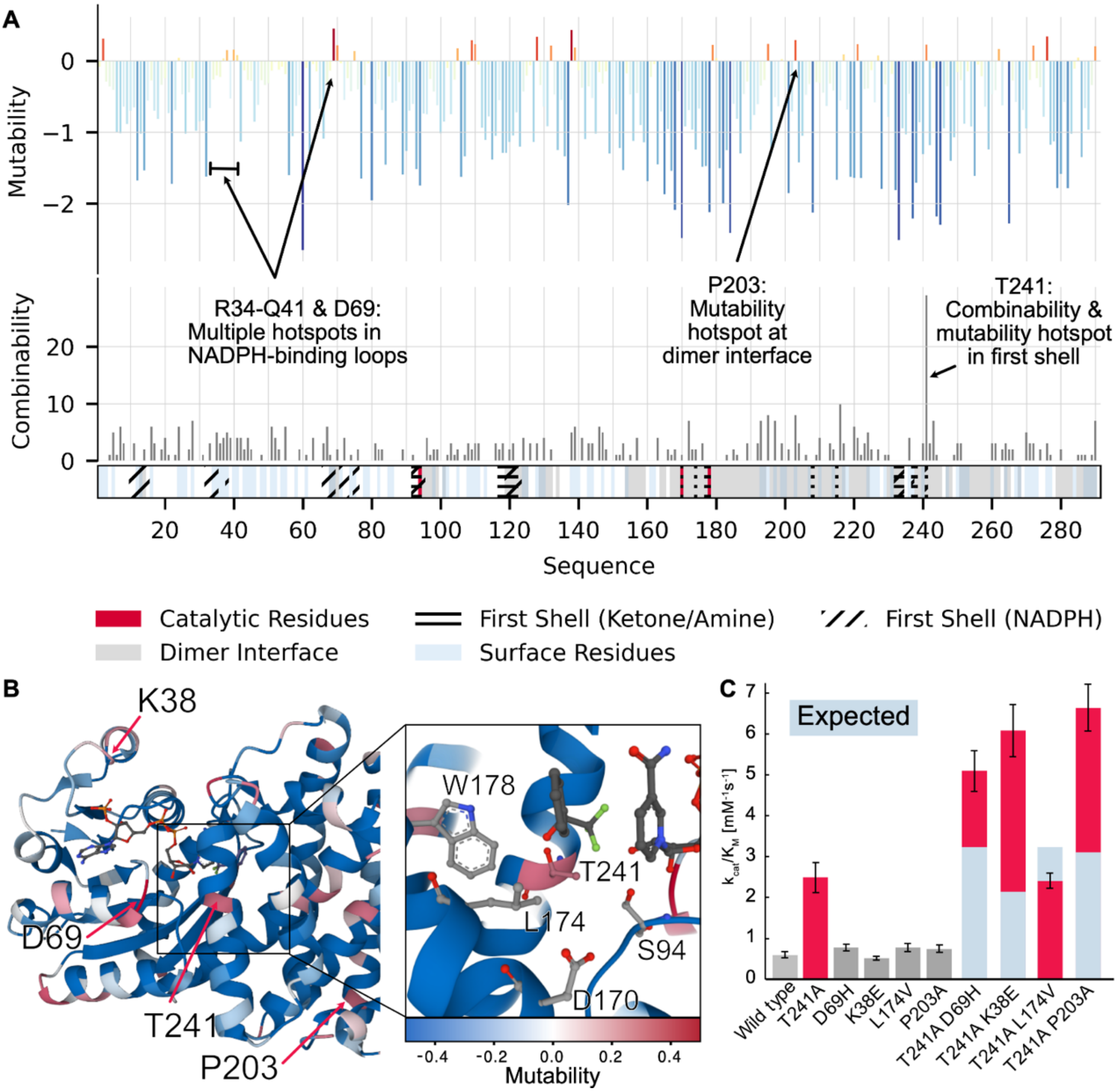

**Figure S23: A mutagenic profile of an IRED enables rational engineering.** (**A**) Median fitness per position (*mutability*) and the number of productive higher-order combinations produced per position (*combinability*) along the *Sr*IRED sequence. Proposed catalytic residues, first shell (around the ketone and amine binding pocket), surface and cofactor binding residues along with residues at the dimer interface are marked as indicated. (**B**) A map of *mutability* onto the IRED structure (5OCM) reveals hotspots e.g. around K38, D69 and at multiple positions central domain, e.g. at position P203. (**C**) Combining T241A with mutations at hotspots K38E, D69H, and P203A yields highly improving synergistic improvements (up to 11-fold $k_{cat}/K_M$).

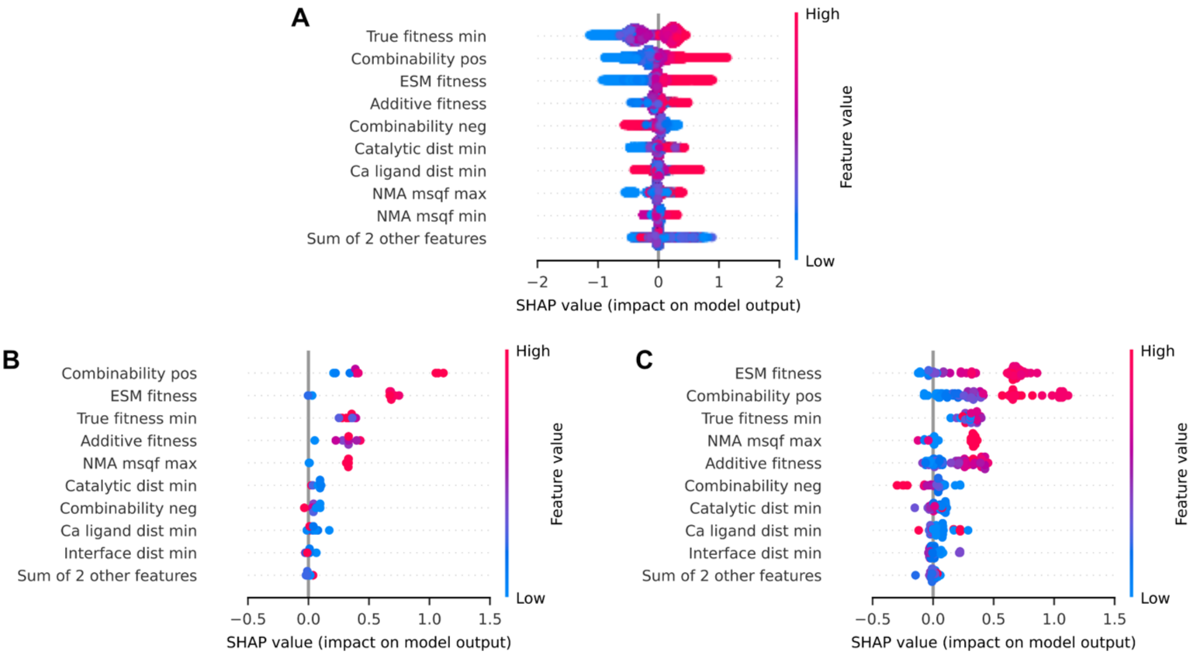

**Figure S24: Bee swarm plots of the signed SHAP values as feature importances for the double mutant model for (a) all considered double mutants (721,870 variants), (b) the top 10 and (c) the top 100 ranked predictions.** For the top predictions, high values of *Combinability* and *ESM fitness* have a particularly large effect on predictions, followed by *additive fitness* and *true fitness min*, the minimum fitness of any of the point mutants that make up the double mutant. This underscores the fact that the model is heavily relying on the assay data to select positions to mutate (*combinability pos* is a position-only feature), and on assay and language model data (a proxy for evolutionary data) to select the mutation at those positions. In comparison, rough a-priori structural information from the wildtype (*Catalytic dist min, …*) plays a minor role for the model's predictions. Note that the definition of combinability used in the double mutant model differs slightly from the one used in rational engineering for historic reasons, as discussed in the **Methods.**

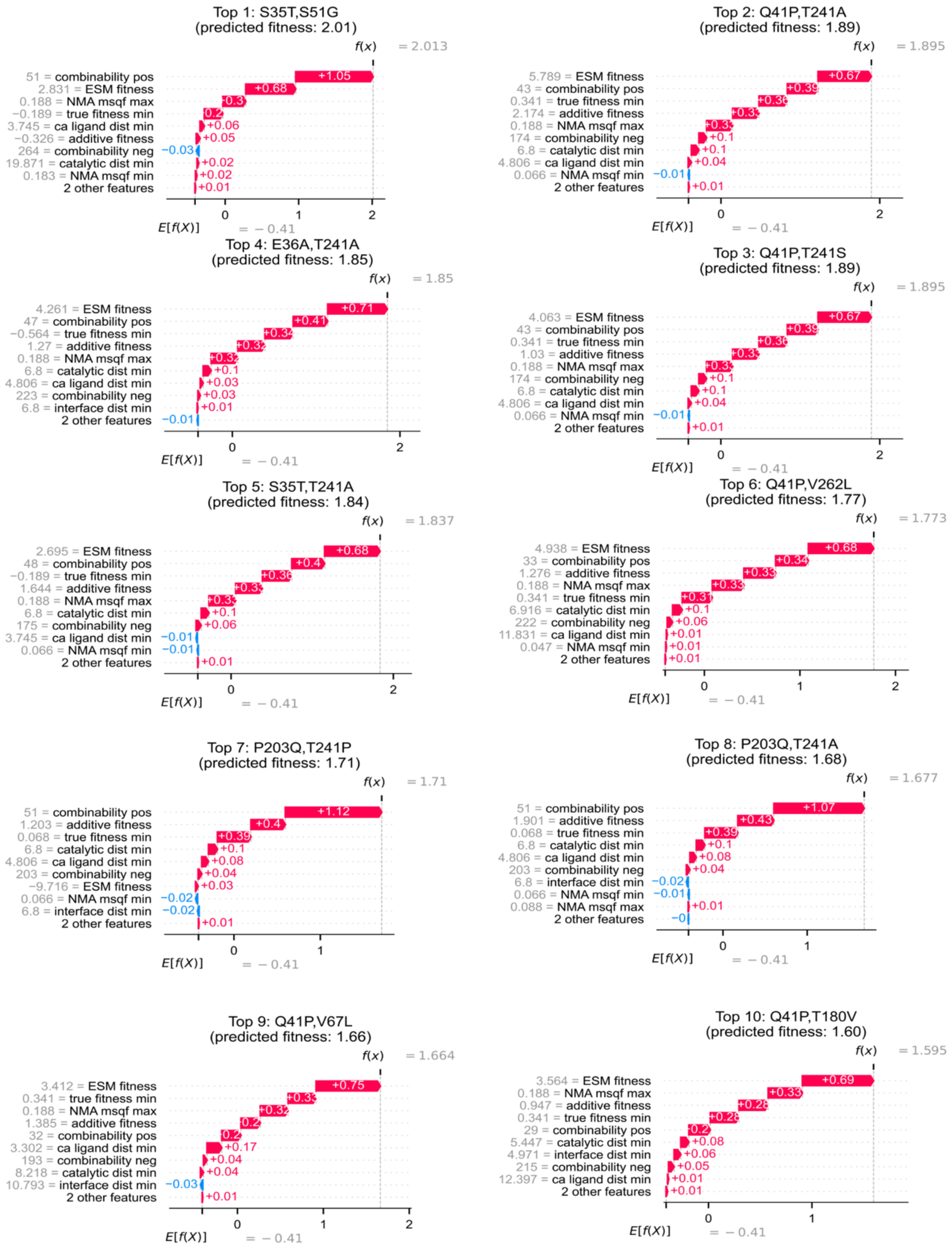

**Figure S25: Waterfall plot of the SHAP feature contributions for the top 10 predictions of the double mutant model.** The individual contributions reveal that the ESM features largely contribute to upweighting the predicted fitness of combinable mutants in the NADPH binding loop (residues 35-41) which were also found in the assay profile (**Supplementary Fig. 17**), while top ranked mutants at the positions 203 and 241, the positions of the best mutants obtained in this study, are strongly driven by assay labelled data (c.f. Top 7 and Top 8). A priori positional information from the wild-type structure on e.g. the distance between the single point mutants, the distance to the catalytic site or the surface (for which NMA mean squared fluctuations is a proxy for) appear to have a weak impact in comparison, indicating that there is little apparent broad structural bias for improving mutations.

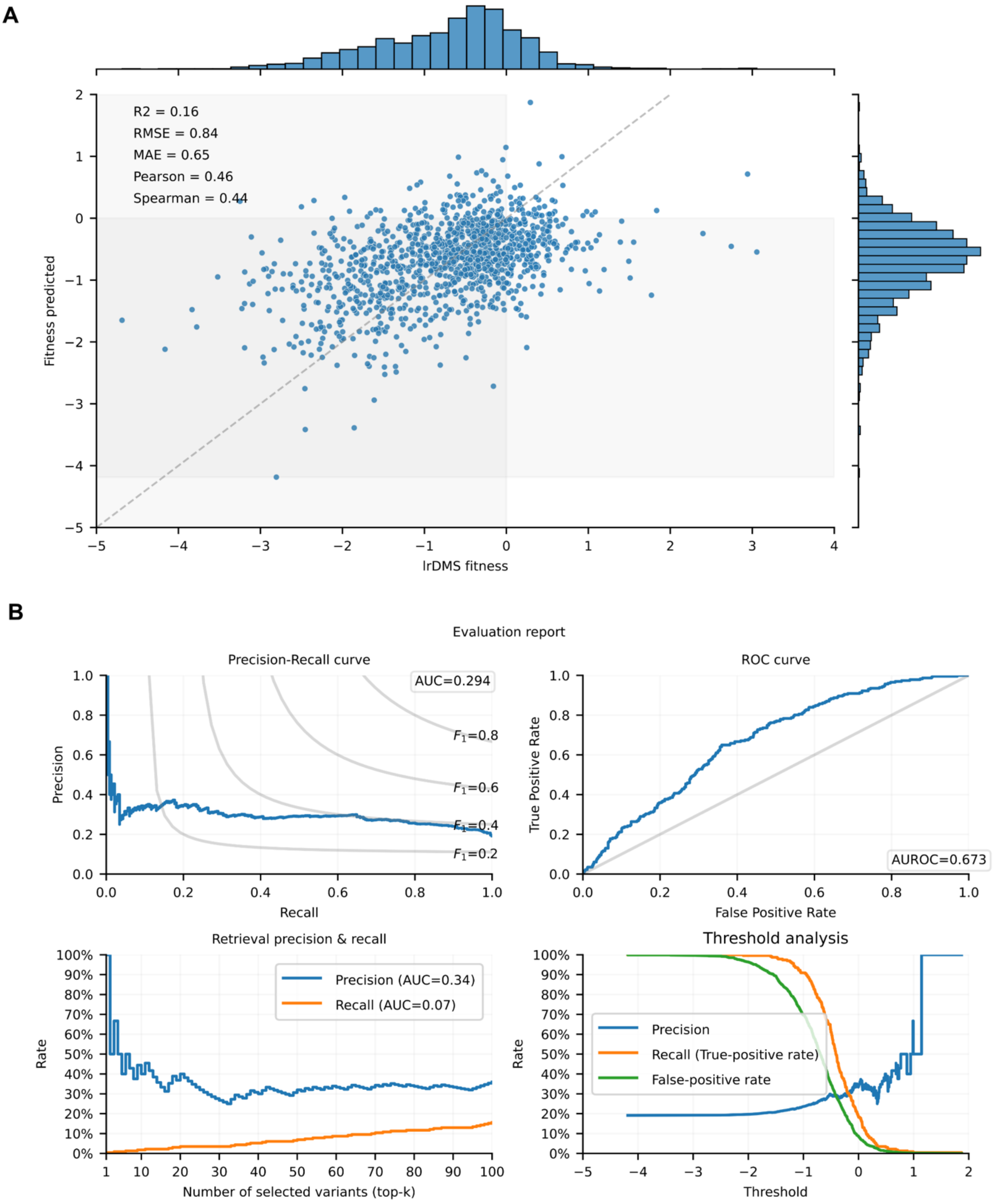

**Figure S26: Cross-validation evaluation of the single mutant model.** The single-mutant model was evaluated on all single mutant data using 10 replicas with 10-fold cross validation each (100 resulting experiments) and the predicted fitness values for each datapoint were averaged across all experiments in which the given datapoint occurred in the test set. **(A)** Scatter plot of the assay labelled ("true") fitness (x-axis) versus the model predicted fitness. The dashed grey line indicates the identity separating the regions of over- and underprediction, and the shaded grey overlays indicate regions of false positive (top left), false negative (bottom right) activity predictions vs. wildtype. Key global performance metrics are indicated in the top left corner. **(B)** Binary classification performance metrics for predicting better-than-wild type activity (fitness > 0) are plotted. The top left panel shows a precision recall curve, while the top right shows a receiver operator curve (ROC). The grey line in the ROC plot corresponds to random predictions. The bottom left plot shows the hit-rate (precision) and recall when selecting only the top-k model predictions in a retrospective analysis on all observed single mutants. Note that the hit rate from choosing at random lies at ~18% for this dataset. The bottom right plot shows the precision, recall and false positive rate as a function of the predicted fitness threshold that is applied for classification.

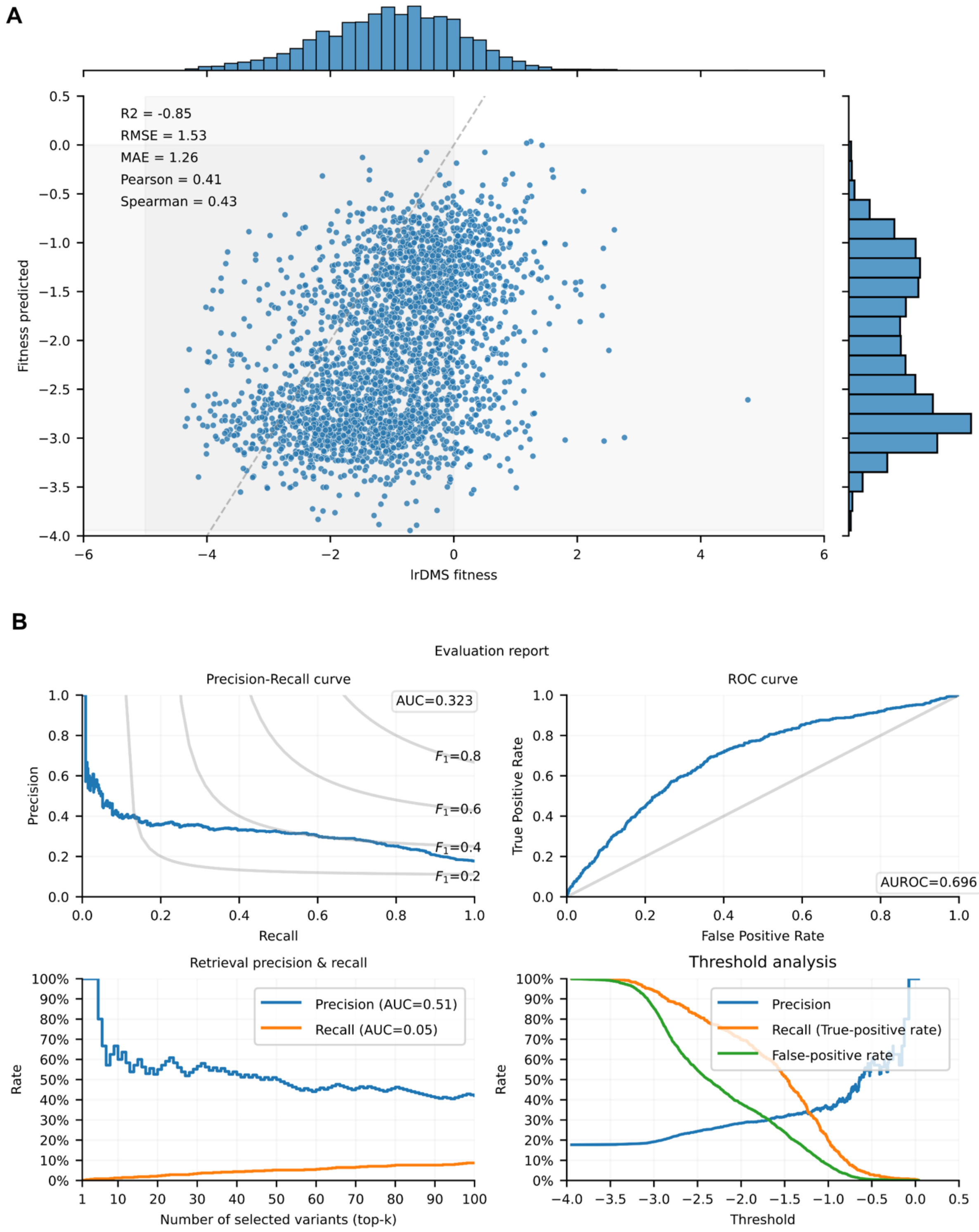

**Figure S27: Cross-validation evaluation of the double mutant model.** The double-mutant model was evaluated on all double mutant data using 6 replicas with 5-fold cross validation each (30 resulting experiments) and the predicted fitness values for each datapoint were averaged across all experiments in which the given datapoint occurred in the test set. **(A)** Scatter plot of the assay labelled ("true") fitness (x-axis) versus the model predicted fitness. The dashed grey line indicates the identity separating the regions of over- and underprediction, and the shaded grey overlays indicate regions of false positive (top left), false negative (bottom right) activity predictions vs. wildtype. Key global performance metrics are indicated in the top left corner. **(B)** Binary classification performance metrics for predicting better-than-wild type activity (fitness > 0) are plotted. The top left panel shows a precision recall curve, while the top right shows a receiver operator curve (ROC). The grey line in the ROC plot corresponds to random predictions. The bottom left plot shows the hit-rate (precision) and recall when selecting only the top-k model predictions in a retrospective analysis on all observed double mutants. Note that the hit rate from choosing at random lies at ~16% for this dataset. The bottom right plot shows the precision, recall and false positive rate as a function of the predicted fitness threshold that is applied for classification.

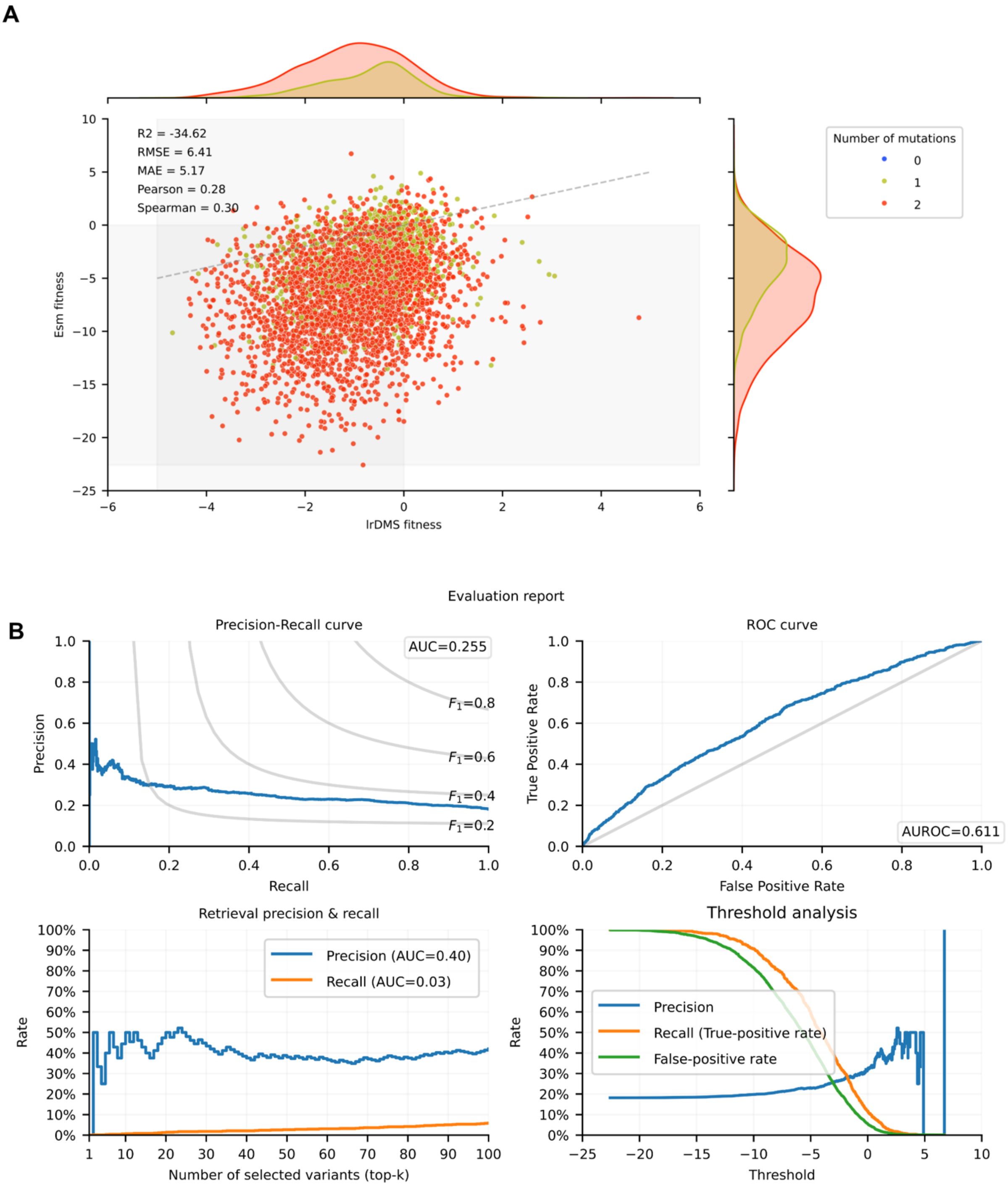

**Figure S28: Zero-shot retrospective evaluation of ESM2 (3B) on lrDMS SrIRED single & double mutant dataset.** The ESM-3B model was evaluated in zero-shot mode on all single and double mutants using PLL as ESM fitness value (**Methods**). (**A**) Scatter plot of the assay labelled fitness (x-axis) versus predicted fitness. The dashed grey line indicates the identity separating the regions of over- and underprediction, and the shaded grey overlays indicate regions of false positive (top left), false negative (bottom right) activity predictions vs. wildtype. Key global performance metrics are indicated in the top left corner. **(B)** Binary classification performance metrics for predicting better-than-wild type activity (fitness > 0) are plotted. The top left panel shows a precision recall curve, while the top right shows a receiver operator curve (ROC). The bottom left plot shows the hit-rate (precision) and recall when selecting only the top-k model predictions in a retrospective analysis. Note that the hit rate from choosing at random lies at ~18%. The bottom right plot shows the precision, recall and false positive rate as a function of the predicted fitness threshold that is applied for classification.

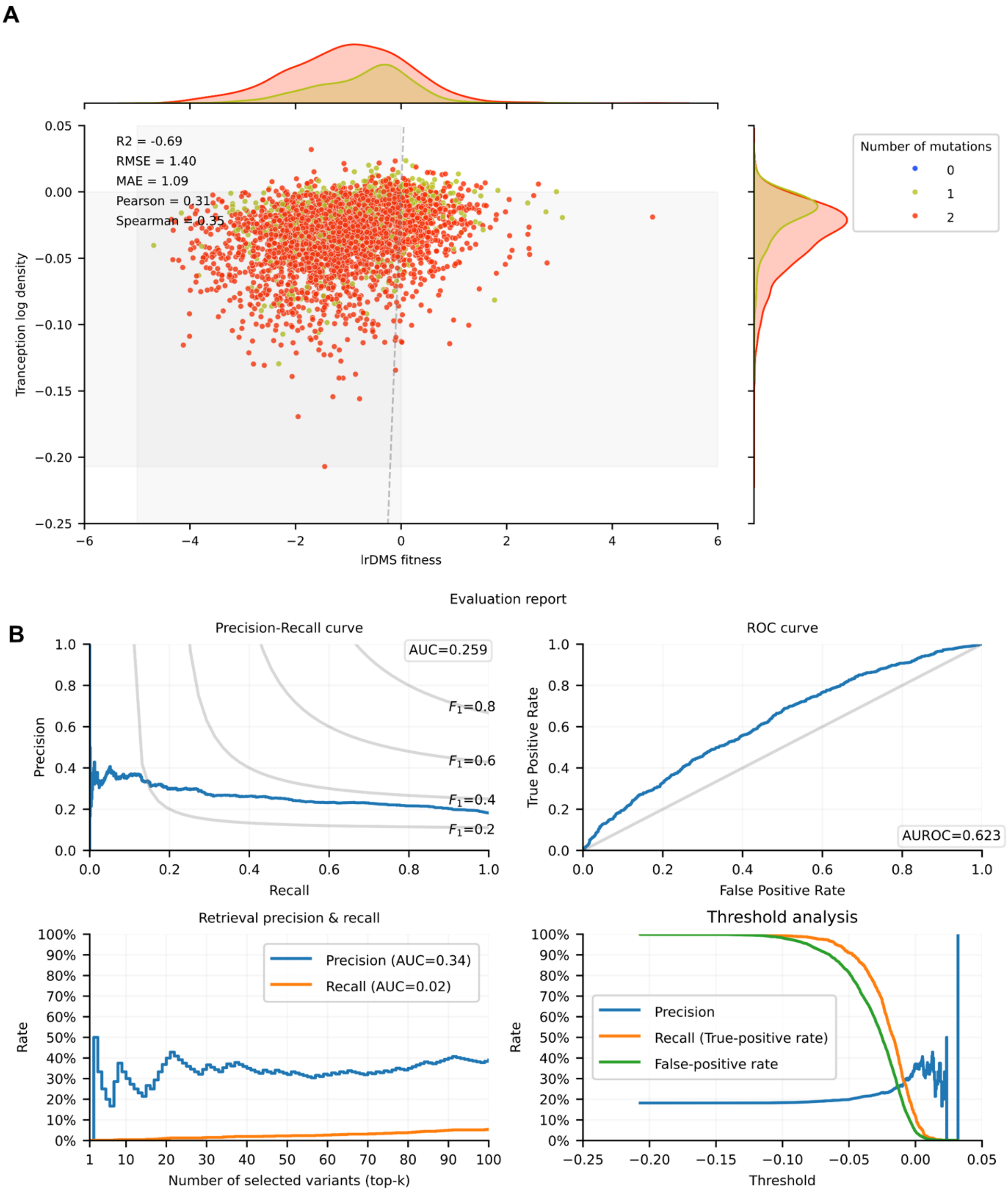

**Figure S29: Zero-shot retrospective evaluation of Tranception on IrDMS SrIRED single & double mutant dataset.** The Tranception model was evaluated in zero-shot mode on all single and double mutants using sequence log likelihood as fitness value. (**A**) Scatter plot of the assay labelled fitness versus predicted fitness. The dashed grey line indicates the identity separating the regions of over- and underprediction, and the shaded grey overlays indicate regions of false positive (top left), false negative (bottom right) activity predictions vs. wildtype. Key global performance metrics are indicated in the top left corner. **(B)** Binary classification performance metrics for predicting better-than-wild type activity (fitness > 0) are plotted. The top left panel shows a precision recall curve, while the top right shows a receiver operator curve (ROC). The bottom left plot shows the hit-rate (precision) and recall when selecting only the top-k model predictions in a retrospective analysis. Note that the hit rate from choosing at random lies at ~18%. The bottom right plot shows the precision, recall and false positive rate as a function of the predicted fitness threshold that is applied for classification.

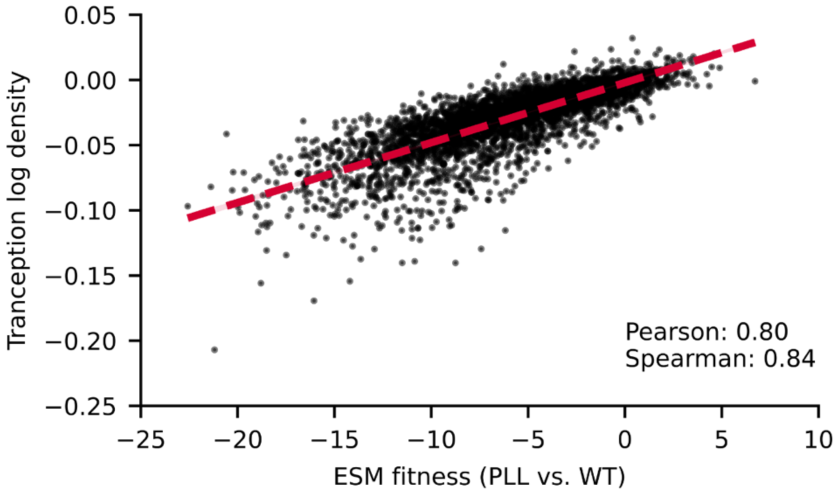

**Figure S30: Correlation analysis between Tranception log likelihoods and ESM pseudo log likelihood (PLL) vs wild type (WT).** The scatterplot between ESM[4] and Tranception[5] fitness values (PLL for ESM, sequence log likelihood for Tranception) shows that ESM and Tranception fitness predictions are highly correlated, indicating that both models likely capture the similar aspects of the underlying evolutionary patterns from related sequences.

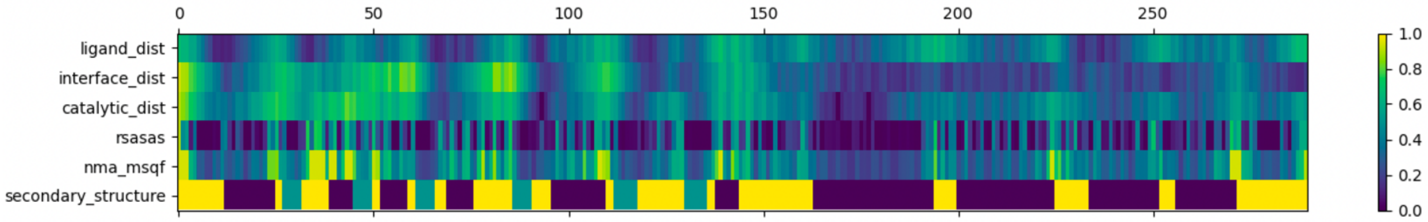

**Figure S31: Profile of structural features used in the construction of the double mutant model along the SrIRED sequence.** All structural features were computed based on the wildtype structure (PDB: 5OCM).

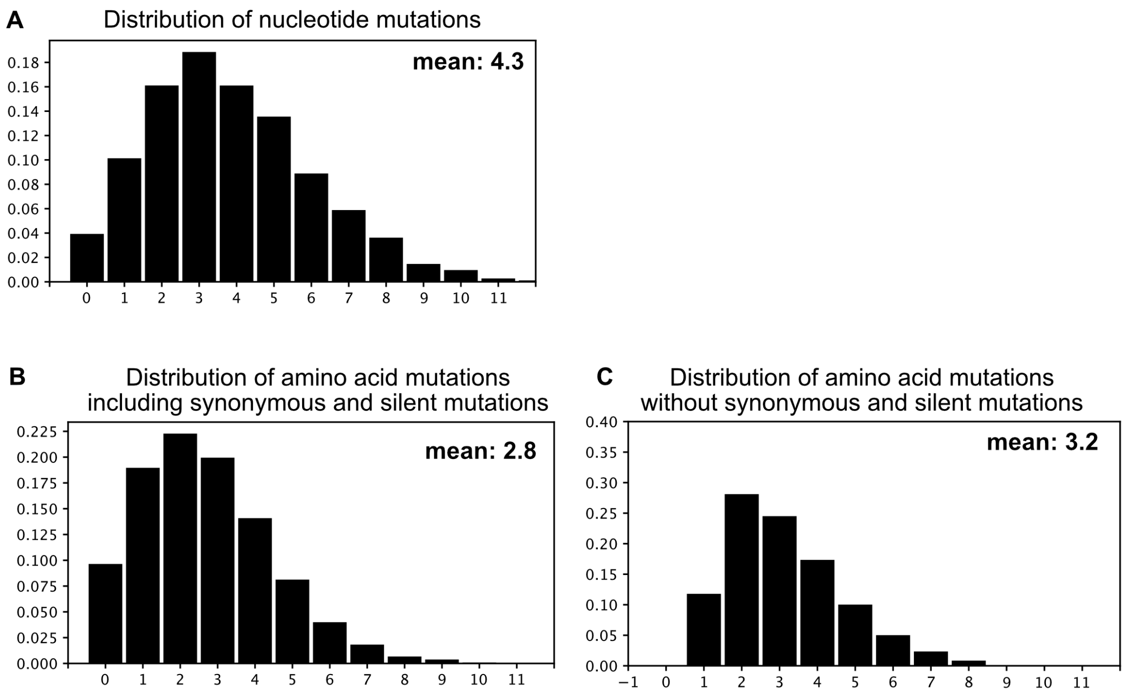

**Figure S32: Mutational load in PcIRED dictionary. (A)** Distribution of all nucleotide mutations for unique plasmids (with a unique UMI) in PcIRED in error-prone PCR library dictionary. For PcIRED we observed a slightly lower mutational load than for SrIRED (Extended data figure 2) with on average 4.3 nucleotide mutations per gene (6.0 for SrIRED). (B) Per unique plasmid we on average observed 2.8 mutations per gene, slightly more than the 2.6 mutations on average observed for SrIRED (Extended data figure 2). **(C)** Removing synonymous and silent mutations, we observe an average mutation number of 3.2 mutations per gene, slightly more than the 3.1 mutations per gene observed for SrIRED (Extended data figure 2). In total, this gives us number 935 single mutations, 2491 double mutations, 2178 triple mutations and 3169 with more than 3 mutations.

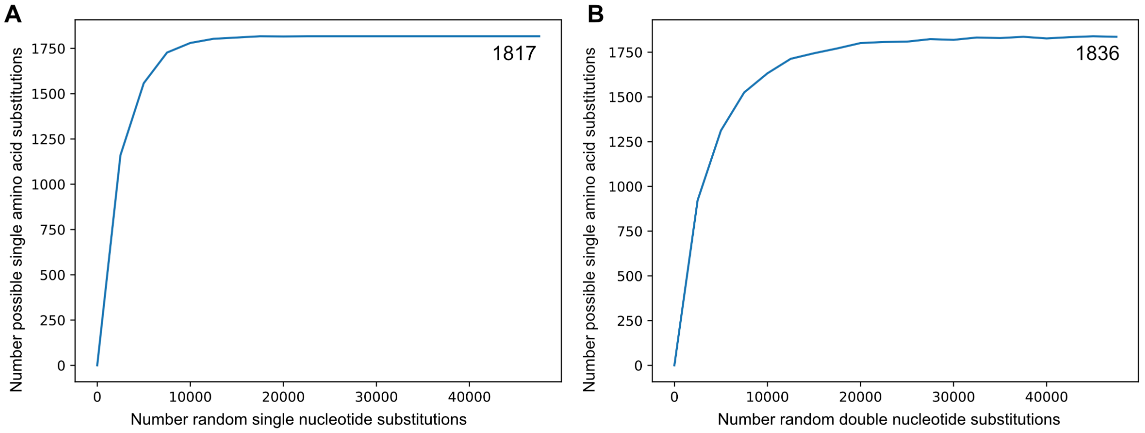

**Figure S33: Modelling the accessible single amino acid mutation space in PcIRED. (A)** Only single nucleotide mutations **(B)** also double nucleotide mutations allowed in modelling the possible space of accessible single mutations (**analogous to Extended Data Figure 2**). Similar to the 1759 maximum single mutations for SrIRED (with 290 total residues), 1817 maximum mutations were observed in PcIRED (with 296 total residues).

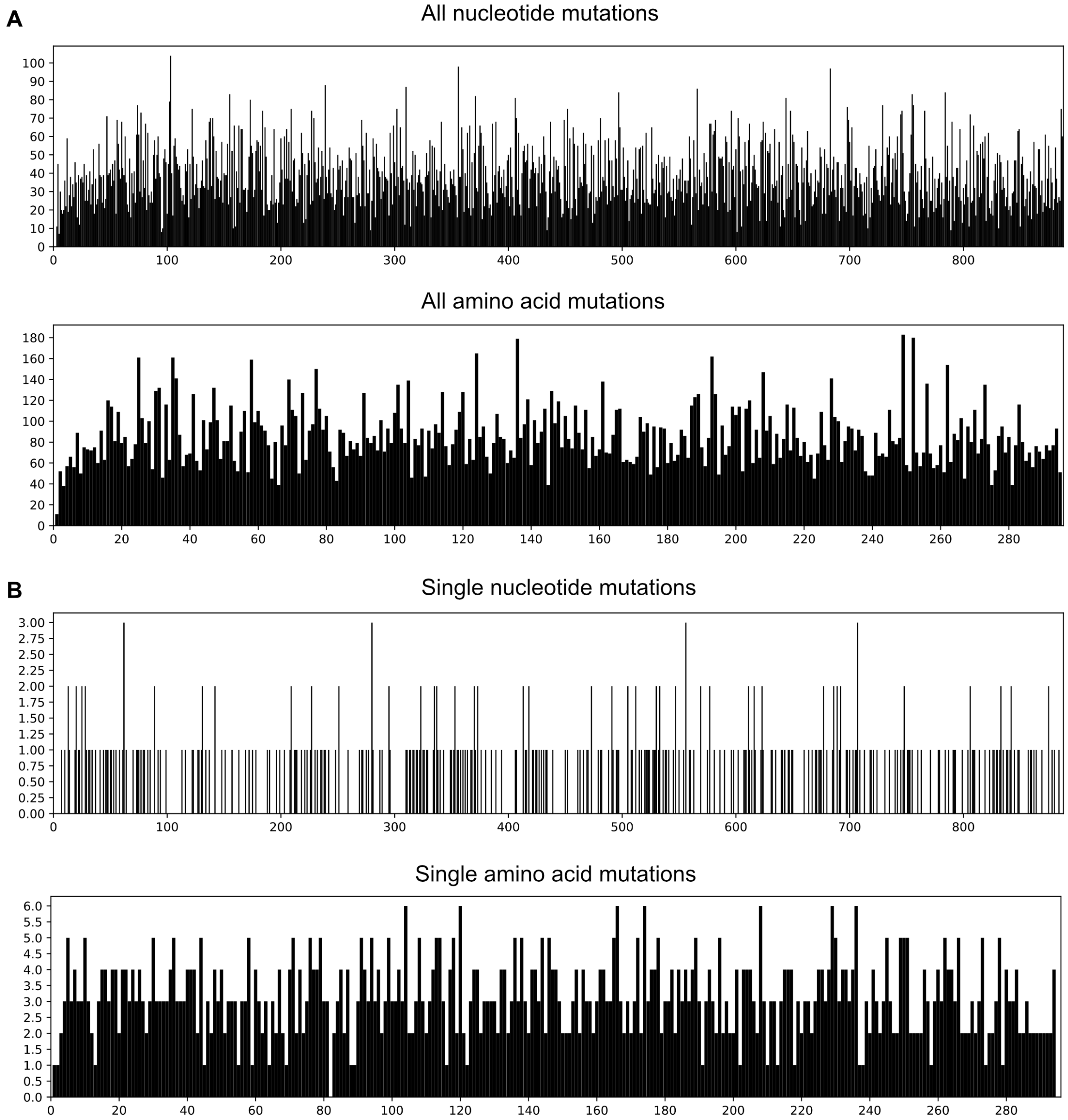

**Figure S34: Distribution of mutations across PcIRED sequence in dictionary.** A total of 12037 unique amino acid mutations were observed in the dictionary. **(A)** Distribution of nucleotide and amino acid mutations over the PcIRED sequence. Nucleotide and amino acid mutations are not equally distributed over the dictionary implying that errors caused by error-prone PCR randomisation are not evenly distributed over the sequence. However, the positions with high occurrence of nucleotide or amino acid mutations do not overlap with the calculated combinability hotspots, suggesting that those are independent of observed biases in the library before sorting. **(B)** Also single nucleotide and amino acid mutations are less equally distributed than in SrIRED. The low amount of observed single mutations can be explained by a high number of higher-order mutations including silent mutations that lead to single amino acid mutations.

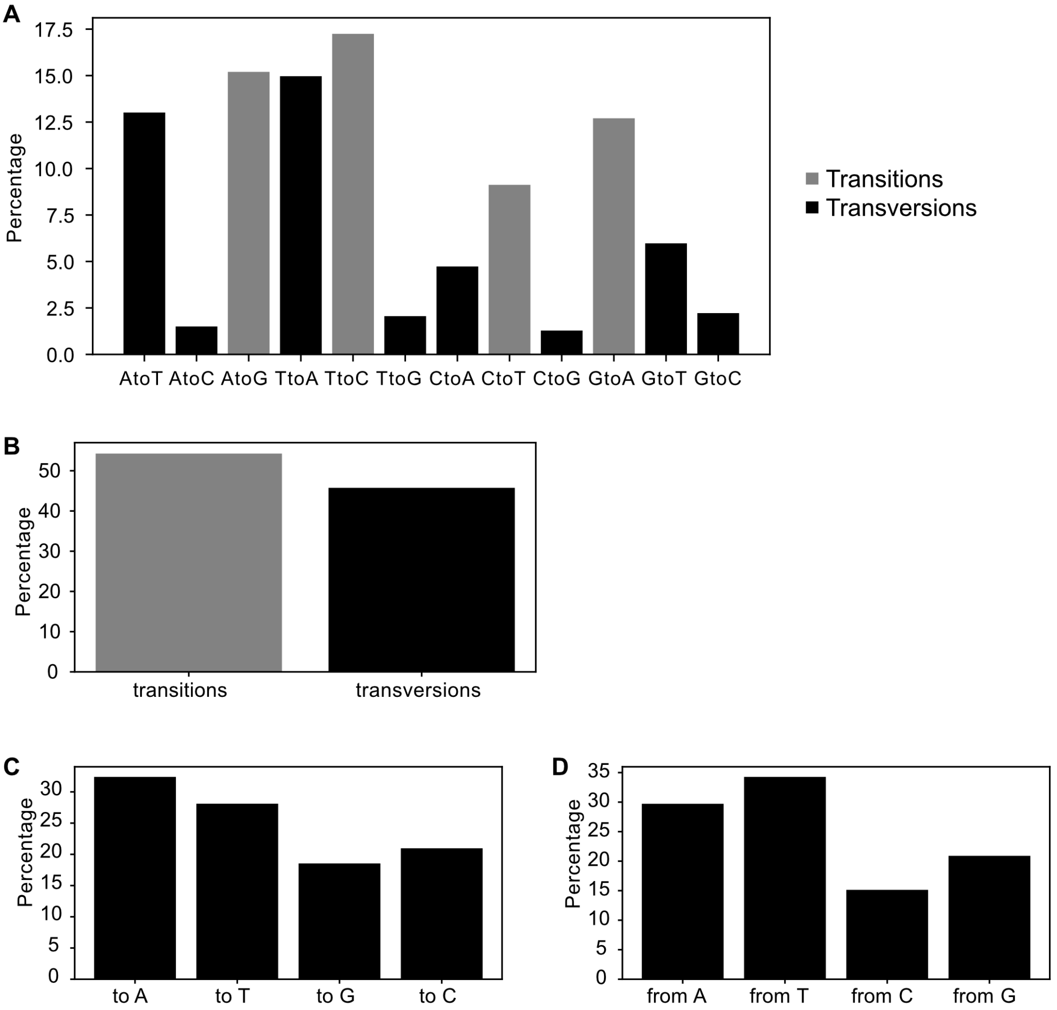

**Figure S35: Analysis of mutational bias in PcIRED epPCR library. (A)** Percentage of mutations from and to all possible nucleotides. Global trends with a bias towards transitions is similar to SrIRED (Figure S6) with a more extreme bias against mutations from A to C, T to G and C to G. **(B)** Similarly to SrIRED the overall fraction of transitions is similar to transversion despite transversions being statistically twice as likely **(C)** Similarly to SrIRED a slight bias towards A and T mutations was observed **(D)** similarly to SrIRED a slight bias towards mutations from A to T to other nucleotides was observed.

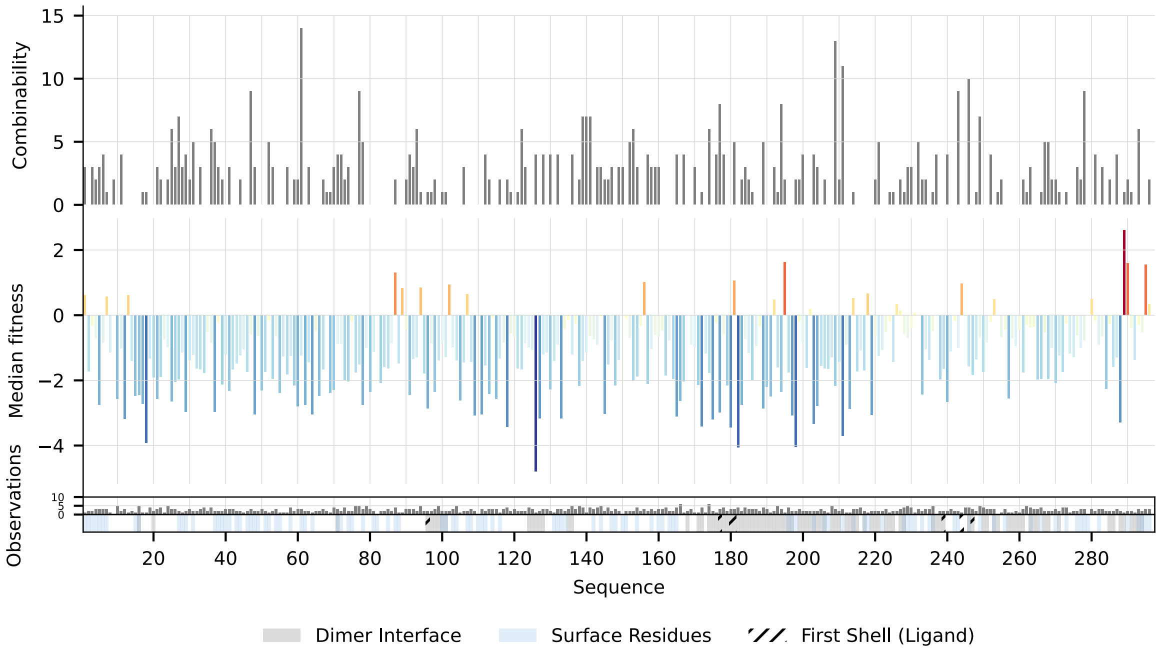

**Figure S36: A mutagenic profile of PcIRED from the microfluidic lrDMS experiment.** The number of productive higher-order combinations (*combinability**) per position, as well as the median fitness value of observed single mutants at that position (*mutability)* are plotted along the PcIRED sequence. The annotations in the bottom two lanes of the figure indicate the number of single mutant observations per position in the dataset and highlight key structural features such as surface exposed residues, residues at the dimer interface or first shell residues. The dimer interface / first shell are defined via a 6Å / 5Å distance cutoff, while surface residues are defined as residues with relative solvent accessible surface area (RSASA) larger than 0.25. Note the combinability hotspots at residue 61, as well as 209 and 211 and towards the C-terminus, as well as the mutability hotspots at the C-terminus and around positions 90, 180 and 200. Compared to the SrIRED campaign, the PcIRED campaign resulted in fewer datapoints. This is especially visible for the single mutants (SrIRED: 1514, PcIRED: 750), with an average coverage of 2.5 mutations per position, which we hypothesize was also a limiting factor for the performance of our single mutant model.

*:Instead of "(*fitness > 0 by 1 std. dev.) & (fitness > expected additive by 1 std. dev.)"* for SrIRED, we used a slightly less stringent filter of "fitness > 2.0" for higher order variants that go into the combinability calculation for the PcIRED campaign. We chose this relaxation after observing that the SrIRED filters would have led to an unnecessarily sparse combinability trace for PcIRED, due to the higher uncertainties in that dataset (discussed in more detail in **Supplementary Fig S38**). For more detail, refer to the **Methods**.

**A**

| | variant | fitness_lr_combo | fitness_esm | pos_n_mut_obs_all | pos_n_replicas_all |
|---|---|---|---|---|---|
| 0 | A135I | 2.935271 | -3.627508 | 5 | 1 |
| 1 | W161T | 2.777232 | 2.573117 | 3 | 1 |
| 2 | V102W | 2.538444 | -5.261409 | 3 | 1 |
| 3 | V289E | 2.451505 | -4.618092 | 1 | 1 |
| 4 | W161C | 2.265623 | -0.788453 | 3 | 1 |
| 5 | W161P | 2.226310 | 1.450728 | 3 | 1 |
| 6 | V289G | 2.197374 | -2.818598 | 1 | 1 |
| 7 | G195R | 2.135817 | 1.954771 | 2 | 4 |
| 8 | V289D | 2.110194 | -5.804620 | 1 | 1 |
| 9 | A135L | 2.035465 | -2.779408 | 5 | 1 |
| 10 | G195Q | 2.010325 | 2.241419 | 2 | 4 |
| 11 | G195E | 2.001048 | 2.252983 | 2 | 4 |
| 12 | G195K | 1.957391 | 2.105302 | 2 | 4 |
| 13 | I251S | 1.850407 | -0.823468 | 4 | 1 |
| 14 | G195H | 1.839689 | 0.490020 | 2 | 4 |

**B**

| | variant | fitness_lr_combo | fitness_esm | pos_n_mut_obs_all | pos_n_replicas_all |
|---|---|---|---|---|---|
| 0 | A135I | 2.935271 | -3.627508 | 5 | 1 |
| 28 | P87G | 1.656170 | 1.595387 | 7 | 3 |
| 51 | A168N | 1.364962 | -2.152996 | 4 | 3 |
| 64 | A181S | 1.181343 | -0.379412 | 6 | 5 |
| 90 | K249G | 1.016656 | 0.338088 | 9 | 4 |

**Figure S37: Top single mutant model predictions for the *Pc*IRED campaign**. We trained the same single mutant model as for the *Sr*IRED campaign (c.f. **Methods**) on the single mutations in the *Pc*IRED campaign, the resulting top 15 predictions of which are shown in **A**. The PcIRED campaign observed fewer single mutants (750) than the *Sr*IRED campaign, leading to an average number of 2.5 single mutants per position (see **Fig S36**). In addition to that, those observations came with higher levels of uncertainty, as mutations to many positions were only seen in one replica (e.g. the positions of predictions 0-6 in **A**), leading to high variance. To mitigate these uncertainties *post-hoc*, we select the 5 mutants to assay in low throughput from among the Top 100 predictions, as the first mutants that satisfy the following criteria:

1. At least 3 single mutants should have been observed at the proposed position (*pos_n_mut_obs_all*)
2. Mutations at the proposed position should have been sampled in at least 3 (of 5) replicas. (*pos_n_replicas_all*)
3. Mutations to proline (P) were excluded.
4. When a position was selected, we exclude other mutations at that position to increase the diversity positions tested at low throughput.

As a single exception to these criteria, we added the top predicted mutant A135I. The resulting top 5 mutants selected for low throughput characterisation are summarized in **B** (c.f. Table **S6** for characterization data), together with their rank in the unfiltered prediction. 2 out of 5 of these predictions resulted in wildtype-like properties (**Extended Data Figure 11**) and one prediction (A181S) showed a 2-fold improved $k_{cat}$, but this was accompanied by an increased $K_M$. While this is still significantly better than what would be expected from random mutagenesis, we attribute the moderate success of the single mutant model for *Pc*IRED as compared to *Sr*IRED to the above mentioned, increased uncertainties, which made the model more susceptible to riskier outliers (see **A,** e.g. A135I, W161T, V102W, …) and introduced higher variance into our small set of 5 low throughput assayed variants.

**A**

| | variant |
|---|---|
| 0 | S178G,K230E |
| 1 | S178G,K249N |
| 2 | P87S,S178I |
| 3 | S178G,P295R |
| 4 | S178G,P295S |
| 5 | S178C,K273E |
| 6 | P87T,S178C |
| 7 | G117S,S178G |
| 8 | M114K,S178I |
| 9 | Q52L,S178C |
| 10 | S178G,K249Q |
| 11 | I138L,K236E |
| 12 | F30Y,P54Q |
| 13 | G134S,S178G |
| 14 | K249N,G267S |
| 15 | E144D,G195S |

**B**

| | variant | combinability_pos | additive_fitness | esm_fitness |
|---|---|---|---|---|
| 0 | Q228R,K249N | 29 | 1.444100 | 1.440451 |
| 1 | I91L,K249N | 29 | -0.360383 | 0.588993 |
| 2 | F25S,K249N | 29 | -1.241989 | -2.294307 |
| 3 | H199L,K249N | 28 | 0.876799 | 1.773486 |
| 4 | I91L,Q228R | 28 | -1.045086 | -0.174499 |
| 5 | F25S,Q228R | 28 | -1.926693 | -3.111042 |
| 6 | F25S,I91L | 28 | -3.731176 | -3.959066 |
| 7 | N36S,K249N | 27 | 1.730353 | -4.338953 |
| 8 | K249N,K296R | 27 | 1.576461 | 0.164372 |
| 9 | K249N,F252S | 27 | 0.938234 | 0.911817 |
| 10 | K249N,E283V | 27 | 0.647052 | -2.742211 |
| 11 | H199Q,Q228R | 27 | 0.639364 | -0.584795 |
| 12 | Q16L,K249N | 27 | -0.407892 | 3.099416 |
| 13 | I91L,H199Q | 27 | -1.165119 | -1.441975 |
| 14 | F25S,H199Q | 27 | -2.046725 | -4.339483 |
| 15 | R61L,K249N | 26 | 2.122588 | -0.105477 |

**C**

| | variant | esm_fitness |
|---|---|---|
| 0 | P87S,M114K | 8.173730 |
| 1 | M114K,L151P | 8.070370 |
| 2 | P87S,L151P | 7.812582 |
| 3 | L151P,K273E | 7.271415 |
| 4 | M114K,K273E | 7.270143 |
| 5 | P87T,M114K | 7.178652 |
| 6 | P87S,K273E | 7.147638 |
| 7 | Q52L,M114K | 7.054793 |
| 8 | Q52L,L151P | 6.963603 |
| 9 | H120D,L151P | 6.933115 |
| 10 | P87T,L151P | 6.908386 |
| 11 | S66T,M114K | 6.893129 |
| 12 | M114K,H120D | 6.855016 |
| 13 | Q52L,P87S | 6.847161 |
| 14 | S66T,L151P | 6.818518 |
| 15 | P87S,H120D | 6.792421 |

**Figure S38: Double mutant model predictions for the *Pc*IRED campaign.**
Top predictions for *Model 1* (**A**), *Model 2* (**B**) and ESM2-650M alone (**C**). Highlighted variants were selected for low throughput characterisation.

Motivated by the strong performance in the *Sr*IRED campaign, we also trained the double mutant model on the *Pc*IRED data. *Model 1* (**A**) represents the same double-mutant model used in the *Sr*IRED campaign with a few minor modifications relevant to *Pc*IRED:

1. We derived the structural features for *Pc*IRED from an AlphaFold3[6] prediction of the wild type sequence, as there is no deposited wild type structure available in the PDB.
2. For reasons explained below, we weighted the training examples in accordance with their estimated standard deviations to better account for noise in the dataset. Hence, loss values for variants with more confident fitness predictions were upweighted compared to noisier datapoints.

The top 5 predictions of *model 1* were selected and tested in low throughput, and 4/5 predictions yield wild type like properties (detailed results in **Extended Data Figure 11**), but improvements were comparatively small. This stands in contrasts to the case of *Sr*IRED, where the same model achieved an over 7-fold improvement in $k_{cat}$ among just 5 predictions.

We attribute the apparent difficulty to bring about larger improvements to two reasons:
(1) The *Pc*IRED dataset has a higher noise level due to the 18-fold lower starting activity of *Pc*IRED compared to *Sr*IRED. The lower starting activity necessitated screening close to the detection limit of absorbance activated droplet sorting (AADS)[7] thereby reducing the signal to noise. While this made extrapolation based on the (AADS) lrDMS data for *Pc*IRED more challenging, future lrDMS campaigns with low starting activity are likely to significantly benefit from a recently developed assay that can be used to link IRED activity to a fluorescence readout, decreasing the limit of detection ~1000-fold[8]. (2) Furthermore, the *Pc*IRED data exhibits significantly fewer *mutation-stacked* double mutants, i.e. double mutants for which both constituting single mutants were observed (PcIRED: 490, SrIRED: 3902). This is relevant, because, even though all double and higher-order mutants factor into the computation of combinability, only mutation-stacked doubles are used as labelled datapoints for training the double mutant model. Paired with the higher noise level, supervised model training on these 490 noisier datapoints is more susceptible to outliers. As a consequence, *model 1* learns to rely more heavily (but not exclusively, about ~33% each) on a priori evolutionary (ESM) and structural biases (see **Supplementary Fiures 39 and 40**). Indeed this is consistent with what we would expect based on the learning curves for *Sr*IRED in **Extended Data Figure 8**, which, for a dataset of this size, suggest that evolutionary features (*ESM*) hold similar predictive power to the assay-derived features. And whilst these general biases were shown to find variants with wild type like enzymatic properties[9], the model struggles to find variants that

improve above these – a feat for which the more contextually rich (exact targeted reaction, pH, buffer, etc) assay labelled data is most relevant.

We therefore conjectured that we would need to further control for noise to avoid overfitting and predict meaningful improvements from the *Pc*IRED data. Inspired by the interpretability analysis of the *Sr*IRED data (**Supplementary Figure 24 & Extended Data Figure 9**) and the salient assay-derived features identified therein, we hypothesised that a baseline that is solely based on assay derived features (*combinability* and *additive fitness*) for picking positions to mutate and only uses evolutionary information (ESM2 650M) as a rejection criterion should do well. We call this baseline *model 2* (**B**), and it simply picks the positions to mutate first based on the summed *combinability* of the considered positions (the more noise-robust property of the data) and then suggests the variant with the highest *additive fitness* of the single mutants at those positions. If the ESM predicted fitness is non-negative, the mutant is selected for assaying. We selected the top 4 variants from this automated procedure (**B**) and found that one variant increased $k_{cat}$ by 2-fold, while the other 3 variants reduced $K_M$ by 2-3-fold, leading to a clearly improved catalytic efficiency ($k_{cat}/K_M$) for 3 of 4 predictions, with the best variant boosting $k_{cat}/K_M$ by 3-fold (detailed results in **Extended Data Figure 11)**, which exceeds the achievements by expert-guided rational engineering. Please note, the preponderance of $K_M$ improvements for *Pc*IRED (where screening happened at $K_M$) vs $k_{cat}$ improvements for SrIRED (where screening happened at substrate saturation) is in line with the "*you get what you screen for*" adage in directed evolution and highlights how lrDMS at different screening conditions may in the future serve the field not only as engineering tool to find better hits, but as a unique tool to elucidate the web of mutations that influence different catalytic parameters in an enzyme, ushering in a new paradigm of data-driven enzymology.

The remarkable success of *model 2* underscores the usefulness of robust summary statistics from an lrDMS study as powerful tool for engineering biocatalytic parameters even in challenging, high-noise scenarios. This is especially impressive, because *model 2* was almost exclusively based on assay features and only relied on evolutionary information to reject overly risky suggestions. This points to significant potential for developing robust algorithms that better leverage assay derived data jointly with evolutionary and structural information in high noise scenarios.

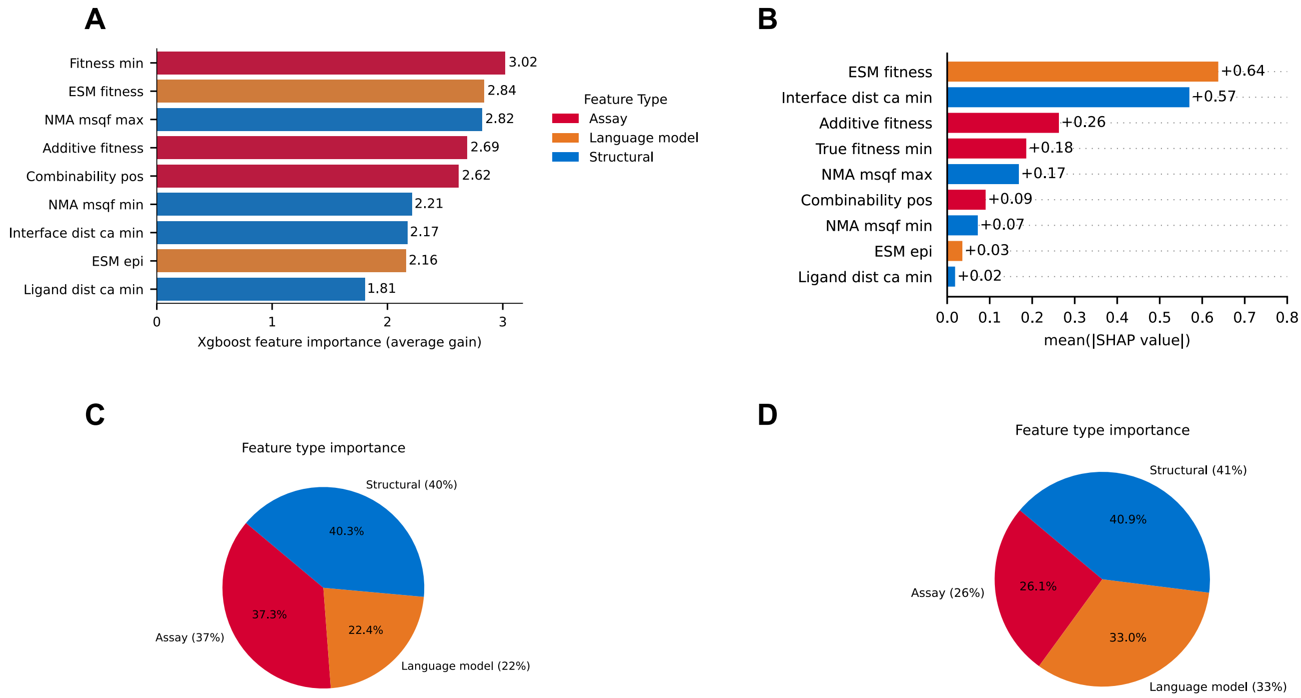

**Figure S39 Feature importance analysis for the double mutant *model 1* in the PcIRED campaign.** While assay derived features are still attributed about one third of impact on the model's predictions, the PcIRED model relies more heavily (but not exclusively, about ~33% each) on a priori evolutionary (ESM) and structural biases than was the case *Sr*IRED (compare **Extended Data Figure 9**). This is consistent with what would be expect based on the learning curves for *Sr*IRED in **Extended Data Figure 8**, which, for a dataset of this size (490 mutation-stacked double mutants), suggest that evolutionary features (*ESM*) hold similar predictive power to the assay-derived features. For more details and analysis of these trends see **Supplementary Figure 38.**

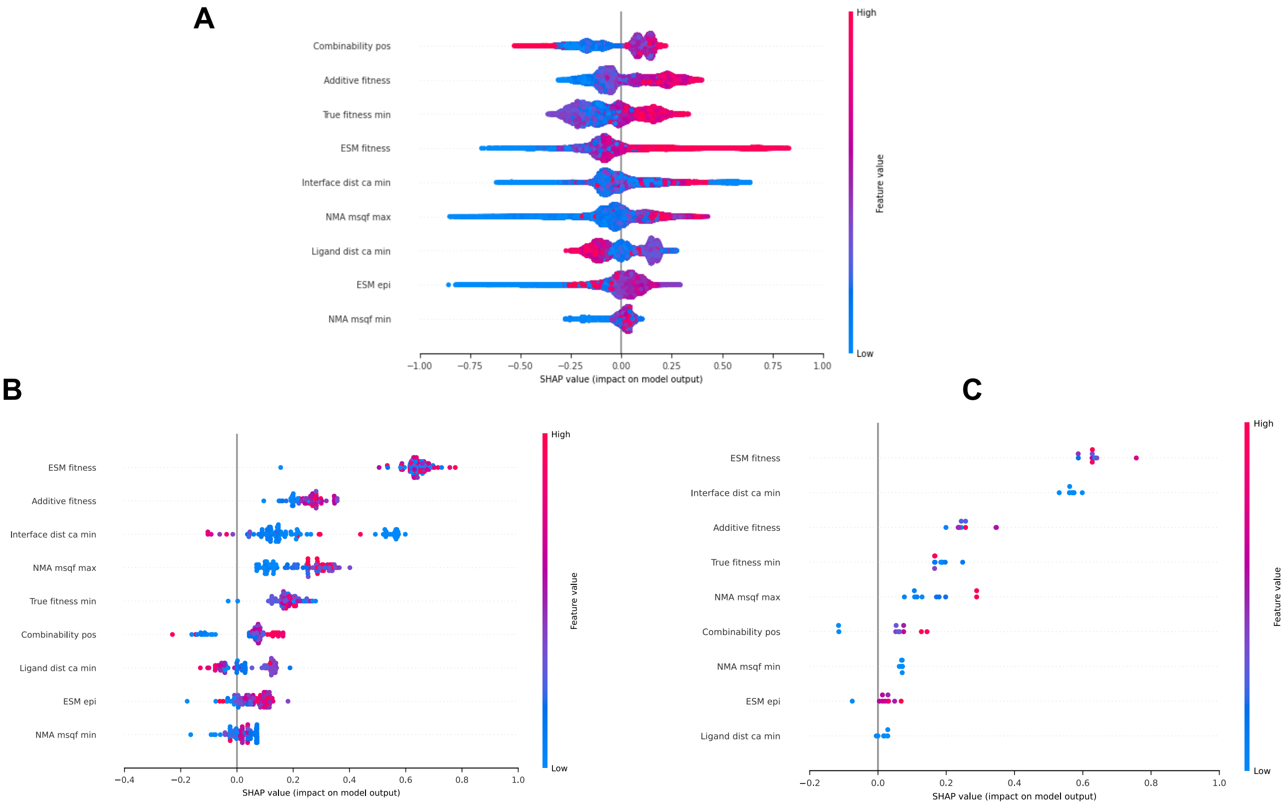

**Figure S40: Bee swarm plots of the signed SHAP values as feature importances for the PcIRED double mutant *model 1* for (a) all considered double mutant predictions (279,610 variants), (b) the top 100 and (c) the top 10 ranked predictions.** After training the model, high values of *ESM fitness* and proximity to the interface (low *interface dist ca min*) contribute most strongly to selecting the top few variants. This positional bias towards the interface that the model picks up from training on double mutant data is consistent with the mutagenic profile based on single mutants in **Supplementary Figure 36**, where the C-terminal domain – which makes up the interface – is found to be more mutable. Beyond these evolutionary (*ESM*) and structural (*interface dist*) biases, the assay derived features *additive fitness* (the expected fitness based on single mutant fitness values assuming additivity), *true fitness min* (the minimum fitness of any of the point mutants that make up the double mutant) and *combinability pos* (position dependent positive combinability) play a role in the model's selection of top mutants. In comparison to the SrIRED campaign, the PcIRED double mutant model therefore relies more on a-priori evolutionary and structural features for reasons discussed in **Supplementary Figure 38.**

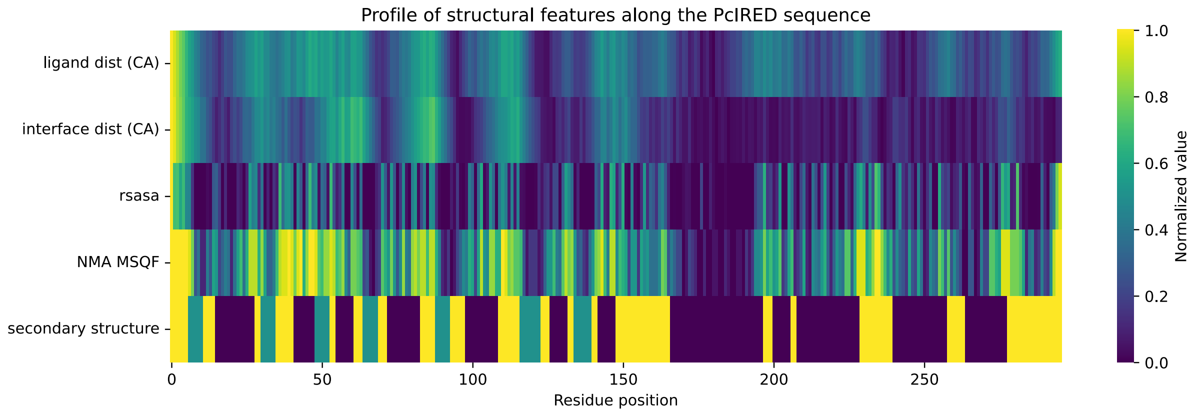

**Figure S41: Profile of structural features used in the construction of the double mutant model along the PcIRED sequence.** All structural features were computed based on the AlphaFold3[6] predicted structure of the wildtype PcIRED sequence. For ligand distance, interface distance & rsasa, the value range (min, max) was re-scaled to the range (0, 1). For NMA mean squared fluctuations we cut the distribution into 20 quantiles (to avoid over-saturation due to high flexibility of the free loop at the N-terminus). To plot the secondary structure on the same axis, we assign alpha helix = 0, beta sheet = 0.5, loop = 1.

### Supplementary Tables

**Table S1: Oversampling of the *Sr*IRED library in screening and next generation sequencing.** 'Droplets screened' refers to the number of droplets assayed in each AADS sample and 'droplets sorted' refers to the number of droplets selected based on the used selection threshold. The 'oversampling' constant refers to the used 60,000 membered *Sr*IRED variant library which was encapsulated with 50% occupancy using the Poisson distribution. NGS reads and depth are calculated for the 'input' library before sorting and the 'output' library after sorting where 'reads' refers to the total number of NGS reads for each sample and 'depth' refers to the coverage of the input library (60,000 variants) and the output library (number of sorted droplets) with NGS reads.

| Sample name | Droplets screened [x $10^6$] | Droplets sorted | Oversampling (with 50% occupancy) | NGS reads input [x $10^6$] | NGS depth input | NGS reads output [x $10^6$] | NGS depth output |
|---|---|---|---|---|---|---|---|
| 3h_1 | 0.9 | 16,738 | 7.5 | 8.0 | 133 | 3.8 | 227 |
| 3h_2 | 1.4 | 26,278 | 11.7 | 10.3 | 171 | 4.0 | 152 |
| 3h_3 | 1.2 | 17,314 | 10.0 | 11.6 | 190 | 6.6 | 390 |
| over_night_1 | 1.2 | 13,859 | 10.0 | 6.9 | 116 | 2.4 | 174 |
| over_night_2 | 1.0 | 11,868 | 8.3 | | | 2.1 | 112 |
| over_night_3 | 1.2 | 13,190 | 10.0 | | | 2.4 | 182 |

**Table S2: Summary table - Characterization of all *Sr*IRED mutants with cyclohexanone 1 cyclopropylamine a.**

| Variant | Lysate activity** [fold-change] | Lysate activity error | Soluble expression [fold-change] | $T_M$ [°C] *** | $T_M$ error | $k_{cat}$ [$s^{-1}$] * | $k_{cat}$ error | $K_M$ [mM] * | $K_M$ error | $k_{cat}/K_M$ [$s^{-1}mM^{-1}$] * | $k_{cat}/K_M$ error |
|---|---|---|---|---|---|---|---|---|---|---|---|
| wt | | | | 38.5 | 0.6 | 2.11 | 0.08 | 3.52 | 0.44 | 0.60 | 0.08 |
| T241A | 1.29 | 0.49 | 0.78 | 50.0 | 0.7 | 5.77 | 0.24 | 2.31 | 0.33 | 2.49 | 0.37 |
| T241S | 2.79 | 0.67 | 1.23 | 37.6 | 0.6 | 6.02 | 0.39 | 2.31 | 0.51 | 2.61 | 0.61 |
| M120L | 3.66 | 0.88 | 1.01 | 33.3 | 0.1 | 2.08 | 0.09 | 5.34 | 0.71 | 0.39 | 0.05 |
| L174V | 3.84 | 1.24 | 0.65 | 27.6 | 0.2 | 10.72 | 0.54 | 13.76 | 1.52 | 0.78 | 0.09 |
| D69H | 3.25 | 1.05 | 1.14 | 39.5 | 0.6 | 4.09 | 0.15 | 5.26 | 0.57 | 0.78 | 0.09 |
| D69G | 5.74 | 1.77 | 1.42 | 38.8 | 0.6 | 3.26 | 0.08 | 6.00 | 0.43 | 0.54 | 0.04 |
| D236G | 2.53 | 0.72 | 1.43 | 27.9 | 0.4 | 2.01 | 0.08 | 3.08 | 0.42 | 0.65 | 0.09 |
| P203A | 0.15 | 0.05 | n.d. | 40.8 | 0.2 | 5.02 | 0.22 | 6.70 | 0.79 | 0.75 | 0.09 |
| P203S | 1.60 | 0.39 | 0.92 | 43.7 | 1.7 | 2.49 | 0.05 | 3.25 | 0.21 | 0.77 | 0.05 |
| D267G | 5.38 | 1.17 | 1.90 | 43.3 | 0.6 | 2.73 | 0.07 | 4.27 | 0.32 | 0.64 | 0.05 |
| K38E | 0.55 | 0.15 | 0.92 | 37.7 | 1.1 | 3.24 | 0.11 | 6.27 | 0.56 | 0.52 | 0.05 |
| D5H | 3.02 | 1.14 | 1.78 | 39.6 | 0.1 | 1.91 | 0.11 | 5.10 | 0.87 | 0.38 | 0.07 |
| V216I | 1.11 | 0.32 | n.d. | 38.8 | 0.9 | 3.48 | 0.11 | 4.29 | 0.39 | 0.81 | 0.08 |
| S51I | 1.18 | 0.24 | 0.84 | 41.2 | 0.6 | 1.92 | 0.05 | 3.35 | 0.31 | 0.57 | 0.06 |
| T193A | 0.77 | 0.28 | n.d. | 33.7 | 0.5 | 2.62 | 0.09 | 2.03 | 0.25 | 1.29 | 0.16 |
| T241A K38E | 3.11 | 0.80 | n.d. | 50.8 | 0.4 | 11.32 | 0.30 | 1.86 | 0.19 | 6.08 | 0.64 |
| T241A D69H | 8.66 | 1.71 | n.d. | 47.0 | 0.6 | 12.09 | 0.33 | 2.37 | 0.22 | 5.10 | 0.49 |
| T241A L174V | 2.79 | 0.24 | n.d. | 37.2 | 0.2 | 12.70 | 0.31 | 5.27 | 0.37 | 2.41 | 0.18 |
| T241A P203A | 1.55 | 0.25 | n.d. | 50.6 | 0.4 | 14.99 | 0.36 | 2.26 | 0.19 | 6.65 | 0.57 |
| V216I K38E | 1.02 | 0.29 | n.d. | 37.5 | 0.8 | 5.20 | 0.23 | 5.30 | 0.70 | 0.98 | 0.14 |

| | | | | | | | | | | | |
|---|---|---|---|---|---|---|---|---|---|---|---|
| V216I D69H | 3.29 | 0.69 | n.d. | 37.8 | 0.1 | 5.39 | 0.21 | 4.28 | 0.50 | 1.26 | 0.15 |
| D69H K38E | 0.30 | 0.02 | n.d. | 36.9 | 0.1 | 4.33 | 0.31 | 12.65 | 2.04 | 0.34 | 0.06 |
| T241A D69H K38E | 1.37 | 0.10 | n.d. | 50.3 | 0.9 | 24.44 | 1.03 | 7.14 | 0.79 | 3.42 | 0.41 |
| S35T S51G | 0.68 | 0.07 | n.d. | 40.1 | 1.0 | 1.93 | 0.05 | 3.56 | 0.27 | 0.54 | 0.04 |
| Q41P T241A | 1.72 | 0.06 | n.d. | 49.7 | 0.7 | 6.45 | 0.16 | 1.61 | 0.14 | 4.00 | 0.37 |
| E36A T241A | 4.23 | 1.13 | n.d. | 49.2 | 0.1 | 9.42 | 0.39 | 2.12 | 0.32 | 4.44 | 0.70 |
| S35T T241A | 2.08 | 0.28 | n.d. | 51.3 | 0.1 | 3.28 | 0.07 | 1.26 | 0.11 | 2.61 | 0.24 |
| Q41P V262L | 2.14 | 0.37 | n.d. | 29.4 | 0.4 | 3.64 | 0.14 | 4.44 | 0.54 | 0.82 | 0.10 |
| M1P | 0.67 | 0.03 | n.d. | 36.7 | 0.4 | 3.58 | 0.11 | 5.14 | 0.46 | 0.70 | 0.07 |
| E146L | 0.44 | 0.14 | n.d. | 38.5 | 0.9 | 2.06 | 0.07 | 2.88 | 0.35 | 0.72 | 0.09 |
| P203G | 0.54 | 0.24 | n.d. | 39.7 | 0.3 | 2.78 | 0.11 | 5.14 | 0.58 | 0.54 | 0.06 |
| T241G | 3.23 | 0.37 | n.d. | 45.8 | 0.6 | 11.73 | 0.37 | 2.98 | 0.30 | 3.94 | 0.42 |
| A251I | 0.01 | 0.13 | n.d. | na | na | na | na | na | na | na | na |
| T241G E36A | 2.95 | 0.20 | n.d. | 45.6 | 0.5 | 13.95 | 0.32 | 2.46 | 0.19 | 5.68 | 0.46 |
| T241G L174V | 6.68 | 0.57 | n.d. | 29.8 | 1.04 | 22.71 | 0.77 | 6.25 | 0.58 | 3.63 | 0.36 |
| T241G P203A | 7.46 | 1.17 | n.d. | 39.3 | 0.00 | 49.31 | 1.50 | 5.08 | 0.44 | 9.70 | 0.90 |
| T241G K38E D69H | 0.50 | 0.05 | n.d. | 40.3 | 0.58 | 12.42 | 0.31 | 4.61 | 0.34 | 2.69 | 0.21 |
| T241G D69H | 9.72 | 1.25 | n.d. | 36.2 | 0.58 | 28.37 | 0.91 | 4.13 | 0.40 | 6.88 | 0.70 |
| T241G K38E | 6.36 | 1.40 | n.d. | 37.7 | 0.29 | 10.19 | 0.61 | 3.88 | 0.74 | 2.63 | 0.53 |
| S35T | 0.57 | 0.04 | n.d. | 33.4 | 0.29 | 2.79 | 0.04 | 3.38 | 0.17 | 0.82 | 0.04 |

| | | | | | | | | | | |
|---|---|---|---|---|---|---|---|---|---|---|
| Q41P | 0.76 | 0.04 | n.d. | 34.3 | 1.53 | 4.23 | 0.06 | 6.31 | 0.26 | 0.67 | 0.03 |
| V262L | 1.35 | 0.09 | n.d. | 30.3 | 1.00 | 5.94 | 0.11 | 9.51 | 0.41 | 0.62 | 0.03 |
| E36A | 0.73 | 0.12 | n.d. | 31.9 | 0.58 | 3.67 | 0.08 | 5.21 | 0.33 | 0.70 | 0.05 |
| S51G | n.d. | n.d. | n.d. | | n.d. | 2.08 | 0.07 | 5.45 | 0.54 | 0.38 | 0.04 |
| E146V | 4.24 | 0.92 | 1.68 | 31.1 | 0.87 | 1.73 | 0.06 | 2.44 | 0.29 | 0.71 | 0.09 |
| V6L V67I | n.d. | n.d. | n.d. | 29.1 | n.d. | 1.68 | 0.04 | 2.27 | 0.18 | 0.74 | 0.06 |

*Michaelis Menten curves were measured by following the decline of NADPH absorbance at 340 nm with 0.5 mM NADPH and 30 mM cyclopropylamine **a** in 100 mM Tris pH 8.0 with varying cyclohexanone **1** concentrations (between 35 and 0.06 mM) in three technical replicates (T=25 °C). Standard errors of the fit and three technical replicates are represented with a 95% confidence interval.

**Lysate activity was determined with 10 mM cyclopropylamine, 20 mM cyclohexanone and 0.5 mM NADPH in 100 mM Tris pH 8.0. Initial rates of NADPH depletion at 340 nm were measured, corrected for background by subtracting the rate of a negative control (pRSF vector expressing beta lactamase instead of SrIRED) and wild-type activity determined at the same plate (all in three biological replicates, errors represent standard deviations).

***The melting temperature $T_M$ was determined in a thermal shift assay (main text, methods) in three technical replicates. Errors represent the standard deviation of three replicates.

**Table S3: All combinable interactions with T241.** A list of all combinable interactions with T241 shows combinable interaction partners across the whole IRED sequence that were observed in the IrDMS data.

| Positions of combinable interaction partners of T241 | |
|---|---|
| 7 | 172 |
| 11 | 195 |
| 20 | 197 |
| 24 | 210 |
| 39 | 216 |
| 42 | 220 |
| 44 | 221 |
| 51 | 243 |
| 117 | 244 |
| 121 | 271 |
| 124 | 272 |
| 128 | 286 |
| 143 | 287 |

**Table S4: Summary table - Characterization of all *Sr*IRED homologs with cyclohexanone 1 and cyclopropylamine a as substrates.**

| Variant | $T_M$ [°C] ** | $T_M$ error | $k_{cat}$ [$s^{-1}$] * | $k_{cat}$ error | $K_M$ [mM] * | $K_M$ error | $k_{cat}/K_M$ [$s^{-1}mM^{-1}$] * | $k_{cat}/K_M$ error |
|---|---|---|---|---|---|---|---|---|
| SrIRED wt | 38.5 | 0.6 | 2.1 | 0.08 | 3.5 | 0.4 | 0.600 | 0.078 |
| SrIRED T241A | 50.0 | 0.7 | 5.8 | 0.24 | 2.3 | 0.3 | 2.492 | 0.368 |
| SmIRED wt | 41.6 | 1.9 | 5.7 | 0.08 | 4.5 | 0.2 | 1.279 | 0.054 |
| SmIRED T241A | 47.1 | 1.4 | 10.3 | 0.42 | 2.8 | 0.4 | 3.698 | 0.518 |
| ScIRED wt | 41.6 | 0.4 | 7.4 | 0.09 | 5.2 | 0.2 | 1.435 | 0.053 |
| ScIRED T241A | 47.8 | 1.3 | 12.1 | 0.52 | 3.6 | 0.5 | 3.310 | 0.464 |
| SbIRED wt | 43.9 | 1.2 | 6.5 | 0.34 | 9.5 | 1.2 | 0.686 | 0.094 |
| SbIRED T241A | 47.1 | 2.2 | 29.6 | 0.48 | 10.0 | 0.4 | 2.973 | 0.123 |
| BtIRED wt | 35.6 | 2.2 | 13.9 | 0.41 | 11.5 | 0.8 | 1.210 | 0.089 |
| BtIRED T241A | 50.9 | 1.3 | 44.5 | 0.80 | 5.3 | 0.3 | 8.366 | 0.453 |
| SlIRED wt | 39.9 | 1.1 | 3.4 | 0.08 | 7.4 | 0.5 | 0.457 | 0.030 |
| SlIRED T241A | 49.1 | 0.3 | 10.8 | 0.43 | 6.3 | 0.7 | 1.719 | 0.197 |
| SxIRED wt | 32.2 | 0.4 | 4.1 | 0.12 | 11.7 | 0.8 | 0.354 | 0.026 |
| SxIRED T241A | 37.3 | 0.7 | 13.1 | 0.50 | 12.3 | 1.0 | 1.068 | 0.099 |
| LpIRED wt | 33.2 | 0.5 | 0.8 | 0.03 | 3.9 | 0.5 | 0.208 | 0.029 |
| LpIRED T241A | 33.9 | 0.6 | 1.1 | 0.07 | 2.2 | 0.6 | 0.493 | 0.148 |
| SoIRED wt | 35.6 | 0.2 | 1.1 | 0.03 | 2.3 | 0.2 | 0.470 | 0.051 |
| SoIRED T241A | 37.2 | 0.9 | 1.8 | 0.04 | 1.5 | 0.1 | 1.166 | 0.106 |
| AsIRED wt | 35.7 | 1.8 | 0.3 | 0.02 | 45.3 | 4.8 | 0.007 | 0.001 |
| AsIRED T241A | 38.6 | 0.6 | 0.0 | 0.01 | 38.2 | 11.3 | 0.001 | 0.000 |
| SuIRED wt | 28.9 | 0.1 | n.a. | n.a. | n.a. | n.a. | n.a. | n.a. |
| SuIRED T241A | 33.6 | 1.3 | **n.a.** | **n.a.** | **n.a.** | **n.a.** | **n.a.** | **n.a.** |

*Michaelis Menten curves were measured by following the decline of NADPH absorbance at 340 nm with 0.5 mM NADPH and 30 mM cyclopropylamine **a** in 100 mM Tris pH 8.0 with varying cyclohexanone **1** concentrations (between 35 and 0.06 mM) in three technical replicates (T=25 °C). Standard errors of the fit and three technical replicates are represented with a 95% confidence interval.

**The melting temperature $T_M$ was determined in a thermal shift assay (main text, methods) in three technical replicates. Errors represent the standard deviation of three replicates.

**Table S5: Summary table - Characterization of all *Sb*IRED mutants with cyclohexanone 1 and cyclopropylamine a as substrates.**

| Variant | $T_M$ [°C] ** | $T_M$ error | $k_{cat}$ [$s^{-1}$] * | $k_{cat}$ error | $K_M$ [mM] * | $K_M$ error | $k_{cat}/K_M$ [$s^{-1}mM^{-1}$] * | $k_{cat}/K_M$ error |
|---|---|---|---|---|---|---|---|---|
| SbIRED wt | 43.9 | 1.2 | 6.5 | 0.3 | 9.5 | 1.2 | 0.7 | 0.1 |
| SbIRED T241A | 47.1 | 2.2 | 29.6 | 0.5 | 10.0 | 0.4 | 3.0 | 0.1 |
| SbIRED D69H | 40.3 | 1.4 | 7.0 | 1.1 | 21.5 | 6.7 | 0.3 | 0.1 |
| SbIRED K38E | 46.9 | 0.6 | 0.4 | 0.0 | 7.9 | 0.9 | 0.0 | 0.0 |
| SbIRED P203A | 46.1 | 0.2 | 7.3 | 0.4 | 18.2 | 1.9 | 0.4 | 0.0 |
| SbIRED D69H T241A | 41.4 | 0.1 | 55.7 | 2.1 | 10.7 | 1.0 | 5.2 | 0.5 |
| SbIRED K38E T241A | 50.0 | 0.4 | 2.4 | 0.1 | 6.7 | 1.1 | 0.4 | 0.1 |
| SbIRED P203A T241A | 45.9 | 0.5 | 46.0 | 1.8 | 6.4 | 0.7 | 7.2 | 0.8 |

*Michaelis Menten curves were measured by following the decline of NADPH absorbance at 340 nm with 0.5 mM NADPH and 30 mM cyclopropylamine **a** in 100 mM Tris pH 8.0 with varying cyclohexanone **1** concentrations (between 35 and 0.06 mM) in three technical replicates (T=25 °C). Standard errors of the fit and three technical replicates are represented with a 95% confidence interval.
**The melting temperature $T_M$ was determined in a thermal shift assay (main text, methods) in three technical replicates. Errors represent the standard deviation of three replicates.

**Table S6: Summary table - Characterization of all *Pc*IRED variants with aldehyde 10 and amine f as substrates.**

| Variant | Delta TM to wt [°C]** | $T_M$ error | $k_{cat}$ [$s^{-1}$] * | $k_{cat}$ error | $K_M$ [mM] * | $K_M$ error | $k_{cat}/K_M$ [$s^{-1}mM^{-1}$] * | $k_{cat}/K_M$ error |
|---|---|---|---|---|---|---|---|---|
| wt | | | 0.1184 | 0.0073 | 2.7030 | 0.4146 | 0.0438 | 0.0072 |
| A181G | -5.1 | 0.3 | 0.1090 | 0.0042 | 1.5000 | 0.1677 | 0.0727 | 0.0086 |
| A181T | -5.0 | 0.2 | 0.2324 | 0.0346 | 8.5360 | 2.0860 | 0.0272 | 0.0078 |
| R61L | -0.5 | 0.1 | 0.1235 | 0.0060 | 2.3900 | 0.2951 | 0.0517 | 0.0069 |
| R61L A181T | -6.0 | 0.6 | 0.2694 | 0.0149 | 3.3120 | 0.4183 | 0.0813 | 0.0112 |
| E107K | -0.2 | 0.1 | 0.1006 | 0.0045 | 1.5320 | 0.1998 | 0.0657 | 0.0091 |
| A135I | No expression | | | | | | | |
| P87G | 0.9 | 0.7 | 0.1054 | 0.0039 | 2.4460 | 0.2263 | 0.0431 | 0.0043 |
| A168N | -7.5 | 0.5 | Activity < 0.8 x wild type | | | | | |
| A181S | -7.6 | 0.7 | 0.2438 | 0.0287 | 8.8190 | 1.7450 | 0.0276 | 0.0064 |
| K249G | -5.0 | 0.9 | 0.1040 | 0.0059 | 3.2850 | 0.4323 | 0.0317 | 0.0045 |
| Q228R K249N | -6.4 | 0.2 | 0.0958 | 0.0025 | 1.2420 | 0.0999 | 0.0771 | 0.0065 |
| I91L K249N | -0.1 | 0.3 | 0.1078 | 0.0053 | 0.9711 | 0.1380 | 0.1110 | 0.0167 |
| H199L K249N | -1.4 | 0.7 | 0.2247 | 0.0162 | 4.2070 | 0.6435 | 0.0534 | 0.0090 |
| K249N K296R | 0.9 | 1.0 | 0.1134 | 0.0046 | 0.8039 | 0.0566 | 0.1411 | 0.0115 |
| S178G K230E | -1.7 | 1.0 | 0.1369 | 0.0035 | 2.8580 | 0.1765 | 0.0479 | 0.0032 |
| S178G K249N | -3.4 | 0.6 | 0.1101 | 0.0049 | 2.2730 | 0.2630 | 0.0484 | 0.0060 |
| P87S S178I | -13.7 | 0.3 | | | | | | |
| S178G P295S | -3.8 | 0.3 | 0.1316 | 0.0042 | 3.1730 | 0.2349 | 0.0415 | 0.0033 |
| S178C K273E | -2.9 | 0.1 | 0.1043 | 0.0021 | 1.8670 | 0.1039 | 0.0559 | 0.0033 |

*Michaelis Menten curves were measured by following the decline of NADPH absorbance at 340 nm with 0.5 mM NADPH and 20 mM amine **f** in 100 mM Tris pH 8.0, 50% DMSO with varying aldehyde **10** concentrations (between 10 and 0.014 mM) in three technical replicates (T=25 °C). Standard errors of the fit and three technical replicates are represented with a 95% confidence interval.
**The melting temperature was determined in a thermal shift assay in three technical replicates. Errors represent the standard deviation of three replicates.

**Table S7: Oversampling of the *Pc*IRED library in screening and next generation sequencing.** 'Droplets screened' refers to the number of droplets assayed in each AADS sample and 'droplets sorted' refers to the number of droplets selected based on the used selection threshold. The 'oversampling' constant refers to the used 60,000 membered *Pc*IRED variant library which was encapsulated with 50% occupancy using the Poisson distribution. NGS reads and depth are calculated for the 'input' library before sorting and the 'output' library after sorting where 'reads' refers to the total number of NGS reads for each sample and 'depth' refers to the coverage of the input library (60,000 variants) and the output library (number of sorted droplets) with NGS reads.

| Sample name | Droplets screened [x $10^6$] | Droplets sorted | Oversampling (with 50% occupancy) | NGS reads input [x $10^6$] | NGS depth input | NGS reads output [x $10^6$] | NGS depth output |
|---|---|---|---|---|---|---|---|
| 20h_1 | 1.1 | 17,100 | 9.2 | 26,408,584 | 440 | 5,754,368 | 337 |
| 20h_2 | 1.1 | 14,532 | 9.2 | 24,843,396 | 414 | 6,091,672 | 419 |
| 20h_3 | 1.3 | 19,000 | 10.8 | 20,332,192 | 339 | 7,677,559 | 404 |
| 40h_1 | 1.1 | 16,800 | 9.2 | 20,332,192 | 339 | 4,880,231 | 290 |
| 40h_2 | 1.2 | 15,494 | 10 | 22,053,928 | 368 | 7,801,163 | 503 |
| 40h_3 | 1.1 | 16,813 | 9.2 | 21,998,584 | 367 | 8,076,957 | 480 |

**Table S8: Organisms and accession numbers of characterised SrIRED homologs**

| IRED short | Organism | Uniref accession number |
|---|---|---|
| SrIRED | Streptosporangium roseum | WP_012887675.1 |
| SmIRED | Streptosporangium minutum | WP_086575968.1 |
| ScIRED | Streptosporangium canum | WP_363977292.1 |
| SbIRED | Streptosporangium brasiliense | WP_306863003.1 |
| BtIRED | Bailinhaonella_thermotolerans | WP_119927492.1 |
| SxIRED | Streptomyces xanthochromogenes | WP_189766438.1 |
| SlIRED | Streptomyces lushanensis | WP_066946639.1 |
| LpIRED | Lentzea pudingi | WP_308426073.1 |
| SoIRED | Streptomyces odonnellii | WP_079023258.1 |
| AsIRED | Amycolatopsis sp. YIM 10 | WP_153033184.1 |
| SuIRED | Streptomyces uncialis | WP_073792574.1 |

**Table S9: Overview over past deep mutational scanning studies with enzymes.** Criteria for consideration in this table are that (i) enzymatic activity is quantitatively assessed or reflected in fitness scores and (ii) the screen considers at least 1000 variants. Three different categories of deep mutational scanning studies can be differentiated showing how short read sequencing has previously restricted access to unbiased epistatic data: (A) Studies assessing only single point mutations considering no epistasis either using site saturation libraries or libraries that contain epispastic interactions which are not measured; (B) Studies resolving epistasis of a subset of possible positions either via single site saturation libraries with multiple different parent variants, combinatorial libraries of short stretches of the enzyme or only looking at consecutive positions; (C) Studies that resolve epistasis in the whole enzyme in an unbiased manner either via nested libraries artificially "stitching" short reads[10] of the same variants which scales poorly with the size of the enzyme or studies that directly sequence the whole gene using long reads (this study). lrDMS is unique in providing a technique directly resolving the full-length gene (scaling well) and allows for unbiased exploration of the whole gene, which in a pooled screen to our knowledge has previously only been shown with the SARS-Cov-2 RBD[11]. The table contains literature published until 07/2023.

| | | Screening / Selection Method | | | | | | Library type | | | Sequencing | | | Epistasis resolution | | | | | | | Downstream experiments | | | | |
|---|---|---|---|---|---|---|---|---|---|---|---|---|---|---|---|---|---|---|---|---|---|---|---|---|---|
| Type | Reaction | Microfluidics | FACS | Selection | Genetic Circuit | Phage Display | Robotic / Manual screening | Single-site saturation | Random mutagenesis | Focused combinatorial | NGS (short reads) | Sanger sequencing | Long-read sequencing | Epistasis resolved? | Unbiased higher-order mutagenesis? | Pairwise epistasis with N parent variants? | Combinatorial library with N positions? | Only sequential mutations? | Directly resolved? | Stitching reads? | Rational Extrapolation? | Machine Learning? | Different substrates? | Data Points | Reference |
| Oxidoreductase | Imine Reductase | ✓ | | | | | | | ✓ | | ✓ | | ✓ | ✓ | | | | | ✓ | | ✓ | ✓ | ✗ | 17143 | This study |
| | | | | | | | ✓ | ✓ | ✓ | | | | ✓ | ✓ | | | | | ✓ | | ✗ | ✓ | ✗ | 11303 | [3] |
| | DHFR | | | ✓ | | | | ✓ | | | ✓ | | | n.a. | | | | | | | ✗ | ✗ | ✗ | 3161 | [12] |
| | TetX | | | ✓ | | | | | ✓ a | | ✓ | | | n.a. | | | | | | | ✗ | ✗ | ✓ | Non. Rep. | [13] |

| | | | | | | | | | | | | | | | | | | | | | | | | | |
|---|---|---|---|---|---|---|---|---|---|---|---|---|---|---|---|---|---|---|---|---|---|---|---|---|---|
| **Hydrolase** | Amidase | | | ✓ | | | | ✓ | | | ✓ | | | n.a. | | | | | | | ✗ | ✗ | ✓ | 6568 | 14 |
| | | | | ✓ | | | | ✓ | | | ✓ | | | ✓ | ✗ | 3 | | | ✓ | | ✗ | ✗ | ✗ | 6391 | 15 |
| | Metallo-β-lactamases | | | ✓ | | | | ✓ | | | ✓ | | | n.a. | | | | | | | ✗ | ✗ | ✓ | 5607 | 16 |
| | TEM-1 β-lactamase | | | | ✓ | | | ✓ | | | ✓ | | | n.a. | | | | | | | ✗ | ✗ | ✗ | 2536 | 17 |
| | | | | ✓ | | | | ✓ d | | | ✓ | | | n.a. | | | | | | | ✗ | ✗ | ✗ | 6000 | 18 |
| | | | | ✓ | | | | | ✓ a | | | ✓ | | ✓ | ✓ | | | | ✓ | | ✗ | ✗ | ✗ | 10000 | 19 |
| | | | | ✓ | | | | ✓ | | | ✓ | | | n.a. | | | | | | | ✗ | ✗ | ✗ | 5000 | 20 |
| | | | | ✓ | | | | ✓ | | | ✓ | | | n.a.[e] | | | | | | | ✗ | ✗ | ✗ | ~5000[b] | 21 |
| | | | | ✓ | | | | ✓ | | | ✓ | | | ✓ | ✗ | 3 | | | ✓ | | ✗ | ✗ | ✗ | ~15000[b] | 22 |
| | | | | ✓ | | | | | | | ✓ | | | ✓ | ✗ | | | ✓ | ✓ | | ✗ | ✗ | ✗ | 12000 | 23 |
| | HSP90 | | | ✓ | | | | | | ✓ | ✓ | | | ✓ | ✗ | | 9 | | ✓ | | ✗ | ✗ | ✗ | 1015 | 24 |
| | Gsp1/Ran GTPase | | | ✓ | | | | ✓ | | | ✓ | | | n.a. | | | | | | | ✗ | ✗ | ✗ | 4315 | 25 |
| | Ras GTPase | | | ✓ | | | | ✓ | | | ✓ | | | ✓ | ✗ | 2 | | | | ✓ | ✗ | ✗ | ✗ | 3000[b] | 26 |
| | GPRC | | ✓ | | | | | | | ✓ | ✓ | | | ✓ | ✗ | | 5 | | | ✓ | ✗ | ✗ | ✗ | < $3.2*10^{6e}$ | 27 |
| | Caspase | ✓ | | | | | | | ✓ a | | ✓ | | | n.a. | | | | | | | ✗ | ✗ | ✗ | Not rep. | 28 |
| | Neuraminidase | | | ✓ | | | | | ✓ a | | ✓ | | | n.a. | | | | | | | ✗ | ✗ | ✗ | Not rep. | 29 |
| | N-acetylglucosamine deacetylase | | | ✓ | | | | ✓ c | | | ✓ | | | n.a. | | | | | | | ✗ | ✗ | ✗ | ~6000 | 30 |
| | Alkaline Phosphatase PafA | ✓ | | | | | | ✓ | | | | ✓ | | n.a. | | | | | | | ✗ | ✗ | ✗ | 1036 | 31 |
| | Cas9 | | | ✓ | | | | | ✓ a | | ✓ | | | n.a. | | | | | | | ✗ | ✗ | ✗ | 8500 | 32 |

| | | | | | | | | | | | | | | | | | | | | | | | | | |
|---|---|---|---|---|---|---|---|---|---|---|---|---|---|---|---|---|---|---|---|---|---|---|---|---|---|
| | β-glucosidase | ✓ | | | | | | | ✓[a] | | ✓ | | | n.a. | | | | | | | ✗ | ✗ | ✗ | 3083 | 33 |
| **Transferase** | Kinase | ✓ | ✓ | | | | | | | ✓ | ✓ | | | ✓ | ✗ | | 6 | | ✓ | | ✗ | ✗ | ✗ | $5*10^5$ | 34 |
| | | | ✓ | ✓ | | | | ✓ | | | ✓ | | | ✓ | ✗ | | 4 | | ✓ | | ✗ | ✗ | ✗ | $1.6*10^5$ | 35 |
| | | | | ✓ | | | | ✓ | | | ✓ | | | n.a. | | | | | | | ✗ | ✗ | ✓ | 4993 | 36 |
| | | | | ✓ | | | | | | | ✓ | | | n.a. | | | | | | | ✗ | ✗ | ✗ | 7000 | 37 |
| | | | | ✓ | | | | ✓ | | | ✓ | | | n.a. | | | | | | | ✗ | ✗ | ✗ | $7*10^5$ | 38 |
| | | | | ✓ | | | | ✓ | | | ✓ | | | n.a. | | | | | | | ✗ | ✗ | ✗ | 6810 | 39 |
| | | | | ✓ | | | | ✓ | | | ✓ | | | n.a. | | | | | | | ✗ | ✗ | ✗ | 8000 | 40 |
| | N-acetylglucosamine 1-carboxyvinyltransferase | | | ✓ | | | | ✓[c] | | | ✓ | | | n.a. | | | | | | | ✗ | ✗ | ✗ | ~8000[b] | 30 |
| | E3 Ubiquitin Ligase | | | ✓ | | | | ✓[c] | | | ✓ | | | n.a. | | | | | | | ✗ | ✗ | ✗ | 3893 | 41 |
| | | | | | | ✓ | | | ✓ | | ✓ | | | ✓ | ✓ | | | | | ✓ | ✗ | ✗ | ✗ | 98289 | 10 |
| | DNA Polymerase | ✓ | ✓ | | | | | ✓ | | | ✓ | | | n.a. | | | | | | | ✗ | ✗ | ✗ | 960 | 42 |
| **Lyase** | Phenylalanine Ammonia-Lyase | | | ✓ | | | | ✓ | | | ✓ | | | n.a. | | | | | | | ✗ | ✗ | ✗ | ~4000[b] | 43 |
| | | | | ✓ | | | | | ✓ | | ✓ | | | ✓ | ✗ | | 7 | | ✓ | | ✗ | ✗ | ✗ | Not rep. | |
| | 3-Hydroxyacyl ACP dehydratase | | | ✓ | | | | ✓[c] | | | ✓ | | | n.a. | | | | | | | ✗ | ✗ | ✗ | ~3000[b] | 30 |
| | Indole-3-glycerol-phosphate synthase | | | ✓ | | | | ✓ | | | ✓ | | | ✓ | ✗ | 3 | | | ✓ | | ✗ | ✗ | ✗ | ~15000[b] | 44 |
| | Cystathionine-β-synthase | | | ✓ | | | | | ✓ | | ✓ | | | n.a. | | | | | | | ✗ | ✗ | ✗ | ~10000[b] | 45 |
| **Translocase** | AcrB multidrug efflux pump | | | ✓ | | | | ✓[c] | | | ✓ | | | n.a. | | | | | | | ✗ | ✗ | ✗ | 4240 | 46 |
| | MurJ Lipid II flippase | | | ✓ | | | | | ✓ | | ✓ | | | n.a. | | | | | | | ✗ | ✗ | ✗ | 1500 | 47 |

[a]Mutagenesis by error-prone PCR
[b]Number of fitness scores not reported – estimated by theoretical library size
[c]CRIPR-based in vivo mutagenesis
[d]InDels
[e]Combinatorial library used but not sequenced

**Table S10: Michaelis Menten parameters of SrIRED from this study, engineering campaigns documented in the literature and wild type IREDs that have been kinetically characterised (for intermolecular reductive amination reactions).** Green: highest performing studies (deeper shading indicates higher performance). Red frame: best performance (within error). This table (combined with **Table S6**) illustrates that (i) our engineered *Sr*IRED has the highest $k_{cat}/K_M$ ever seen in reductive aminations (ii) only one RedAm[48] has $k_{cat}$ values in the region of our engineered *Sr*IRED (but required multiple rounds of directed evolution) (iii) we report the highest improvements in activity over one round of directed evolution (iv) most campaigns start with extremely low activities (these are easier to improve) while we improve an already proficient enzyme. (v) the only other machine learning study of an IRED[49] shows only a 1.2-fold (20%) improvement – we show a 17-fold higher improvement (includes all literature published until 04/2024).

| IRED | Variant | Method | $k_{cat}$ [$s^{-1}$] | $K_M$ ketone [mM] | $k_{cat}/K_M$ [$s^{-1}mM^{-1}$] | Maximal improvement per round[a] | Substrate | Reference |
|---|---|---|---|---|---|---|---|---|
| SrIRED | wt | lrDMS profiling in µdroplets | 2.1 | 3.5 | 0.6 | **23-fold** for $k_{cat}$ **16-fold** for $k_{cat}/K_M$ | Cyclohexanone + Cyclopropylamine | This paper |
| SrIRED | T241A | | 5.8 | 2.3 | 2.5 | | Cyclohexanone + Cyclopropylamine | This paper |
| SrIRED | T241G P203A | | **49.3** | 5.1 | **9.7** | | Cyclohexanone + Cyclopropylamine | This paper |
| AspRedAm | wt | Homologue exploration | 1.47 | 1.90 | 0.73 | n/a | Cyclohexanone + Methylamine | 50 |
| AspRedAm | wt | | 5.0 | 2.3 | 2.17 | | Cyclohexanone + Allylamine | 50 |
| AtRedAm | wt | | 0.11 | 2.1 | 0.05 | | Cyclohexanone + Allylamine | 51 |
| AdRedAm | wt | | 2.1 | 3.8 | 0.56 | | Cyclohexanone + Allylamine | 51 |
| (S)-IRED | wt | | 0.02 | 3 | 0.007 | | Cyclohexanone + Allylamine | 51 |
| IRED 17 | wt | | 4.74 | 57.2 | 0.08 | | Cyclohexanone + Cyclopropylamine | 52 |
| IRED 18 | wt | | 0.62 | 20.2 | 0.03 | | Cyclohexanone + Cyclopropylamine | 52 |
| IRED 33 | wt | | 5.84 | 69.8 | 0.08 | | Cyclohexanone + Cyclopropylamine | 52 |

| | | | | | | | | |
|---|---|---|---|---|---|---|---|---|
| IRED 69 | wt | | 6.93 | 29.6 | 0.23 | | Cyclohexanone + Cyclopropylamine | 52 |
| SpRedAM | wt | Directed evolution by robotic plate screening (3 rounds) | 2.7 | 2.5 | 1.1 | **2.7-fold** for $k_{cat}$ [b] **1.5-fold** for $k_{cat}/K_M$ [c] | API building block | 53 |
| SpRedAm R1V1 | R1 | | 14.6 | 5.58 | 2.6 | | API building block | 53 |
| SpRedAm R1V2 | R1 | | 28.9 | 14.2 | 1.97 | | API building block | 53 |
| SpRedAm R2V3 | R2 | | 17.9 | 15.0 | 1.2 | | API building block | 53 |
| SpRedAm R2V4 | R2 | | 16.0 | 8.94 | 1.8 | | API building block | 53 |
| SpRedAm R3V5 | R3 | | 22.8 | 8.5 | 2.7 | | API building block | 53 |
| SpRedAm R3V6 | R3 | | **52.2** | 13.8 | 3.8 | | API building block | 53 |
| IR-77 | wt | Structure-based engineering | 0.24 | 15 | 0.016 | **1.8-fold** for $k_{cat}$ **2.8-fold** for $k_{cat}/K_M$ | Cyclohexanone + Pyrrolidine | 54 |
| IR-77 | A208S | | 0.43 | 9.6 | 0.045 | | Cyclohexanone + Pyrrolidine | 54 |
| IR-G02 | wt | n/a | 19.8 | 6.58 | 3.00 | n/a | Cyclohexanone + Cyclopropylamine | 55 |
| IR-G36 | wt | Directed evolution (5 rounds) | $7 \times 10^{-4}$ | 32.3 | $2.2 \times 10^{-5}$ | **4.7-fold** for $k_{cat}$ [d] **5-fold** for $k_{cat}/K_M$ [e] | Cyclohexanone derivative + Benzylamine derivative | 55 |
| IR-G36 M1 | 2 mutations | | $5.3 \times 10^{-3}$ | 33.0 | 0.00017 | | Cyclohexanone derivative + Benzylamine derivative | 55 |
| IR-G36 M2 | 4 mutations | | 0.17 | 32.2 | 0.0052 | | Cyclohexanone derivative + Benzylamine derivative | 55 |
| IR-G36 M3 | 6 mutations | | 1.08 | 34.8 | 0.031 | | Cyclohexanone derivative + Benzylamine derivative | 55 |

| | | | | | | | | |
|---|---|---|---|---|---|---|---|---|
| IR-G36 M4 | 8 mutations | | 1.54 | 27.5 | 0.056 | | Cyclohexanone derivative + Benzylamine derivative | 55 |
| IR-G36 M5 | 10 mutations | | 1.58 | 22.6 | 0.07 | | Cyclohexanone derivative + Benzylamine derivative | 55 |
| pIR23 | wt | Exploration of multiple substrates | 3.47 | 10.1 | 0.34 | n/a | Cyclohexanone + Cyclopropylamine | 56 |
| pIR23 | wt | | 1.84 | 0.64 | 2.88 | | Hydrocinnamaldehydw + pyrrolidine | 56 |
| pIR23 | wt | | 0.77 | 0.25 | 3.08 | | Cinnamaldehyde +pyrrolidine | 56 |
| BacRedAm | wt | | 5.12 | 13.3 | 0.38 | | Cyclohexanone + Cyclopropylamine | 56 |
| BacRedAm | wt | | 2.01 | 3.32 | 0.61 | | Hydrocinnamaldehyde + Allylamine | 56 |
| BacRedAm | wt | | 0.11 | 2.58 | 0.04 | | Cinnamaldehyde + Cyclopropylamine | 56 |

[a] Assuming equal improvements for multi-round campaigns

[b] $(52.2/2.7)^{1/3} = 2.7$
[c] $(3.8/1.1)^{1/3} = 1.5$
[d] $(1.58/0.0007)^{1/5} = 4.7$
[e] $(0.07/0.000022)^{1/5} = 5$

**Table S11: Specific activities in U/mg of SrIRED from this study and engineering campaigns documented in the literature** (for intermolecular reductive amination reactions). Green: highest performing studies (deeper shading indicates higher performance). Red frame: best performance. This table (combined with **Table S5**) illustrates that (i) our engineered *Sr*IRED has the highest $k_{cat}/K_M$ ever seen in reductive aminations (ii) only one RedAm[48] has $k_{cat}$ values in the region of our engineered *Sr*IRED (but required multiple rounds of directed evolution) (iii) we report the highest improvements in activity over one round of directed evolution (iv) most campaigns start with extremely low activities (these are easier to improve) while we improve an already proficient enzyme. (v) the only other machine learning study of an IRED[49] shows only a 1.2-fold (20%) improvement – we show a 17-fold higher improvement. (includes all literature published until 04/2024).

| IRED | Variant | Specific activity [U/mg] | $k_{cat}$ [$s^{-1}$] | Maximal improvement per round[a] | Substrates | Reference |
|---|---|---|---|---|---|---|
| SrIRED | wt | - | 2.1 | - | Cyclohexanone + Cyclopropylamine | This paper |
| SrIRED | T241A: DMS alone | - | 5.8 | 3-fold | Cyclohexanone + Cyclopropylamine | This paper |
| SrIRED | T241G P203A: lrDMS profiling in µdroplets | - | 49.3 | **23-fold** | Cyclohexanone + Cyclopropylamine | This paper |
| IRED-88 | wt | 45 | - | - | API building block | [3] |
| IRED-88 | DMS alone | 88 | - | 2-fold | API building block | [3] |
| IRED-88 | Machine learning | 54 | - | 1.2-fold | API building block | [3] |
| IRED-88 | 3 rounds of directed evolution | 106 | - | 1.3-fold [b] | API building block | [3] |
| GSK IR-46 | wt | 0.75 | - | - | API building block | [57] |
| GSK IR-46 M3 | 13 mutations accumulated in 3 rounds of directed evolution | 9.76 | - | 2.4-fold [c,d] | API building block | [57] |
| PcIRED | wt | 0.017 | - | - | Bulky aldehyde + bulky amine | [58] |
| PcIRED M3* | 8 mutations accumulated in 3 rounds of directed evolution | 8.14 | - | **7.8-fold** [e] | Bulky aldehyde + bulky amine | [58] |

[a] Assuming equal improvements for multi-round campaigns. [b] $(106/45)^{1/3} = 1.3$ [c] $(9.76/0.75)^{1/3} = 2.4$ [d] Schober et al. report a 38,000-fold improvement which is however calculated cumulatively by only considering conversion. The improvement in specific activity is only 13-fold in total and 2.4-fold per round of directed evolution. [e] $(8.14/0.017)^{1/3} = 7.8$

### Supplementary Notes

#### 1 Quality control: Coverage, accuracy, and bias of the lrDMS dictionary

Sanger sequencing of random library members showed that the dictionary covers 85% of the library (76 of 89 examined variants were found in the dictionary) and is highly accurate with an error rate of 0.4% (only one error was found among 235 mutations). Among the dictionary members, we observed a slight bias towards transitions (interchange of purines and pyrimidines; **Supplementary Figure 5A**) mirroring trends in natural mutagenesis[59] and a slight bias of mutations to A or T and from C or G. On average, variants have 3.1 amino acid mutations with 1514 unique single point mutations (**Extended Data Figure 2A and B**) which are distributed across the whole IRED sequence (**Supplementary Figure 4**). This corresponds to 26% of all possible 5800 single mutations approaching the maximum possible diversity of single point mutations given the constraints of the genetic code (~1750 variants; **Extended Data Figure 2C**).

#### 2 Quality control: lrDMS is accurate, quantitative, and chemoselective

To validate the screening strategy in droplets, we conducted a mock-sorting experiment with a 1:100 dilution of *E.coli* expressing *Sr*IRED compared to *E. coli* containing an empty (but otherwise identical) vector (**Extended Data Figure 1A**). After testing the sorted population for activity in a plate assay, we found that 93% of variants are active IREDs, showing a 93-fold enrichment of active variants [60] (**Extended Data Figure 1B**). This shows that our novel AADS screening strategy is a reliable and accurate method to capture variant-specific fitness information for lrDMS. To test the chemoselectivity of hits after sequencing we tested a plate of hits for activity with ketone and found that we are not enriching for ketoreductase activity (**Supplementary Figure 2**). To test whether the fitness information quantitatively reflects on the proficiencies of the respective variants, 14 of them were picked randomly from the input library and, for comparison, from the library after sorting (7 and 7). Their conversion was measured in a lysate assay in plates. The logarithmic activity in plates correlated well with the fitness score (Pearson r = 0.92) showing that our fitness score captures quantitative fitness information of variants with larger and smaller than wild type activity (**Figure 1E**).

#### 3 Literature comparison: One lrDMS round from an optimised starting point achieves comparable conversion fold-changes as three cycles of traditional directed evolution for a challenging drug precursor

The *Pc*IRED engineering campaign demonstrates that our lrDMS workflow can successfully be applied to challenging substrates (a bulky carbonyl compound and a bulky amine) as well as to hard-to-evolve starting points (the final variant of a previously stalled evolution campaign). The challenge of the *Pc*IRED engineering scenario was confirmed by the obtained sequence-function dataset which reports a lower than average mutability in the M3 *Pc*IRED when compared to *Sr*IRED. To grant comparability between our results and the results reported by Chen *et al.*, we normalised the reported specific activity improvements to the used reaction conditions (**Table SN1**). *Chen et al*. had used 10 mM ketone and 10 mM amine concentrations for wild-type kinetics but 100 mM ketone and 100 mM amine concentrations for the kinetic measurements of the evolved variants M1, M2 and M3. Our normalisation helped to disentangle any effect from substrate concentration [S] on enzymatic rate as described by the Michaelis-Menten law. Therefore, we distinguish two regimes, a regime with a near-linear dependence on [S] around and below $K_M$ and a substrate independent regime at [S] >> $K_M$. After normalisation, Chen *et al.* achieved specific activity improvements of 4.8-fold in three rounds of extensive screening of more than 47 site saturation libraries. This corresponds to 3.2-fold improvements in conversion as reported by Chen *et al.* In our study, we reach similar results in specific activity as well as conversion improvements in only one round of engineering using lrDMS.

**Table SN1: Comparison of our engineering campaign for *Pc*IRED with literature data from Chen *et al*.** Specific activity improvements reported by Chen *et al.* are normalised to the used reaction conditions to grant comparability with improvements reported in our work. The comparison highlights that our fold-change improvement in one round is comparable to the improvements achieved by Chen *et al.* in three rounds. The campaign by Chen *et al.* suffered from diminishing returns and stalled with nearly all improvements observed in round 1. In this work, we rescue the stalled evolution campaign with a 2.8-fold improvement in conversion on top of Chen *et al.*'s final variant.

| **Round** | **Specific activity (fold-change to previous round)** | | **Conversion (fold-change to previous round)**[2, 3] |
|---|---|---|---|
| | Reported | Adjusted for reaction conditions | |
| M1 (Chen *et al.*), Cinacalcet[4] | 101[1] | 1.01 (assumes fully linear regime) – 101 (assumes full substrate saturation) | 2.96 |
| M2 (Chen *et al.*), Cinacalcet[4] | 2.5 | 2.5 | 1.05 |
| M3 (Chen *et al.*), Cinacalcet [4] **Starting point in this work** | 1.9 | 1.9 | 1.01 |
| Cumulative improvement over **three** rounds Chen *et al.*, Cinacalcet | 488 | 4.8 | 3.16 |
| **Per-round** improvement Chen *et al.*, Cinacalcet | 7.9 | 1.7 | 1.5 |
| **One** round of engineering *This work*, Tecalcet[4] | 2.3 | | 2.80 |

[1]Data as represented in Chen *et al.* Table 1 footnote '*b*' (panel B). Note: 10-fold lower concentration of aldehyde and amine substrates were used for wild type compared to mutant kinetic measurements.
[2]More reliable as same conditions were used for all mutants.
[3]Comparison of 25%, 74%, 78% and 79% for *Pc*IRED wild type, M1, M2 and M3 by Chen *et al.* All measured at 10 mM concentration of the substrates.
[4]Cinacalcet and Tecalcet are very similar in structure with substrates classified as bulky.

### Supplementary Methods

**1 Manufacturing of microfluidic devices** (see Figure S1)

Master molds were prepared via standard Su-8 photolithography. First, the designs of the microfluidic chips were prepared using AutoCAD (Autodesk) and patterned on a high-resolution film photomask (Micro Lithography Services). Then, microfluidic molds were fabricated via photolithography using MJB4 mask aligner (SÜSS MicroTec) to UV expose 3" silicon wafers (Microchemicals) spin-coated with SU-8 photoresists (Kayaku Advanced Materials). Final microfluidic devices were prepared by soft lithography by mixing of PDMS elastomer base and crosslinking agent at a 10:1 ratio (Sylgard 184, Dow), pouring on the master mold and curing overnight at 65°C. After cutting chips using a scalpel, holes were punched using 1 mm biopsy punchers (Kai Medical). After treatment with plasma (Diener Femto), chips were bonded onto microscope glass slides, and channels were treated with 1% (v/v) trichloro(1H,1H,2H,2H-perfluorooctyl)silane (Sigma Aldrich) in fluorinated oil (HFE-7500, Novec) and incubated at 65°C overnight.

**2 Substrate scoping of *Sr*IRED wt and selected mutants based on initial rates** (see Figure S7)

The substrate scope of *Sr*IRED wt and the variants T241A, D69H, K38E, E146V, D236G and D5H was determined using substrates **1** – **4**, **9** and **a** – **e**. Activities were tested as initial rates with 10 mM ketone and 20 mM amine, or 10 mM imine, as well as 0.5 mM NADPH in 100 mM Tris buffer pH 8.0 at room temperature. Enzyme concentrations varied depending on the used substrates between 0.05 µM and 3 µM. Activities were measured in triplicates in 96-well-plates as the initial rate of NADPH depletion at 340 nm in a spectrophotometer (SpectraMax 190, Molecular Devices). Substrate depletion was calculated using the Beer-Lambert law, specific activity was derived by normalising on the used enzyme concentration, and fold-changes were calculated compared to wild type.

**3 Biotransformations of *Sr*IRED wt and T241A with different substrates** (see Figure S8)

**3.1 Reaction conditions**

Biocatalytic reactions were performed on a 1 or 2 mL-scale in 2 mL Eppendorf tubes and contained: buffer (KPi, 100 mM, pH 8), amine donor (1.6 eq., 40 mM final concentration; preparation stock solution: 80 mM of the amine donor were dissolved in 100 mM KPi buffer, pH 8; the pH was then re-adjusted to 8.0 with phosphoric acid), sodium formate (3 eq. 75 mM, 68.01 g/mol), $NADP^+$ (0.5 mM, 801.4 g/mol), FDH-QRN (10 µM) and *Sr*IRED wt or T241A (20 µM). The substrate was added last (25 mM final concentration as 1 M stock solution in DMSO). The reactions were incubated in a horizontal shaker at 30 °C for 4 h and 24 h, respectively. After the incubation time, 950 µL of the reaction mixture was taken and basified with KOH (10 M, 100 µL). The organic compounds were extracted with MTBE that contained the internal standard (IS toluene 20 mM, extraction with 2x500 µL), dried over $MgSO_4$ and measured by GC-FID. The conversions were calculated based on the substrate consumption with a calibration curve using substrate and toluene.

**3.2 Preparation of calibration curves for GC-FID**

1 mL samples containing buffer ($K_3PO_4$, 100 mM, pH 8) and substrates **1** - **4** or **6** - **8** (25 mM, 12.5 mM, 6 mM, 2.5 mM, 1 mM, and 0.5 mM final concentrations from 1 M stock solution in DMSO) were incubated for 24 h at 30 °C. After that, 950 µL of the mixtures were taken and brought to basic pH with KOH (10 M, 100 µL). The organic compound was extracted with MTBE that contained the internal standard (IS: toluene 20 mM, extraction with 2 x 500 µL), dried over $MgSO_4$ and measured by GC-FID. As the corresponding reference amine products

were not available, the product formation was confirmed by GC-MS. Example calibration curves are shown for substrates **1** to **4**.

**Substrate 1**

| | substrate (mM) | area IS | area substrate | area substrate/ area IS |
|---|---|---|---|---|
| cal1A | 25 | 3615 | 3522 | 0.97 |
| cal1B | 12.5 | 3553 | 1632 | 0.46 |
| cal1C | 6 | 3605 | 774 | 0.21 |
| cal1D | 2.5 | 3596 | 316 | 0.09 |
| cal1E | 1 | 3614 | 138 | 0.04 |

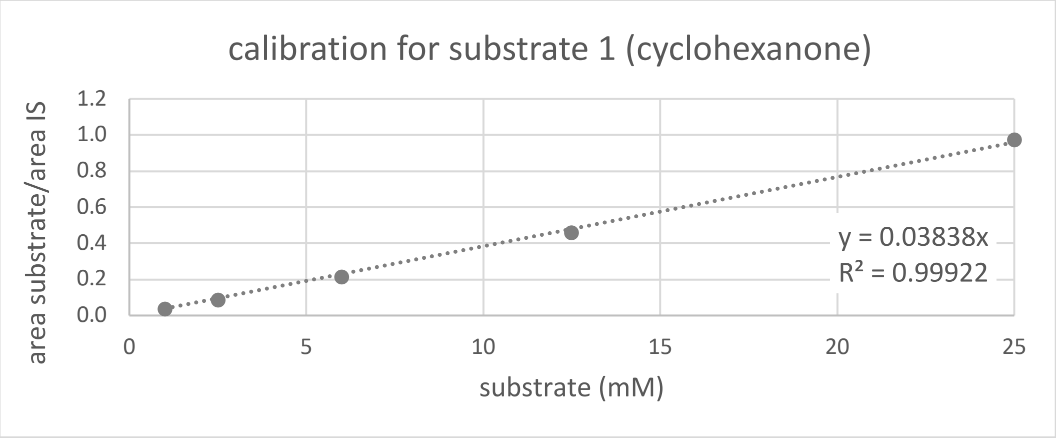

**Substrate 2**

| | substrate (mM) | area IS | area substrate | area substrate/ area IS |
|---|---|---|---|---|
| cal2A | 25 | 3604 | 2351 | 0.65 |
| cal2B | 12.5 | 3615 | 1190 | 0.33 |
| cal2C | 6 | 3621 | 563 | 0.16 |
| cal2D | 2.5 | 3543 | 218 | 0.06 |
| cal2E | 1 | 3673 | 105 | 0.03 |

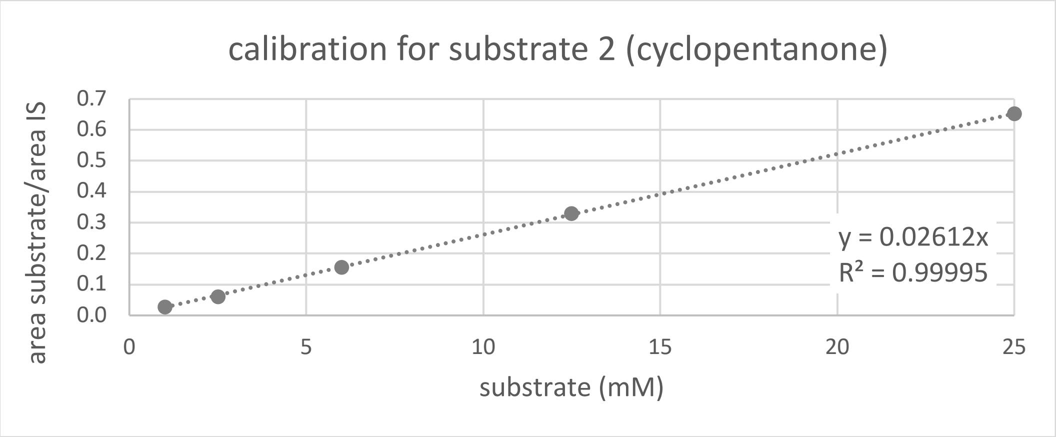

**Substrate 3**

| | substrate (mM) | area IS | area substrate | area substrate/ area IS |
|---|---|---|---|---|
| cal3A | 25 | 3763 | 4674 | 1.24 |
| cal3B | 12.5 | 3549 | 2146 | 0.60 |
| cal3C | 6 | 3625 | 1028 | 0.28 |
| cal3D | 2.5 | 3632 | 447 | 0.12 |
| cal3E | 1 | 4975 | 276 | 0.06 |

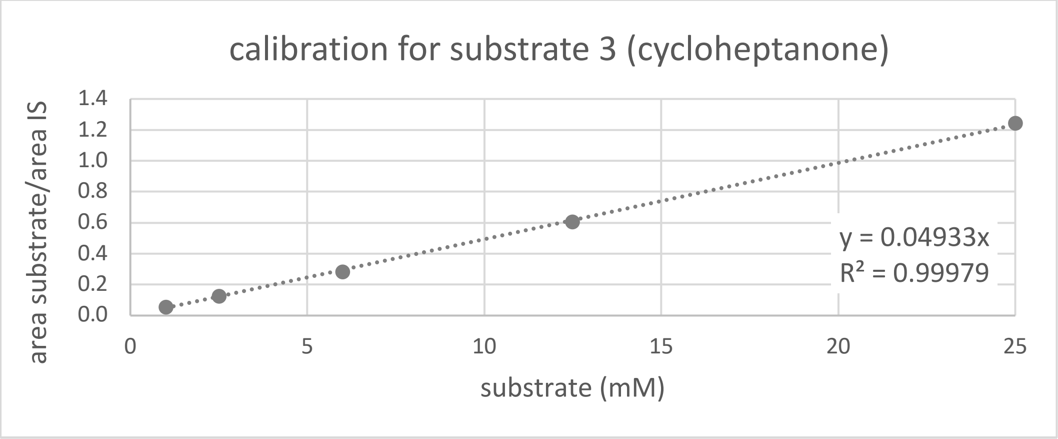

**Substrate 4**

| | substrate (mM) | area IS | area substrate | area substrate/ area IS |
|---|---|---|---|---|
| cal4A | 25 | 3808 | 3522 | 0.92 |
| cal4B | 12.5 | 3716 | 1704 | 0.46 |
| cal4C | 6 | 4125 | 927 | 0.22 |
| cal4D | 2.5 | 6819 | 675 | 0.10 |
| cal4E | 1 | 5303 | 226 | 0.04 |

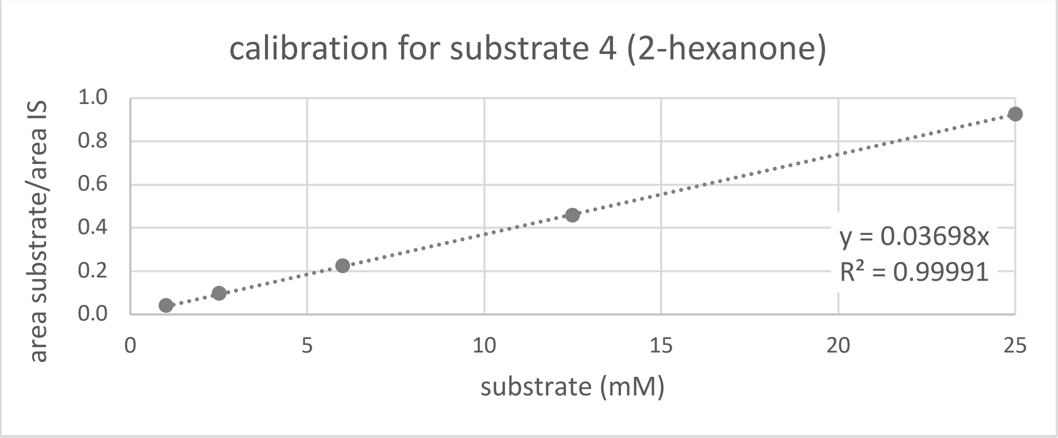

##### 3.3 GC-FID methods and retention times

Column: The conversions were measured by GC-FID using an Agilent J&W DB-1701 (30 m, 250 µm, 0.25 µm) column. Carrier gas: $H_2$
Method: constant pressure 6.9 psi; temperature program: 80 °C, hold 6.5 min; 10 °C min$^{-1}$ to 160 °C, hold 0 min; 20 °C min$^{-1}$ to 280 °C, hold 0 min.

**Example retention times in GC-FID for substrates 1 to 4 (**internal standard toluene: 2.9 min; cosolvent DMSO: 7.7 min; n.m. = not measured)

| primary amine | alcohol | substrate | unidentified (only visible in GC-FID) | amine product | imine |
|---|---|---|---|---|---|
| 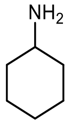 4.1 min | 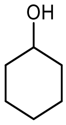 5.7 min | 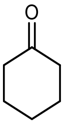 **1** 6.2 min | 8.4 min | 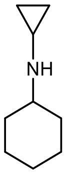 10.0 min | 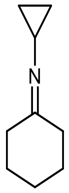 11.9 min |
| n.m. | n.m. | 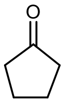 **2** 3.9 min | 5.3 min | 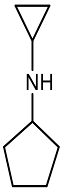 7.2 min | 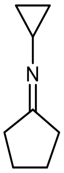 10.1 min |
| n.m. | n.m. | 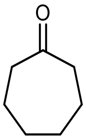 **3** 9.7 min | 11.5 min | 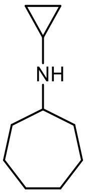 12.9 min | 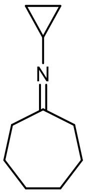 14.1 min |
| n.m. | 3.7 min | 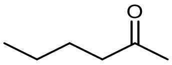 **4** 3.5 min | 5.7 min | 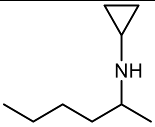 7.0 min | 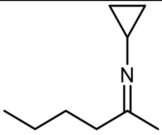 8.8 min |

**Additional primary amines measured**

| | | | |
|---|---|---|---|
| 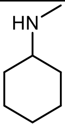 5.1 min | 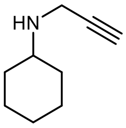 10.8 min | 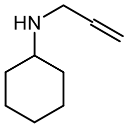 9.5 min | 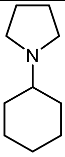 12.2 min |

#### 3.4 GC-FID chromatograms for the biocatalytic reactions

**Substrate 1-a**

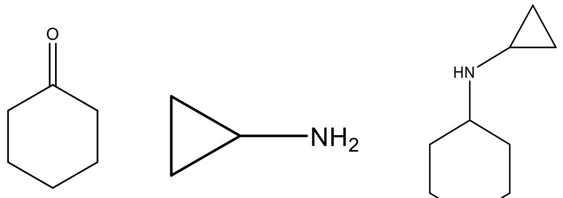

m/z: 98.07 (100.0%), 99.08 (6.6%)

m/z: 57.06 (100.0%), 58.06 (3.3%)

m/z: 139.14 (100.0%), 140.14 (9.9%)

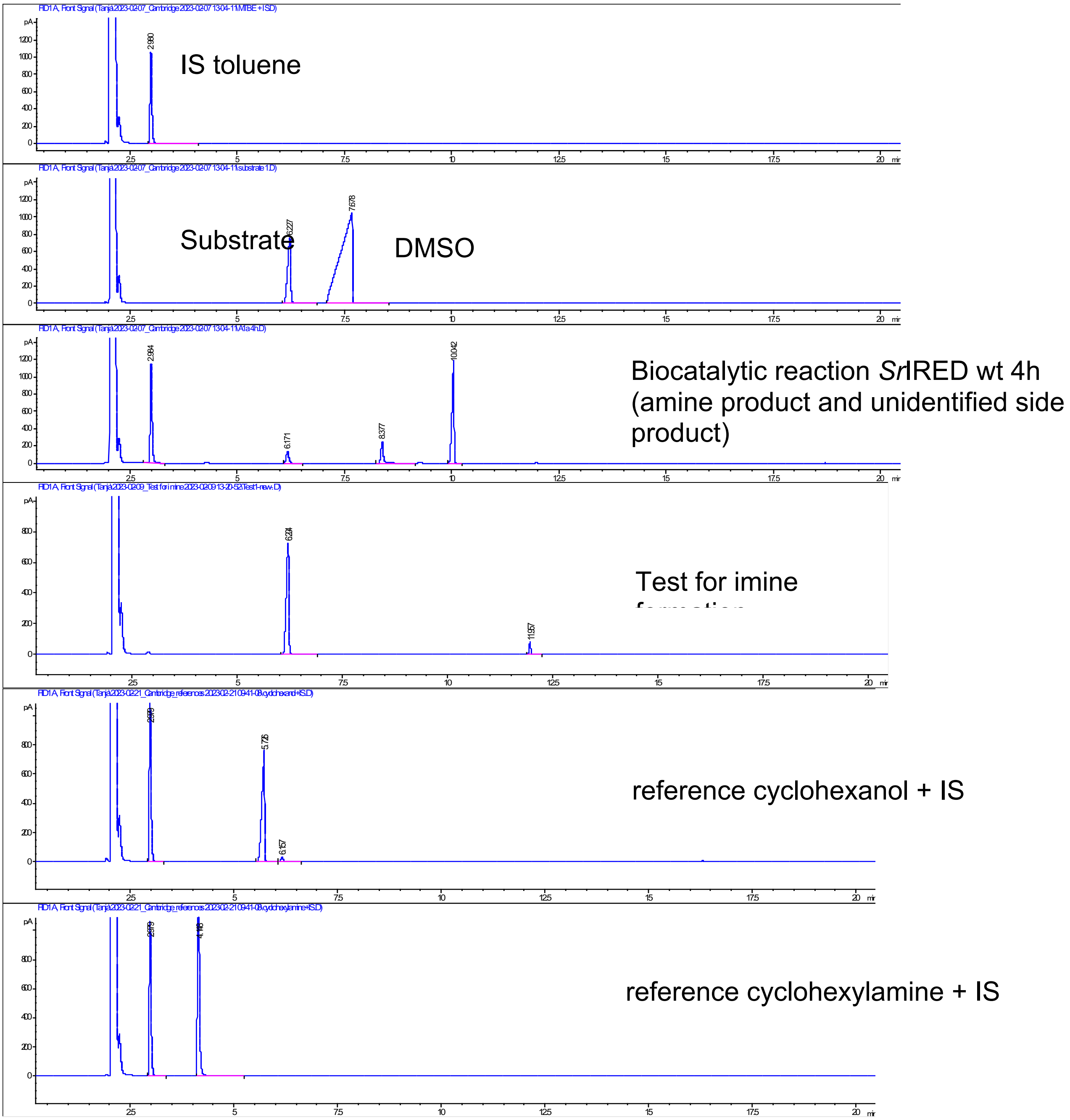

#### **Substrate 2-a**

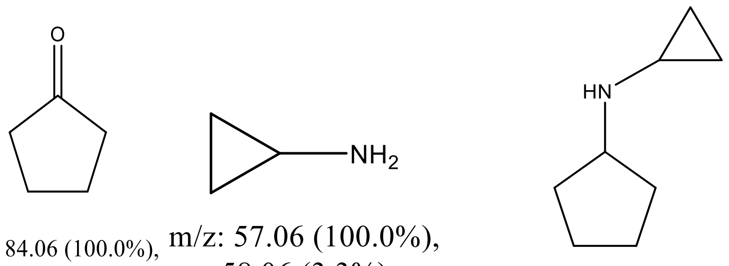

m/z: 84.06 (100.0%), 85.06 (5.5%)

m/z: 57.06 (100.0%), 58.06 (3.3%)

m/z: 125.12 (100.0%), 126.12 (9.0%)

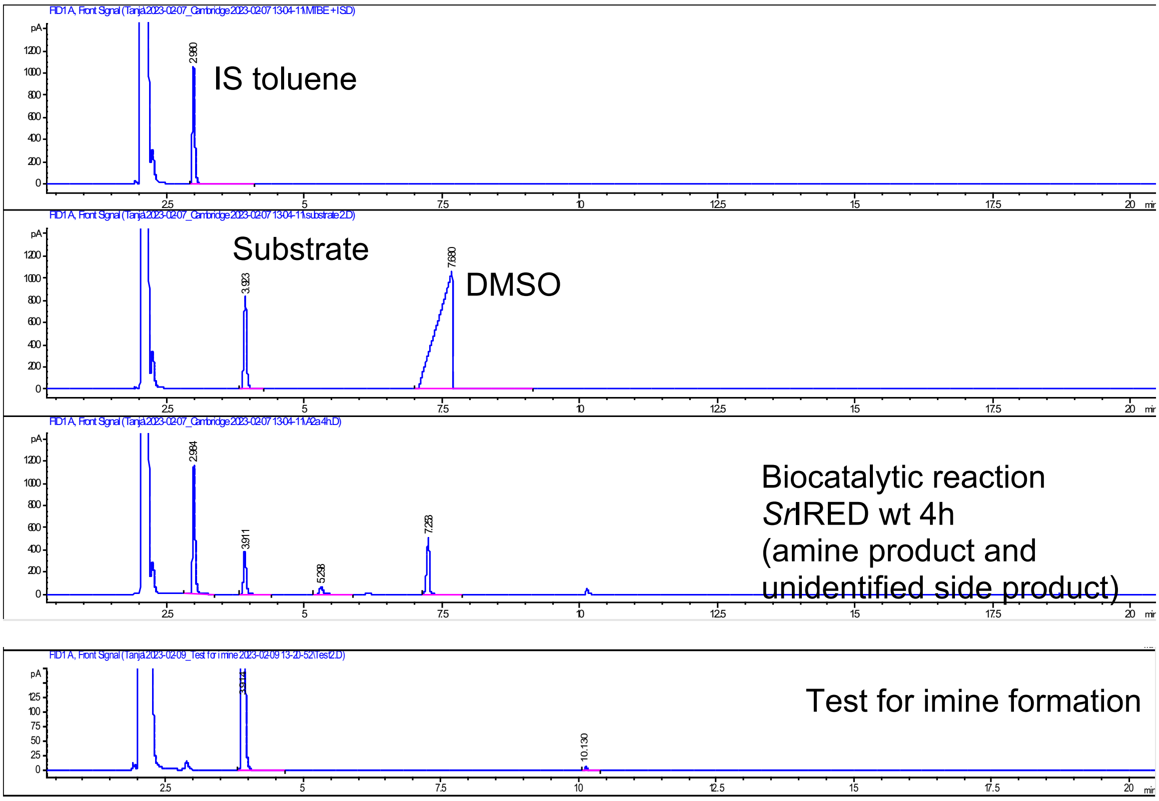

#### Substrate 3-a

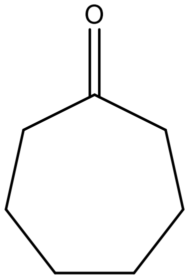

m/z: 112.09 (100.0%), 113.09 (7.6%)

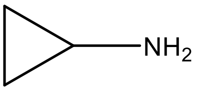

m/z: 57.06 (100.0%), 58.06 (3.3%)

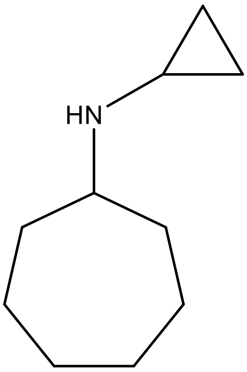

m/z: 153.15 (100.0%), 154.16 (11.0%)

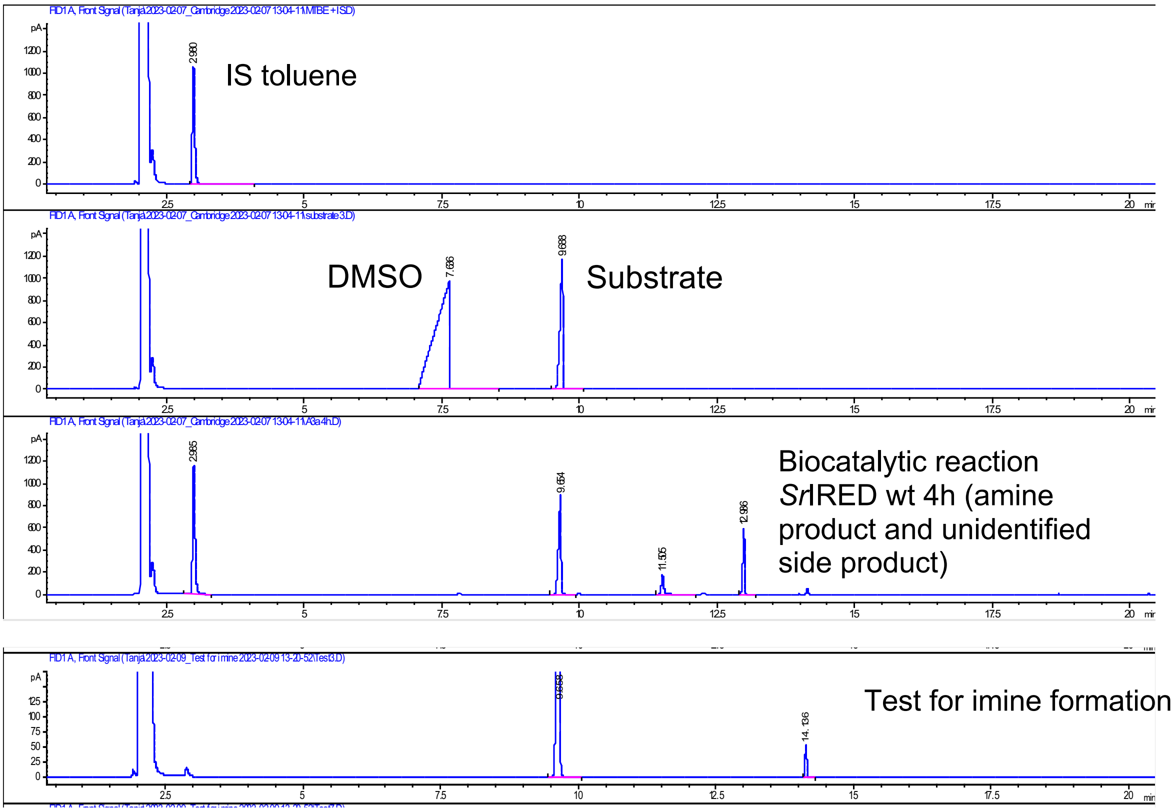

#### Substrate 4-a

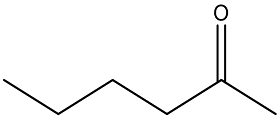

m/z: 100.09 (100.0%), 101.09 (6.5%)

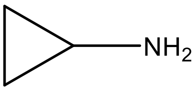

m/z: 57.06 (100.0%), 58.06 (3.3%)

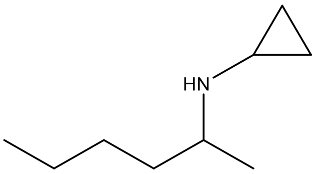

m/z: 141.2 (100.0%), 142.2 (10.0%)

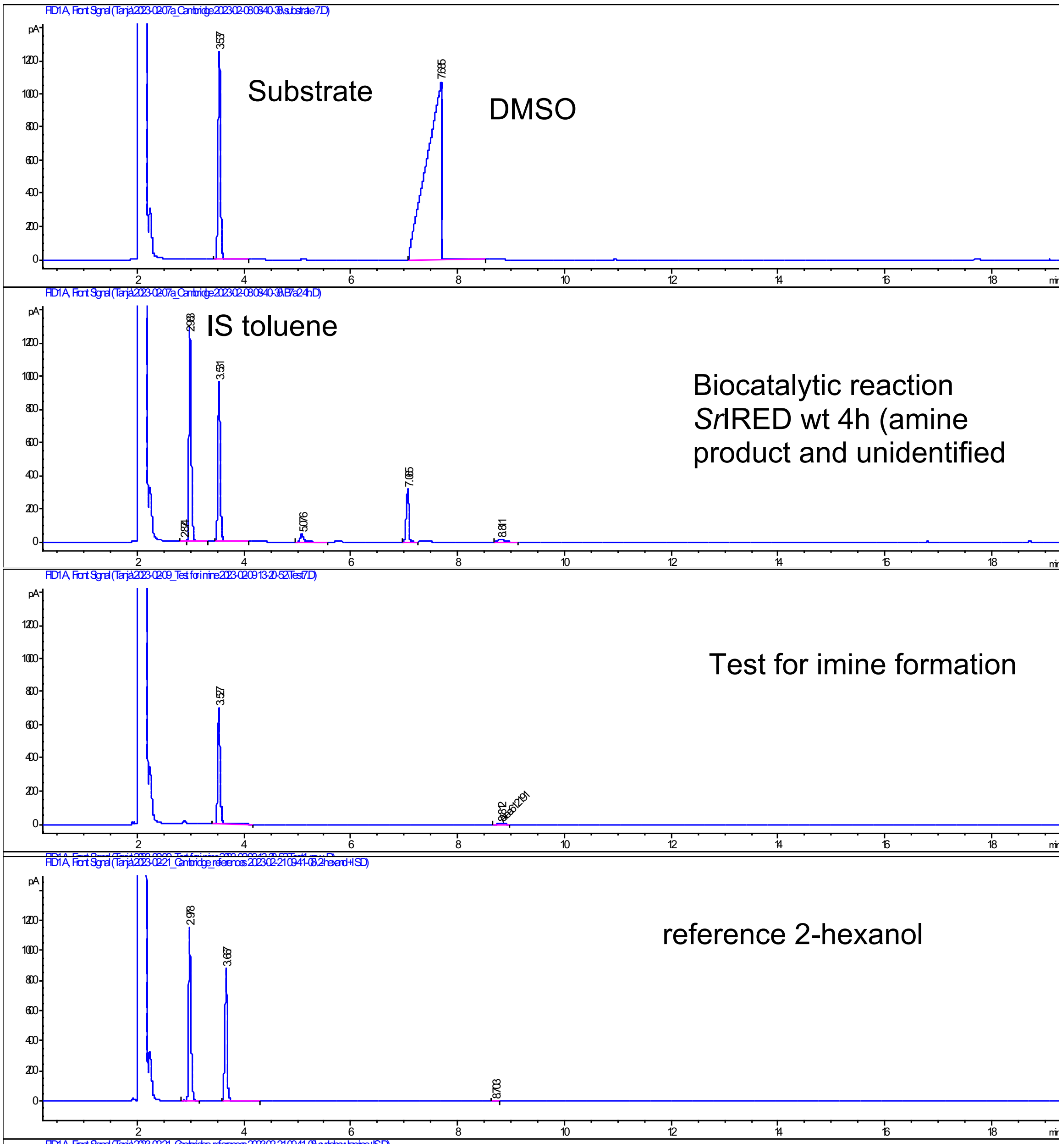

#### Substrate 1-b

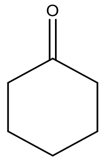

m/z: 98.07 (100.0%), 99.08 (6.6%)

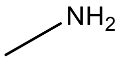

m/z: 31.0 (100.0%), 32.0 (1.5%)

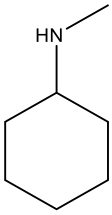

m/z: 113.12 (100.0%), 114.12 (7.9%)

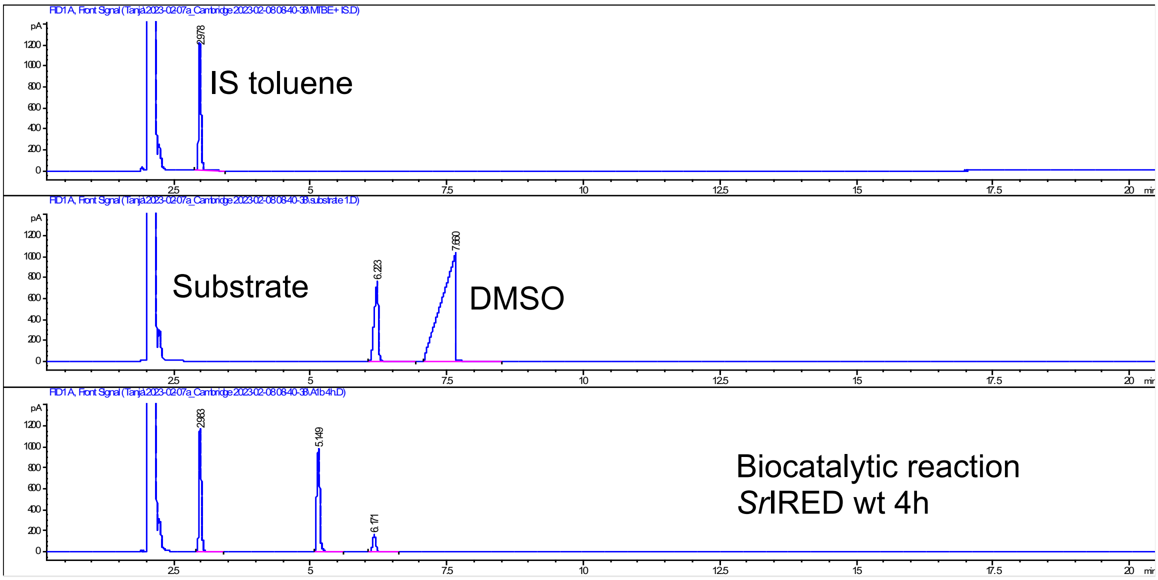

#### Substrate 1-c

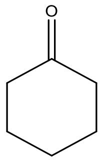

m/z: 98.07 (100.0%), 99.08 (6.6%)

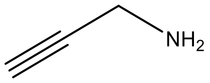

m/z: 55.0 (100.0%), 56.0 (3.7%)

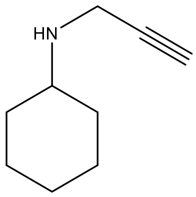

m/z: 137.12 (100.0%), 138.12 (10.1%)

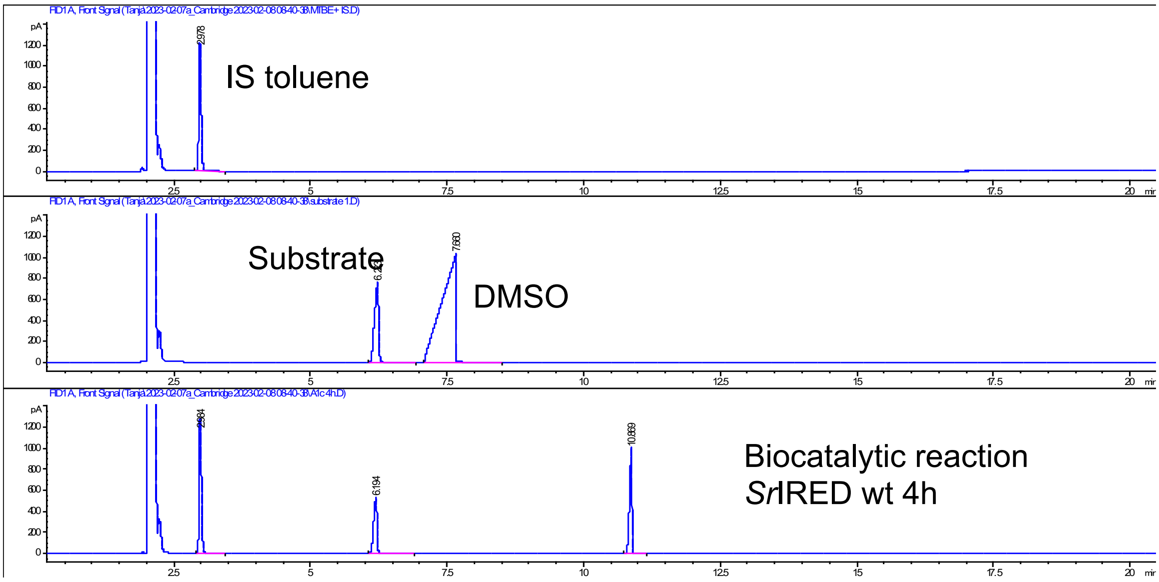

#### Substrate 1-d

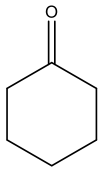

m/z: 98.07 (100.0%), 99.08 (6.6%)

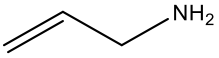

m/z: 57.1 (100.0%), 58.1 (3.7%)

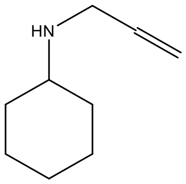

m/z: 139.1 (100.0%), 140.1 (10.3%)

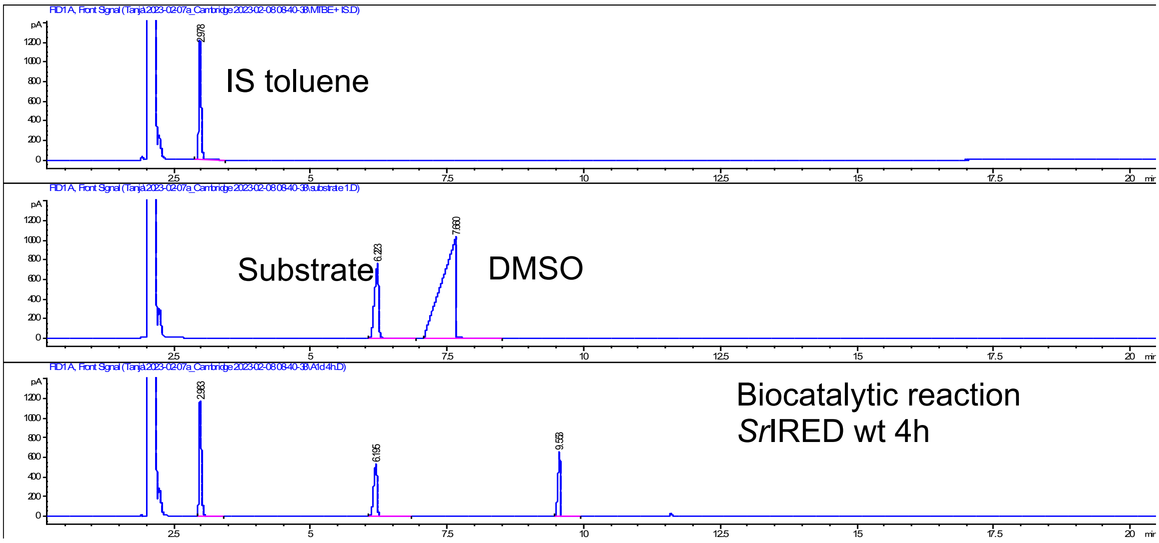

#### Substrate 1-e

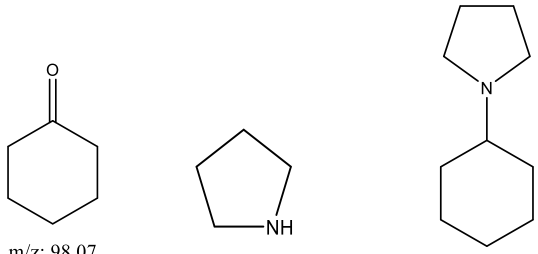

m/z: 98.07 (100.0%), 99.08 (6.6%)

m/z: 71.1 (100.0%), 72.1 (4.8%)

m/z: 153.2 (100.0%), 154.2 (11.0%)

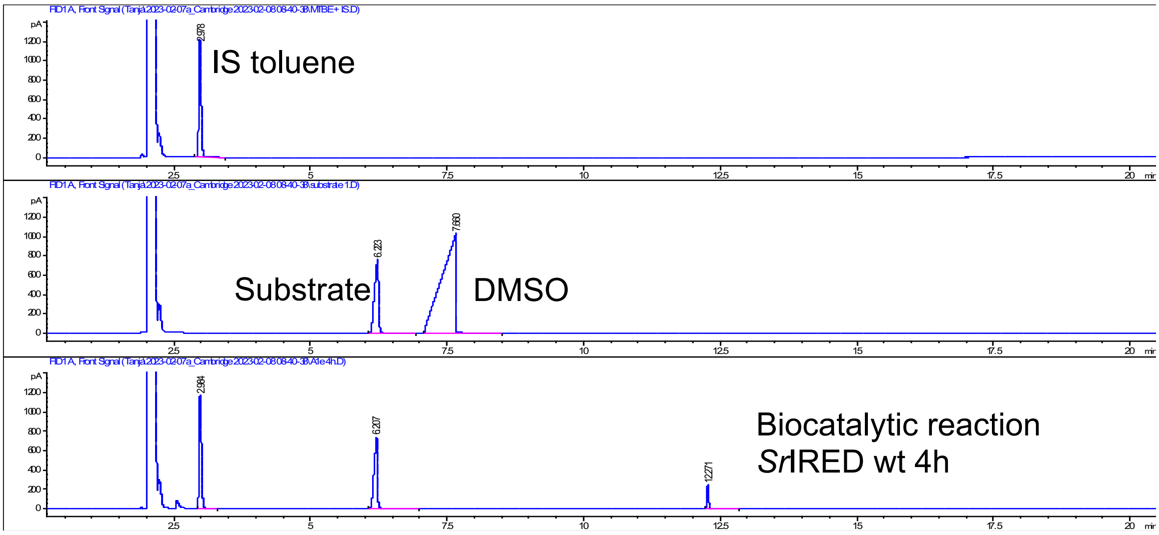

#### Substrate 8a

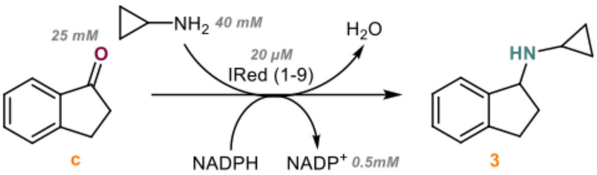

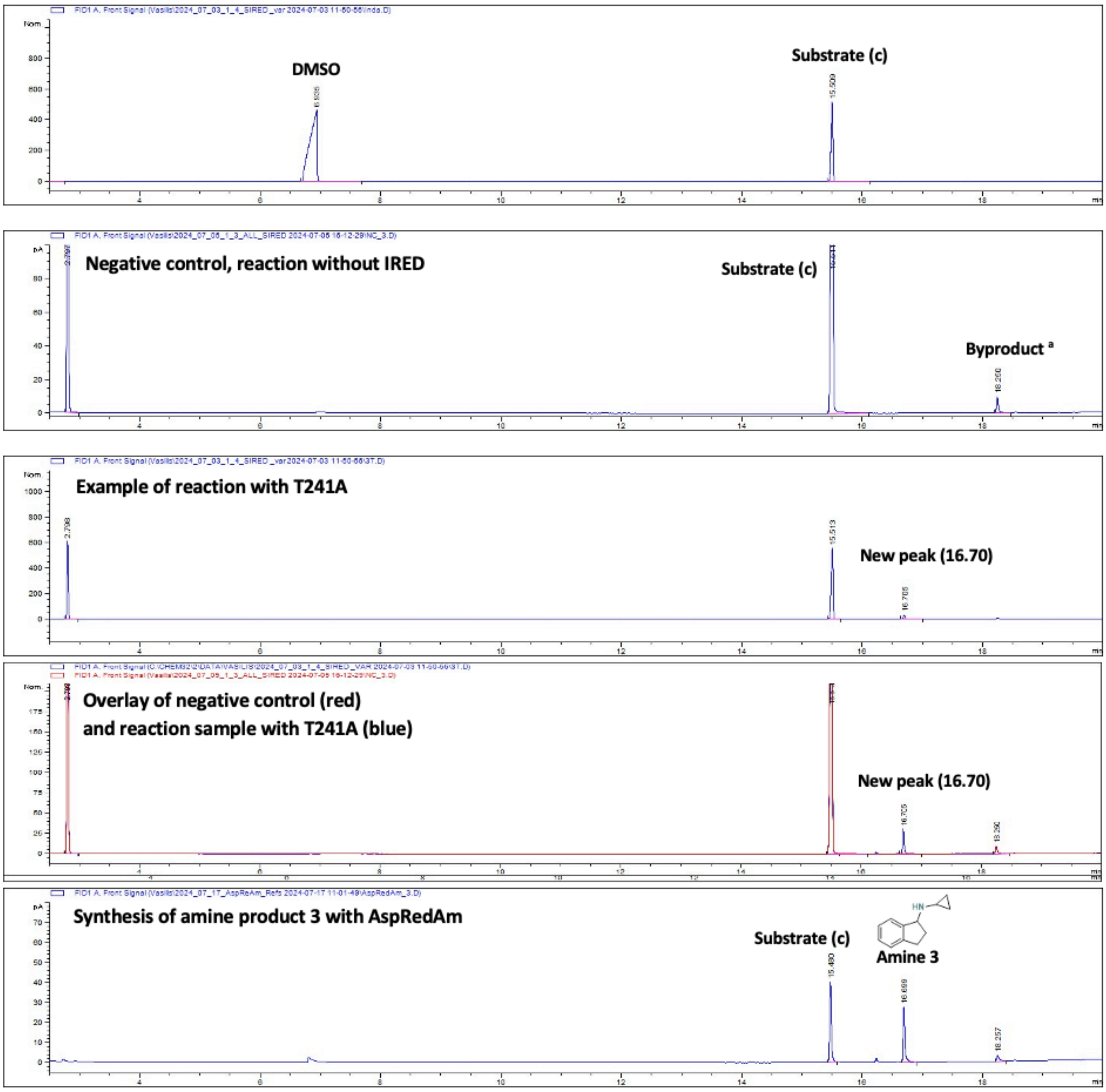

[a] The byproduct has been identified as the imine intermediate form during extraction in organic solvent. The peak of the imine is normally visible also in GC-MS and the MS peak is the one of the mass of the imine.

#### Substrate 7a

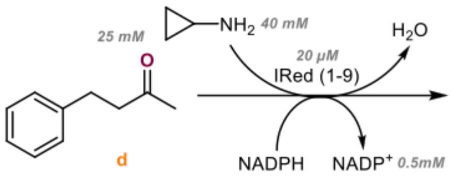

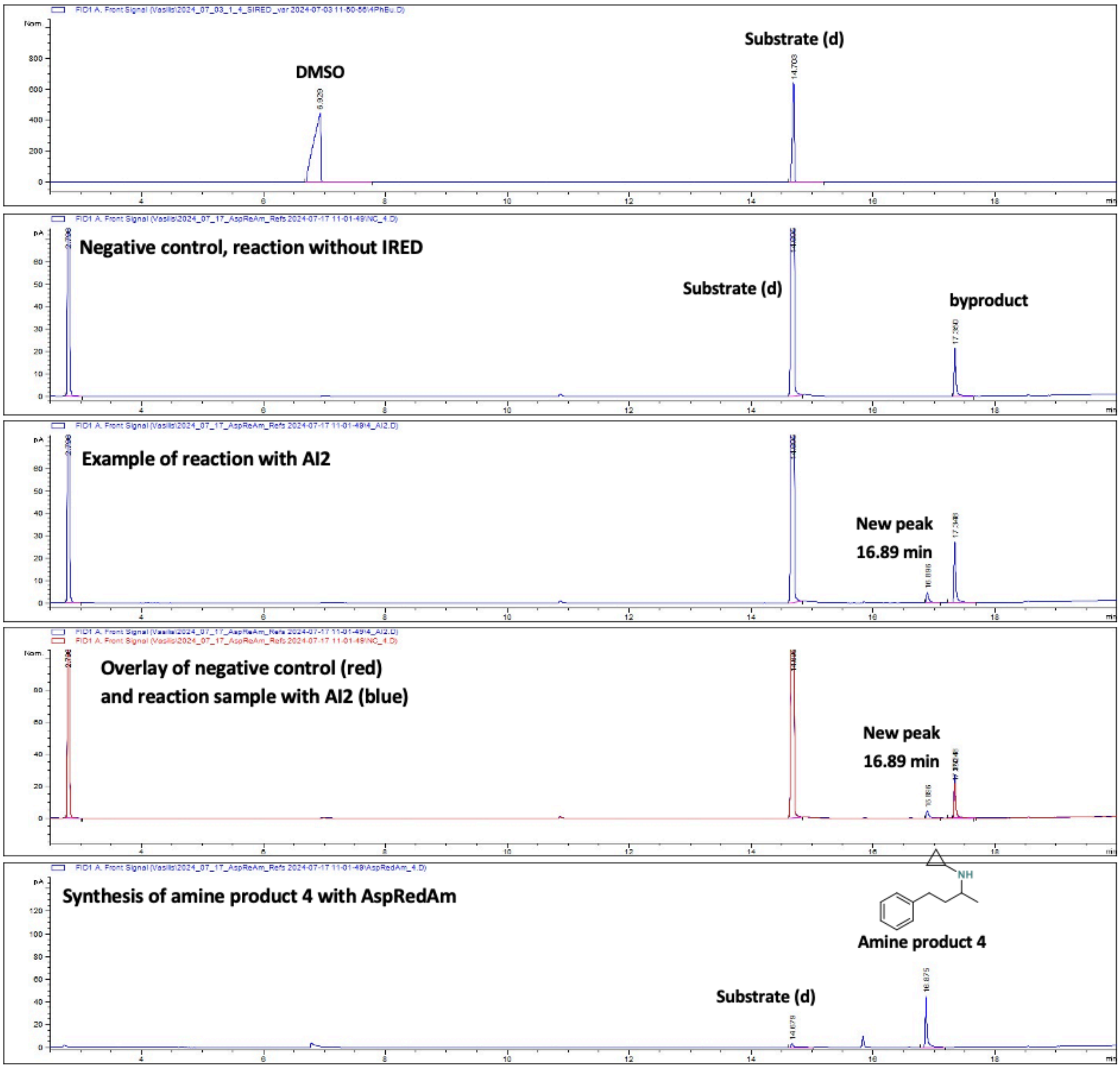

#### 3.5 GC-MS spectra to confirm/identify the peaks of substrate and product

**GC-MS spectrum of Sample *Sr*IRED T241A with 1a (4h)**

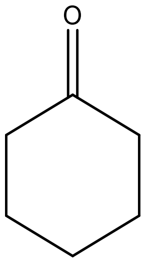

m/z: 98.07 (100.0%), 99.08 (6.6%)

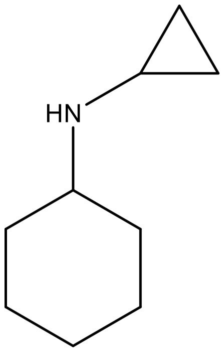

m/z: 139.14 (100.0%), 140.14 (9.9%)

GC-FID: 6.2 min substrate, 8.4 min new peak 1, 10.0 min new peak 2

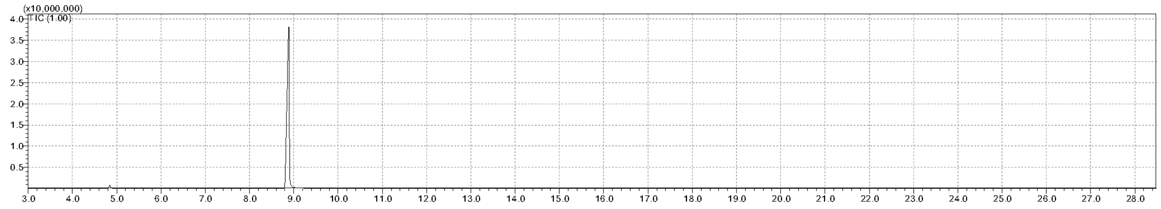

Zoom in

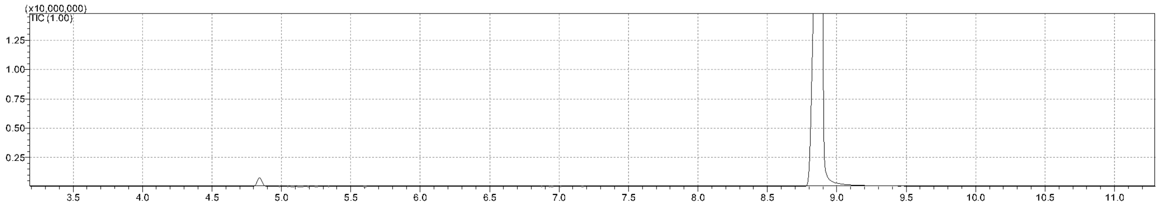

Peak 1: 4.8 min substrate

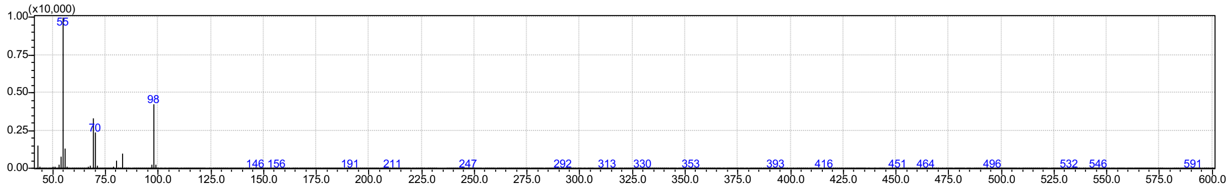

Peak 2: 8.9 min amine product confirmed

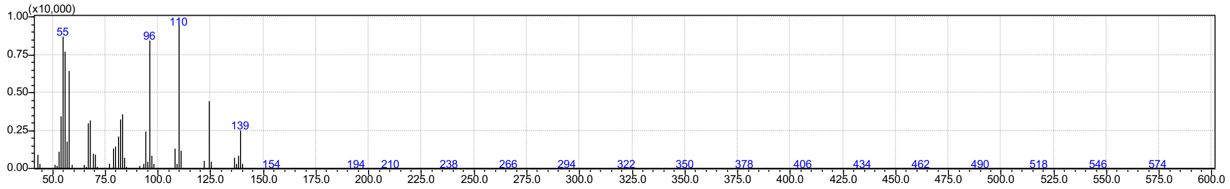

In the GC-MS chromatogram we did not detect the side-product peak that was visible in the GC-FID chromatogram (and elutes before the desired product).

#### GC-MS spectrum of Sample *Sr*IRED T241A with 2a (4h)

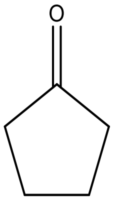

m/z: 84.06 (100.0%), 85.06 (5.5%)

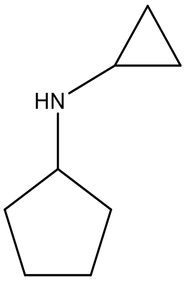

m/z: 125.12 (100.0%), 126.12 (9.0%)

GC-FID: 3.9 min substrate, 5.3 min new peak 1, 7.3 min new peak 2

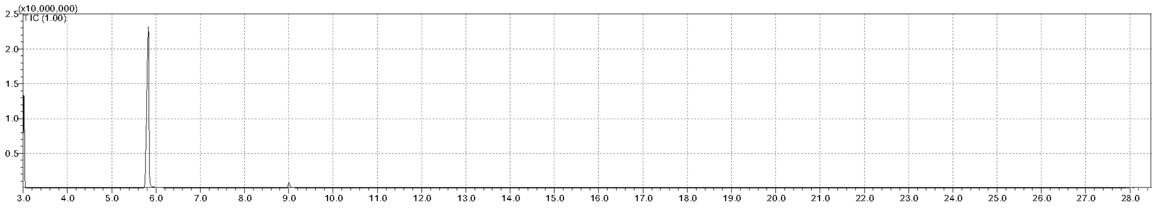

Zoom in:

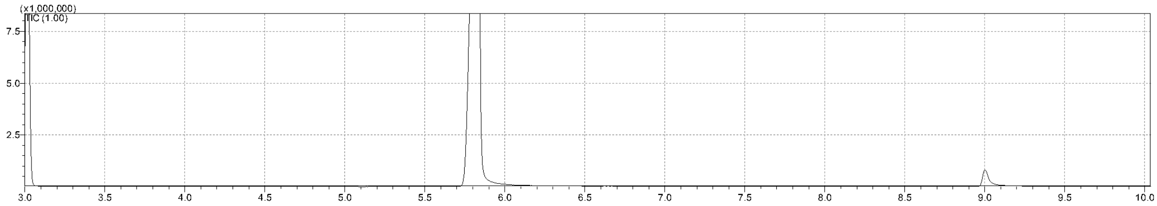

Peak 1: 3.0 min substrate

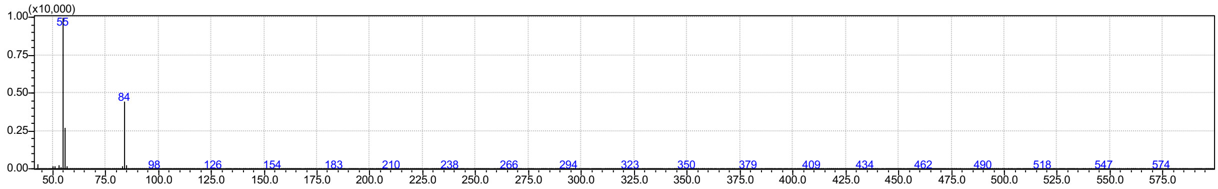

Peak 2: 5.8 min amine product confirmed

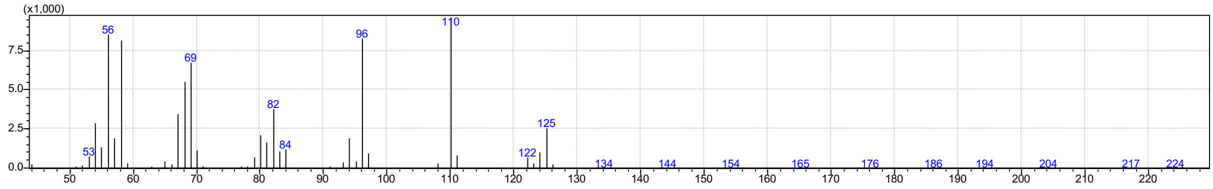

We did not detect in this GC-MS chromatogram the side-product peak that was visible in the GC-FID chromatogram (and elutes before the desired product).

#### GC-MS spectrum of Sample *Sr*IRED T241A with 3a (4h)

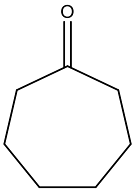

m/z: 112.09 (100.0%), 113.09 (7.6%)

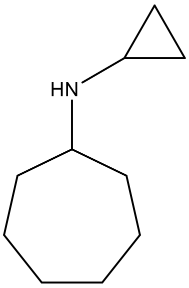

m/z: 153.15 (100.0%), 154.16 (11.0%)

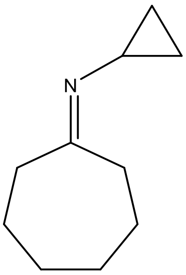

m/z: 151.1 (100.0%), 152.1 (11.4%)

GC-FID 9.7 min substrate, 11.5 min product 1, 13.0 min product 2

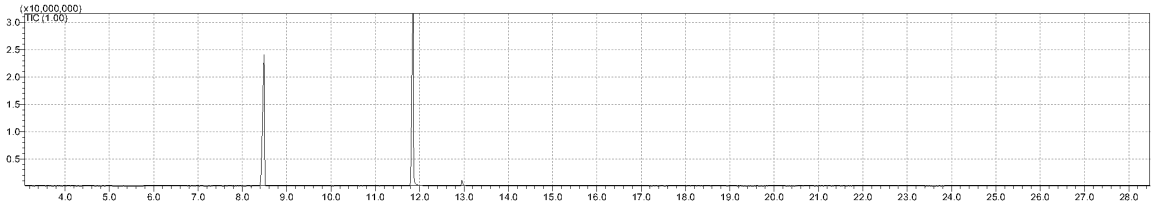

Peak 1: 8.5 min substrate

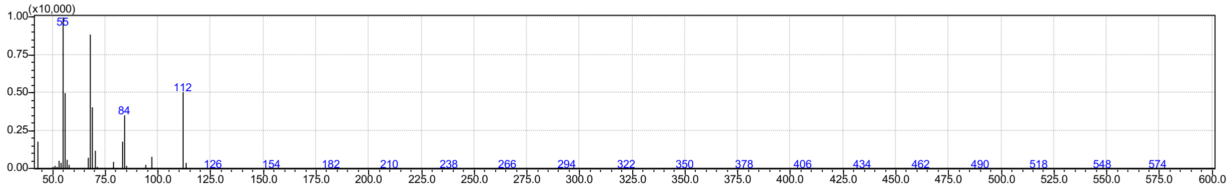

Peak 2: 11.8 min amine product confirmed

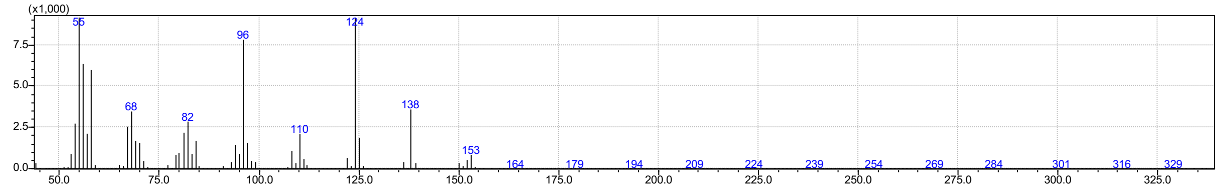

Peak 3: 13 min imine

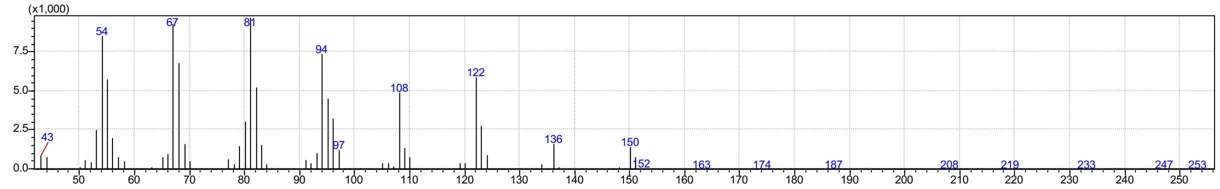

#### GC-MS spectrum of Sample *Sr*IRED T241A with 4a (4h)

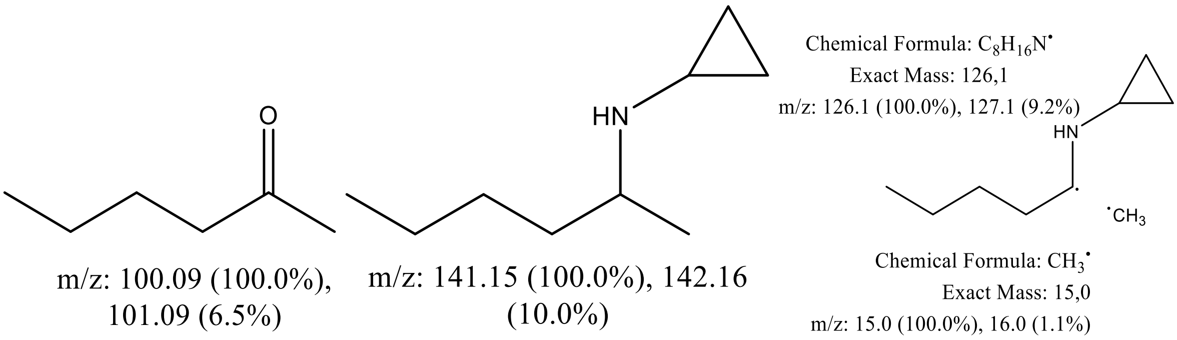

GC-FID: 3.5 min substrate, 5.1 min product 1, 7.1 min product 2

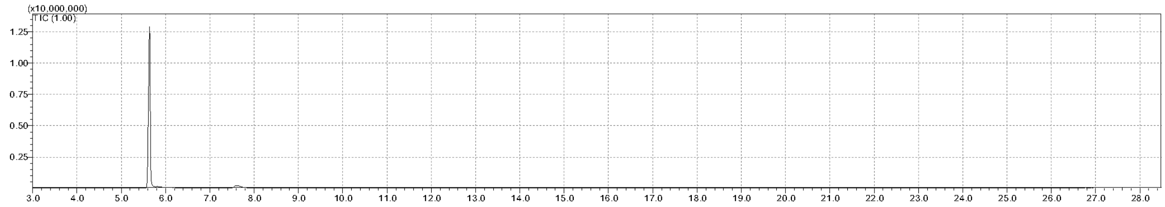

Zoom in:

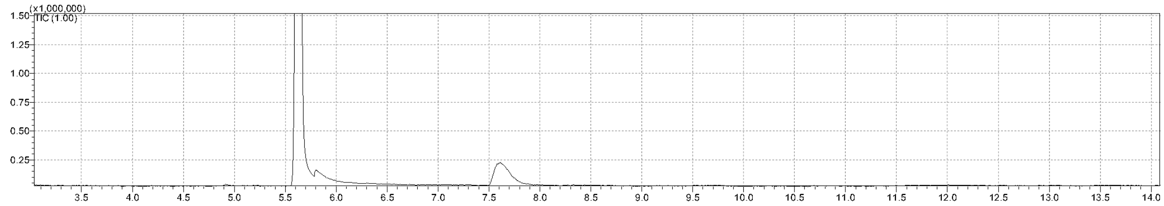

Peak 1: 5.6 min, m/z – [Me] = 126,

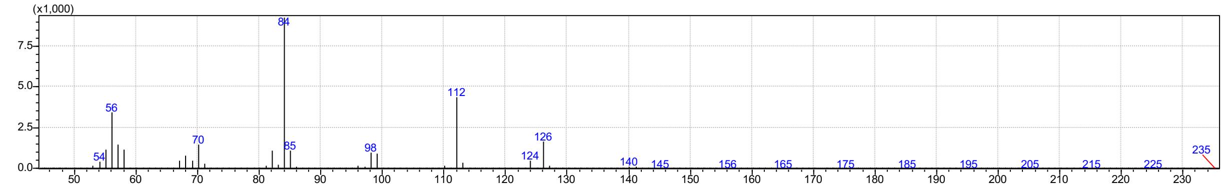

Peak 2 Small peak after peak 1: at ca. 5.7 min, m/z – [Me] – [$H_2$] = 124

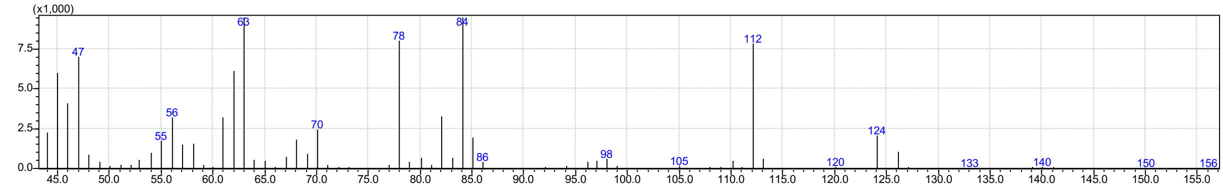

Peak 3: 7.6 min imine

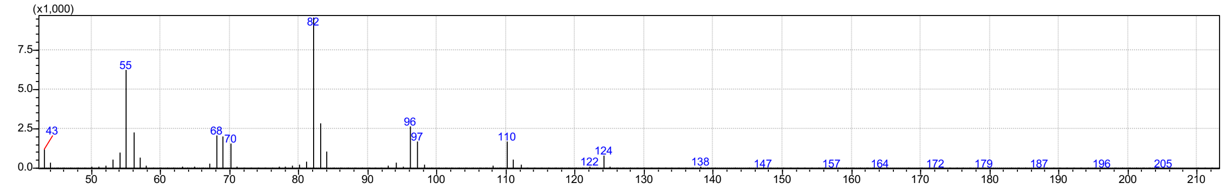

#### GC-MS spectrum of Sample *Sr*IRED wt with 1b (4h)

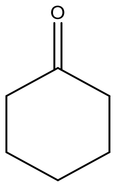

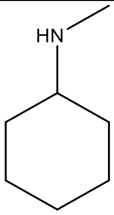

m/z: 98.1 (100.0%), 99.1 (6.6%) m/z: 113.12 (100.0%), 114.12 (7.9%)

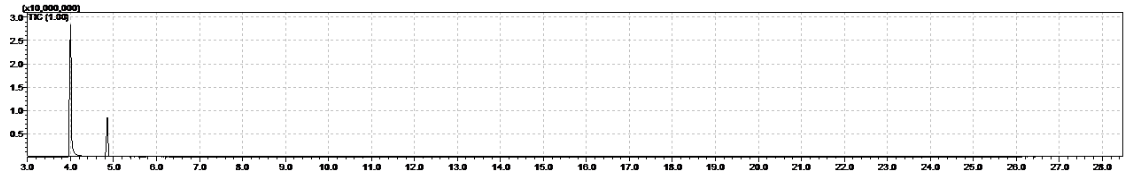

Peak 1: 4.0 min amine product confirmed

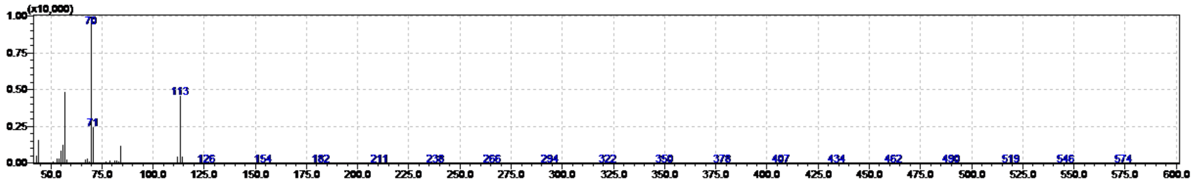

Peak 2: 4.9 min substrate

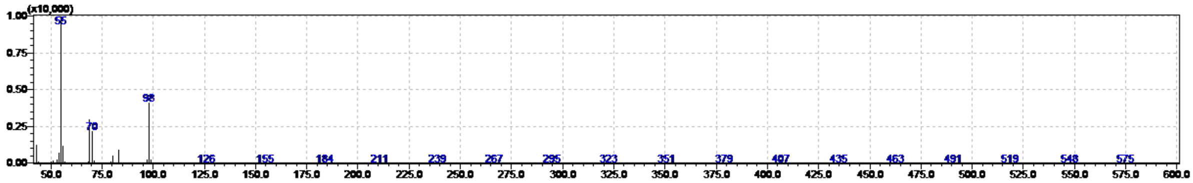

### GC-MS spectrum of Sample *Sr*IRED wt with 1c (4h)

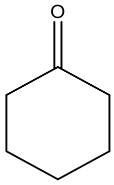

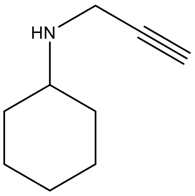

m/z: 98.1 (100.0%), 99.1 (6.6%) m/z: 137.12 (100.0%), 138.12 (10.1%)

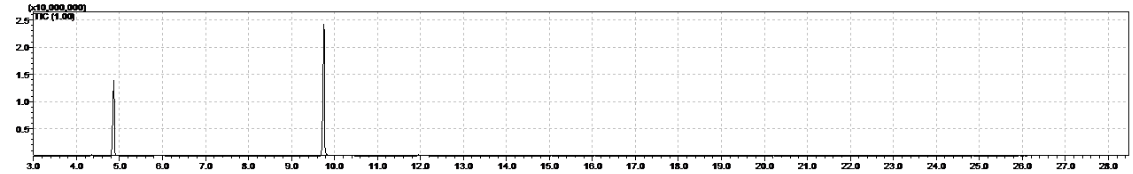

Peak 1: 4.9 min substrate

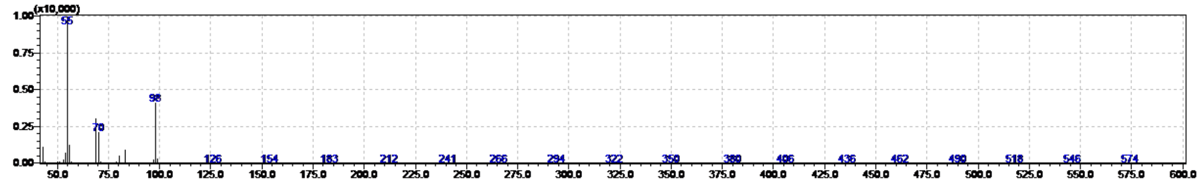

Peak 2: 9.7 min amine product confirmed

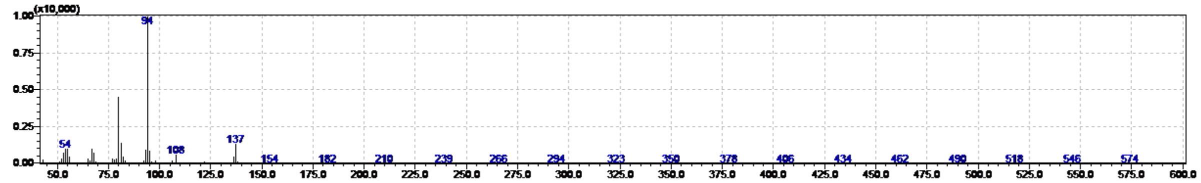

### GC-MS spectrum of Sample *Sr*IRED wt with 1d (24h)

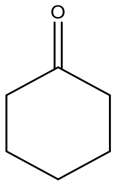

m/z: 98.1 (100.0%), 99.1 (6.6%)

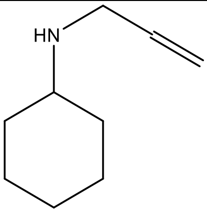

m/z: 139.1 (100.0%), 140.1 (10.3%)

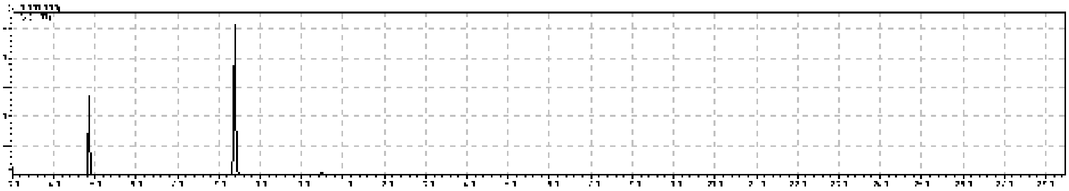

Peak 1: 4.9 min substrate

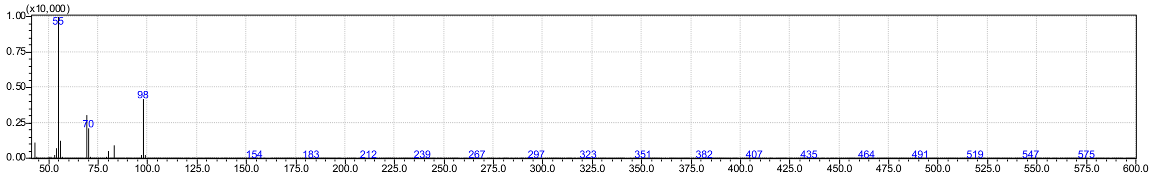

Peak 2: 8.4 min amine product confirmed

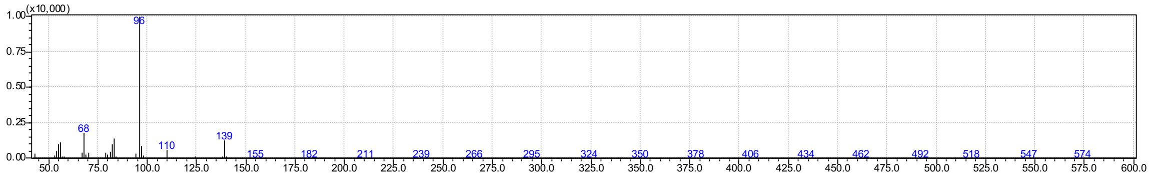

#### GC-MS spectrum of Sample *Sr*IRED wt with 1e (24h)

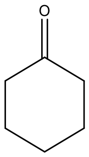

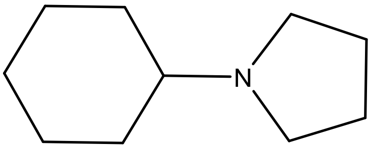

m/z: 98.1 (100.0%), 99.1 (6.6%) m/z: 153.2 (100.0%), 154.2 (11.0%)

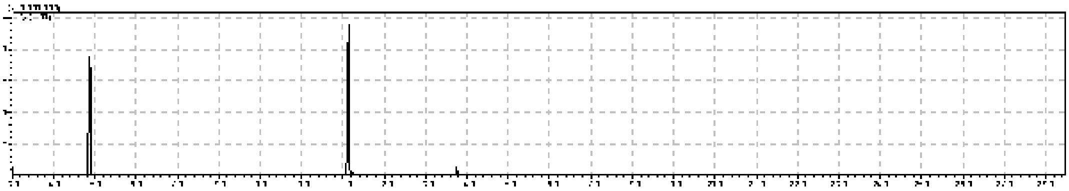

Peak 1: 4.9 min substrate

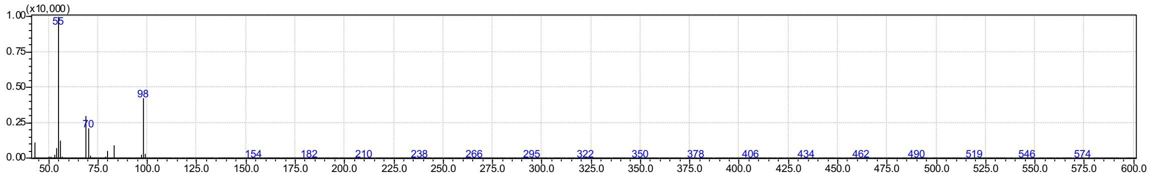

Peak 2: 11.1 min amine product confirmed

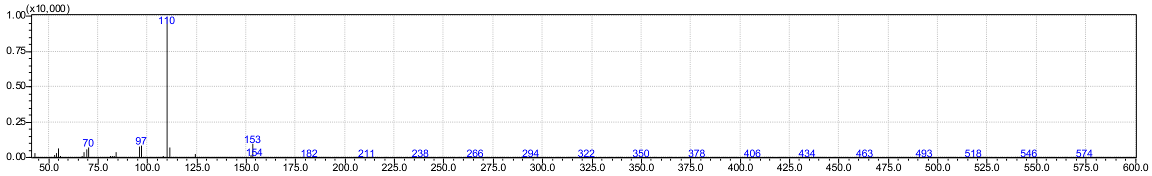

#### GC-MS spectrum 8a

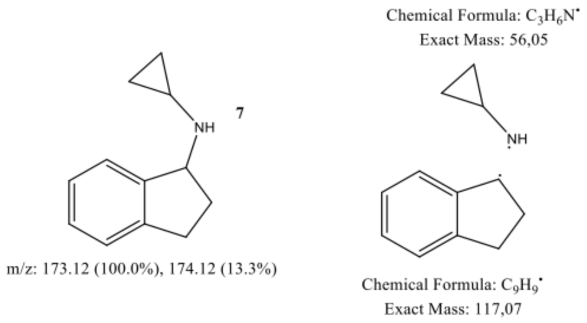

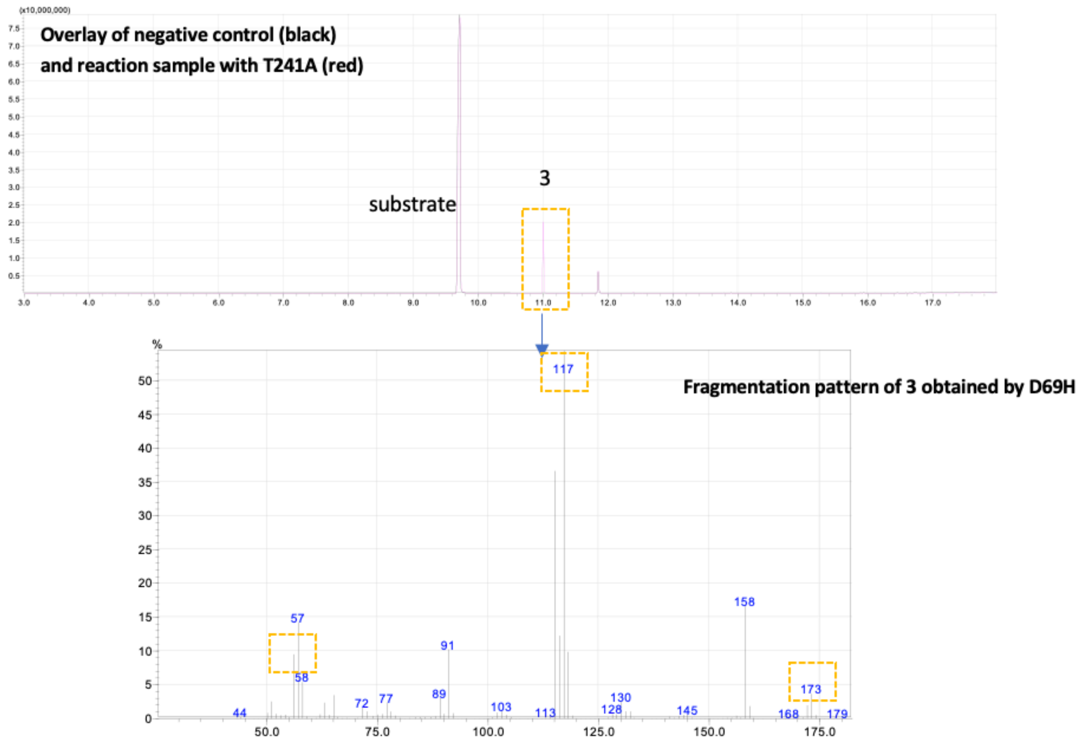

The same fragmentation pattern for the reference product (**11 min**) synthesized by AspRedAm

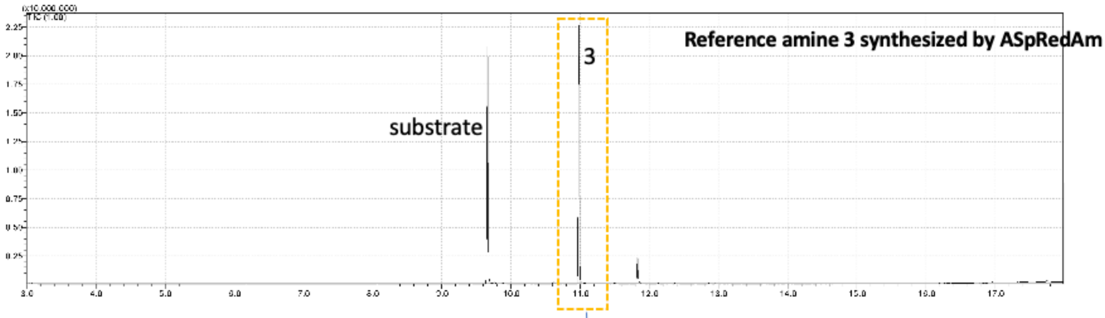

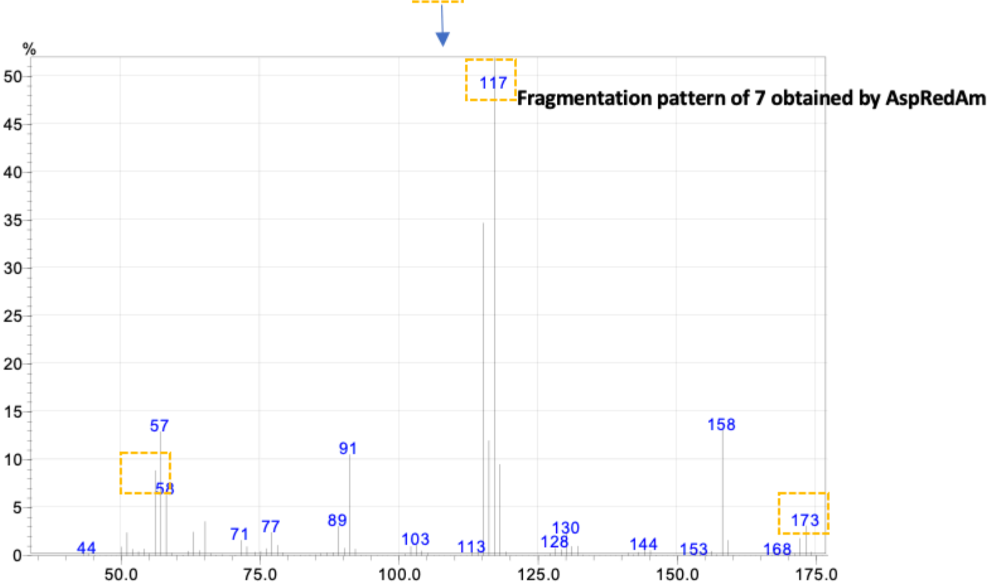

#### GC-MS spectrum 7a:

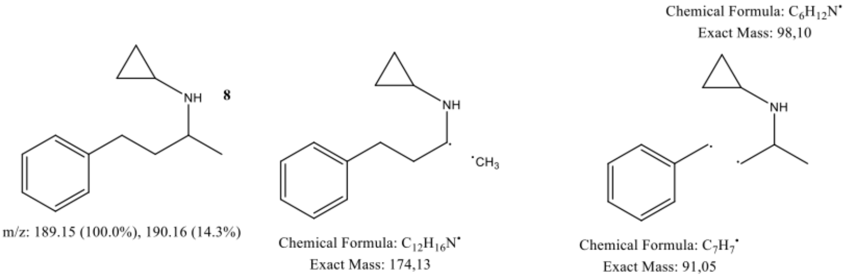

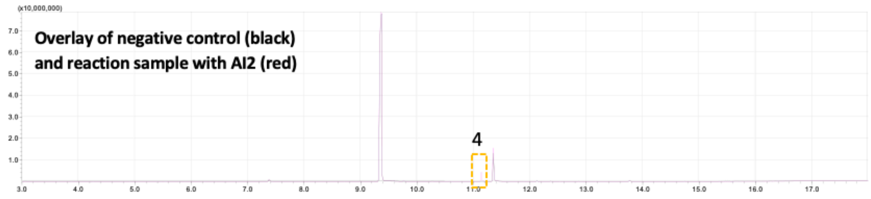

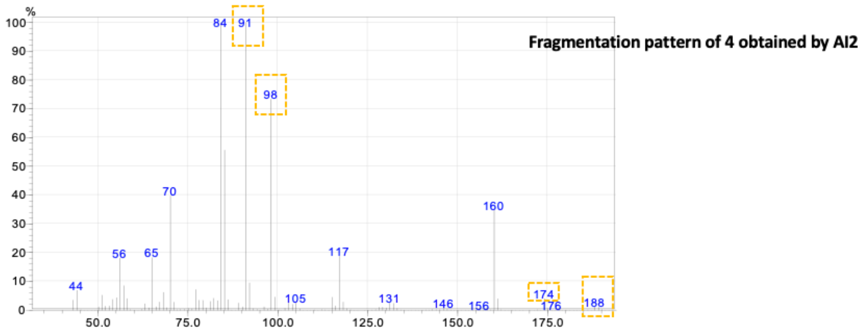

The same fragmentation pattern for the reference product (**11.2 min**) synthesized by AspRedAm

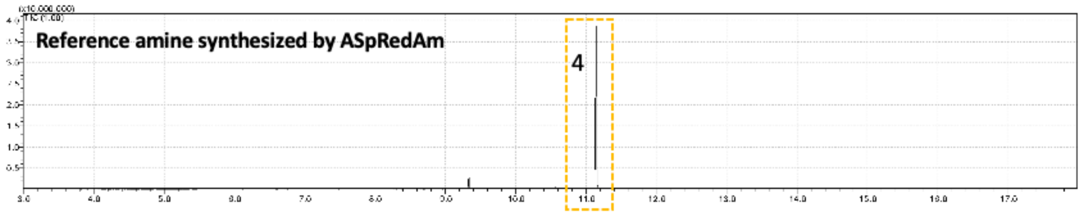

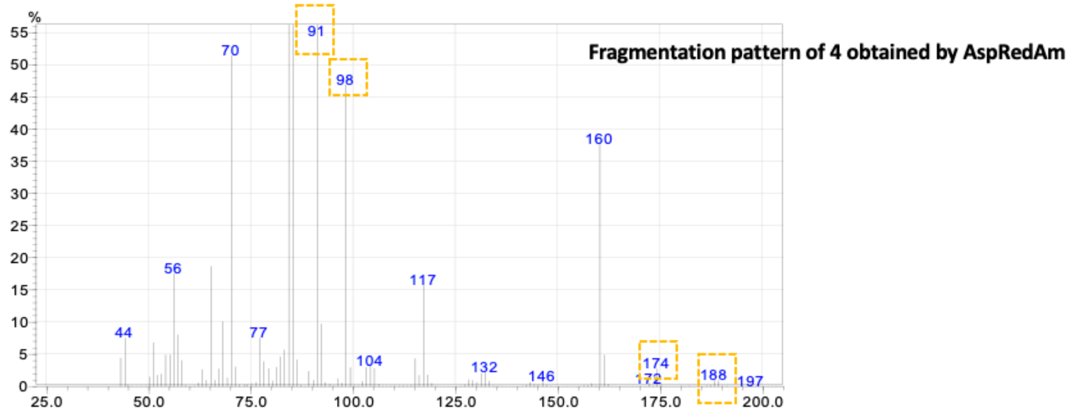

#### 4 Biotransformations of *Sr*IERD wt and selected mutants with 1 a (see Figure S15)

##### 4.1 Reaction conditions

Biocatalytic reactions were performed on a 5 mL-scale in 5 mL Eppendorf tubes or 1 mL scale in 2 mL Eppendorf tubes and contained: buffer (KPi, 100 mM, pH 8), amine donor (1.6 eq., 40 mM final concentration; preparation of the stock solution: 80 mM of the amine donor were dissolved in 100 mM KPi buffer, pH 8; the pH was then re-adjusted to 8.0 with phosphoric acid), sodium formate (3 eq. 75 mM, 68.01 g/mol), $NADP^+$ (0.5 mM or 1 mM, 801.4 g/mol), FDH-QRN (10 µM) and *Sr*IRED wt or one of the variants (10 and 5 µM and also 2.5 µM for T241G P203A). The substrate was added last (25 mM final concentration as 1 M stock solution in DMSO). The reactions were incubated in a horizontal shaker at 30 °C. After the incubation time (see tables and figures), 950 µL of the reaction mixture were taken and basified with KOH (10 M, 100 µL). The organic compounds were extracted with MTBE that contained the internal standard (IS toluene 20 mM, extraction with 2 x 500 µL), dried over $MgSO_4$ and measured by GC-FID. The conversions were calculated based on the substrate consumption with a calibration curve using substrate and toluene.

##### 4.2 Preparation of calibration curves for GC-FID

1 mL samples containing buffer ($K_3PO_4$, 100 mM, pH 8) and substrates (25 mM, 12.5 mM, 6 mM, 2.5 mM and 1 mM final concentrations from 1 M stock solution in DMSO) were incubated for 24 h. After that, 950 µL of the mixtures were taken and brought to basic pH with KOH (10 M, 100 µL). The organic compound was extracted with MTBE that contained the internal standard (IS toluene 20 mM, extraction with 2 x 500 µL), dried over $MgSO_4$ and measured by GC-FID. As the corresponding reference amine products were not available, the product formation was confirmed by GC-MS.

##### 4.3 GC-FID methods

*Column*: the conversions were measured by GC-FID using an Agilent J&W DB-1701 (30 m, 250 µm, 0.25 µm) column. Carrier gas: $H_2$

*Method A:* for the amination of cyclohexanone with cyclopropylamine: constant pressure 6.9 psi; temperature program: 80 °C, hold 6.5 min; 10 °C $min^{-1}$ to 160 °C, hold 0 min; 20 °C $min^{-1}$ to 280 °C, hold 0 min.

*Method B:* for the amination of (*R*)-3-methylcyclohexanone with cyclopropylamine: constant pressure 6.9 psi; temperature program: 60 °C, hold 6.5 min; 20 °C $min^{-1}$ to 100 °C, hold 1 min; 20 °C $min^{-1}$ to 280 °C, hold 1 min.

Internal standard toluene: method A: 2.8 min, method B: 3.6 min; cosolvent DMSO: method A: 7.1 min, method B: 9.5 min.

#### 4.4 GC-FID chromatograms for the biocatalytic reactions

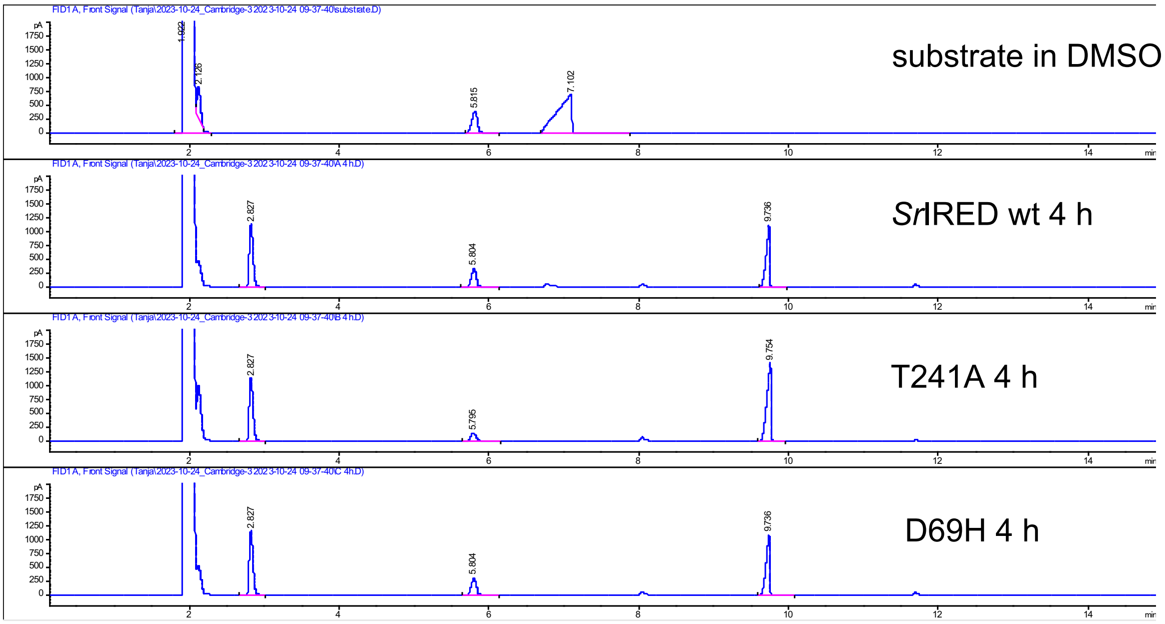

#### 5 Stereoselectivity of *Sr*IRED wt and selected mutants with (*R*)-3-methylcyclohexanone (Figure S16)

##### 5.1 Reaction conditions

The reaction conditions described in **Supplementary Methods 4.1** were used. IRED5 and IRED20 were used as reference enzymes, as described in [61].

##### 5.2 Preparation of calibration curves for GC-FID

**Substrate (*R*)-3-methylcyclohexanone**

| | substrate (mM) | area IS | area substrate | area substrate/area IS |
|---|---|---|---|---|
| calA | 25 | 1930 | 2521 | 1.31 |
| calB | 12.5 | 1991 | 1219 | 0.61 |
| calC | 6 | 2022 | 583 | 0.29 |
| calD | 2.5 | 2002 | 244 | 0.12 |
| calE | 1 | 2017 | 109 | 0.05 |

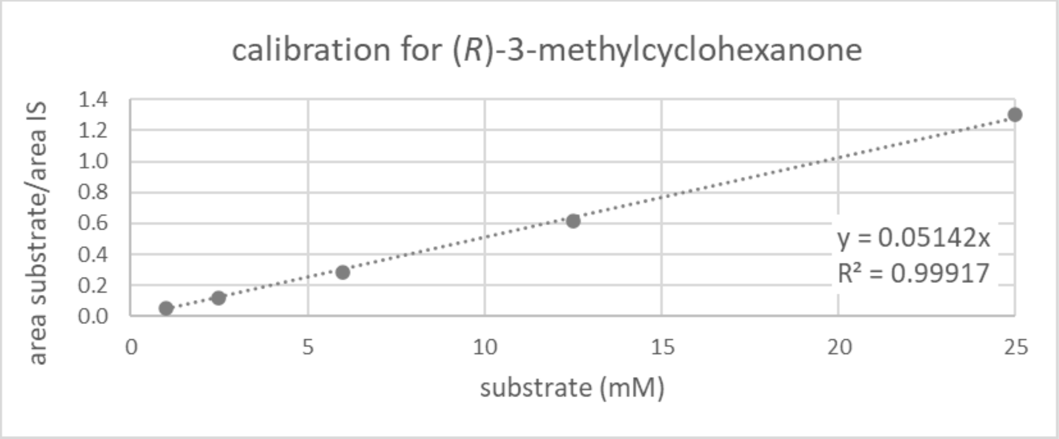

#### 5.3 GC-FID chromatograms for the biocatalytic reactions

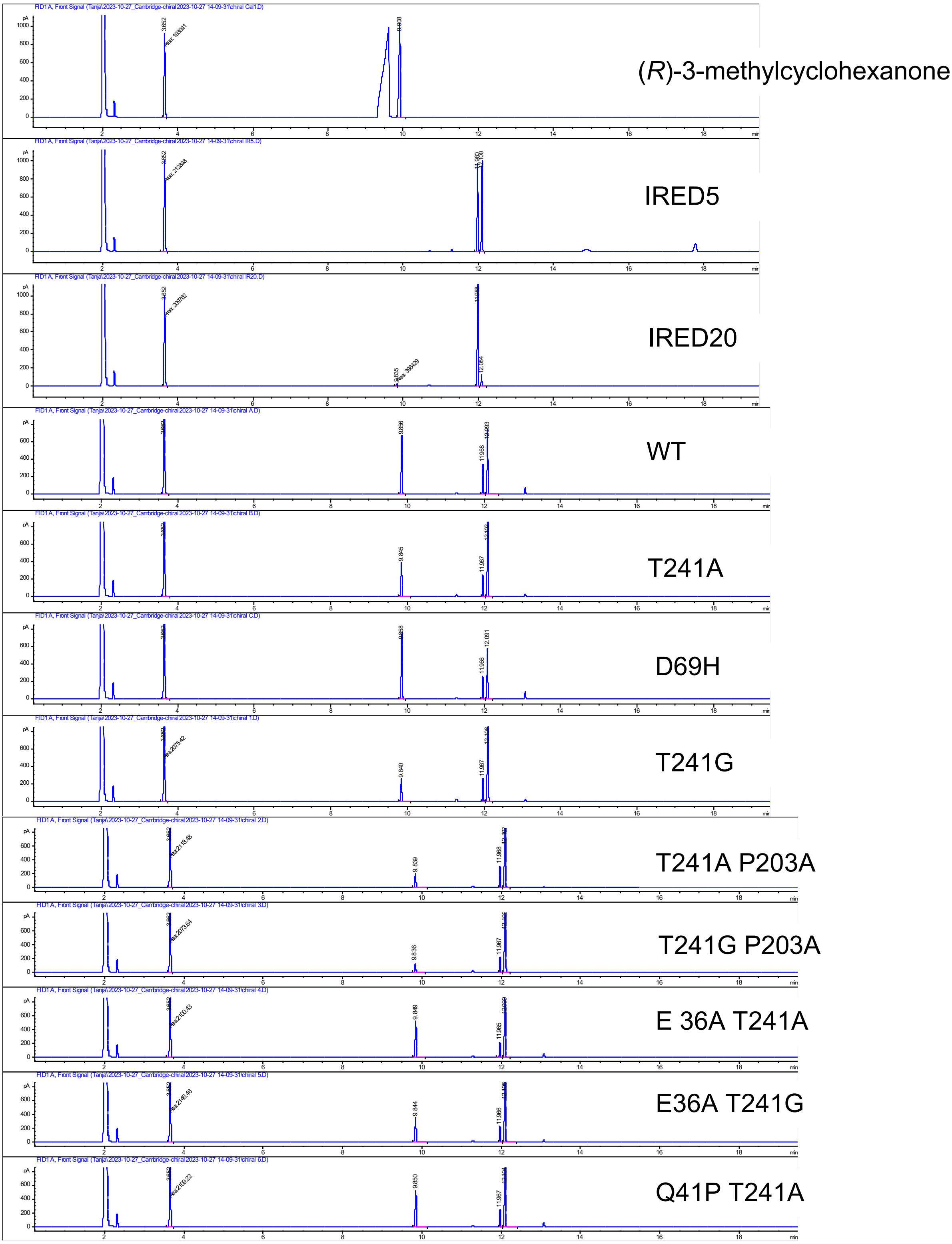

#### 5.4 GC-MS spectrum for the reaction of D69H with (R)-3-methylcyclohexanone and cyclopropylamine

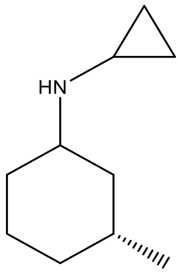

m/z: 153.2 (100.0%), 154.2 (11.0%)

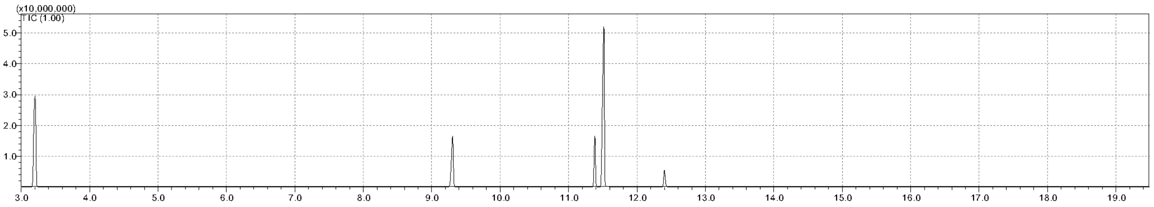

Peak at 11.4 min amine product

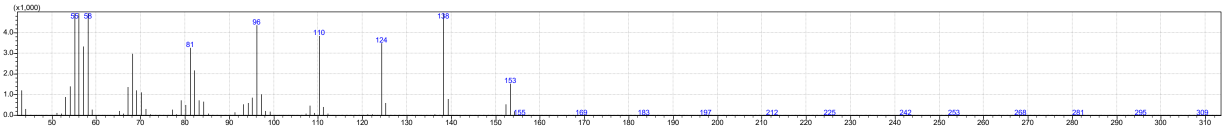

Peak at 11.5 min amine product

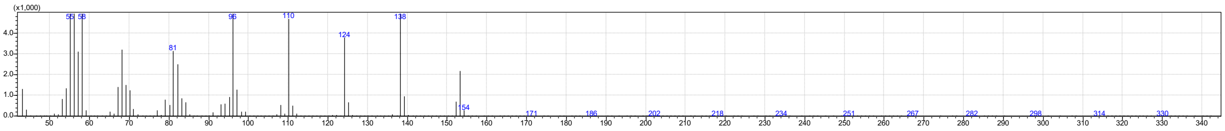

### 6 Biotransformations with *Pc*IRED and its mutants

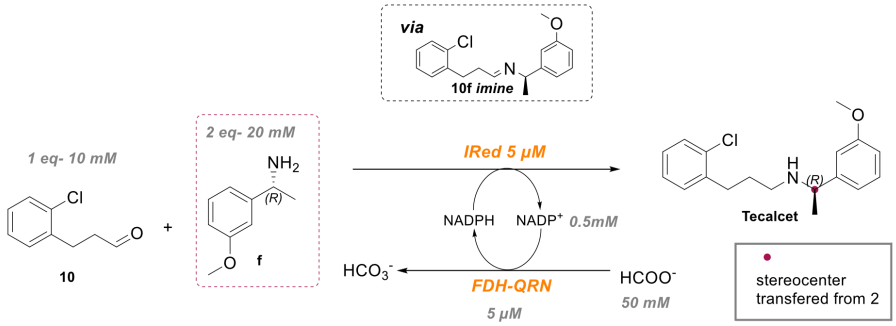

#### 6.1 PcIRED: Biocatalytic reactions

Biocatalytic reactions using enantiopure amine (*R*-2a) were performed on a 0.5 mL scale in 2 mL Eppendorf tubes containing the following components: buffer (KPi, 100 mM, pH 8) supplemented with 50 mM sodium formate, $NADP^+$ (0.5 mM), FDH-QRN (5 µM), and IRED (5 µM). The aldehyde substrate (10 mM) and amine substrate (20 mM) were added last. Specifically, 100 µL of the aldehyde substrate (prepared as a 50 mM stock solution in DMSO) and 150 µL of the amine substrate (prepared as a 66.6 mM stock solution in DMSO) were added, resulting in a final DMSO concentration of 50% v/v.
Biocatalytic reactions using racemic amine 2 were conducted over 24 hours with the following adjustments: 10 µM FDH-QRN, 20 µM IRED, 5 mM aldehyde substrate, and 10 mM amine substrate.
The reactions were incubated in a horizontal shaker at 30 °C. After incubation, the reaction mixtures were basified with 50 µL of 10 M KOH. Organic compounds were then extracted using DCM containing an internal standard (toluene, 10 mM) with two 500 µL extractions. The organic phase was dried over $MgSO_4$ and analyzed using GC-FID or GC-MS.

#### 6.2 PcIRED: Analytical methods

##### 6.2.1 GC-FID columns and methods

Column: the conversions were measured by GC-FID using an Agilent J&W HP-5 (30 m, 320 µm, 0.25 µm) column. Carrier gas: $H_2$
Method: pressure 5.1 psi; constant flow 1.8 mL/ min temperature program: 80 °C, hold 5 min; 10 °C $min^{-1}$ to 160 °C, hold 0 min; 20 °C $min^{-1}$ to 300 °C, hold 5 min.
Column: The ee values were measured by GC-FID using a Hydrodex-β-TBDAC column from Macherey-Nagel (L: 50m, OD: 0.40 mm, ID: 0.25 mm). Carrier gas: $H_2$
Method: for the determination of the *ee* of 3: constant pressure 11.7 psi; temperature program: 100 °C, hold 2 min; 1 °C $min^{-1}$ to 220 °C, hold 2 min.

##### 6.2.2 GC-MS columns and methods

Column The products of enzymatic reactions were identified using a QP2010SE GCMS system (Shimadzu), with He as carrier gas using an HP-5 column from Agilent (30 m, 250 µm, 0.25 µm).
Method: GC program parameters: Linear Velocity: 45.1 cm/sec, pressure 18.4 KPa, Flow: 1.53 mL/min, split ratio 20:1, T injector 250 °C. Temperature Program: T initial 60 °C, hold 10 min, gradient 10 °C/min up to 300 °C; hold 1 min. MS program, parameters: Ion Source Temperature: 200 °C, Detector Voltage: 0.1 Kv, Start Time: 3 min, End Time: 35 min, Start m/z: 43, End m/z: 600.

#### 6.3 PcIRED: Spectra

### GC-MS

Overlap of enzymatic reaction with wt camIred (black) and negative control (pink) without enzyme

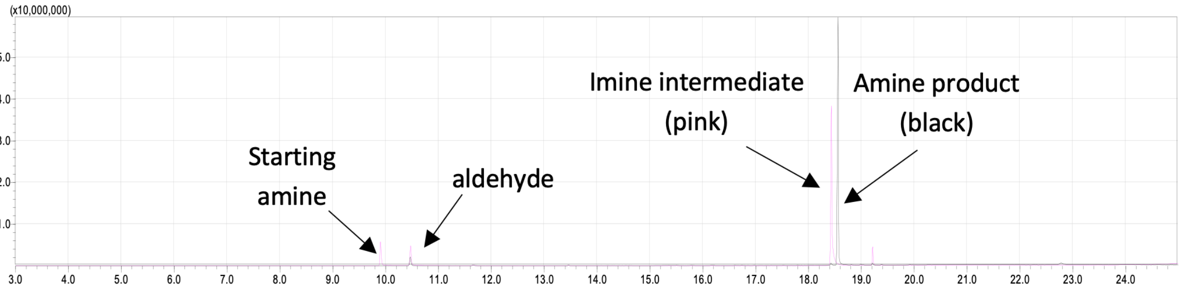

Mass fragmentation at peak 18.557 (**amine product**) from Wt PcIRED

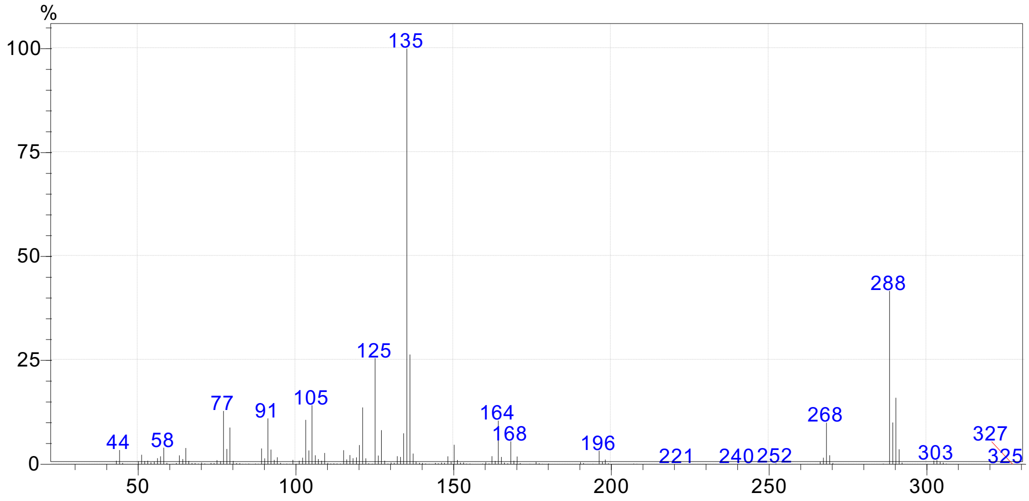

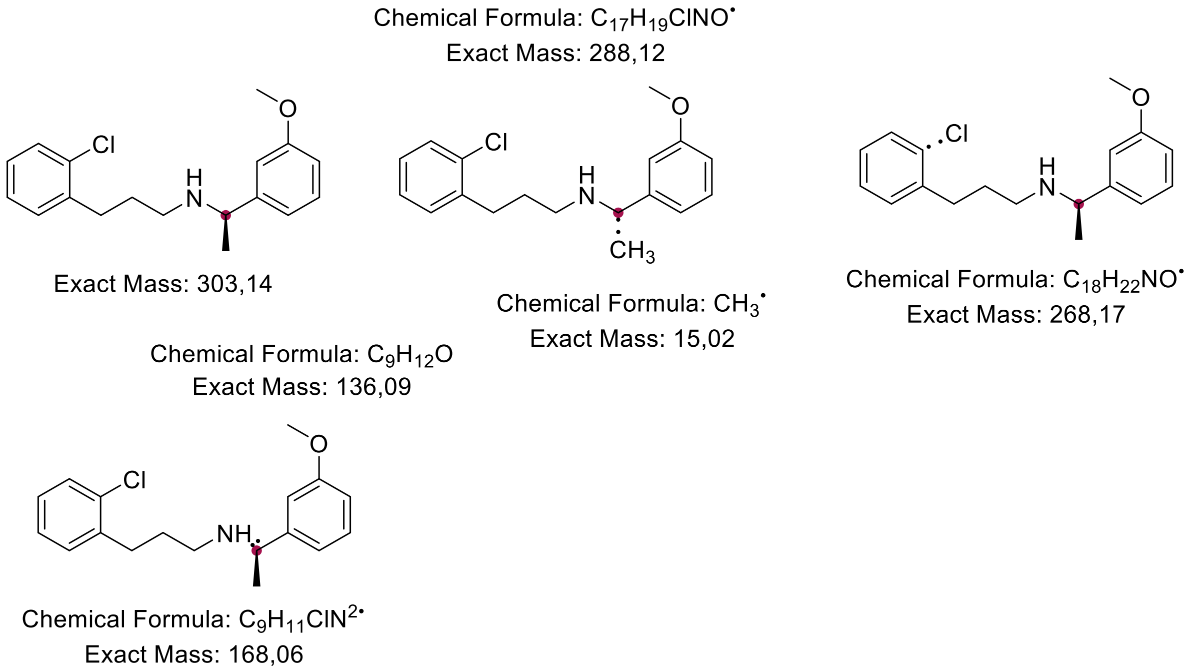

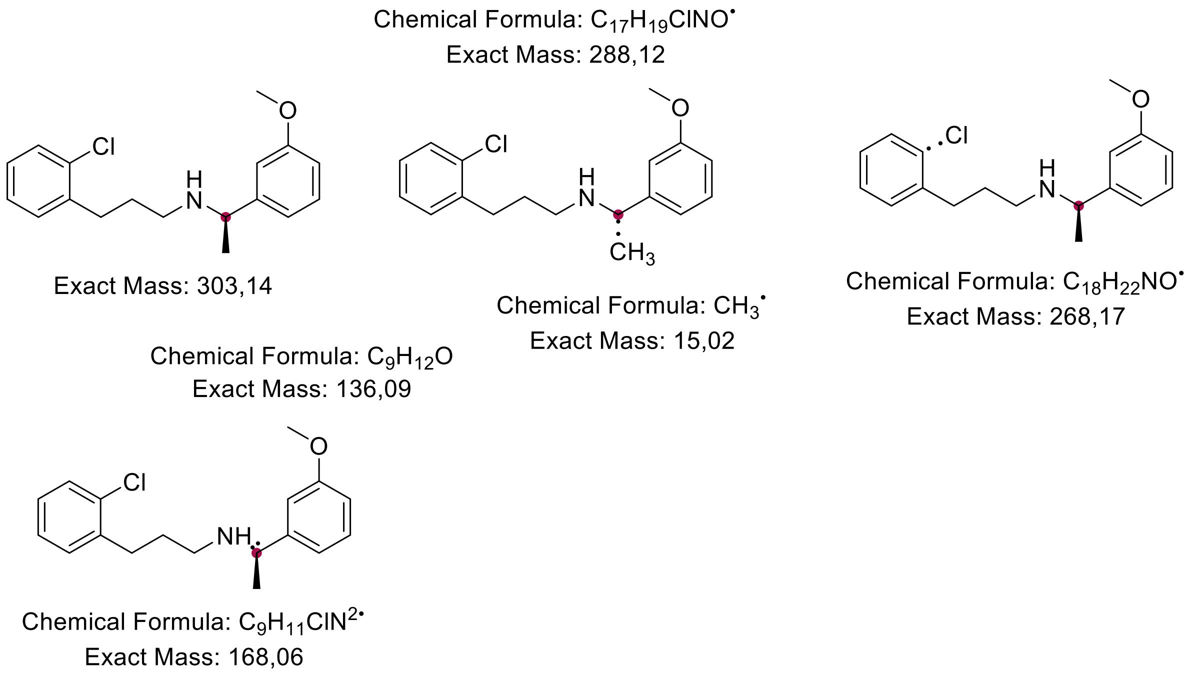

Mass fragmentation at peak 18.427 () from NC2

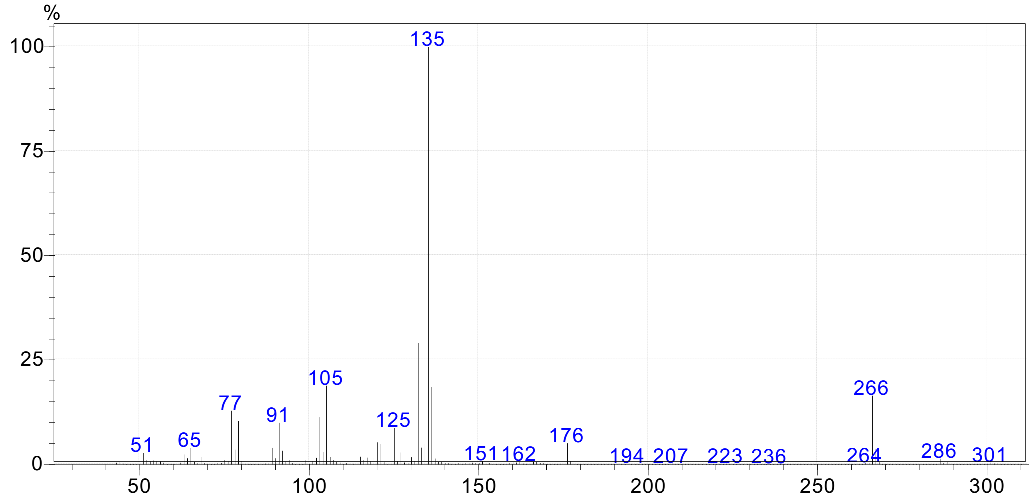
%
100
75
50
25
0
135
105
77
91
125
51
65
151
162
176
194
207
223
236
266
264
286
301
50
100
150
200
250
300

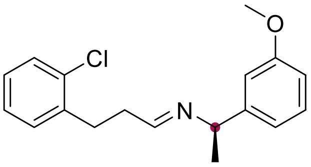

Exact Mass: 301,12

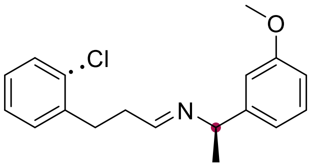

Chemical Formula: $C_{18}H_{20}NO^{\bullet}$
Exact Mass: 266,15

Chemical Formula: $C_{17}H_{17}ClNO^{\bullet}$
Exact Mass: 286,10

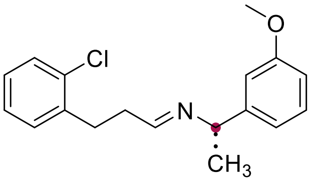

Chemical Formula: $CH_3^{\bullet}$
Exact Mass: 15,02

Chemical Formula: $C_9H_{12}O$
Exact Mass: 136,09

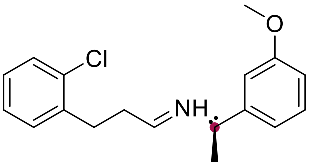

Chemical Formula: $C_9H_{10}ClN^{2\bullet}$
Exact Mass: 167,05

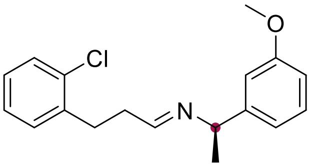

Exact Mass: 301,12

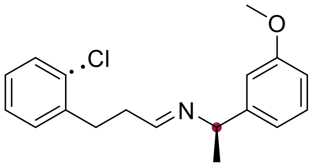

Chemical Formula: $C_{18}H_{20}NO^{\bullet}$
Exact Mass: 266,15

Chemical Formula: $C_{17}H_{17}ClNO^{\bullet}$
Exact Mass: 286,10

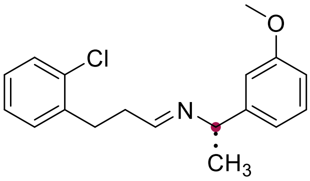

Chemical Formula: $CH_3^{\bullet}$
Exact Mass: 15,02

Chemical Formula: $C_9H_{12}O$
Exact Mass: 136,09

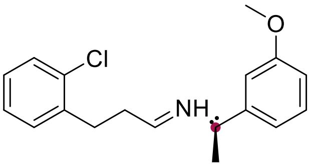

Chemical Formula: $C_9H_{10}ClN^{2\bullet}$
Exact Mass: 167,05

#### GC-FID for determination of the relative yield (with wild type and R61L A181T)

Aldehyde reference

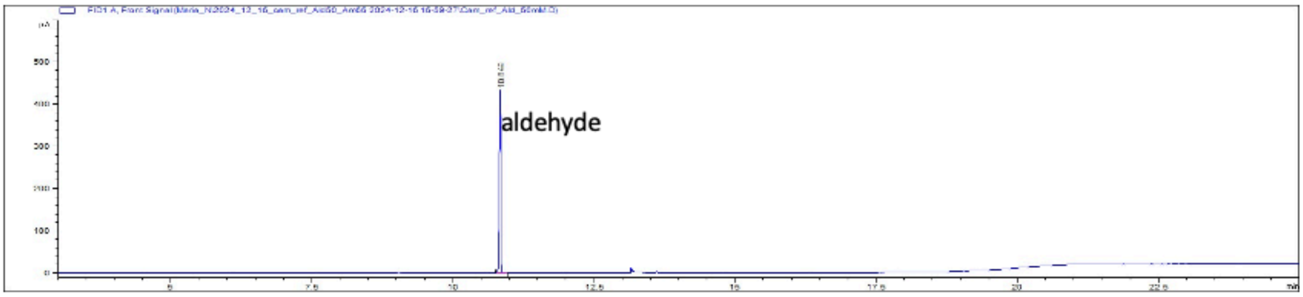

Amine reference

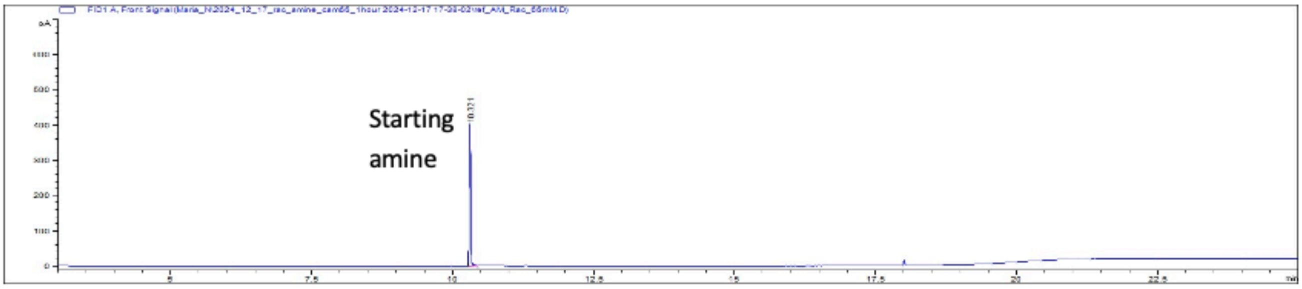

Reaction without enzyme

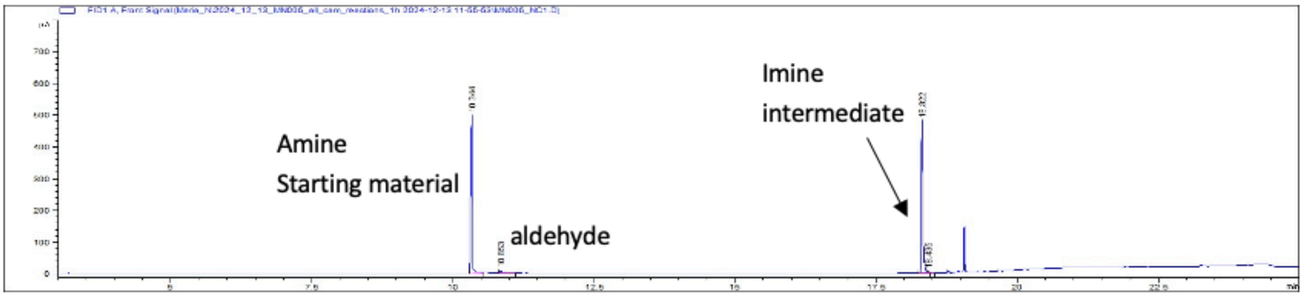

Reaction with cam 66

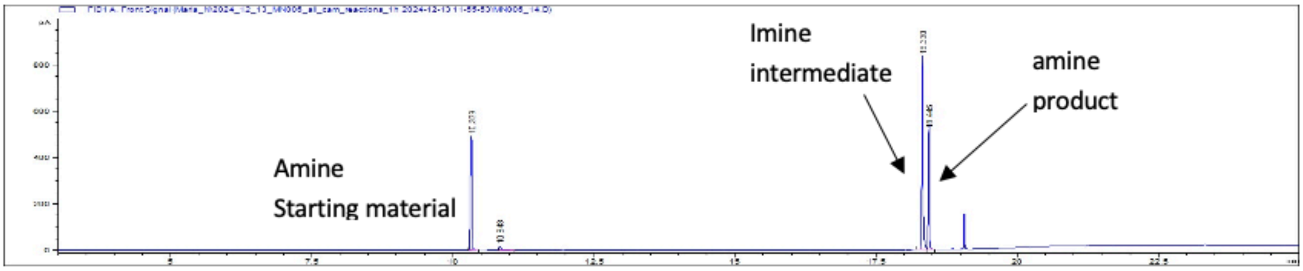

Reaction with Wt

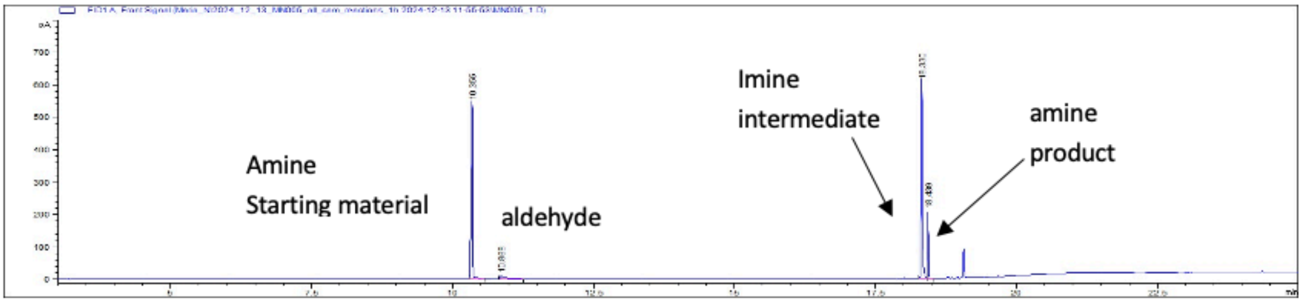

#### 7 Primers for cloning epPCR libraries for lrDMS with *Srl*RED and *Pc*IRED

##### 7.1 *Srl*RED

The following fragments were generated by PCR (Q5 polymerase, standard conditions) for assembly of the library: Promoter with primers G1 and G5 and vector with primers G2 and G6. Primers G3 + G4 were used for error-prone PCR.

Primers for Gibson assembly of SrIRED epPCR library

| | |
|---|---|
| G1 | atcccgcgaaattaatacgactcactatatgcgactcctgcattaggaaattaatacGACTCACTATAGGGG |
| G2 | agatctcgatcctctacgccggacgcatctaagggagagcgtcgagatcc |
| G3 | CCGCGCGGCAGCCATATG |
| G4 | caaatttcgcaGCAGCGGTTTCTTTACCAGATTA |
| G5 | CATATGGCTGCCGCGCGGCACCAG |
| G6 | TAATCTGGTAAAGAAACCGCTGCtgc |

The UMI sequence printed below was ordered as oligonucleotide from IDT. Double-stranded DNA was produced by 3 cycles of PCR with the primer printed below.

**UMI sequence:**
gatgcgtccggcgtagaggatcgagatctgNNNNNNNNNNGATCNNNNNNNNNNGATCNNNNNNNN
NNGATCNNNNNNNNNNGATCNNNNNNNNNNgatcccgcgaaattaatacgactcactata

**Primer:**
tatagtgagtcgtattaatttcgcgg

##### 7.2 *Pc*IRED

Vector (GGA1 & GGA2) and promoter fragment were amplified using standard PCR with Q5 polymerase
Promoter: GGA3 & GGA4
Gene: GGA5 & GGA6

Primers for Golden Gate assembly of SrIRED epPCR library

| | |
|---|---|
| GGA1 | ctatGCTCTTCattgaacgccagcacatggac |
| GGA2 | cgttGCTCTTCtagctGCTCGGAGATTGAGAAAGCAgcatctaagggagagcgtcgag |
| GGA3 | attgGCTCTTCgatatgcgactcctgcattaggaaat |
| GGA4 | gataGCTCTTCaGGAACCAGGCCGCTGCTG |
| GGA5 | ctagGCTCTTCaTCCGCGCGGCAGCCATATG |
| GGA6 | cggtGCTCTTCacaaatttcgcaGCAGCGGTTTCTTTACCAGATTA |

**UMI sequence:**
cactGCTCTTCagctgNNNNRYNNNNGATCNNNNRYNNNNGATCNNNNRYNNNNGATCNN
NNRYNNNNGATCNNNNRYNNNNgatcacgcgtcatgcatgcgactcactataaGAAGAGCacag

**Primer:**
ctgtGCTCTTCttatagtgagtcgcatgcatgacgcgtgatc

### 8 Prepraration of DNA for Nanopore sequencing

#### 8.1 SrIRED

For dictionary generation with the SrIRED epPCR library, the SIRED gene together with the UMI region were cleaved out using the enzymes AcuI and Bsu36I (NEB). The fragement was separated from the vectore by agarose gel electrophoresis.

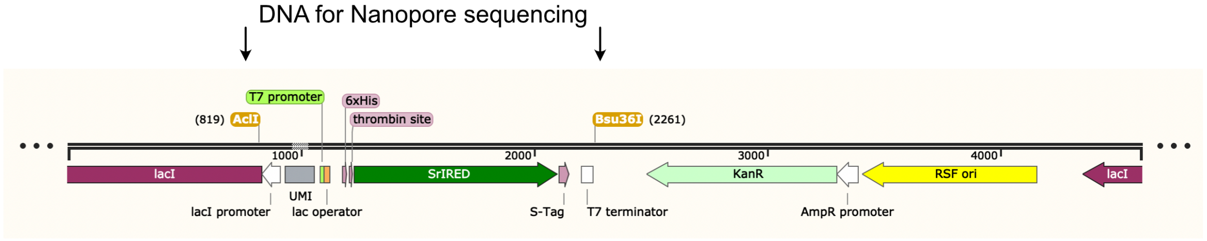

#### 8.2 *Pc*IRED

For dictionary generation with PcIRED, the whole plasmid was read. We linearised the plasmid using the restriction enzyme HpaI (NEB) and purified the DNA using SPRI beads (Beckman).

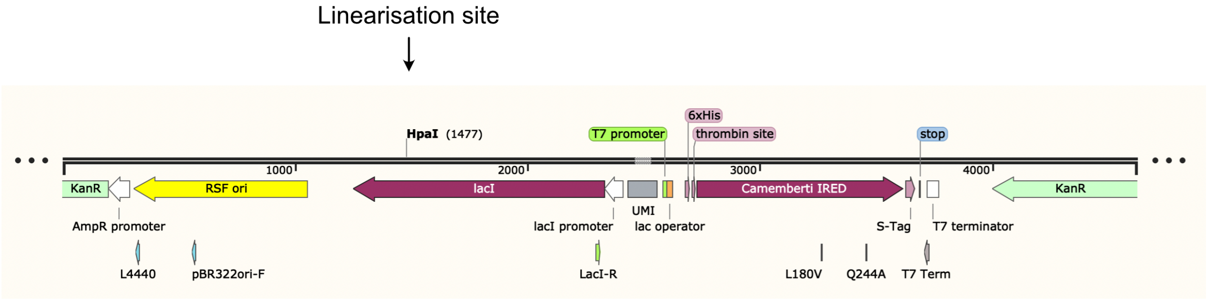

#### 9 Example curves for melting temperature determination via thermal shift assay

Below, example curves in 3 technical replicates of thermal shift assays (Methods) with BtIRED wild type (left) and T241A (right) are shown. Data was analysed using paulsbond.co.uk.

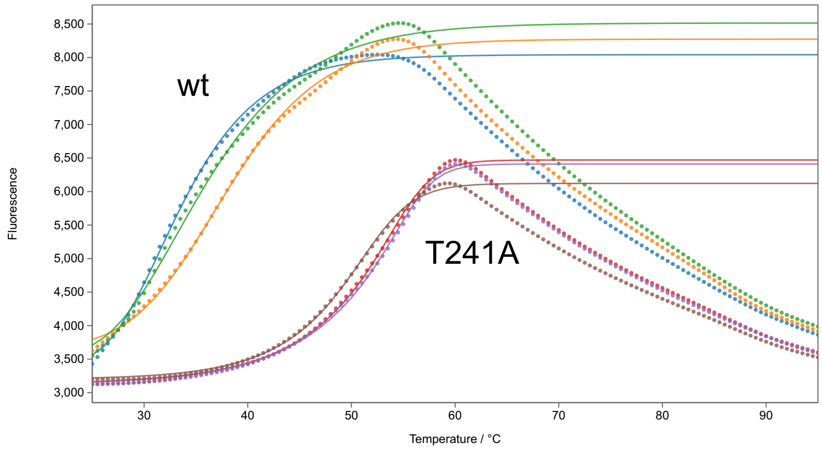

#### Sequences

##### SrIRED nucleotide sequence

ATGGGCAGCAGCCATCATCATCATCATCACAGCAGCGGCCTGGTGCCGCGCGGCAGC
CATATGCGGGACACCGATGTCACTGTCCTCGGCCTGGGTCTGATGGGCCAGGCTTTGG
CAGGCGCCTTTCTCAAAGATGGGCATGCGACGACCGTTTGGAATCGTTCTGAAGGCAA
AGCCGGTCAGCTGGCGGAACAGGGTGCGGTACTGGCCAGCAGCGCCCGGGATGCGG
CCGAGGCGAGCCCACTGGTAGTTGTTTGCGTTTCAGACCACGCCGCGGTTCGCGCGG
TGCTGGATCCTCTGGGTGATGTTCTGGCGGGGCGTGTTCTGGTGAACCTCACCTCTGG
AACTAGCGAACAGGCCCGCGCCACGGCGGAGTGGGCCGCTGAACGCGGGATTACATA
CCTTGACGGAGCTATCATGGCAATCCCGCAGGTTGTCGGCACAGCAGATGCATTCTTA
CTGTACTCTGGGCCGGAAGCCGCTTATGAGGCGCACGAACCGACCTTACGGAGCCTG
GGGGCGGGTACCACTTATTTGGGCGCCGACCACGGCCTGTCATCTTTGTATGACGTCG
CGTTGCTCGGCATCATGTGGGGCACGCTGAATTCTTTCCTCCACGGGGCGGCATTGTT
GGGCACAGCCAAAGTGGAAGCGACGACCTTTGCACCGTTCGCGAACCGTTGGATTGAA
GCGGTAACCGGGTTTGTATCCGCGTATGCTGGTCAGGTGGACCAGGGGGCCTACCCG
GCACTGGACGCGACTATCGATACCCACGTGGCTACGGTAGATCATCTGATTCACGAGT
CCGAGGCCGCTGGCGTGAACACGGAGCTCCCACGCCTCGTGCGCACGCTGGCCGAC
CGTGCCCTGGCGGGAGGCCAGGGCGGTCTTGGTTACGCCGCTATGATTGAGCAGTTT
CGGTCTCCTAGCTAA

##### SrIRED amino acid sequence

MGSSHHHHHHSSGLVPRGSHMRDTDVTVLGLGLMGQALAGAFLKDGHATTVWNRSEGK
AGQLAEQGAVLASSARDAAEASPLVVVCVSDHAAVRAVLDPLGDVLAGRVLVNLTSGTSE
QARATAEWAAERGITYLDGAIMAIPQVVGTADAFLLYSGPEAAYEAHEPTLRSLGAGTTYLG
ADHGLSSLYDVALLGIMWGTLNSFLHGAALLGTAKVEATTFAPFANRWIEAVTGFVSAYAG
QVDQGAYPALDATIDTHVATVDHLIHESEAAGVNTELPRLVRTLADRALAGGQGGLGYAAM
IEQFRSPS

##### SrIRED reference for numbering in data analysis

MRDTDVTVLGLGLMGQALAGAFLKDGHATTVWNRSEGKAGQLAEQGAVLASSARDAAEA
SPLVVVCVSDHAAVRAVLDPLGDVLAGRVLVNLTSGTSEQARATAEWAAERGITYLDGAIM
AIPQVVGTADAFLLYSGPEAAYEAHEPTLRSLGAGTTYLGADHGLSSLYDVALLGIMWGTLN
SFLHGAALLGTAKVEATTFAPFANRWIEAVTGFVSAYAGQVDQGAYPALDATIDTHVATVD
HLIHESEAAGVNTELPRLVRTLADRALAGGQGGLGYAAMIEQFRSPS*

##### SrIRED whole plasmid sequence

ctgtaacatcattggcaacgctacctttgccatgtttcagaaacaactctggcgcatcgggcttcccatacaatcgatagattgtcgc
acctgattgcccgacattatcgcgagcccatttatacccatataaatcagcatccatgttggaatttaatcgcggcctagagcaaga
cgtttcccgttgaatatggctcatactcttcctttttcaatattattgaagcatttatcagggttattgtctcatgagcggatacatatttgaat
gtatttagaaaaataaacaaataggcatgcagcgctcttccgcttcctcgctcactgactcgctacgctcggtcgttcgactgcggc
gagcggtgtcagctcactcaaaagcggtaatacggttatccacagaatcaggggataaagccggaaagaacatgtgagcaa
aaagcaaagcaccggaagaagccaacgccgcaggcgtttttccataggctccgcccccctgacgagcatcacaaaaatcga
cgctcaagccagaggtggcgaaacccgacaggactataaagataccaggcgtttccccctggaagctccctcgtgcgctctcct
gttccgaccctgccgcttaccggatacctgtccgcctttctcccttcgggaagcgtggcgctttctcatagctcacgctgttggtatctc
agttcggtgtaggtcgttcgctccaagctgggctgtgtgcacgaaccccccgttcagcccgaccgctgcgccttatccggtaactat
cgtcttgagtccaacccggtaagacacgacttatcgccactggcagcagccattggtaactgatttagaggactttgtcttgaagtt
atgcacctgttaaggctaaactgaaagaacagattttggtgagtgcggtcctccaacccacttaccttggttcaaagagttggtagc
tcagcgaaccttgagaaaaccaccgttggtagcggtggtttttctttatttatgagatgatgaatcaatcggtctatcaagtcaacga
acagctattccgttactctagatttcagtgcaatttatctcttcaaatgtagcacctgaagtcagccccatacgatataagttgtaattct

catgttagtcatgccccgcgcccaccggaaggagctgactgggttgaaggctctcaagggcatcggtcgagatcccggtgccta
atgagtgagctaacttacattaattgcgttgcgctcactgcccgctttccagtcgggaaacctgtcgtgccagctgcattaatgaatc
ggccaacgcgcggggagaggcggtttgcgtattgggcgccagggtggtttttcttttcaccagtgagacgggcaacagctgattg
cccttcaccgcctggccctgagagagttgcagcaagcggtccacgctggtttgccccagcaggcgaaaatcctgtttgatggtgg
ttaacggcgggatataacatgagctgtcttcggtatcgtcgtatcccactaccgagatgtccgcaccaacgcgcagcccggactc
ggtaatggcgcgcattgcgcccagcgccatctgatcgttggcaaccagcatcgcagtgggaacgatgccctcattcagcatttgc
atggtttgttgaaaaccggacatggcactccagtcgccttcccgttccgctatcggctgaatttgattgcgagtgagatatttatgcca
gccagccagacgcagacgcgccgagacagaacttaatgggcccgctaacagcgcgatttgctggtgacccaatgcgaccag
atgctccacgcccagtcgcgtaccgtcttcatgggagaaaataatactgttgatgggtgtctggtcagagacatcaagaaataac
gccggaacattagtgcaggcagcttccacagcaatggcatcctggtcatccagcggatagttaatgatcagcccactgacgcgtt
gcgcgagaagattgtgcaccgccgctttacaggcttcgacgccgcttcgttctaccatcgacaccaccacgctggcacccagttg
atcggcgcgagatttaatcgccgcgacaatttgcgacggcgcgtgcagggccagactggaggtggcaacgccaatcagcaac
gactgtttgcccgccagttgttgtgccacgcggttgggaatgtaattcagctccgccatcgccgcttccactttttcccgcgttttcgca
gaaacgtggctggcctggttcaccacgcgggaaacggtctgataagagacaccggcatactctgcgacatcgtataacgttact
ggtttcacattcaccaccctgaattgactctcttccgggcgctatcatgccataccgcgaaaggttttgcgccattcgatggtgtccgg
gatctcgacgctctcccttatgcgactcctgcattaggaaattaatacgactcactataggggaattgtgagcggataacaattcC
CCTGTAGAAATAATTTTGTTTAACTTTAATAAGGAGATATACCATGGGCAGCAGCCATCA
TCATCATCATCACAGCAGCGGCCTGGTGCCGCGCGGCAGCCATATGCGGGACACCGA
TGTCACTGTCCTCGGCCTGGGTCTGATGGGCCAGGCTTTGGCAGGCGCCTTTCTCAAA
GATGGGCATGCGACGACCGTTTGGAATCGTTCTGAAGGCAAAGCCGGTCAGCTGGCG
GAACAGGGTGCGGTACTGGCCAGCAGCGCCCGGGATGCGGCCGAGGCGAGCCCACT
GGTAGTTGTTTGCGTTTCAGACCACGCCGCGGTTCGCGCGGTGCTGGATCCTCTGGGT
GATGTTCTGGCGGGGCGTGTTCTGGTGAACCTCACCTCTGGAACTAGCGAACAGGCCC
GCGCCACGGCGGAGTGGGCCGCTGAACGCGGGATTACATACCTTGACGGAGCTATCA
TGGCAATCCCGCAGGTTGTCGGCACAGCAGATGCATTCTTACTGTACTCTGGGCCGGA
AGCCGCTTATGAGGCGCACGAACCGACCTTACGGAGCCTGGGGGCGGGTACCACTTA
TTTGGGCGCCGACCACGGCCTGTCATCTTTGTATGACGTCGCGTTGCTCGGCATCATG
TGGGGCACGCTGAATTCTTTCCTCCACGGGGCGGCATTGTTGGGCACAGCCAAAGTGG
AAGCGACGACCTTTGCACCGTTCGCGAACCGTTGGATTGAAGCGGTAACCGGGTTTGT
ATCCGCGTATGCTGGTCAGGTGGACCAGGGGGCCTACCCGGCACTGGACGCGACTAT
CGATACCCACGTGGCTACGGTAGATCATCTGATTCACGAGTCCGAGGCCGCTGGCGTG
AACACGGAGCTCCCACGCCTCGTGCGCACGCTGGCCGACCGTGCCCTGGCGGGAGG
CCAGGGCGGTCTTGGTTACGCCGCTATGATTGAGCAGTTTCGGTCTCCTAGCTAATCT
GGTAAAGAAACCGCTGCtgcgaaatttgaacgccagcacatggactcgtctactagcgcagcttaattaacctaggct
gctgccaccgctgagcaataactagcataaccccttggggcctctaaacgggtcttgaggggttttttgctgaaacctcaggcattt
gagaagcacacggtcacactgcttccggtagtcaataaaccggtaaaccagcaatagacataagcggctatttaacgaccctg
ccctgaaccgacgacaagctgacgaccgggtctccgcaagtggcacttttcggggaaatgtgcgcggaacccctatttgtttatttt
tctaaatacattcaaatatgtatccgctcatgaattaattcttagaaaaactcatcgagcatcaaatgaaactgcaatttattcatatc
aggattatcaataccatatttttgaaaaagccgtttctgtaatgaaggagaaaactcaccgaggcagttccataggatggcaagat
cctggtatcggtctgcgattccgactcgtccaacatcaatacaacctattaatttcccctcgtcaaaaataaggttatcaagtgagaa
atcaccatgagtgacgactgaatccggtgagaatggcaaaagtttatgcatttctttccagacttgttcaacaggccagccattacg
ctcgtcatcaaaatcactcgcatcaaccaaaccgttattcattcgtgattgcgcctgagcgagacgaaatacgcggtcgctgttaa
aaggacaattacaaacaggaatcgaatgcaaccggcgcaggaacactgccagcgcatcaacaatattttcacctgaatcagg
atattcttctaatacctggaatgctgttttcccggggatcgcagtggtgagtaaccatgcatcatcaggagtacggataaaatgcttg
atggtcggaagaggcataaattccgtcagccagtttagtctgaccatctcat

**SrIRED whole plasmid sequence with UMI**

gtcttcggtatcgtcgtatcccactaccgagatgtccgcaccaacgcgcagcccggactcggtaatggcgcgcattgcgcccag
cgccatctgatcgttggcaaccagcatcgcagtgggaacgatgccctcattcagcatttgcatggtttgttgaaaaccggacatgg
cactccagtcgccttcccgttccgctatcggctgaatttgattgcgagtgagatatttatgccagccagccagacgcagacgcgcc
gagacagaacttaatgggcccgctaacagcgcgatttgctggtgacccaatgcgaccagatgctccacgcccagtcgcgtacc
gtcttcatgggagaaaataatactgttgatgggtgtctggtcagagacatcaagaaataacgccggaacattagtgcaggcagct
tccacagcaatggcatcctggtcatccagcggatagttaatgatcagcccactgacgcgttgcgcgagaagattgtgcaccgcc
gctttacaggcttcgacgccgcttcgttctaccatcgacaccaccacgctggcacccagttgatcggcgcgagatttaatcgccgc
gacaatttgcgacggcgcgtgcagggccagactggaggtggcaacgccaatcagcaacgactgtttgcccgccagttgttgtgc

cacgcggttgggaatgtaattcagctccgccatcgccgcttccactttttcccgcgttttcgcagaaacgtggctggcctggttcacc
acgcgggaaacggtctgataagagacaccggcatactctgcgacatcgtataacgttactggtttcacattcaccaccctgaattg
actctcttccgggcgctatcatgccataccgcgaaaggttttgcgccattcgatggtgtccgggatctcgacgctctcccttagatgc
gtccggcgtagaggatcgagatctgNNNNNNNNNNGATCNNNNNNNNNNGATCNNNNNNNNNNGA
TCNNNNNNNNNNGATCNNNNNNNNNNgatcccgcgaaattaatacgactcactatatgcgactcctgcattag
gaaattaatacGACTCACTATAGGGGaattgtgagcggataacaattcCCCTGTAGAAATAATTTTGTTT
AACTTTAATAAGGAGATATACCATGGGCAGCAGCCATCATCATCATCATCACAGCAGCG
GCCTGGTGCCGCGCGGCAGCCATATGCGGGACACCGATGTCACTGTCCTCGGCCTGG
GTCTGATGGGCCAGGCTTTGGCAGGCGCCTTTCTCAAAGATGGGCATGCGACGACCGT
TTGGAATCGTTCTGAAGGCAAAGCCGGTCAGCTGGCGGAACAGGGTGCGGTACTGGC
CAGCAGCGCCCGGGATGCGGCCGAGGCGAGCCCACTGGTAGTTGTTTGCGTTTCAGA
CCACGCCGCGGTTCGCGCGGTGCTGGATCCTCTGGGTGATGTTCTGGCGGGGCGTGT
TCTGGTGAACCTCACCTCTGGAACTAGCGAACAGGCCCGCGCCACGGCGGAGTGGGC
CGCTGAACGCGGGATTACATACCTTGACGGAGCTATCATGGCAATCCCGCAGGTTGTC
GGCACAGCAGATGCATTCTTACTGTACTCTGGGCCGGAAGCCGCTTATGAGGCGCACG
AACCGACCTTACGGAGCCTGGGGGCGGGTACCACTTATTTGGGCGCCGACCACGGCC
TGTCATCTTTGTATGACGTCGCGTTGCTCGGCATCATGTGGGGCACGCTGAATTCTTTC
CTCCACGGGGCGGCATTGTTGGGCACAGCCAAAGTGGAAGCGACGACCTTTGCACCG
TTCGCGAACCGTTGGATTGAAGCGGTAACCGGGTTTGTATCCGCGTATGCTGGTCAGG
TGGACCAGGGGGCCTACCCGGCACTGGACGCGACTATCGATACCCACGTGGCTACGG
TAGATCATCTGATTCACGAGTCCGAGGCCGCTGGCGTGAACACGGAGCTCCCACGCCT
CGTGCGCACGCTGGCCGACCGTGCCCTGGCGGGAGGCCAGGGCGGTCTTGGTTACG
CCGCTATGATTGAGCAGTTTCGGTCTCCTAGCTAATCTGGTAAAGAAACCGCTGCtgcga
aatttgaacgccagcacatggactcgtctactagcgcagcttaattaacctaggctgctgccaccgctgagcaataactagcata
accccttggggcctctaaacgggtcttgaggggttttttgctgaaacctcaggcatttgagaagcacacggtcacactgcttccggt
agtcaataaaccggtaaaccagcaatagacataagcggctatttaacgaccctgccctgaaccgacgacaagctgacgaccg
ggtctccgcaagtggcacttttcggggaaatgtgcgcggaacccctatttgtttatttttctaaatacattcaaatatgtatccgctcatg
aattaattcttagaaaaactcatcgagcatcaaatgaaactgcaatttattcatatcaggattatcaataccatatttttgaaaaagcc
gtttctgtaatgaaggagaaaactcaccgaggcagttccataggatggcaagatcctggtatcggtctgcgattccgactcgtcca
acatcaatacaacctattaatttcccctcgtcaaaaataaggttatcaagtgagaaatcaccatgagtgacgactgaatccggtga
gaatggcaaaagtttatgcatttctttccagacttgttcaacaggccagccattacgctcgtcatcaaaatcactcgcatcaaccaa
accgttattcattcgtgattgcgcctgagcgagacgaaatacgcggtcgctgttaaaaggacaattacaaacaggaatcgaatgc
aaccggcgcaggaacactgccagcgcatcaacaatattttcacctgaatcaggatattcttctaatacctggaatgctgttttcccg
gggatcgcagtggtgagtaaccatgcatcatcaggagtacggataaaatgcttgatggtcggaagaggcataaattccgtcagc
cagtttagtctgaccatctcatctgtaacatcattggcaacgctacctttgccatgtttcagaaacaactctggcgcatcgggcttccc
atacaatcgatagattgtcgcacctgattgcccgacattatcgcgagcccatttatacccatataaatcagcatccatgttggaattta
atcgcggcctagagcaagacgtttcccgttgaatatggctcatactcttcctttttcaatattattgaagcatttatcagggttattgtctc
atgagcggatacatatttgaatgtatttagaaaaataaacaaataggcatgcagcgctcttccgcttcctcgctcactgactcgcta
cgctcggtcgttcgactgcggcgagcggtgtcagctcactcaaaagcggtaatacggttatccacagaatcaggggataaagc
cggaaagaacatgtgagcaaaaagcaaagcaccggaagaagccaacgccgcaggcgtttttccataggctccgcccccctg
acgagcatcacaaaaatcgacgctcaagccagaggtggcgaaacccgacaggactataaagataccaggcgtttccccctg
gaagctccctcgtgcgctctcctgttccgaccctgccgcttaccggatacctgtccgcctttctcccttcgggaagcgtggcgctttct
catagctcacgctgttggtatctcagttcggtgtaggtcgttcgctccaagctgggctgtgtgcacgaaccccccgttcagcccgac
cgctgcgccttatccggtaactatcgtcttgagtccaacccggtaagacacgacttatcgccactggcagcagccattggtaactg
atttagaggactttgtcttgaagttatgcacctgttaaggctaaactgaaagaacagattttggtgagtgcggtcctccaacccactt
accttggttcaaagagttggtagctcagcgaaccttgagaaaaccaccgttggtagcggtggtttttctttatttatgagatgatgaat
caatcggtctatcaagtcaacgaacagctattccgttactctagatttcagtgcaatttatctcttcaaatgtagcacctgaagtcagc
cccatacgatataagttgtaattctcatgttagtcatgccccgcgcccaccggaaggagctgactgggttgaaggctctcaagggc
atcggtcgagatcccggtgcctaatgagtgagctaacttacattaattgcgttgcgctcactgcccgctttccagtcgggaaacctgt
cgtgccagctgcattaatgaatcggccaacgcgcggggagaggcggtttgcgtattgggcgccagggtggtttttcttttcaccagt
gagacgggcaacagctgattgcccttcaccgcctggccctgagagagttgcagcaagcggtccacgctggtttgccccagcag
gcgaaaatcctgtttgatggtggttaacggcgggatataacatgagct

**PcIRED amino acid sequence - reference for numbering**

MSTSEPKSISILGLGQMGHAIASNFVSQGFKTFVWNRTPSKAADLVEQGAIQSPSSTECIRS
SPLSILCVNTDDIVMDILAAAGDIPGHTIVNIVNGSPQQVRKTAEKSISQYMAAGYLHGSVMA
SPGLVRSGGAMTIFAGSSETFKKWELTLQPLGVTLWLLDDVGAAPLYDNSLLSIVAGIFSGF
MQALAMIGAAGHSETEFARGFVIPLLGQTEEWLVRTAEEVQNKDYVAEKNGSPIAVALDAT
KNIFETAKELGVSSRLLQGFLDVVKEGVQRGQGREEISGLVRLLREPK

**PcIRED amino acid sequence - expression construct**
MGSSHHHHHHSSGLVPRGSHMSTSEPKSISILGLGQMGHAIASNFVSQGFKTFVWNRTPS
KAADLVEQGAIQSPSSTECIRSSPLSILCVNTDDIVMDILAAAGDIPGHTIVNIVNGSPQQVRK
TAEKSISQYMAAGYLHGSVMASPGLVRSGGAMTIFAGSSETFKKWELTLQPLGVTLWLLDD
VGAAPLYDNSLLSIVAGIFSGFMQALAMIGAAGHSETEFARGFVIPLLGQTEEWLVRTAEEV
QNKDYVAEKNGSPIAVALDATKNIFETAKELGVSSRLLQGFLDVVKEGVQRGQGREEISGLV
RLLREPK

**PcIRED nucleotide sequence - expression construct**
ATGGGCAGCAGCCATCATCATCATCATCACAGCAGCGGCCTGGTGCCGCGCGGCAGC
CATATGTCTACCTCTGAACCGAAATCAATTAGCATTCTGGGTTTAGGCCAGATGGGTCA
TGCAATTGCCTCTAATTTTGTGTCACAGGGCTTTAAAACCTTTGTTTGGAATCGTACCCC
GTCTAAAGCAGCCGATCTGGTTGAACAGGGCGCAATTCAGTCTCCGTCAAGTACCGAA
TGTATTCGCTCAAGCCCGCTGTCTATTCTGTGTGTGAATACCGATGATATTGTGATGGA
TATTCTGGCAGCAGCCGGCGATATTCCGGGTCATACCATTGTTAATATTGTTAATGGTA
GTCCGCAGCAGGTGCGCAAAACCGCAGAAAAATCAATTTCACAGTATATGGCAGCCGG
CTATCTGCATGGCTCAGTTATGGCAAGTCCGGGCCTGGTGCGTAGTGGCGGTGCCATG
ACCATTTTTGCCGGTAGTAGCGAAACCTTTAAAAAATGGGAACTGACCTTACAGCCGTT
AGGCGTGACCCTGTGGCTGCTGGATGATGTTGGTGCCGCCCCGCTGTATGATAATAGC
TTACTGAGCATTGTGGCGGGCATTTTTAGCGGTTTTATGCAGGCACTGGCCATGATTGG
TGCCGCAGGTCATAGTGAAACCGAATTTGCACGCGGCTTTGTGATTCCGTTACTGGGC
CAGACTGAAGAATGGTTAGTTCGTACCGCCGAAGAAGTTCAGAATAAAGATTATGTTGC
AGAAAAAAATGGCTCTCCGATTGCCGTTGCGTTAGATGCGACCAAAAATATTTTTGAAA
CCGCCAAAGAACTGGGCGTATCCAGTCGCTTATTACAGGGCTTTCTGGATGTTGTTAAA
GAAGGCGTTCAGCGCGGCCAGGGTCGTGAAGAAATTAGCGGTTTAGTTCGCCTGCTGC
GCGAACCGAAATAA

**PcIRED whole plasmid sequence:**
ctgtaacatcattggcaacgctacctttgccatgtttcagaaacaactctggcgcatcgggcttcccatacaatcgatagattgtcgc
acctgattgcccgacattatcgcgagcccatttatacccatataaatcagcatccatgttggaatttaatcgcggcctagagcaaga
cgtttcccgttgaatatggctcatactcttcctttttcaatattattgaagcatttatcagggttattgtctcatgagcggatacatatttgaat
gtatttagaaaaataaacaaataggcatgcagcgctcttccgcttcctcgctcactgactcgctacgctcggtcgttcgactgcggc
gagcggtgtcagctcactcaaaagcggtaatacggttatccacagaatcaggggataaagccggaaagaacatgtgagcaa
aaagcaaagcaccggaagaagccaacgccgcaggcgtttttccataggctccgcccccctgacgagcatcacaaaaatcga
cgctcaagccagaggtggcgaaacccgacaggactataaagataccaggcgtttccccctggaagctccctcgtgcgctctcct
gttccgaccctgccgcttaccggatacctgtccgcctttctcccttcgggaagcgtggcgctttctcatagctcacgctgttggtatctc
agttcggtgtaggtcgttcgctccaagctgggctgtgtgcacgaaccccccgttcagcccgaccgctgcgccttatccggtaactat
cgtcttgagtccaacccggtaagacacgacttatcgccactggcagcagccattggtaactgatttagaggactttgtcttgaagtt
atgcacctgttaaggctaaactgaaagaacagatttggtgagtgcggtcctccaacccacttaccttggttcaaagagttggtagc
tcagcgaaccttgagaaaaccaccgttggtagcggtggtttttctttatttatgagatgatgaatcaatcggtctatcaagtcaacga
acagctattccgttactctagatttcagtgcaatttatctcttcaaatgtagcacctgaagtcagccccatacgatataagttgtaattct
catgttagtcatgccccgcgcccaccggaaggagctgactgggttgaaggctctcaagggcatcggtcgagatcccggtgccta
atgagtgagctaacttacattaattgcgttgcgctcactgcccgctttccagtcgggaaacctgtcgtgccagctgcattaatgaatc
ggccaacgcgcggggagaggcggtttgcgtattgggcgccagggtggtttttcttttcaccagtgagacgggcaacagctgattg
cccttcaccgcctggccctgagagagttgcagcaagcggtccacgctggtttgccccagcaggcgaaaatcctgtttgatggtgg
ttaacggcgggatataacatgagctgtcttcggtatcgtcgtatcccactaccgagatgtccgcaccaacgcgcagcccggactc
ggtaatggcgcgcattgcgcccagcgccatctgatcgttggcaaccagcatcgcagtgggaacgatgccctcattcagcatttgc
atggtttgttgaaaaccggacatggcactccagtcgccttcccgttccgctatcggctgaatttgattgcgagtgagatatttatgcca
gccagccagacgcagacgcgccgagacagaacttaatgggcccgctaacagcgcgatttgctggtgacccaatgcgaccag

atgctccacgcccagtcgcgtaccgtcttcatgggagaaaataatactgttgatgggtgtctggtcagagacatcaagaaataac
gccggaacattagtgcaggcagcttccacagcaatggcatcctggtcatccagcggatagttaatgatcagcccactgacgcgtt
gcgcgagaagattgtgcaccgccgctttacaggcttcgacgccgcttcgttctaccatcgacaccaccacgctggcacccagttg
atcggcgcgagatttaatcgccgcgacaatttgcgacggcgcgtgcagggccagactggaggtggcaacgccaatcagcaac
gactgtttgcccgccagttgttgtgccacgcggttgggaatgtaattcagctccgccatcgccgcttccactttttcccgcgttttcgca
gaaacgtggctggcctggttcaccacgcgggaaacggtctgataagagacaccggcatactctgcgacatcgtataacgttact
ggtttcacattcaccaccctgaattgactctcttccgggcgctatcatgccataccgcgaaaggttttgcgccattcgatggtgtccgg
gatctcgacgctctcccttatgcgactcctgcattaggaaattaatacgactcactataggggaattgtgagcggataacaattcC
CCTGTAGAAATAATTTTGTTTAACTTTAATAAGGAGATATACCATGGGCAGCAGCCATCA
TCATCATCATCACAGCAGCGGCCTGGTGCCGCGCGGCAGCCATATGTCTACCTCTGAA
CCGAAATCAATTAGCATTCTGGGTTTAGGCCAGATGGGTCATGCAATTGCCTCTAATTTT
GTGTCACAGGGCTTTAAAACCTTTGTTTGGAATCGTACCCCGTCTAAAGCAGCCGATCT
GGTTGAACAGGGCGCAATTCAGTCTCCGTCAAGTACCGAATGTATTCGCTCAAGCCCG
CTGTCTATTCTGTGTGTGAATACCGATGATATTGTGATGGATATTCTGGCAGCAGCCGG
CGATATTCCGGGTCATACCATTGTTAATATTGTTAATGGTAGTCCGCAGCAGGTGCGCA
AAACCGCAGAAAAATCAATTTCACAGTATATGGCAGCCGGCTATCTGCATGGCTCAGTT
ATGGCAAGTCCGGGCCTGGTGCGTAGTGGCGGTGCCATGACCATTTTTGCCGGTAGTA
GCGAAACCTTTAAAAAATGGGAACTGACCTTACAGCCGTTAGGCGTGACCCTGTGGCT
GCTGGATGATGTTGGTGCCGCCCCGCTGTATGATAATAGCTTACTGAGCATTGTGGCG
GGCATTTTTAGCGGTTTTATGCAGGCACTGGCCATGATTGGTGCCGCAGGTCATAGTG
AAACCGAATTTGCACGCGGCTTTGTGATTCCGTTACTGGGCCAGACTGAAGAATGGTTA
GTTCGTACCGCCGAAGAAGTTCAGAATAAAGATTATGTTGCAGAAAAAAATGGCTCTCC
GATTGCCGTTGCGTTAGATGCGACCAAAAATATTTTTGAAACCGCCAAAGAACTGGGCG
TATCCAGTCGCTTATTACAGGGCTTTCTGGATGTTGTTAAAGAAGGCGTTCAGCGCGGC
CAGGGTCGTGAAGAAATTAGCGGTTTAGTTCGCCTGCTGCGCGAACCGAAATAATCTG
GTAAAGAAACCGCTGCtgcgaaattgaacgccagcacatggactcgtctactagcgcagcttaattaacctaggctg
ctgccaccgctgagcaataactagcataaccccttggggcctctaaacgggtcttgaggggttttttgctgaaacctcaggcatttg
agaagcacacggtcacactgcttccggtagtcaataaaccggtaaaccagcaatagacataagcggctatttaacgaccctgc
cctgaaccgacgacaagctgacgaccgggtctccgcaagtggcacttttcggggaaatgtgcgcggaacccctatttgtttattttt
ctaaatacattcaaatatgtatccgctcatgaattaattcttagaaaaactcatcgagcatcaaatgaaactgcaatttattcatatca
ggattatcaataccatatttttgaaaaagccgtttctgtaatgaaggagaaaactcaccgaggcagttccataggatggcaagatc
ctggtatcggtctgcgattccgactcgtccaacatcaatacaacctattaatttcccctcgtcaaaaataaggttatcaagtgagaaa
tcaccatgagtgacgactgaatccggtgagaatggcaaaagtttatgcatttctttccagacttgttcaacaggccagccattacgc
tcgtcatcaaaatcactcgcatcaaccaaaccgttattcattcgtgattgcgcctgagcgagacgaaatacgcggtcgctgttaaa
aggacaattacaaacaggaatcgaatgcaaccggcgcaggaacactgccagcgcatcaacaatattttcacctgaatcagga
tattcttctaatacctggaatgctgttttcccggggatcgcagtggtgagtaaccatgcatcatcaggagtacggataaaatgcttga
tggtcggaagaggcataaattccgtcagccagtttagtctgaccatctcat

**ScIRED amino acid sequence - reference for numbering**
MRDTDVTVLGLGLMGQALAGAFLKDGHATTVWNRSEGKAGRLAEQGAVLASSARDAAEA
SPLVVVCVSDHAAVRAVLDPLGDVLAGRVLVNLTSGTSDQARATAEWAAERGITYLDGAIM
AIPQVVGTADAFLLYSGPEAAYEAHEPTLRSLGAGTTYLGADHGLSSLYDVALLGIMWGTLN
SFLHGAALLGTAKVEATTFAPFANRWIAAVTGFVSAYADQVDQGAYPALDATIDTHVA**T**VDH
LIHESEAAGVNTELPRLVRTLADRALADGQGGLGYAAMIEQFRRPSA
*T241 equivalent in bold

**ScIRED amino acid sequence - expression construct**
MRDTDVTVLGLGLMGQALAGAFLKDGHATTVWNRSEGKAGRLAEQGAVLASSARDAAEA
SPLVVVCVSDHAAVRAVLDPLGDVLAGRVLVNLTSGTSDQARATAEWAAERGITYLDGAIM
AIPQVVGTADAFLLYSGPEAAYEAHEPTLRSLGAGTTYLGADHGLSSLYDVALLGIMWGTLN
SFLHGAALLGTAKVEATTFAPFANRWIAAVTGFVSAYADQVDQGAYPALDATIDTHVA**T**VDH
LIHESEAAGVNTELPRLVRTLADRALADGQGGLGYAAMIEQFRRPSA*
*T241 equivalent in bold

**ScIRED nucleotide sequence - expression construct**
ATGGGCAGCAGCCATCATCATCATCATCACAGCAGCGGCCTGGTGCCGCGCGGCAGC
CATATGCGTGATACCGATGTTACCGTTTTAGGTCTGGGTCTGATGGGTCAAGCACTGGC
AGGCGCATTTCTGAAAGATGGTCATGCAACCACCGTTTGGAATCGTAGCGAAGGTAAA
GCAGGTCGTCTGGCAGAACAGGGTGCAGTTCTGGCAAGCAGCGCACGTGATGCAGCA
GAAGCAAGTCCGCTGGTTGTTGTTTGTGTTAGCGATCATGCAGCAGTTCGTGCCGTTCT
GGATCCGCTGGGTGATGTTCTGGCAGGTCGTGTTCTGGTTAATCTGACCAGCGGCACC
AGCGATCAGGCACGTGCAACCGCAGAATGGGCAGCAGAACGTGGTATTACCTATCTGG
ATGGTGCAATTATGGCAATTCCGCAGGTTGTGGGCACCGCAGATGCATTTCTGCTGTAT
AGCGGTCCGGAAGCAGCATATGAAGCACATGAACCGACACTGCGTAGCTTAGGTGCAG
GCACCACATATCTGGGTGCAGATCATGGTCTGAGCAGCCTGTATGATGTTGCACTGCT
GGGTATTATGTGGGGCACCCTGAATAGCTTTCTGCATGGTGCAGCCCTGCTGGGCACA
GCAAAAGTTGAAGCCACCACCTTTGCACCGTTTGCAAATCGTTGGATTGCAGCCGTTAC
CGGTTTTGTTAGCGCATATGCCGATCAGGTTGATCAGGGTGCATATCCGGCACTGGAT
GCAACCATTGATACCCATGTTGCAACCGTTGATCATCTGATTCATGAAAGCGAAGCAGC
CGGTGTTAATACCGAACTGCCTCGTCTGGTTCGTACCCTGGCAGATCGTGCACTGGCC
GATGGTCAAGGTGGTCTGGGTTATGCAGCAATGATTGAACAGTTTCGTCGTCCGAGCG
CATAA

**SmIRED amino acid sequence - reference for numbering**
MRDTDVTVLGLGLMGQALAGAFLKDGHATTVWNRSAGKAGRLAEQGAVLASSARDAAEA
SPLVVVCVSDHTAVRAVLDPLGDVLAGRVLVNLTSGTSEQARATAEWAAERGITYLDGAIM
AIPQVVGTADAFLLYSGPEAAYEAHEPTLRSLGAGTTYLGADHGLSSLYDVALLGIMWGTLN
SFLHGAALLGTAKVEATTFAPFANRWIEAVTGFVSAYAGQVDQGAYPALDATIDTHVA**T**VD
HLIHESEAAGVNTELPRLVRTLADRALAEGQGGLGYAAMIEQFRRPSA*
*T241 equivalent in bold

**SmIRED amino acid sequence - expression construct**
MGSSHHHHHHSSGLVPRGSHMRDTDVTVLGLGLMGQALAGAFLKDGHATTVWNRSAGK
AGRLAEQGAVLASSARDAAEASPLVVVCVSDHTAVRAVLDPLGDVLAGRVLVNLTSGTSEQ
ARATAEWAAERGITYLDGAIMAIPQVVGTADAFLLYSGPEAAYEAHEPTLRSLGAGTTYLGA
DHGLSSLYDVALLGIMWGTLNSFLHGAALLGTAKVEATTFAPFANRWIEAVTGFVSAYAGQ
VDQGAYPALDATIDTHVA**T**VDHLIHESEAAGVNTELPRLVRTLADRALAEGQGGLGYAAMIE
QFRRPSA*
*T241 equivalent in bold

**SmIRED nucleotide sequence - expression construct**
ATGGGCAGCAGCCATCATCATCATCATCACAGCAGCGGCCTGGTGCCGCGCGGCAGC
CATATGCGTGATACCGATGTTACCGTTTTAGGTCTGGGTCTGATGGGTCAAGCACTGGC
AGGCGCATTTCTGAAAGATGGTCATGCAACCACCGTTTGGAATCGTAGCGCAGGTAAA
GCAGGTCGTCTGGCAGAACAGGGTGCAGTTCTGGCAAGCAGCGCACGTGATGCAGCA
GAAGCAAGTCCGCTGGTTGTTGTTTGTGTTAGCGATCATACCGCAGTTCGTGCCGTTCT
GGATCCGCTGGGTGATGTTCTGGCAGGTCGTGTTCTGGTTAATCTGACCAGCGGCACC
AGCGAACAGGCACGTGCAACCGCAGAATGGGCAGCAGAACGTGGTATTACCTATCTGG
ATGGTGCAATTATGGCAATTCCGCAGGTTGTGGGCACCGCAGATGCATTTCTGCTGTAT
AGCGGTCCGGAAGCAGCATATGAAGCACATGAACCGACACTGCGTAGCTTAGGTGCAG
GCACCACATATCTGGGTGCAGATCATGGTCTGAGCAGCCTGTATGATGTTGCACTGCT
GGGTATTATGTGGGGCACCCTGAATAGCTTTCTGCATGGTGCAGCCCTGCTGGGCACA
GCAAAAGTTGAAGCCACCACCTTTGCACCGTTTGCAAATCGTTGGATTGAAGCAGTTAC
CGGTTTTGTTAGCGCATATGCAGGTCAGGTTGATCAGGGTGCATATCCGGCACTGGAT
GCAACCATTGATACCCATGTTGCAACCGTTGATCATCTGATTCATGAAAGCGAAGCAGC
CGGTGTTAATACCGAACTGCCTCGTCTGGTTCGTACCCTGGCAGATCGTGCACTGGCC
GAAGGTCAAGGTGGTCTGGGTTATGCAGCAATGATTGAACAGTTTCGTCGTCCGAGCG
CATAA

**SbIRED amino acid sequence - reference for numbering**
MGQSPVTVIGLGLMGQALAGTFLGNGHPTTVWNRSAEKADELVARGATLAASVRDAVEAS
PLVIVCVSDYAAAHGLLAPLGDALAGRVLVNLTSGTSEQARESAEWAAGEGVTYLDGAIMAI
PPVIGTADAFLLYSGPQDAYDAHEPTLKALGAGTTYLGADHGLASLYDVALLGIMWGTLNSF
LHGAALVGTAKVDATVFAPFANRWIEAVTGFVSAYAGQIDEGAYPALDATIDTHVA**T**MDHLI
HESESGGVNTELPRLVKALADRAVAEGRGGDGYAAMIEQFRRPSA*
*T241 equivalent in bold

**SbIRED amino acid sequence - expression construct**
MGSSHHHHHHSSGLVPRGSHMGQSPVTVIGLGLMGQALAGTFLGNGHPTTVWNRSAEKA
DELVARGATLAASVRDAVEASPLVIVCVSDYAAAHGLLAPLGDALAGRVLVNLTSGTSEQA
RESAEWAAGEGVTYLDGAIMAIPPVIGTADAFLLYSGPQDAYDAHEPTLKALGAGTTYLGAD
HGLASLYDVALLGIMWGTLNSFLHGAALVGTAKVDATVFAPFANRWIEAVTGFVSAYAGQID
EGAYPALDATIDTHVA**T**MDHLIHESESGGVNTELPRLVKALADRAVAEGRGGDGYAAMIEQ
FRRPSA*
*T241 equivalent in bold

**SbIRED nucleotide sequence - expression construct**
ATGGGCAGCAGCCATCATCATCATCATCACAGCAGCGGCCTGGTGCCGCGCGGCAGC
CATATGGGTCAGAGTCCGGTTACCGTTATTGGTCTGGGTCTGATGGGTCAAGCACTGG
CAGGCACCTTTTTAGGTAATGGTCATCCGACCACCGTTTGGAATCGTAGCGCAGAAAAA
GCAGATGAACTGGTTGCACGTGGTGCAACCCTGGCAGCAAGCGTTCGTGATGCAGTTG
AAGCAAGTCCGCTGGTTATTGTTTGTGTTAGCGATTATGCAGCAGCACATGGTCTGCTG
GCACCGCTGGGTGATGCCCTGGCAGGTCGTGTTCTGGTTAATCTGACCAGCGGCACCA
GCGAACAGGCACGTGAAAGCGCAGAATGGGCAGCCGGTGAAGGTGTTACCTATCTGG
ATGGTGCAATTATGGCAATTCCGCCTGTTATTGGCACCGCAGATGCATTTCTGCTGTAT
AGCGGTCCGCAGGATGCATATGATGCACATGAACCGACACTGAAAGCATTAGGTGCAG
GCACCACATATCTGGGTGCAGATCATGGTCTGGCAAGCCTGTATGATGTTGCACTGCT
GGGTATTATGTGGGGCACCCTGAATAGCTTTCTGCATGGTGCAGCACTGGTTGGTACA
GCAAAAGTTGATGCAACCGTTTTTGCACCGTTTGCAAATCGTTGGATTGAAGCAGTTAC
CGGTTTTGTTAGCGCATATGCCGGTCAGATTGATGAAGGTGCATATCCGGCACTGGAT
GCAACCATTGATACCCATGTTGCCACCATGGATCATCTGATTCATGAAAGCGAAAGCGG
TGGTGTTAATACCGAACTGCCTCGTCTGGTTAAAGCGCTGGCAGATCGTGCAGTTGCC
GAAGGTCGTGGTGGTGATGGTTATGCAGCCATGATTGAACAGTTTCGTCGTCCGAGCG
CATAA

**BtIRED amino acid sequence - reference for numbering**
MRETDVTVLGLGLMGQAIAGAFLKGGREVTVWNRTAAKADALVAQGAQRGETAAEAVGA
SGLVVVVLTDYTAVTGVLEPLADTLAGRTVVNLTSGTAAQAREFAEWAAKRDITYLDGAIMA
IPQVVGTDDAFLLYGGAKEVYDAHEAVLRDLGASGTIHLGADHGLASLYDVALLGIMWGTLN
SFLHGAALVGTAGVDAKGFSEFANKWIGAITGFVSAYAAQIDEGSYTALDASIDTHAA**T**VDH
LIEESEAAGVNAEVPKLVKSFTDRARAEGHGLDSYAAMITQFRRP*
*T241 equivalent in bold

**BtIRED amino acid sequence - expression construct**
MGSSHHHHHHSSGLVPRGSHMRETDVTVLGLGLMGQAIAGAFLKGGREVTVWNRTAAKA
DALVAQGAQRGETAAEAVGASGLVVVVLTDYTAVTGVLEPLADTLAGRTVVNLTSGTAAQA
REFAEWAAKRDITYLDGAIMAIPQVVGTDDAFLLYGGAKEVYDAHEAVLRDLGASGTIHLGA
DHGLASLYDVALLGIMWGTLNSFLHGAALVGTAGVDAKGFSEFANKWIGAITGFVSAYAAQI
DEGSYTALDASIDTHAA**T**VDHLIEESEAAGVNAEVPKLVKSFTDRARAEGHGLDSYAAMITQ
FRRP*
*T241 equivalent in bold

**BtIRED nucleotide sequence - expression construct**

ATGGGCAGCAGCCATCATCATCATCATCACAGCAGCGGCCTGGTGCCGCGCGGCAGC
CATATGCGTGAAACCGATGTTACCGTTTTAGGTCTGGGTCTGATGGGTCAAGCAATTGC
CGGTGCATTTCTGAAAGGTGGTCGTGAAGTTACCGTGTGGAATCGTACCGCAGCAAAA
GCAGATGCACTGGTTGCACAGGGTGCACAGCGTGGTGAAACCGCAGCCGAAGCAGTT
GGTGCAAGCGGTCTGGTTGTTGTTGTTCTGACCGATTATACCGCAGTTACCGGTGTTCT
GGAACCGCTGGCAGATACCCTGGCAGGTCGTACCGTTGTTAATCTGACCAGCGGCACC
GCAGCACAGGCACGTGAATTTGCAGAATGGGCAGCAAAACGTGATATTACCTATCTGG
ATGGTGCAATTATGGCAATTCCGCAGGTTGTTGGCACCGATGATGCCTTTCTGCTGTAT
GGTGGTGCAAAAGAAGTTTACGACGCACACGAAGCCGTTCTGCGTGATCTGGGTGCGA
GCGGCACCATTCACCTGGGTGCAGATCATGGTCTGGCAAGCCTGTATGATGTTGCACT
GCTGGGTATTATGTGGGGCACCCTGAATAGCTTTCTGCATGGTGCAGCCCTGGTTGGT
ACAGCCGGTGTTGATGCAAAAGGTTTTAGCGAATTTGCCAACAAATGGATTGGTGCCAT
TACCGGTTTTGTTAGCGCATATGCAGCACAGATTGATGAAGGTAGCTATACCGCACTGG
ATGCAAGCATTGATACCCATGCAGCAACGGTTGATCATCTGATTGAAGAAAGCGAAGCA
GCGGGTGTTAATGCAGAAGTTCCGAAACTGGTTAAAAGCTTTACCGATCGTGCACGTG
CCGAAGGTCATGGCCTGGATAGCTATGCAGCAATGATTACCCAGTTTCGTCGTCCGTAA

**SIIRED amino acid sequence - reference for numbering**
MNRTPVTVIGLGLMGQALAGAFLKAGHPTTVWNRSAEKAADLVSEGAVLAPSAEDAVKAS
GLVVLCVTDYDVVHTVLDPLSGELSGKTLVNLTSGHSEGARETAEWAAKLGAAYLDGAIMAI
PPVIGTEHAVLLYAGSKEVYDANESALQAVAPAGTTHLGEDHGLASLYDVALLGIMWGVLN
SFLHGAALLGAANVKASTFAPFANNWIGAVTNFVTAYAGQIDEGDFTAHDATIDTHLA**T**MIHL
IHESEATGISPELPEFVKALTDRAVAAGQNGLSYAAMIDQFRKPSK*
*T241 equivalent in bold

**SIIRED amino acid sequence - expression construct**
MGSSHHHHHHSSGLVPRGSHMNRTPVTVIGLGLMGQALAGAFLKAGHPTTVWNRSAEKA
ADLVSEGAVLAPSAEDAVKASGLVVLCVTDYDVVHTVLDPLSGELSGKTLVNLTSGHSEGA
RETAEWAAKLGAAYLDGAIMAIPPVIGTEHAVLLYAGSKEVYDANESALQAVAPAGTTHLGE
DHGLASLYDVALLGIMWGVLNSFLHGAALLGAANVKASTFAPFANNWIGAVTNFVTAYAGQI
DEGDFTAHDATIDTHLA**T**MIHLIHESEATGISPELPEFVKALTDRAVAAGQNGLSYAAMIDQF
RKPSK*
*T241 equivalent in bold

**SIIRED nucleotide sequence - expression construct**
ATGGGCAGCAGCCATCATCATCATCATCACAGCAGCGGCCTGGTGCCGCGCGGCAGC
CATATGAATCGTACACCGGTTACCGTTATTGGTCTGGGTCTGATGGGTCAAGCACTGGC
AGGCGCATTTCTGAAAGCAGGTCATCCGACCACCGTTTGGAATCGTAGCGCAGAAAAA
GCAGCAGATCTGGTTAGCGAAGGTGCAGTTCTGGCACCGAGCGCAGAAGATGCAGTTA
AAGCAAGCGGTCTGGTTGTTCTGTGTGTTACCGATTATGATGTTGTTCATACCGTTCTG
GATCCGCTGAGCGGTGAACTGAGCGGTAAAACCCTGGTTAATCTGACCAGCGGTCATA
GTGAAGGTGCACGTGAAACCGCAGAATGGGCAGCAAAACTGGGTGCAGCATATCTGGA
TGGTGCAATTATGGCAATTCCGCCTGTTATTGGCACCGAACATGCCGTTCTGCTGTATG
CAGGTAGCAAAGAAGTTTACGACGCAAACGAAAGCGCACTGCAGGCAGTTGCACCGGC
AGGTACAACCCATCTGGGTGAAGATCATGGTCTGGCAAGCCTGTATGATGTGGCACTG
CTGGGTATTATGTGGGGTGTTCTGAATAGCTTTCTGCATGGTGCAGCCCTGCTGGGTG
CCGCAAATGTTAAAGCCAGCACCTTTGCACCGTTTGCCAATAATTGGATTGGTGCCGTT
ACCAATTTTGTTACCGCATATGCAGGTCAGATTGATGAAGGTGATTTTACCGCACATGAT
GCAACCATTGATACACATCTGGCAACCATGATTCATCTGATTCATGAAAGCGAAGCAAC
CGGTATTAGTCCGGAACTGCCGGAATTTGTTAAAGCACTGACCGATCGTGCAGTTGCA
GCGGGTCAGAATGGTCTGAGCTATGCAGCAATGATTGATCAGTTTCGTAAACCGAGCA
AATAA

**SxIRED amino acid sequence - reference for numbering**
MNHTPVTVVGLGLMGQALAGAFLKAGHATTVWNRSSDKAAGLVGEGATLAPSLKEAIEAS
PLIVLCVTDYDVVRSLLDSVSGDLAGKTLVNLTSGHSEGARETAEWVLEHDAQYLDGAIMAI
PPVIGTEHATLLYGGSKEAYDAHESALKALGGATHLGTDYQLASLYDVALLGIMWGVLNSFL
HGAALVGTAGVDASTFAPFANNWIGAVTNFVSAYAGQIDQGDFTAHDATIDTHLA**T**MIHLIH
ESKAKGISSELPEFVKAVTDRAVAAGEGKSSYAAMFEQFKNAAK*
*T241 equivalent in bold

**SxIRED amino acid sequence - expression construct**
MGSSHHHHHHSSGLVPRGSHMNHTPVTVVGLGLMGQALAGAFLKAGHATTVWNRSSDK
AAGLVGEGATLAPSLKEAIEASPLIVLCVTDYDVVRSLLDSVSGDLAGKTLVNLTSGHSEGA
RETAEWVLEHDAQYLDGAIMAIPPVIGTEHATLLYGGSKEAYDAHESALKALGGATHLGTDY
QLASLYDVALLGIMWGVLNSFLHGAALVGTAGVDASTFAPFANNWIGAVTNFVSAYAGQID
QGDFTAHDATIDTHLA**T**MIHLIHESKAKGISSELPEFVKAVTDRAVAAGEGKSSYAAMFEQF
KNAAK*
*T241 equivalent in bold

**SxIRED nucleotide sequence - expression construct**
ATGGGCAGCAGCCATCATCATCATCATCACAGCAGCGGCCTGGTGCCGCGCGGCAGC
CATATGAATCATACACCGGTTACCGTTGTTGGTCTGGGTCTGATGGGTCAAGCACTGGC
AGGCGCATTTCTGAAAGCAGGTCATGCAACCACCGTTTGGAATCGTAGCAGCGATAAA
GCAGCAGGTCTGGTTGGTGAAGGTGCAACCCTGGCACCGAGCCTGAAAGAAGCAATT
GAAGCAAGTCCGCTGATTGTTCTGTGTGTTACCGATTATGATGTTGTTCGTAGCCTGCT
GGATAGCGTTAGCGGTGATCTGGCAGGTAAAACCCTGGTTAATCTGACCAGCGGTCAT
AGCGAAGGTGCACGTGAAACCGCAGAATGGGTTTTAGAACATGATGCACAGTATCTGG
ATGGTGCAATTATGGCAATTCCGCCTGTTATTGGCACCGAACATGCGACCCTGCTGTAT
GGTGGTAGCAAAGAAGCATACGACGCACACGAAAGCGCACTGAAAGCATTAGGTGGTG
CGACCCATCTGGGCACCGATTATCAGCTGGCAAGCCTGTATGATGTGGCACTGCTGGG
TATTATGTGGGGTGTTCTGAATAGCTTTCTGCATGGTGCAGCACTGGTTGGTACAGCCG
GTGTTGATGCAAGCACCTTTGCACCGTTTGCAAATAATTGGATTGGTGCCGTTACCAAT
TTTGTTAGCGCATATGCAGGTCAGATTGATCAGGGCGATTTTACCGCACATGATGCCAC
CATTGATACACATCTGGCAACCATGATTCATCTGATCCATGAAAGCAAAGCCAAAGGTA
TTAGCAGCGAACTGCCGGAATTTGTTAAAGCAGTTACCGATCGTGCAGTTGCAGCCGG
TGAAGGTAAAAGCAGCTATGCAGCAATGTTTGAGCAGTTTAAAAACGCAGCCAAATAA

**LpIRED amino acid sequence - reference for numbering**
MEPVTVIGLGLMGSALAAAFVEAGYPTTIWNRTPGKADELVARGAVLVPFVADALRASSLIV
VCLSDAASVEDVLGPHRDELDSRTLVNLTSGTSDEARGISKWAGDYVDGAIMAIPEVIGRPE
AFLLFSGSAEAYARHRESLGRLGTTTFFGEDVGLASLYDVALLGVMWGTLNSFLHGAALLG
AAGVDAAEFAPFANQWVSSVTGFVNAYADQIDRGVYPAEDASLETHLA**T**MKYLVRESSAA
EVNTEWPARIQAMTERAIRAGHAGASYASLIEVFRSRA*
*T241 equivalent in bold

**LpIRED amino acid sequence - expression construct**
MGSSHHHHHHSSGLVPRGSHMEPVTVIGLGLMGSALAAAFVEAGYPTTIWNRTPGKADEL
VARGAVLVPFVADALRASSLIVVCLSDAASVEDVLGPHRDELDSRTLVNLTSGTSDEARGIS
KWAGDYVDGAIMAIPEVIGRPEAFLLFSGSAEAYARHRESLGRLGTTTFFGEDVGLASLYDV
ALLGVMWGTLNSFLHGAALLGAAGVDAAEFAPFANQWVSSVTGFVNAYADQIDRGVYPAE
DASLETHLA**T**MKYLVRESSAAEVNTEWPARIQAMTERAIRAGHAGASYASLIEVFRSRA*
*T241 equivalent in bold

**LpIRED nucleotide sequence - expression construct**
ATGGGCAGCAGCCATCATCATCATCATCACAGCAGCGGCCTGGTGCCGCGCGGCAGC
CATATGGAACCGGTTACCGTTATTGGTCTGGGTCTGATGGGTAGCGCACTGGCAGCAG
CATTTGTTGAAGCAGGTTATCCGACCACCATTTGGAATCGTACACCGGGTAAAGCAGAT

GAACTGGTTGCACGTGGTGCAGTTCTGGTTCCGTTTGTTGCAGATGCACTGCGTGCAA
GCAGCCTGATTGTTGTTTGTCTGAGTGATGCAGCAAGCGTTGAAGATGTTCTGGGTCC
GCATCGTGATGAGCTGGATAGTCGTACCCTGGTTAATCTGACCAGCGGCACCAGTGAT
GAAGCACGTGGTATTAGCAAATGGGCAGGCGATTATGTTGATGGTGCAATTATGGCAAT
TCCGGAAGTGATTGGTCGTCCGGAAGCATTTCTGCTGTTTAGCGGTAGTGCCGAAGCA
TATGCACGTCATCGTGAAAGCCTGGGTCGTCTGGGTACAACCACCTTTTTCGGCGAAG
ATGTTGGTCTGGCAAGCCTGTATGATGTTGCACTGCTGGGTGTTATGTGGGGCACCCT
GAATAGCTTTCTGCATGGTGCAGCCCTGCTGGGAGCAGCCGGTGTTGATGCAGCCGAA
TTTGCACCGTTTGCAAATCAGTGGGTTAGCAGCGTTACCGGTTTTGTTAATGCATATGC
CGATCAGATTGATCGTGGTGTTTATCCGGCAGAAGATGCAAGCCTGGAAACCCATCTG
GCAACCATGAAATATCTGGTTCGTGAAAGTAGCGCAGCCGAAGTTAATACCGAATGGC
CTGCACGTATTCAGGCAATGACCGAACGTGCAATTCGTGCAGGTCATGCCGGTGCAAG
CTATGCAAGTCTGATTGAAGTTTTTCGTAGCCGTGCATAA

**SolRED amino acid sequence - reference for numbering**
MTARTAVAVIGLGLMGQALAQALLKQGHTTTVWNRSPEKADALAAQGAVRAGTAAEAVTA
APLVLICVSTYEVVDELLAPLTGALAGRTVVNLTSGTPEQARTTAQWAARHGIGYLDGAVM
SVPEGVGEPGTILLYSGPHESFETHRATLELLGGGTTYLSSDAGLASLYDVSLLGLMWSTM
SGYVHAAALVGTEGVDAETFTRVGNMWLATISGYLTAYAGQIDSRTYPADATLHTQAM**T**M
DHVIHASEERGIDSVIPRAIKALTERGIAAGHGDDSFASLIEVVRNSKPVE*
*T241 equivalent in bold

**SolRED amino acid sequence - expression construct**
MGSSHHHHHHSSGLVPRGSHMTARTAVAVIGLGLMGQALAQALLKQGHTTTVWNRSPEK
ADALAAQGAVRAGTAAEAVTAAPLVLICVSTYEVVDELLAPLTGALAGRTVVNLTSGTPEQA
RTTAQWAARHGIGYLDGAVMSVPEGVGEPGTILLYSGPHESFETHRATLELLGGGTTYLSS
DAGLASLYDVSLLGLMWSTMSGYVHAAALVGTEGVDAETFTRVGNMWLATISGYLTAYAG
QIDSRTYPADATLHTQAM**T**MDHVIHASEERGIDSVIPRAIKALTERGIAAGHGDDSFASLIEV
VRNSKPVE*
*T241 equivalent in bold

**SolRED nucleotide sequence - expression construct**
ATGGGCAGCAGCCATCATCATCATCATCACAGCAGCGGCCTGGTGCCGCGCGGCAGC
CATATGACCGCACGTACCGCAGTTGCAGTTATTGGTCTGGGTCTGATGGGTCAAGCAC
TGGCACAGGCACTGCTGAAACAGGGTCATACCACCACCGTTTGGAATCGTAGTCCGGA
AAAAGCAGATGCACTGGCAGCACAGGGTGCAGTTCGTGCAGGCACCGCAGCAGAAGC
AGTTACCGCAGCACCGCTGGTTCTGATTTGTGTTAGCACCTATGAAGTTGTTGATGAAC
TGCTGGCACCGCTGACCGGTGCGCTGGCAGGTCGTACCGTTGTTAATCTGACCAGCG
GTACACCGGAACAGGCACGTACCACCGCACAGTGGGCAGCACGTCATGGTATTGGTTA
TCTGGATGGTGCAGTTATGAGCGTTCCGGAAGGTGTTGGTGAACCGGGTACAATTCTG
CTGTATAGCGGTCCGCATGAAAGCTTTGAAACCCATCGTGCAACCCTGGAACTGTTAG
GCGGTGGCACCACCTATCTGAGCAGTGATGCAGGTCTGGCAAGCCTGTATGATGTTAG
CCTGCTGGGTTTAATGTGGTCAACCATGAGCGGTTATGTTCATGCAGCAGCACTGGTTG
GCACCGAAGGTGTGGATGCAGAAACCTTTACACGTGTTGGTAATATGTGGCTGGCCAC
CATTTCAGGTTATCTGACCGCCTATGCAGGCCAGATTGATAGCCGTACCTATCCGGCAG
ATGCGACCCTGCATACCCAGGCAATGACCATGGATCATGTTATTCATGCAAGCGAAGAA
CGTGGTATTGATAGCGTTATTCCGCGTGCAATTAAAGCACTGACCGAACGCGGTATTGC
AGCCGGTCATGGTGATGATAGCTTTGCAAGCCTGATTGAAGTTGTGCGTAATAGCAAAC
CGGTGGAATAA

**AsIRED amino acid sequence - reference for numbering**
MTPVTFIGLGPMGQAMVRSLLAGGHPVTVWNRTASRADGVVADGAKRADSAALAIAENEL
VILSLTDYQAMYDILGPVGDALRGRVVVNLSSDTPSRTREAAEWLAERGAELITGGVMVPA
EWVGKDASYVFYSGPEAVFERFRDTLVLIGRTDYLGADHALAQLFYQAQLDIFLTALSAYLH

ASALLEAHGVTAEKFWPYAESNLNTIAPMMVEAPGQLDSGEYPGDQANAKMMGA**T**ADHIV
QASYEAGIDTGLPRAVKDLYDRAIAAGDGGKSWTSLFEQLRRRG*
*T241 equivalent in bold

**AsIRED amino acid sequence - expression construct**
MGSSHHHHHHSSGLVPRGSHMTPVTFIGLGPMGQAMVRSLLAGGHPVTVWNRTASRAD
GVVADGAKRADSAALAIAENELVILSLTDYQAMYDILGPVGDALRGRVVVNLSSDTPSRTRE
AAEWLAERGAELITGGVMVPAEWVGKDASYVFYSGPEAVFERFRDTLVLIGRTDYLGADHA
LAQLFYQAQLDIFLTALSAYLHASALLEAHGVTAEKFWPYAESNLNTIAPMMVEAPGQLDSG
EYPGDQANAKMMGA**T**ADHIVQASYEAGIDTGLPRAVKDLYDRAIAAGDGGKSWTSLFEQL
RRRG*
*T241 equivalent in bold

**AsIRED nucleotide sequence - expression construct**
ATGGGCAGCAGCCATCATCATCATCATCACAGCAGCGGCCTGGTGCCGCGCGGCAGC
CATATGACACCGGTTACCTTTATTGGTCTGGGTCCGATGGGTCAAGCAATGGTTCGTAG
CCTGCTGGCAGGCGGTCATCCGGTTACCGTTTGGAATCGTACCGCAAGCCGTGCAGAT
GGTGTTGTTGCCGATGGTGCAAAACGTGCAGATAGCGCAGCACTGGCAATTGCAGAAA
ATGAACTGGTTATTCTGAGCCTGACCGATTATCAGGCAATGTATGACATTCTGGGTCCT
GTTGGTGATGCACTGCGTGGTCGTGTTGTTGTTAATCTGAGCAGCGATACCCCGAGTC
GTACCCGTGAAGCAGCAGAATGGCTGGCAGAACGTGGTGCAGAACTGATTACCGGTG
GTGTTATGGTTCCGGCAGAATGGGTTGGTAAAGATGCAAGCTATGTGTTTTATAGCGGT
CCGGAAGCAGTTTTTGAACGTTTTCGTGATACCCTGGTTCTGATTGGTCGTACAGATTA
TCTGGGTGCAGATCATGCACTGGCACAGCTGTTTTATCAGGCCCAGCTGGATATTTTTC
TGACCGCACTGAGCGCATATCTGCATGCAAGCGCACTGCTGGAAGCACATGGTGTTAC
CGCAGAAAAATTCTGGCCGTATGCAGAAAGCAATCTGAATACCATTGCACCGATGATGG
TTGAAGCACCGGGTCAGCTGGATTCAGGTGAATATCCGGGTGATCAGGCAAATGCCAA
AATGATGGGTGCAACCGCAGATCATATTGTTCAGGCCAGCTATGAAGCAGGTATTGATA
CCGGTCTGCCTCGTGCAGTTAAAGATCTGTATGATCGTGCCATTGCTGCCGGTGATGG
TGGTAAAAGCTGGACCAGCCTGTTTGAACAGCTGCGTCGTCGTGGTTAA

**SuIRED amino acid sequence - reference for numbering**
MRAKVTVLGLGPMGAALASAFLAAGHPTTVWNRTPGRAGALIDRGAREESSAAAAVTAGP
LVVICLVSYDAVEEVLAPLSAQLRGRTIVNLTSGSPVQAREAAALAQRCGADYLDGVIMTTP
PGIGRPESLLLHGGAPEVFAAHRDTLAALGDPVHVGADPALSSVYDTALLSQMWGTLTGW
LHAVALIGSDGPGGGVTAREYTGIADRWMGSVSAFMNAYAPHVDSGRYPGGDFTLDLHLR
**T**MEVLSHASELRGVASGLPQVFEELTRRAVDAGHGDDSYARLVEFMRVGGHV*
*T241 equivalent in bold

**SuIRED amino acid sequence - expression construct**
MGSSHHHHHHSSGLVPRGSHMRAKVTVLGLGPMGAALASAFLAAGHPTTVWNRTPGRAG
ALIDRGAREESSAAAAVTAGPLVVICLVSYDAVEEVLAPLSAQLRGRTIVNLTSGSPVQARE
AAALAQRCGADYLDGVIMTTPPGIGRPESLLLHGGAPEVFAAHRDTLAALGDPVHVGADPA
LSSVYDTALLSQMWGTLTGWLHAVALIGSDGPGGGVTAREYTGIADRWMGSVSAFMNAY
APHVDSGRYPGGDFTLDLHLR**T**MEVLSHASELRGVASGLPQVFEELTRRAVDAGHGDDSY
ARLVEFMRVGGHV*
*T241 equivalent in bold

**SuIRED nucleotide sequence - expression construct**
ATGGGCAGCAGCCATCATCATCATCATCACAGCAGCGGCCTGGTGCCGCGCGGCAGC
CATATGCGTGCAAAAGTTACCGTTTTAGGTCTGGGTCCGATGGGTGCAGCACTGGCAA
GCGCATTTCTGGCAGCAGGTCATCCGACCACCGTTTGGAATCGTACACCGGGTCGTGC
CGGTGCACTGATTGATCGTGGTGCACGTGAAGAAAGCAGCGCAGCAGCAGCCGTTAC
CGCAGGTCCGCTGGTTGTTATTTGTCTGGTTAGCTATGATGCCGTTGAAGAAGTTCTGG
CACCGCTGAGCGCACAGCTGCGTGGTCGTACCATTGTTAATCTGACCAGCGGTAGTCC

GGTTCAGGCACGCGAAGCAGCCGCACTGGCCCAGCGTTGTGGTGCAGATTATCTGGA
TGGTGTTATTATGACCACACCGCCTGGTATTGGTCGTCCGGAAAGCCTGCTGTTACATG
GTGGTGCACCGGAAGTTTTTGCAGCACATCGTGATACCCTGGCAGCCCTGGGTGATCC
GGTTCATGTTGGTGCCGATCCGGCACTGAGCAGCGTTTATGATACCGCACTGCTGAGC
CAGATGTGGGGCACCCTGACCGGTTGGCTGCATGCCGTTGCGCTGATTGGTAGTGATG
GTCCTGGTGGTGGTGTTACCGCACGTGAATATACCGGCATTGCAGATCGTTGGATGGG
TAGCGTTAGCGCATTTATGAATGCCTATGCACCGCATGTTGATAGCGGTCGTTATCCAG
GTGGTGATTTTACCCTGGATCTGCATCTGCGTACCATGGAAGTTCTGAGCCATGCAAGC
GAACTGCGTGGTGTTGCAAGCGGTCTGCCGCAGGTTTTTGAAGAACTGACCCGTCGTG
CAGTTGATGCAGGTCATGGTGATGATAGCTATGCACGTCTGGTTGAATTTATGCGTGTT
GGTGGTCATGTCTAA
